# DCAF16-Based Covalent Degradative Handles for the Modular Design of Degraders

**DOI:** 10.1101/2025.04.25.650514

**Authors:** Lauren M. Orr, Sydney J. Tomlinson, Hannah R. Grupe, Melissa Lim, Emily Ho, Halime Yilmaz, Grace Zhou, Barbara Leon, James A. Olzmann, Daniel K. Nomura

## Abstract

Targeted protein degradation (TPD) is a powerful strategy for targeting and eliminating disease-causing proteins. While heterobifunctional Proteolysis-Targeting Chimeras (PROTACs) are more modular, the rational design of monovalent or molecular glue degraders remains challenging. In this study, we generated a small library of BET-domain inhibitor JQ1 analogs bearing elaborated electrophilic handles to identify permissive covalent degradative handles and E3 ligase pairs. We identified an elaborated fumaramide handle that, when appended onto JQ1, led to the proteasome-dependent degradation of BRD4. Further characterization revealed that the E3 ubiquitin ligase CUL4^DCAF16^—a common E3 ligase target of electrophilic degraders—was responsible for BRD4 loss by covalently targeting C173 on DCAF16. While this original fumaramide handle, when appended onto other protein-targeting ligands, did not accommodate the degradation of other neo-substrates, a truncated version of this handle attached to JQ1 was still capable of degrading BRD4, now through targeting both C173 and C178. This truncated fumaramide handle, when appended on various protein targeting ligands, and was also more permissive in degrading other neo-substrates, including CDK4/6, SMARCA2/4, and the androgen receptor (AR). We further demonstrated that this optimized truncated fumaramide handle, when transplanted onto an AR DNA binding domain-targeting ligand, could degrade both AR and the undruggable truncation variant of AR, AR-V7, in androgen-independent prostate cancer cells in a DCAF16-dependent manner. Overall, we have identified a unique DCAF16-targeting covalent degradative handle that can be transplanted across several protein-targeting ligands to induce the degradation of their respective targets for the modular design of monovalent or bifunctional degraders.

## Introduction

Targeted protein degradation (TPD) with heterobifunctional Proteolysis-Targeting Chimeras (PROTACs) or molecular glue degraders has emerged as a popular strategy for degrading and eliminating disease-causing proteins, including those deemed “undruggable.” Covalent chemistry and covalent chemoproteomics approaches have also enabled the identification of new covalent degradative handles and E3 ligase pairs beyond cereblon and VHL that can be exploited for TPD applications. Electrophilic handles have enabled the covalent recruitment of several components of the ubiquitin-proteasome system for both heterobifunctional and monovalent degraders, including E3 ligases CUL4^DCAF16^, CUL4^DCAF11^, CUL1^FBXO22^, CUL1^FBXW7(R465C)^, CUL4^DCAF1^, RNF114, RNF4, and RNF126, Cullin adaptor proteins DDB1 and SKP1, as well as E2 ubiquitin-conjugating enzymes UBE2D ^1–21^. In hunting for permissive electrophilic degradative handles, there has been a convergence in specific “frequent hitting” E3 ligases that appear to be particularly amenable for covalent recruitment in TPD applications, including CUL4^DCAF16^, CUL4^DCAF11^, and CUL1^FBXO22 1,2,4,5,9–11,22–28^ While several electrophilic handles have been identified that accommodate the degradation of individual neo-substrates, identifying permissive covalent degradative handles that can be broadly used against many different ligand and target pairs beyond recruiters that target cereblon or VHL ^29–31^, for both heterobifunctional and monovalent degraders has been challenging.

In this study, we generated a small library of monovalent BET-domain inhibitor JQ1 analogs bearing elaborated electrophilic handles to identify permissive covalent degradative handles and E3 ligase pairs. We identified an initial electrophilic handle that degraded BRD4 through the previously identified electrophile-sensitive E3 ligase CUL4^DCAF16^. However, this initial handle proved not to be particularly permissive to other ligand and target pairs, prompting further optimization of our initial handle to generate a truncated fumaramide handle that we demonstrated to be more accommodating in degrading neo-substrate proteins, still through CUL4^DCAF16^. We have thus identified a unique covalent degradative handle that exploits CUL4^DCAF16^, which is transplantable across multiple protein-targeting ligands to enable the degradation of their respective neo-substrate protein targets.

## Results

### Screening of a Covalent JQ1 Library for BRD4 Degraders

To identify unique electrophilic handles that can induce the degradation of neo-substrate proteins, we generated a small collection of 16 BET inhibitor JQ1 analogs bearing elaborated fumaramide or acrylamide-based electrophilic handles **(Figure 1a)**. We subsequently screened these compounds for BRD4 degradation in HEK293T cells. We identified five compounds—HRG034, HRG038, HRG073, HRG078, and HRG083—that led to significant reductions in the short isoform of BRD4. Notably, these five compounds bore elaborated fumaramide handles, while the remaining compounds (except HRG075) contained elaborated acrylamide warheads and were much less active. HRG038 showed the highest degree of degradation, with an 84 % loss of BRD4 **(Figure 1b-1d)**. Given that HRG038 also did not impair cell viability in HEK293T cells, we prioritized follow-up of this compound **(Figure S1a)**. We synthesized a negative control analog of HRG038 with the same covalent handle attached to the inactive JQ1 enantiomer, LO-3-60, demonstrating that this compound did not degrade BRD4 **(Figure S1b-S1c)**.

**Figure 1.**
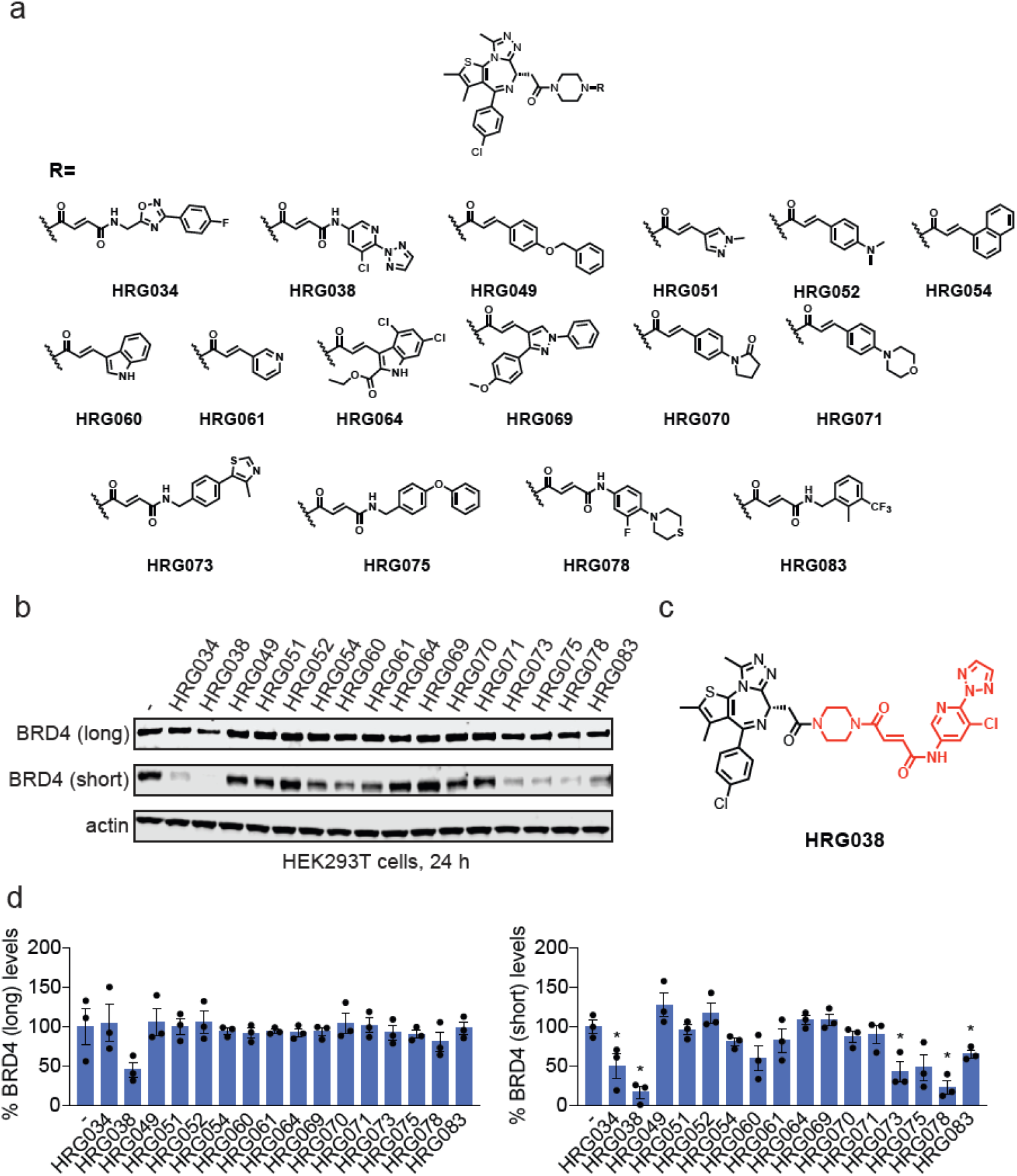
Screening of JQ1 analogs bearing electrophilic handles for BRD4 degradation. **(a)** Structures of JQ1 analogs bearing electrophilic handles. **(b)** Screening JQ1 electrophilic library in HEK293T cells for BRD4 degradation. HEK293T cells were treated with DMSO vehicle or compounds (1 μM) for 24 h and BRD4 and loading control actin levels were assessed by SDS/PAGE and Western blotting. **(c)** Structure of best hit HRG038 with the elaborated electrophilic handle in red. **(d)** Quantification of the experiment is described in **(b)**. Blot in **(b)** represents n=3 biologically independent replicates per group. Data in **(d)** shows individual replicates and average ± sem from n=3 biologically independent replicates per group. Significance expressed as *p<0.05 compared to vehicle-treated controls.

HRG038 showed selective loss of the short BRD4 isoform over the long BRD4 isoform in HEK293T cells with nanomolar potency **(Figure 2a)**. We have previously observed this preference for short BRD4 isoform degradation over the long isoform in HEK293T cells with covalent degraders, which was not evident in other cell lines ^5,19,32^. Similarly, we observed potent degradation of both long and short isoforms of BRD4 with HRG038 in MDA-MB-231 breast cancer cells **(Figure 2b)**. The HRG038-mediated degradation of BRD4 in HEK293T cells was attenuated upon pre-treatment of cells with inhibitors of the proteasome or NEDDylation with bortezomib (BTZ) and MLN4924, respectively, demonstrating proteasome- and Cullin E3 ligase-dependence of BRD4 loss **(Figure 2c-2d)**. Quantitative proteomic profiling of HRG038 in MDA-MB-231 cells demonstrated moderately selective degradation of BRD4 with 89 other proteins significantly downregulated **(Figure 2e, Table S1)**.

**Figure 2.**
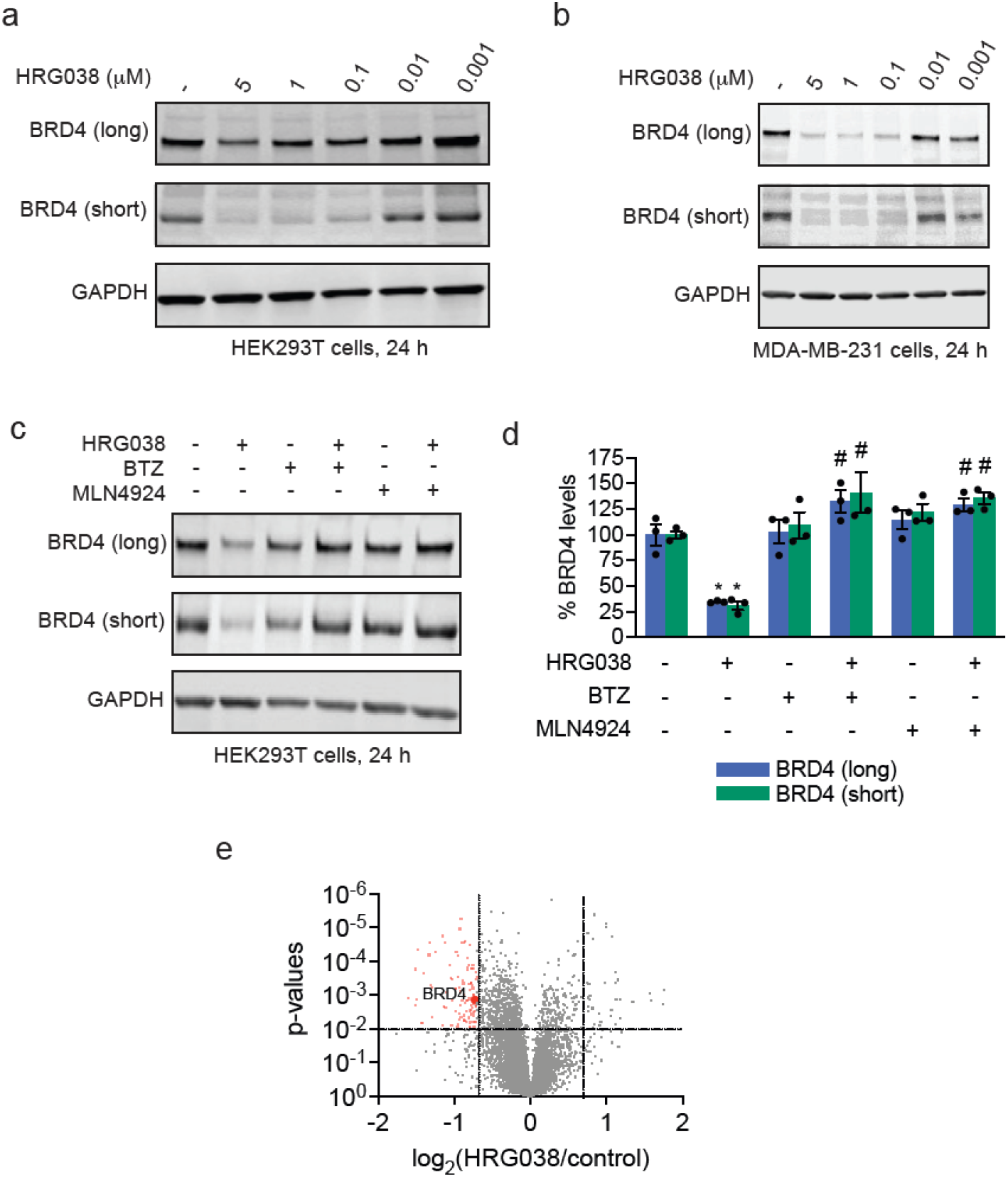
Characterization of HRG038 as a BRD4 degrader. **(a, b)** Dose-response of BRD4 degradation with HRG038 treatment. HEK293T cells **(a)** or MDA-MB-231 cells **(b)** were treated with DMSO vehicle or HRG038 for 24 h, after which BRD4 and loading control GAPDH levels were assessed by SDS/PAGE and Western blotting. **(c)** Proteasome-dependence of BRD4 degradation by HRG038. HEK293T cells were pre-treated with DMSO vehicle, BTZ (1 μM), or MLN4924 (1 μM) for 1 h prior to treatment of cells with DMSO or HRG038 (100 nM) for 24 h after which BRD4 and loading control GAPDH levels were assessed by SDS/PAGE and Western blotting. **(d)** Quantification of data from the experiment described in **(c). (e)** Quantitative tandem mass tagging (TMT)-based proteomic profiling of HRG038 in MDA-MB-231 cells. MDA-MB-231 cells were treated with DMSO vehicle or HRG038 (1 μM) for 24 h. Shown in red are proteins significantly lowered in levels with log_2_ < 0.6 with p < 0.01 with BRD4 highlighted. Full proteomics data can be found in **Table S1**. Blots in **(a-c)** are representative of n=3 biologically independent replicates per group. Bar graph in **(d)** shows individual replicate values and average ± sem from n=3 biologically independent replicates per group. Data in **(e)** are from n=3 biologically independent replicates per group. Significance in **(d)** shown as *p<0.01 compared to vehicle-treated controls and #p<0.05 compared to HRG038-treated groups.

### Functional Genomic Screening of a Ubiquitin Ligase Library

To identify the E3 ligase responsible for HRG038-mediated BRD4 degradation, we performed parallel functional CRISPR-Cas9 screens with an sgRNA library targeting ∼2000 genes associated with ubiquitin, autophagy, and lysosomal (UBAL) degradation pathways (10 sgRNAs per gene) in HEK293T cells stably expressing BRD4-GFP and Cas9 **(Figure 3a)**. Fluorescent reporter cells with high and low amounts of BRD4-GFP were subsequently isolated by fluorescence-activated cell sorting and deep sequencing performed to identify either endogenous factors (vehicle-treated condition) or degrader-specific factors (HRG038-treated condition) involved in BRD4 degradation. Through this analysis, we identified SPOP as an E3 ligase associated with the native turnover of BRD4 **(Figure 3b)**, which is consistent with previous reports regarding the degradation pathway for BRD4 ^33^ and the overall efficacy of our screening approach to reveal essential degradation factors. CUL4^DCAF16^ and the CUL4 adaptor protein DDB1 were identified as HRG038-specific targets responsible for BRD4-GFP degradation **(Figure 3c-3d)**. Multiple sgRNAs targeting SPOP, DCAF16, and DDB1 were significantly enriched in the high GFP cells, confirming these genes as high-confidence candidate regulators of BRD4 stability **(Figure 3e)**. Thus, our data indicated that the E3 ligase CUL4^DCAF16^ was responsible for HRG038-mediated BRD4 degradation.

**Figure 3.**
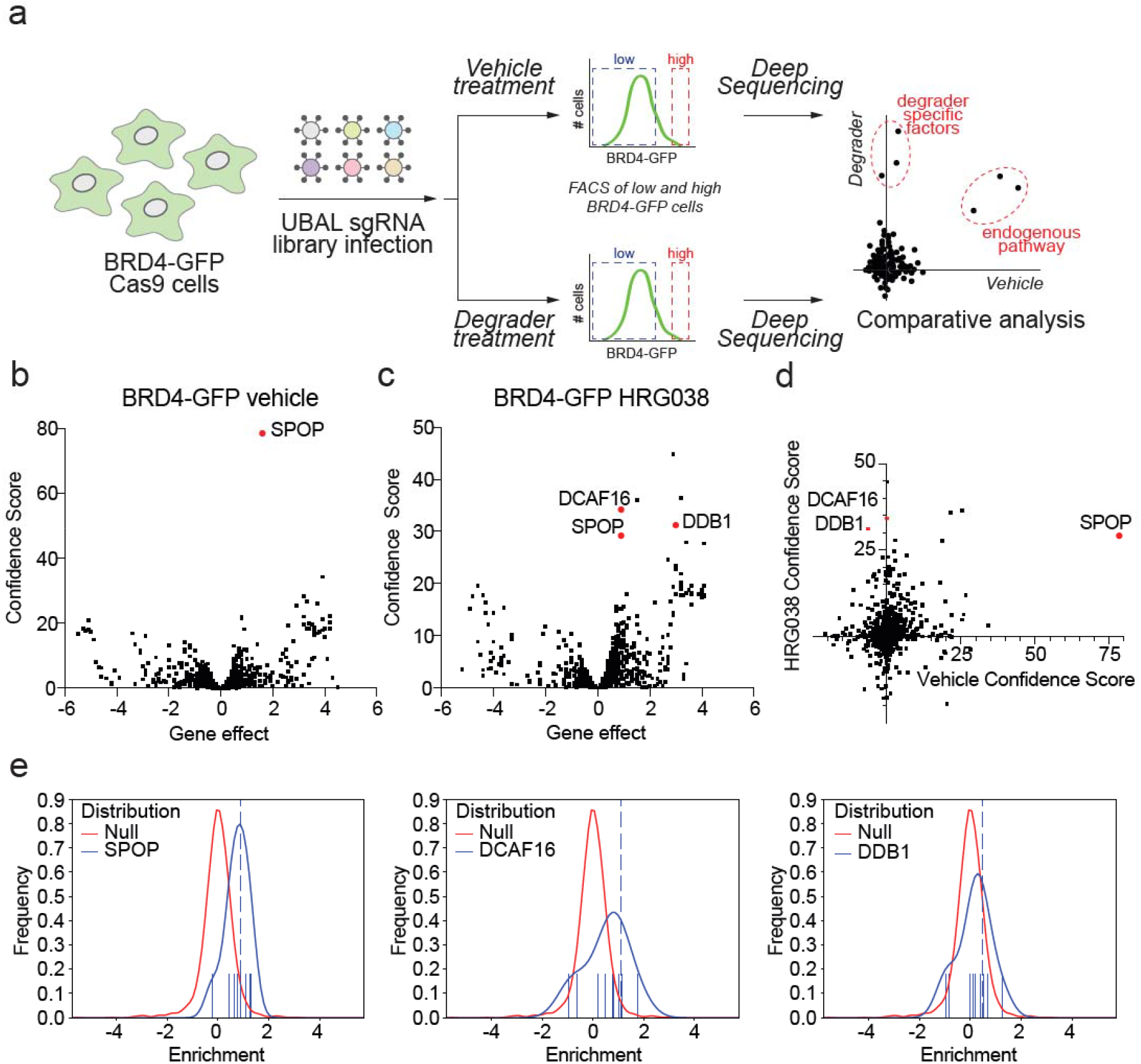
Functional CRISPR screen with UBAL library to identify E3 ligase responsible for HRG038-mediated BRD4 degradation. **(a)** A schematic for the CRISPR-Cas9 screen design. We performed parallel screens with the BRD4-GFP reporter cells expressing Cas9 treated with vehicle or HRG038 (in duplicate). **(b**,**c)** Volcano plots of the screen data for the vehicle **(b)** and HRG038 **(c)**. The y-axis is a confidence score, and the x-axis indicates the maximum effect (phenotype) size, with more positive values indicating an enrichment of the sgRNAs in the high GFP population, while more negative values reflecting an enrichment in the low GFP population. The confidence score is based on the Cas9 High-Throughput Maximum Likelihood Estimator (casTLE) score ^43^. **(d)** Comparative analyses of the confidence scores in **(b)** and **(c)**. This analysis highlights hits that are part of the endogenous pathway (e.g., SPOP) versus hits that are selective to the degrader molecule (e.g., hits that sit on the y-axis, DCAF16 / DDB1). **(e)** Histograms showing the enrichment of multiple sgRNAs per gene within cells with high levels of BRD4-GFP (i.e., degradation inhibited by gene KO). Functional CRISPR screen data can be found in **Table S2**.

### Validation of DCAF16 as E3 Ligase Responsible for HRG038-Mediated BRD4 Degradation

To determine whether our handle interacts directly with CUL4^DCAF16^, we synthesized an alkyne-functionalized probe, LO-3-44 bearing the fumaramide handle from HRG038 **(Figure 4a)**. We demonstrated dose-responsive and covalent labeling of pure human CUL4^DCAF16^ in the DCAF16-DDB1-DDA1 complex, with no labeling observed on DDB1 **(Figure 4b; Figure S1d)**. Confirming our results from the functional CRISPR screen, we demonstrated that HRG038-mediated degradation of BRD4 was fully attenuated in CUL4^DCAF16^ knockout (KO) HEK293T cells **(Figure 4c-4d)**. While we do not possess a DCAF16 antibody, we previously showed DCAF16 loss in this KO cell line by quantitative proteomics ^5^. Previous studies with CUL4^DCAF16^-mediated covalent PROTACs or molecular glue degraders had identified several distinct cysteines in CUL4^DCAF16^ that could contribute to neo-substrate degradation, including C177 and C179 with KB02-SLF, C58 with TMX1, and C119 with ML1-50 ^5,23,28,34^. To assess which cysteine in CUL4^DCAF16^ may be responsible for the effects of HRG038, we first mapped the site of reactivity of HRG038 with the recombinant DCAF16-DDA1-DDB1 complex by mass spectrometry detection of the compound adducted tryptic digest. The primary site of modification on CUL4^DCAF16^ was C173, but not C177, C178, or C179 **(Figure S1e)**. Further corroborating the importance of this residue, mutation of C173 to serine (C173S) in CUL4^DCAF16^, but not C58S, C119S, or C178S mutations, completely attenuated HRG038-mediated degradation of BRD4 **(Figure 4e-4f)**. Given that the previously published KB02-SLF acted through CUL4^DCAF16^ C177 and C179, we also tried to generate cells expressing C177S and C179S mutants, but in our hands we were not able to generate cells that could express these proteins, potentially because C177 and C179 coordinate zinc and may be necessary for the structural stability of CUL4^DCAF16 34^.

**Figure 4.**
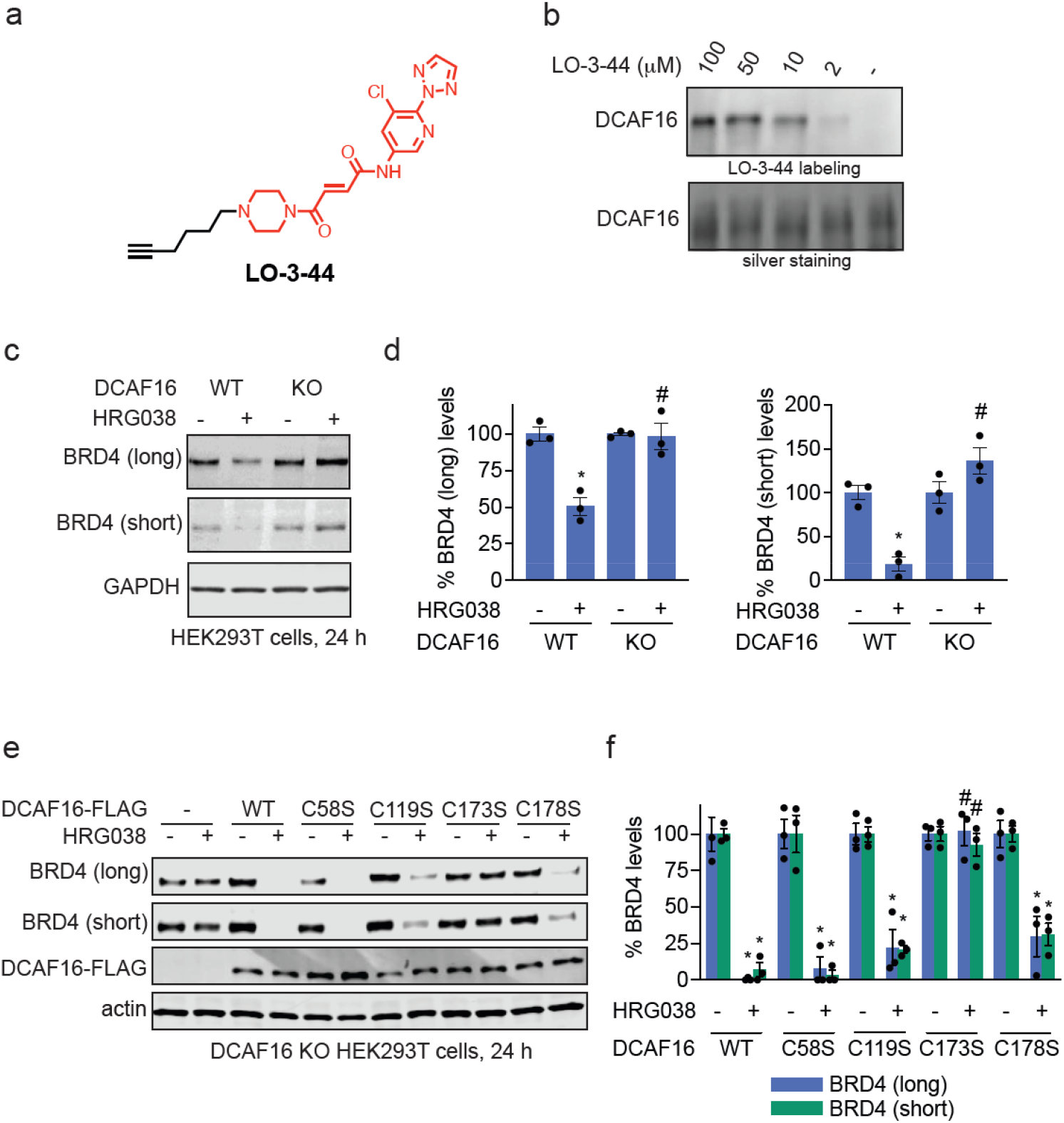
Characterizing CUL4^DCAF16^ as the responsible target for HRG038-mediated BRD4 degradation. **(a)** Alkyne-functionalized probe of degradative handle, LO-3-44. **(b)** LO-3-44 labeling of pure human CUL4^DCAF16^. The CUL4^DCAF16^-DDB1-DDA1 complex was labeled with LO-3-44 for 1h, after which probe-labeled proteins were conjugated with an azide-functionalized rhodamine by copper-mediated azide-alkyne cycloaddition (CuAAC). Proteins were separated by SDS/PAGE and visualized by in-gel fluorescence or loading was assessed by silver staining. **(c)** BRD4 degradation in CUL4^DCAF16^ WT and KO cells. CUL4^DCAF16^ WT and KO HEK293T cells were treated with DMSO vehicle or HRG038 (100 nM) for 24 h and BRD4 and loading control GAPDH levels were assessed by SDS/PAGE and Western blotting. **(d)** Quantification for the experiment described in **(c). (e)** HRG038-mediated BRD4 degradation in cells expressing WT and mutant CUL4^DCAF16^. CUL4^DCAF16^ KO HEK293T cells expressing either empty vector, FLAG-CUL4^DCAF16^ WT, C58S, C119S, C173S, or C178S were treated with DMSO vehicle or HRG038 (100 nM) for 24 h and BRD4 and loading control actin levels were assessed by SDS/PAGE and Western blotting. **(f)** Quantification for the experiment described in **(d)**. Gels and blots in **(b**,**c**,**e)** are representative of n=3 biologically independent replicates/group, of which individual replicates and average ± sem are shown in **(d**,**f)**. Significance is expressed as *p<0.05 compared to vehicle-treated WT controls in **(d)** or respective vehicle-treated groups **(f)** and #p<0.05 compared to CUL4^DCAF16^ WT cells treated with HRG038 in **(d)** or CUL4^DCAF16^ KO cells expressing FLAG-CUL4^DCAF16^ WT treated with HRG038 **(f)**.

### Transplanting the Fumaramide Handle onto Other Protein-Targeting Ligands

To assess whether this elaborated fumaramide handle was permissive towards the degradation of other neo-substrate proteins, we next generated compounds linking this handle onto several different types of protein-targeting ligands, including the CDK4/6 inhibitor ribociclib^35^, the SMARCA2/4 bromodomain inhibitor SGC SMARCA-BRDVIII^36,37^, and an androgen receptor (AR) antagonist from the AR PROTAC ARV-110^38^ to generate LO-3-20, LO-3-25, and LO-3-48, respectively **(Figure S2a-S2f)**. While LO-3-20 showed modest degradation of CDK4 and CDK6, LO-3-25, and LO-3-48 did not degrade their respective targets SMARCA2 or AR, respectively **(Figure S2a-S2f)**. These data indicated that this elaborated fumaramide handle was not permissive to other neo-substrates.

### Optimizing DCAF16 Handle to Accommodate Degradation of BRD4 and Other Neo-Substrates

Given that CUL4^DCAF16^ appears to have so many cysteines that can be accessed for targeted protein degradation of neo-substrates and that C173 is spatially located near C58 and the cysteine cluster C177-C179^28^, we postulated that our original handle might be restricted from accessing cysteines near C173 due to steric hindrance, thus limiting the ability of this handle to accommodate an expanded neo-substrate scope. We truncated our original fumaramide handle on HRG038 to address this possibility by removing the triazole moiety from the chloropyridine substituent to generate LO-3-61 **(Figure 5a)**. Gratifyingly, LO-3-61 treatment led to the proteasome-mediated degradation of BRD4 **(Figure 5b)**. Quantitative proteomic profiling showed selectivity for BRD4 degradation in K562 cells **(Figure 5c; Table S3)**. LO-3-61, like with HRG038, did not show impairments in cell viability **(Figure S3a)**. As was observed with HRG038, the loss of BRD4 observed with LO-3-61 was wholly attenuated in DCAF16 KO cells, thus still showing CUL4^DCAF16^-dependence of BRD4 degradation **(Figure 5d-5e)**.

**Figure 5.**
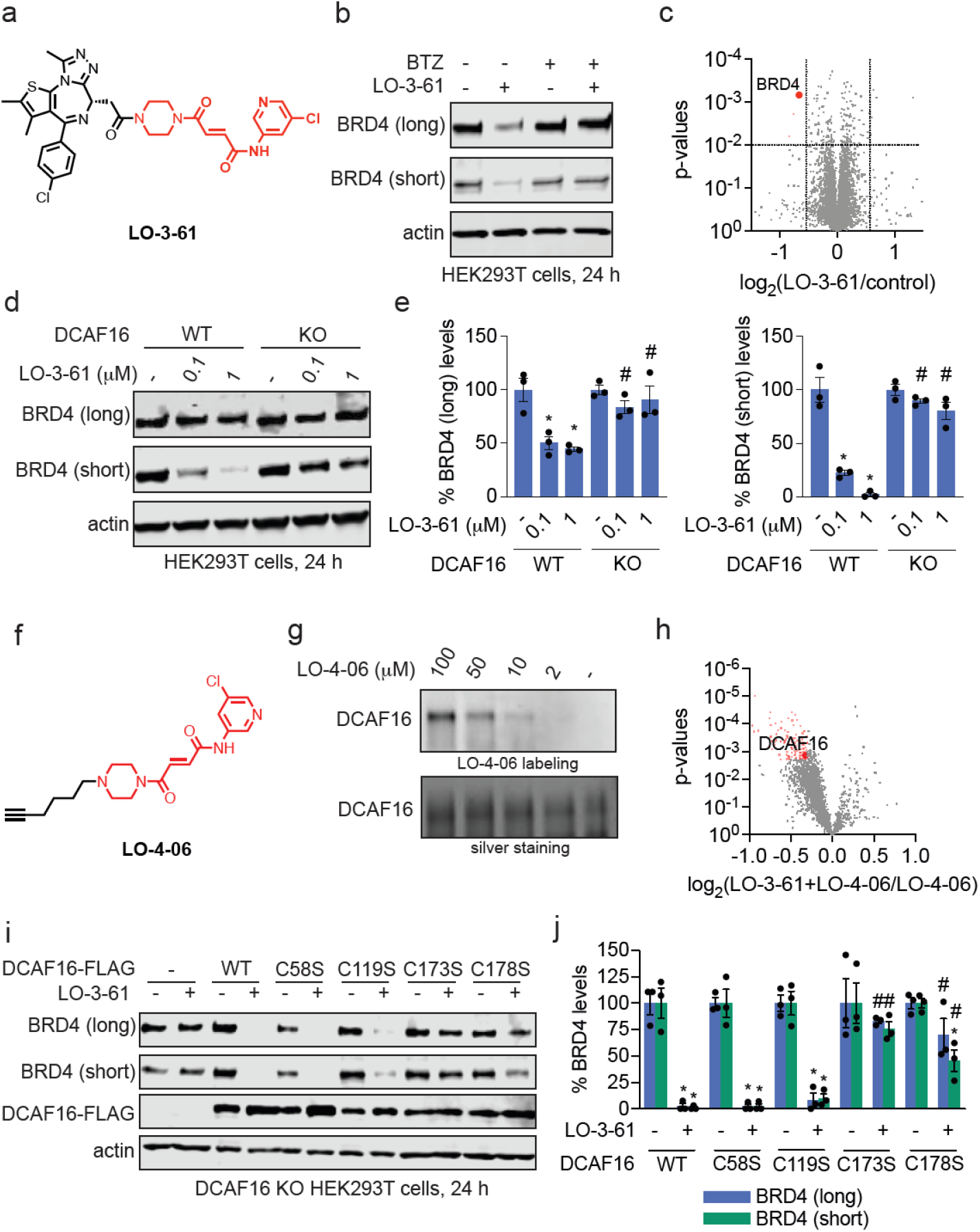
Assessing the degradative potential of truncated fumaramide handle. **(a)** Structure of LO-3-61, a JQ1 analog bearing a truncated fumaramide handle in red. **(b)** Proteasome-dependence of BRD4 degradation by HRG038. HEK293T cells were pre-treated with DMSO vehicle, BTZ (1 μM) for 1 h prior to treatment of cells with DMSO or LO-3-61 (100 nM) for 24 h after which BRD4 and loading control GAPDH levels were assessed by SDS/PAGE and Western blotting. **(c)** Quantitative tandem mass tagging (TMT)-based proteomic profiling of LO-3-61 in K562 cells. K562 cells were treated with DMSO vehicle or LO-3-61 (100 nM) for 24 h. Shown in red are proteins significantly lowered in levels with log_2_ < 0.6 with p < 0.01 with BRD4 highlighted. Full proteomics data can be found in **Table S3. (d)** BRD4 degradation in CUL4^DCAF16^ WT and KO cells. CUL4^DCAF16^ WT and KO HEK293T cells were treated with DMSO vehicle or LO-3-61 for 24 h, and BRD4 and loading control actin levels were assessed by SDS/PAGE and Western blotting. **(e)** Quantification of the experiment described in **(d). (f)** The structure of LO-4-06, an alkyne-functionalized probe based on the truncated fumaramide handle shown in red. **(g)** LO-3-44 labeling of pure human CUL4^DCAF16^. The CUL4^DCAF16^-DDB1-DDA1 complex was labeled with LO-3-44 for 1h, after which probe-labeled proteins were conjugated with an azide-functionalized rhodamine by copper-mediated azide-alkyne cycloaddition (CuAAC). Proteins were separated by SDS/PAGE and visualized by in-gel fluorescence or loading was assessed by silver staining. **(h)** Pulldown chemoproteomics profiling with the probe. HEK293T cell lysates were pre-treated with DMSO vehicle or LO-3-61 (50 µM) 1 h prior to treatment with LO-4-06 probe (10 µM). Probe-modified proteins were appended with azide-functionalized biotin by copper-mediated azide-alkyne cycloaddition (CuAAC), after which probe-modified proteins were avidin-enriched, tryptically digested, and quantitatively analyzed by TMT-based proteomics. Shown in red are proteins in which LO-4-06 labeling was significantly out-competed by LO-3-61, with DCAF16 highlighted. **(i)** LO-3-61-mediated BRD4 degradation in cells expressing WT and mutant CUL4^DCAF16^. CUL4^DCAF16^ KO HEK293T cells expressing either empty vector, FLAG-CUL4^DCAF16^ WT, C58S, C119S, C173S, or C178S were treated with DMSO vehicle or LO-3-61 (100 nM) for 24 h and BRD4 and loading control actin levels were assessed by SDS/PAGE and Western blotting. **(j)** Quantification for the experiment described in **(i)**. Gels and blots in **(b**,**d**,**g**,**i)** are representative of n=3 biologically independent replicates per group with individual replicates and average ± sem are shown in **(e**,**j)**. Data in **(c**,**h)** are from n=3 biologically independent replicates per group and proteomics data can be found in **Table S3** and **Table S4**, respectively.

To further confirm CUL4^DCAF16^ engagement, we generated an alkyne-functionalized probe based on the truncated fumaramide handle, LO-4-06 **(Figure 5f)**. As expected, we observed dose-responsive LO-4-06 covalent labeling of pure human CUL4^DCAF16^ protein in the DCAF16-DDB1-DDA1 complex, with no observed labeling of DDB1 **(Figure 5g; Figure S3b)**. Using this probe, we performed pulldown chemoproteomics profiling to identify proteins enriched by LO-4-06 treatment and outcompeted by LO-3-61 pre-treatment. This chemoproteomics profiling revealed significant enrichment of DCAF16 by the probe and competition of this enrichment with LO-3-61, demonstrating moderately selective targeting of CUL4^DCAF16^ with the handle and the BRD4 degrader LO-3-61 **(Figure 5h; Table S4)**. Site mapping of LO-3-61 labeling on the recombinant DCAF16-DDA1-DDB1 complex showed labeling on C173 **(Figure S3c)**. However, unlike HRG038, we observed significant rescue of BRD4 degradation in cells expressing CUL4^DCAF16^ C173S and cells expressing the C178S mutant **(Figure 5i-5j)**. We still observed equivalent BRD4 degradation in cells expressing CUL4^DCAF16^ C58S and C119S mutant as in CUL4^DCAF16^ wild-type expressing cells **(Figure 5i-5j)**. These data indicated that the reduced size of LO-3-61 may facilitate more flexibility in binding conformations in the CUL4^DCAF16^ substrate binding site, compared to HRG038, to enable access to not only C173 but also C178 for the degradation of BRD4.

The structure of CUL4^DCAF16^-DDB1 has been previously solved in a ternary complex with BRD4 and another electrophilic BRD4 molecular glue degrader, MMH2, that acts through C58 on CUL4^DCAF16 34^. This structure did not include the disordered loop that includes C173. Given that previous CUL4^DCAF16^-mediated electrophilic degraders were shown to act through C58 or C177/C179 ^23,34^, we wanted to understand the relation of C173 to C58 and C178. We modeled this disordered loop onto the previously reported structure. While a predicted model may not represent the actual conformation of this loop, the model showed relatively close proximity of C173, C58, and C178 within the binding region CUL4^DCAF16^ to BRD4 **(Figure 6; Figure S4)**. C177 and C179 are both coordinated to zinc, while C173, C178, and C58 are solvent-exposed, where C173 is within a disordered and likely flexible loop **(Figure 6; Figure S4)**. While we do not yet understand the structural basis for how HRG038 and LO-3-61 interact with C173 or C173 and C178, respectively, our model, consistent with previous structural work and commentary by Zhang and Cravatt ^28,34^, demonstrated multiple cysteines within the CUL4^DCAF16^ neo-substrate binding interface.

**Figure 6.**
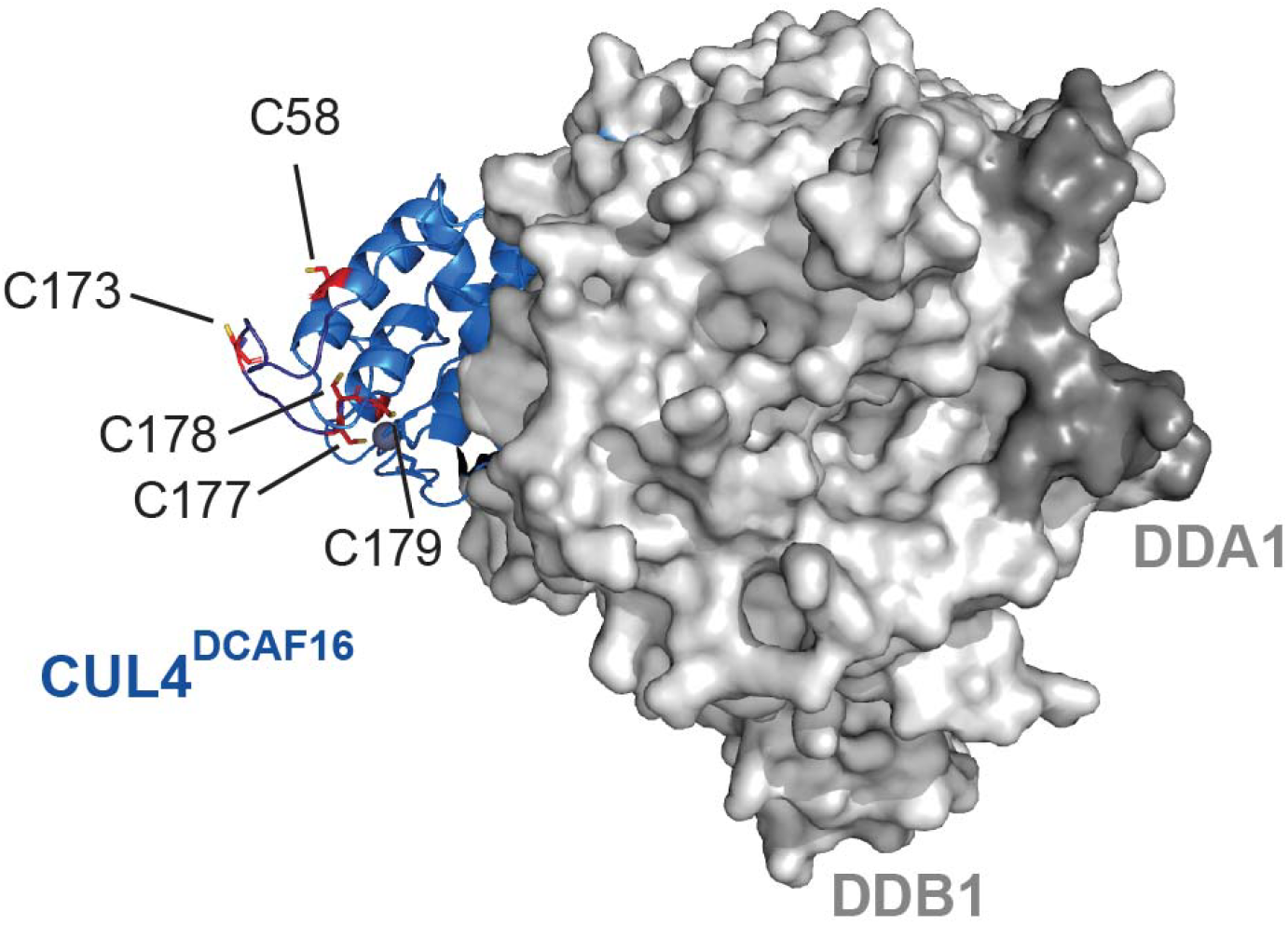
Model of CUL4^DCAF16^-DDB1-DDA1 complex showing C173, C177-C179, and C58. The model is based on a structure previously solved of DCAF16-DDB1-DDA1 in complex with BRD4 in the presence of an electrophilic BRD4 molecular glue degrader MMH2 (PDB: 8g46)^34^ that did not include the disordered loop containing C173. We have removed MMH2 and BRD4. To add the loop containing C173 not resolved in the reported structure, the PyMol builder function was used to add the residues to the model structure, and the loop was then fit to a potential conformation via homology modeling with ModLoop ^44,45^.

Encouraged by the finding that LO-3-61 can act through multiple cysteines in DCAF16, we surmised that the truncated handle could accommodate the formation of more diverse ternary complexes with DCAF16 to degrade a broader scope of neo-substrates when appended onto various protein-targeting ligands. When conjugated onto the CDK4/6 inhibitor ribociclib, LO-3-63 treatment exhibited much more robust degradation of CDK4 and CDK6 in HEK293T cells, compared to LO-3-20 after 24 h of treatment **(Figure 7a-7b; Figure S2a-S2b)**. This degradation of CDK4 and CDK6 was attenuated in CUL4^DCAF16^ KO cells, thus still showing CUL4^DCAF16^-dependence **(Figure 7c)**. While we did not observe degradation of SMARCA2 with LO-3-25 and AR with LO-3-48 **(Figure S2c-S2f)**, we observed degradation of SMARCA2/4 and AR with LO-3-62 and LO-4-12, respectively, with the truncated fumaramide handle **(Figure 7d-7e; Figure S5a-S5b)**. These data indicated that this truncated handle was more permissive in accommodating different ligand and target pairs.

**Figure 7.**
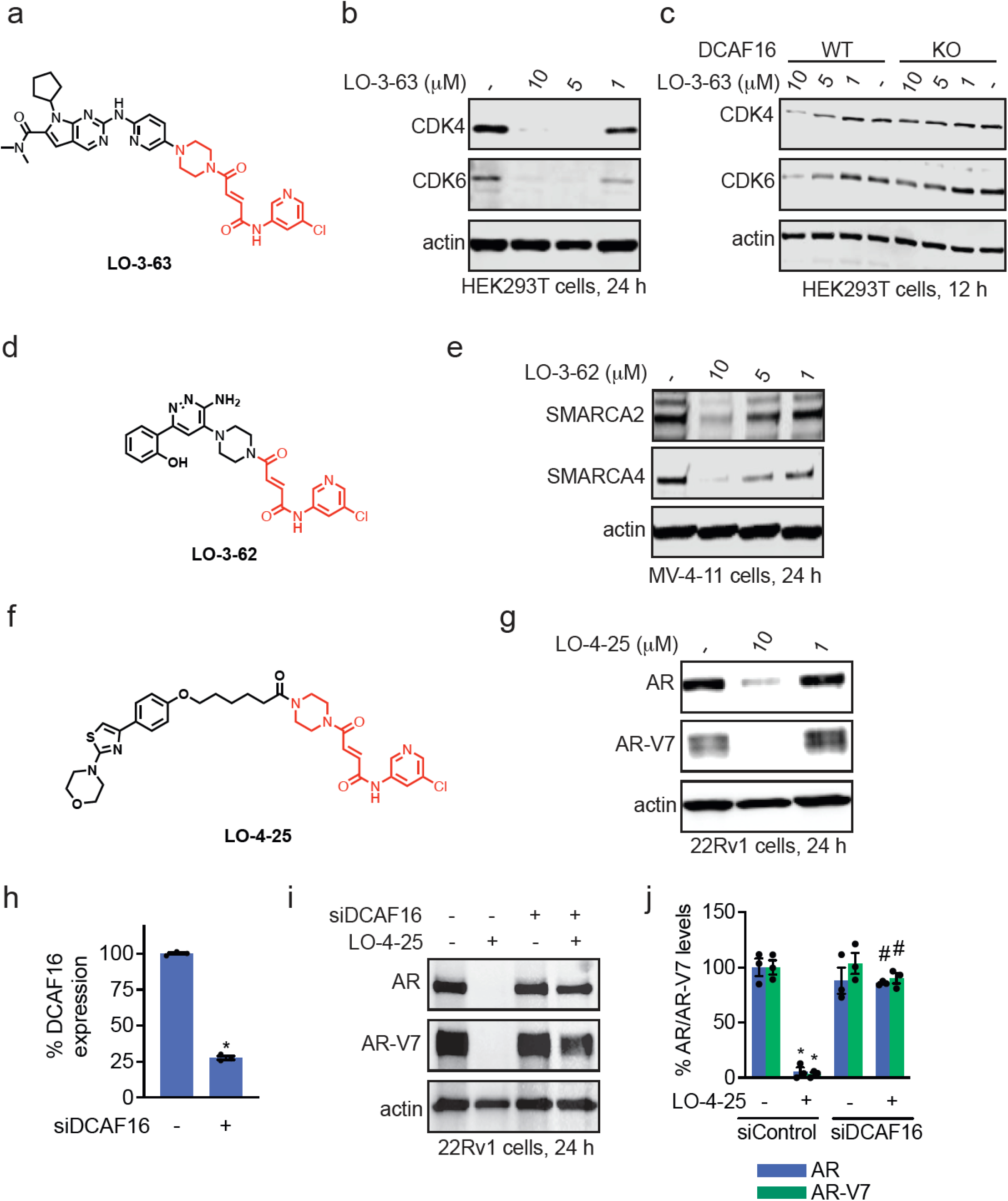
Permissiveness of truncated fumaramide handle in degrading multiple neo-substrates. **(a)** Structure of LO-3-63, a CDK4/6 inhibitor ribociclib appended to truncated fumaramide handle in red. **(b)** CDK4/6 degradation in HEK293T cells. HEK293T cells were treated with DMSO vehicle or LO-3-63 for 24 h after which CDK4/6 and loading control actin levels were assessed by SDS/PAGE and Western blotting. **(c)** CDK4/6 degradation CUL4^DCAF16^ WT and KO cells. CUL4^DCAF16^ WT and KO HEK293T cells were treated with DMSO vehicle or LO-3-63 for 12 h after which CDK4/6 and loading control actin levels were assessed by SDS/PAGE and Western blotting. **(d)** Structure of LO-3-62, a SMARCA2/4 inhibitor linked to truncated fumaramide handle in red. **(e)** SMARCA2/4 degradation in MV-4-11 cells. MV-4-11 cells were treated with DMSO vehicle or LO-3-62 for 24 h after which SMARCA2/4 and loading control actin levels were assessed by SDS/PAGE and Western blotting. **(f)** Structure of LO-4-25, a AR DNA binding domain ligand linked via a linker to the truncated fumaramide handle in red. **(g)** AR and AR-V7 degradation in 22Rv1 cells. 22Rv1 cells were treated with DMSO vehicle or LO-4-25 for 24 h after which AR and AR-V7 levels were assessed by SDS/PAGE and Western blotting. **(h)** DCAF16 mRNA levels. 22Rv1 cells were transiently transfected with siControl (−) or siDCAF16 (+) oligonucleotides for 48 h after which DCAF16 mRNA levels were assessed by qPCR. **(i)** AR and AR-V7 degradation in siControl and siDCAF16 22Rv1 cells. 22Rv1 siControl or siDCAF16 cells were treated with DMSO vehicle or LO-4-25 for 24 h and AR, AR-V7, and loading control actin levels were assessed by SDS/PAGE and Western blotting. **(j)** Quantification of the experiment described in **(i)**. Blots in **(b**,**c**,**e**,**g**,**i)** are representative of n=3 biologically independent replicates/group. Bar graphs in **(h**,**j)** show individual replicate values and average ± sem. Significance expressed as *p<0,05 compared to siControl in **(h)** or to vehicle-treated siControl groups in **(j)** and #p<0.05 compared to LO-4-25-treated siControl groups in **(j)**.

We next sought to test this truncated handle against a more challenging target, such as the truncation variant of the androgen receptor, AR-V7, that drives the pathogenesis of certain androgen-independent prostate cancers ^39,40^. AR-V7 has been particularly difficult to therapeutically target due to the lack of a steroid-binding domain to which most AR-targeting ligands bind, and the remaining portion of the protein is highly intrinsically disordered. Previous studies have identified a ligand that binds to the AR DNA binding domain ^41,42^. We previously identified a covalent RNF126-targeting degradative handle that, when appended onto this AR DNA binding domain ligand, led to the degradation of both AR and AR-V7 in 22Rv1 androgen-independent prostate cancer cells ^18^. We have also previously discovered a vinyl sulfonylpiperazine CUL4^DCAF16^ recruiting handle that acts through covalent targeting of C119 that showed versatile degradation of several neo-substrate targets, including BRD4, SMARCA4, CDK4, BTK, and BCR-ABL and c-ABL ^5^. However, we did not observe degradation of AR and AR-V7 when this handle was appended to the AR DNA binding domain ligand in 22Rv1 cells **(Figure S5c-S5d)**. Next, we generated a degrader linking the same AR DNA binding domain ligand to our new truncated fumaramide handle to yield LO-4-25 **(Figure 7f)**. LO-4-25 showed robust degradation of both AR and AR-V7 that was completely attenuated upon DCAF16 knockdown in 22Rv1 cells **(Figure 7f-7j)**. LO-4-25 exhibited very modest impairments in cell viability in 22Rv1 cells, possibly due to on- or off-target effects **(Figure S5e)**. Overall, we demonstrated that our truncated fumaramide handle that acts through targeting C173 and C178 on CUL4^DCAF16^ is more permissive in accommodating the degradation of multiple neo-substrate proteins when appended onto a diversity of protein-targeting ligands.

## Discussion

In this study, we report the identification and characterization of a versatile covalent degradative handle targeting the E3 ubiquitin ligase CUL4^DCAF16^, expanding the toolkit available for TPD. We screened a library of BET inhibitor JQ1 analogs with electrophilic handles and discovered an elaborated fumaramide derivative capable of potently degrading BRD4 via CUL4^DCAF16^. While initial iterations of this fumaramide handle displayed limited flexibility in targeting neo-substrates beyond BRD4, subsequent optimization yielded a truncated version with significantly improved permissiveness. This truncated fumaramide handle maintained robust degradation of BRD4 and expanded the scope of neo-substrate targets, including CDK4/6, SMARCA2/4, and notably, the androgen receptor (AR) and its therapeutically challenging truncation variant AR-V7 in androgen-independent prostate cancer cells.

While PROTAC and molecular glue degrader strategies commonly rely on cereblon and VHL, identifying alternative ligases, particularly covalently targetable ones, opens new avenues for expanding therapeutic reach, especially towards historically undruggable targets. Prior studies have established CUL4^DCAF16^ as a promising yet relatively underutilized ligase in covalent TPD approaches beyond BRD4 and FKBP12, primarily due to the challenge of designing handles with sufficient flexibility and permissiveness. Our optimized truncated fumaramide handle addresses this challenge, providing a generalizable and modular platform that can be transplanted across various protein-targeting ligands.

An intriguing aspect of our findings involves the differential cysteine targeting by the original and truncated fumaramide handles. Initial data indicated exclusive involvement of C173 in CUL4^DCAF16^ by the elaborated fumaramide handle. However, truncation enabled additional engagement with C178, broadening substrate permissivity and highlighting the importance of spatial and structural considerations in covalent degrader design. This structural flexibility appears crucial, given the diversity of accessible cysteines within DCAF16 previously described for other degraders ^2,5,23,34^. While we demonstrated that LO-3-61, bearing our truncated fumaramide handle, degraded BRD4 through both C173 and C178 on CUL4^DCAF16^, and not through C58, given the proximity of all three residues in our CUL4^DCAF16^ model, our other degraders may operate through these or other cysteines on CUL4^DCAF16^.

Despite the promising findings, several caveats and future considerations warrant mention. First, while the truncated fumaramide handle significantly broadens neo-substrate compatibility, its selectivity profile, as evidenced by proteomic data, requires further refinement to limit off-target effects. Additionally, while cell viability remained largely unaffected in the cell lines tested, broader evaluations across multiple cellular contexts are necessary to assess potential cytotoxicity or off-target liabilities *in vivo* fully. Further medicinal chemistry is also required to improve potency, overall selectivity, metabolic stability, and pharmacokinetic and pharmacodynamic parameters for any *in vivo* use.

Our work introduces an optimized, flexible fumaramide-based degradative handle exploiting the CUL4^DCAF16^ ligase for efficient, covalently targeted degradation. This handle’s modularity and broad applicability represent an advance in the rational design of monovalent and bifunctional degraders, paving the way toward therapeutic targeting of challenging proteins previously deemed undruggable.

## Acknowledgment

We thank the members of the Nomura Research Group and Novartis BioMedical Research for critically reading the manuscript. This work was also supported by Novartis Biomedical Research, the National Science Foundation Molecular Foundations for Biotechnology (MFB) grant (2127788), the UC Berkeley Molecular Therapeutics Initiative (MTI), the Mark Foundation for Cancer Research ASPIRE Award, the National Institutes of Health (R35CA263814, R01CA240981), the Bakar Award, and the Bakar Innovation Fellows. We also thank Hasan, Lund, and the UC Berkeley NMR facility in the College of Chemistry (CoC-NMR) for spectroscopic assistance. Instruments in the College of Chemistry NMR facility are partly supported by NIH S10OD024998.

