## Supporting Information for "DCAF16-Based Covalent Degradative Handles for the Modular Design of Degraders"

[1] Department of Chemistry. University of California, Berkeley, Berkeley, CA 94720 USA

[2] Department of Molecular and Cell Biology, University of California, Berkeley, Berkeley, CA 94720 USA

[3] Department of Nutritional Sciences and Toxicology, University of California, Berkeley, Berkeley, CA 94720 USA

[4] Innovative Genomics Institute, Berkeley, CA 94720 USA

[5] Novartis-Berkeley Translational Chemical Biology Institute, Berkeley, CA 94720 USA

[6] Novartis Biomedical Research, Emeryville, CA; Cambridge, MA; Basel, Switzerland

**Supporting Table Legends**

**Table S1. Quantitative TMT-based proteomic profiling of HRG038 in MDA-MB-231 cells.** MDA-MB-231 cells were treated with DMSO vehicle or HRG038 (1 μM) for 24 h. Data are from n=3 biologically independent replicates per group.

**Table S2. Functional CRISPR screen with UBAL library to identify E3 ligase responsible for HRG038-mediated BRD4 degradation.** We performed parallel screens with the BRD4-GFP cells expressing Cas9 treated with vehicle or HRG038 (in duplicate). Data show confidence scores and phenotype, with more positive numbers indicating an enrichment of the sgRNAs in the high GFP and negative numbers indicating a depletion in the sgRNAs in high GFP. The confidence score is based on the Cas9 High-Throughput Maximum Likelihood Estimator (casTLE) score.

**Table S3. Quantitative tandem mass tagging (TMT)-based proteomic profiling of LO-3-61 in K562 cells.** K562 cells were treated with DMSO vehicle or LO-3-61 (100 nM) for 24 h. Data are from n=3 biologically independent replicates per group.

**Table S4. Pulldown chemoproteomics profiling with the LO-4-06 probe.** HEK293T cell lysates were pre-treated with DMSO vehicle or LO-3-61 (50 μM) 1 h prior to treatment with LO-4-06 probe (10 μM). Probe-modified proteins were appended with azide-functionalized biotin by copper-mediated azide-alkyne cycloaddition (CuAAC), after which probe-modified proteins were avidin-enriched, tryptically digested, and quantitatively analyzed by TMT-based proteomics. Data are from n=3 biologically independent replicates per group.

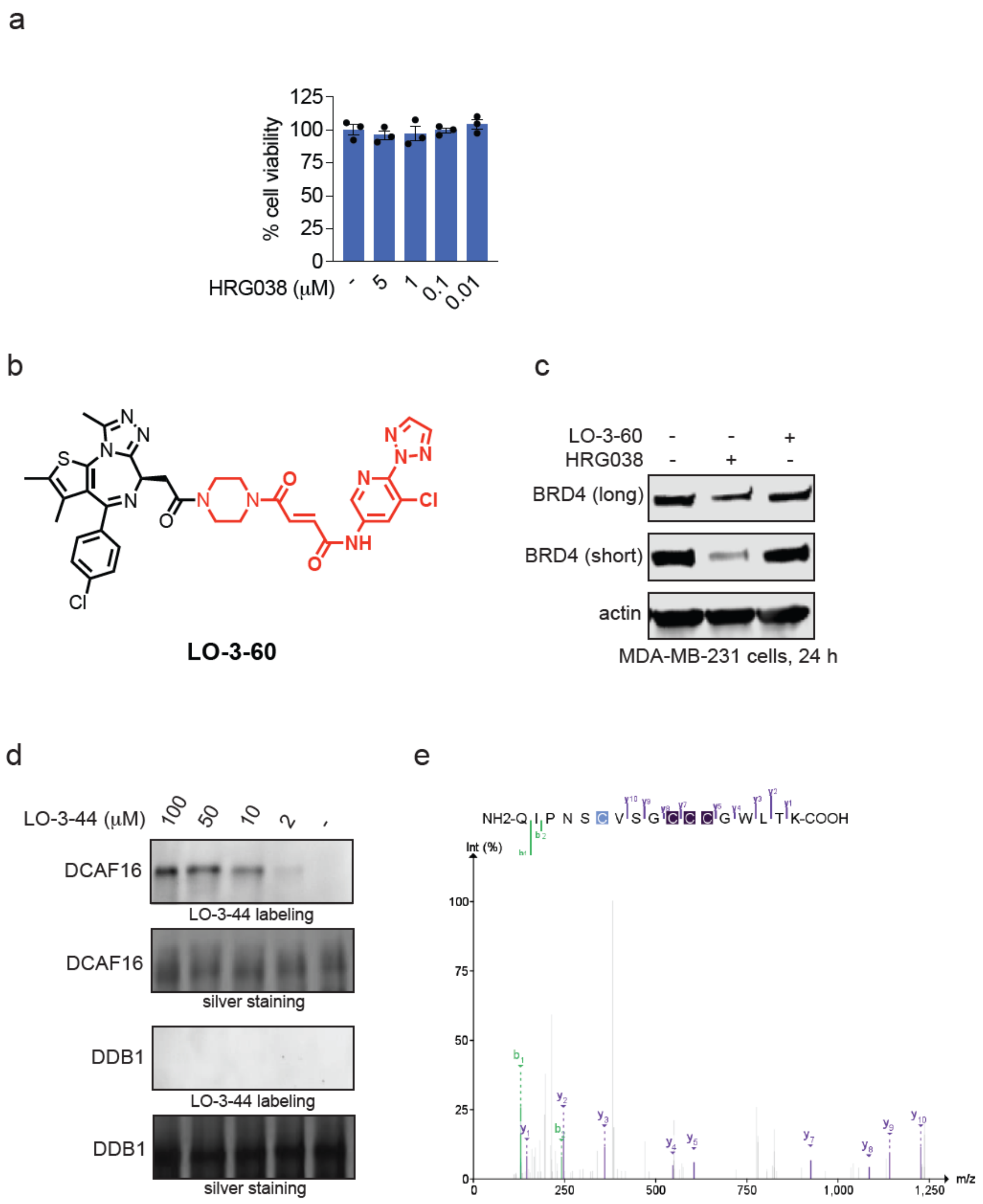


**Figure S1. Characterization of HRG038 and negative control enantiomer LO-3-60. (a)** HRG038 effects on cell viability. HEK293T cells were treated with DMSO vehicle or HRG038 for 24 h, and cell viability was assessed with CellTiter-Glo. **(b)** Structure of negative control enantiomer LO-3-60 with covalent fumaramide handle in red. **(c)** Degradation of BRD4 with HRG038 and LO-3-60. MDA-MB-231 cells were treated with DMSO vehicle, HRG038 (100 nM), or LO-3060 (100 nM) for 24 h, after which BRD4 and loading control actin levels were assessed by SDS/PAGE and Western blotting. **(d)** LO-3-44 labeling of pure human CUL4$^{DCAF16}$. The CUL4$^{DCAF16}$-DDB1-DDA1 complex was labeled with LO-3-44 for 1h, after which probe-labeled proteins were conjugated with an azide-functionalized rhodamine by copper-mediated azide-alkyne cycloaddition (CuAAC). Proteins were separated by SDS/PAGE, and labeling of DCAF16 and DDB1 was visualized by in-gel fluorescence, or loading was assessed by silver staining. **(e)** The human DCAF16-DDB1-DDA1 pure protein complex was treated with HRG038 (10 μM) for 1 h, after which the HRG038 site of modification on DCAF16 was assessed after tryptic digestion of the complex and analysis by LC-MS/MS. Shown are the spectra of C173 on DCAF16 modified by HRG038. Data in **(a,c,d)** are from n=3 biologically independent replicates per group. **(a)** shows individual replicate values and average ± sem. Blot and gel in **(c,d)** are representative.

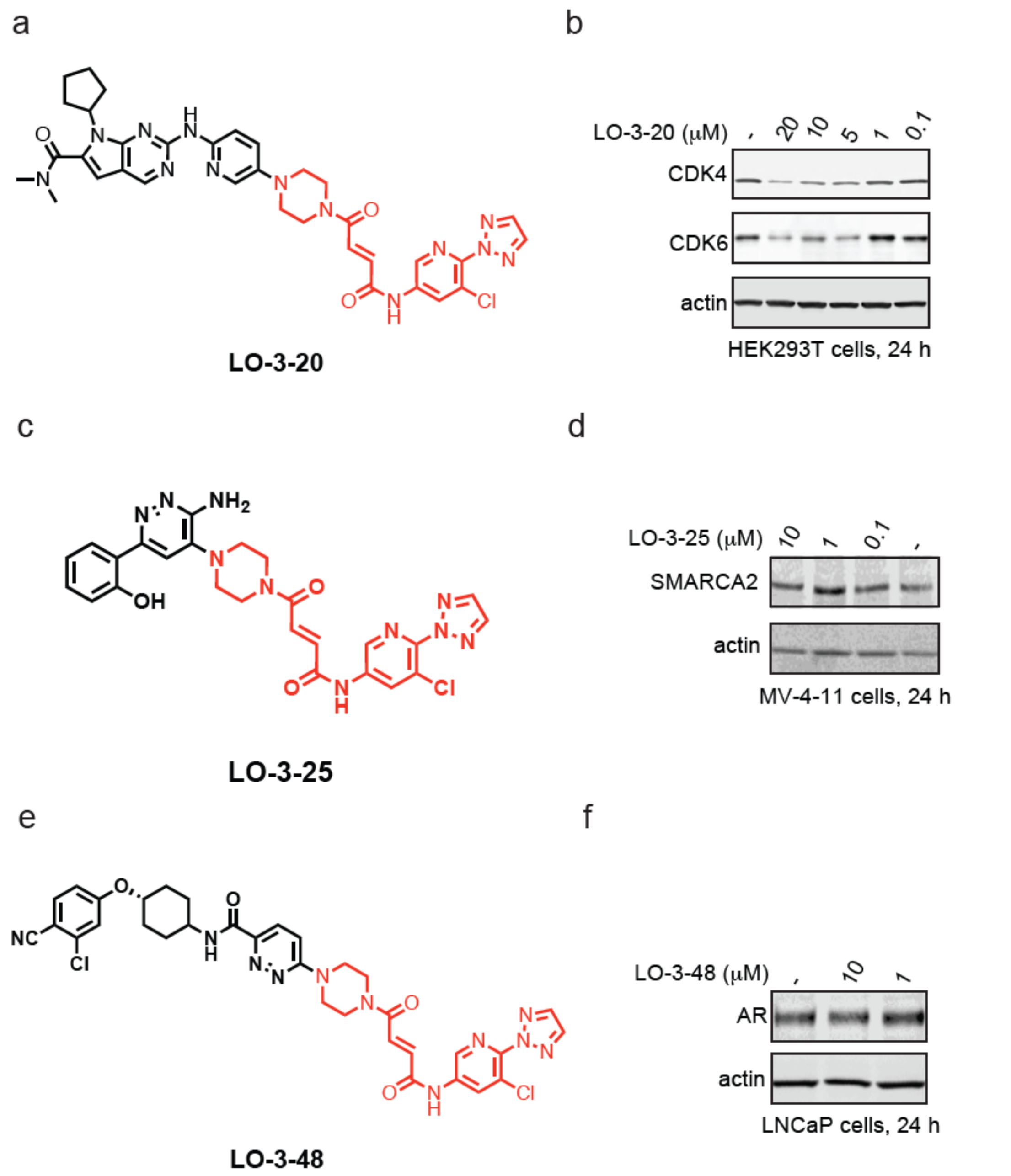


**Figure S2. Characterization of LO-3-20, LO-3-25, and LO-3-48. (a)** Structure, LO-3-20, ribociclib linked to the elaborated fumaramide handle in red. **(b)** CDK4/6 degradation by LO-3-20. HEK293T cells were treated with DMSO vehicle or LO-3-20 for 24 h after which CDK4/6 and loading control actin levels were assessed by SDS/PAGE and Western blotting. **(c)** Structure of LO-3-25, a SMARCA2/4 inhibitor linked to the elaborated fumaramide handle in red. **(d)** SMARCA2 degradation in MV-4-11 cells. MV-4-11 cells were treated with DMSO vehicle or LO-3-25 for 24 h and SMARCA2 and loading control actin levels were assessed by SDS/PAGE and Western blotting. **(e)** Structure of LO-3-48, an AR targeting ligand from the ARV110 PROTAC linked to the elaborated fumaramide handle in red. **(f)** AR degradation in LNCaP cells. LNCaP cells were treated with DMSO vehicle or LO-3-48 for 24 h and AR and loading control actin levels were assessed by SDS/PAGE and Western blotting. Blots in **(b,d,f)** are representative of n=3 biologically independent replicates per group.

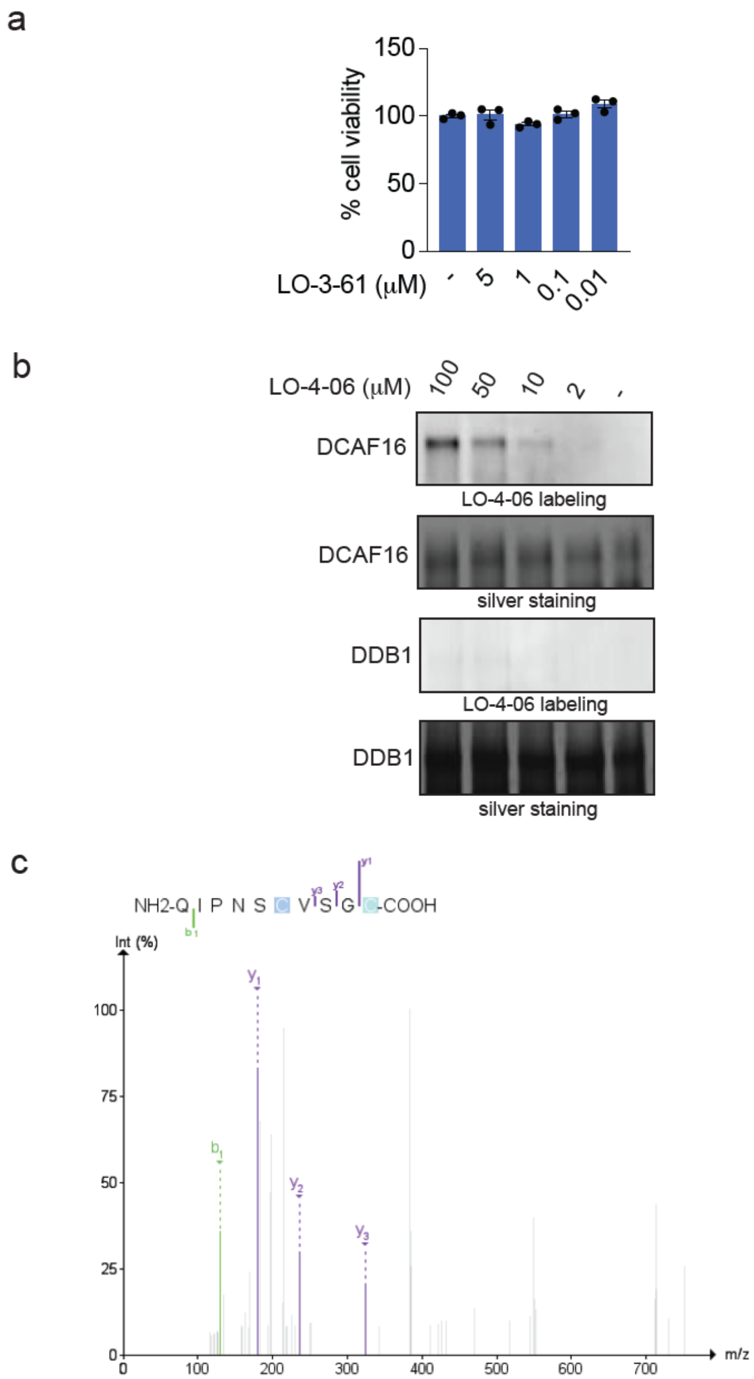


**Figure S3. Characterization of LO-3-61. (a)** HRG038 effects on cell viability. HEK293T cells were treated with DMSO vehicle or LO-3-61 for 24 h, and cell viability was assessed with CellTiter-Glo. **(b)** LO-3-44 labeling of pure human $CUL4^{DCAF16}$. The $CUL4^{DCAF16}$-DDB1-DDA1 complex was labeled with LO-3-44 for 1h, after which probe-labeled proteins were conjugated with an azide-functionalized rhodamine by copper-mediated azide-alkyne cycloaddition (CuAAC). Proteins were separated by SDS/PAGE, and DCAF16 or DDB1 labeling was visualized by in-gel fluorescence, and loading was assessed by silver staining. **(c)** The human DCAF16-DDB1-DDA1 pure protein complex was treated with LO-3-61 (50 μM) for 1 h, after which the LO-3-61 site of modification on DCAF16 was assessed after tryptic digestion of the complex and analysis by LC-MS/MS. The spectra of C173 on DCAF16 modified by LO-3-61 are shown. Data in **(a)** are from n=3 biologically independent replicates per group and show individual replicate values and average ± sem. Gel in **(b)** is representative of n=3 biologically independent replicates per group.

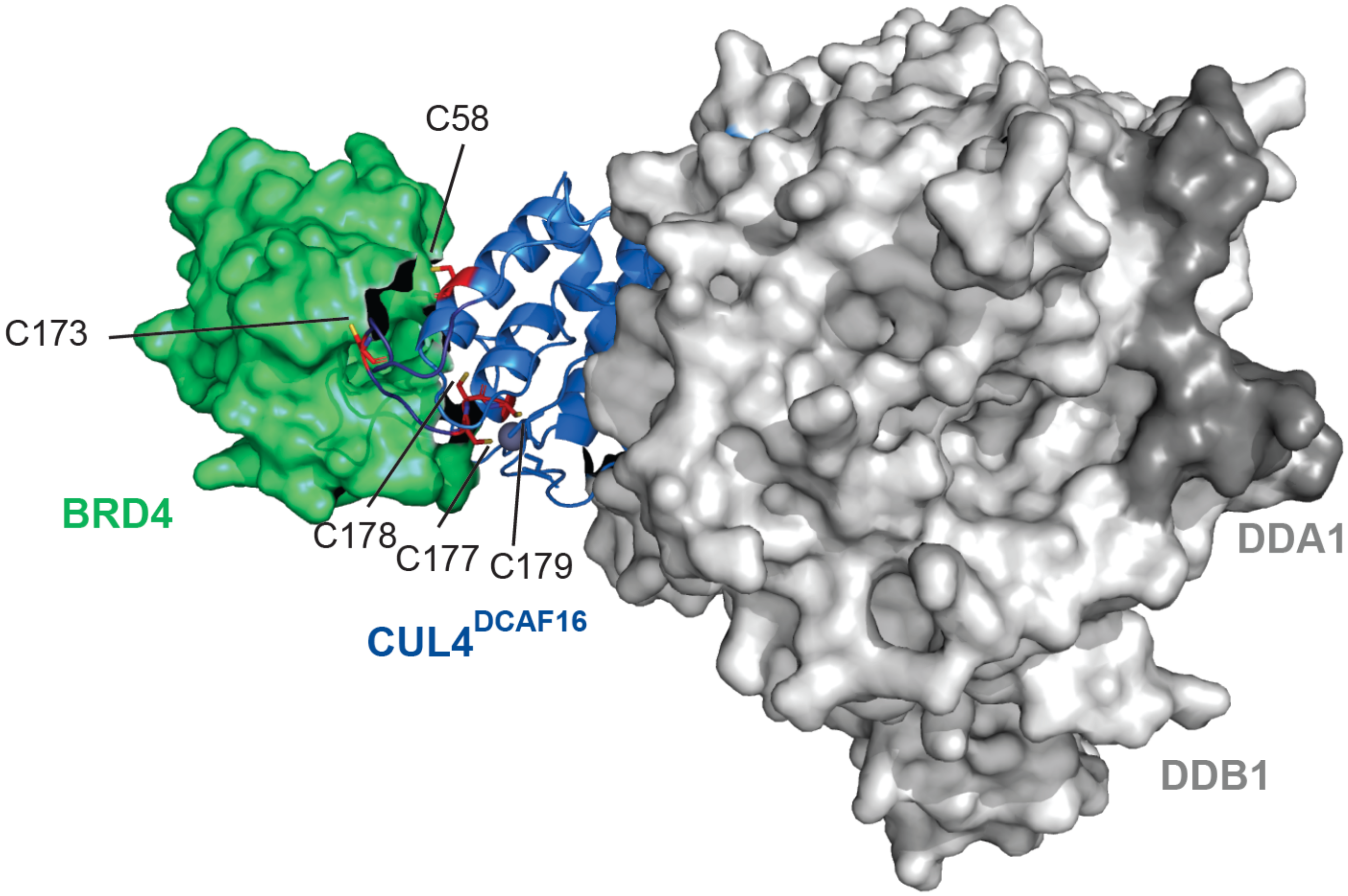


**Figure S4. Model of CUL4$^{DCAF16}$-DDB1-DDA1 complex with BRD4 showing C173, C177-179, and C58.** The model is based on a structure previously solved of DCAF16-DDB1-DDA1 in complex with BRD4 in the presence of an electrophilic BRD4 molecular glue degrader MMH2 (PDB: 8g46)[34] that did not include the disordered loop containing C173. We have removed MMH2. To add the loop containing C173 not resolved in the reported structure, the Pymol builder function was used to add the residues to the model structure, and the loop was then fit to a potential conformation via homology modeling with ModLoop [44,45].

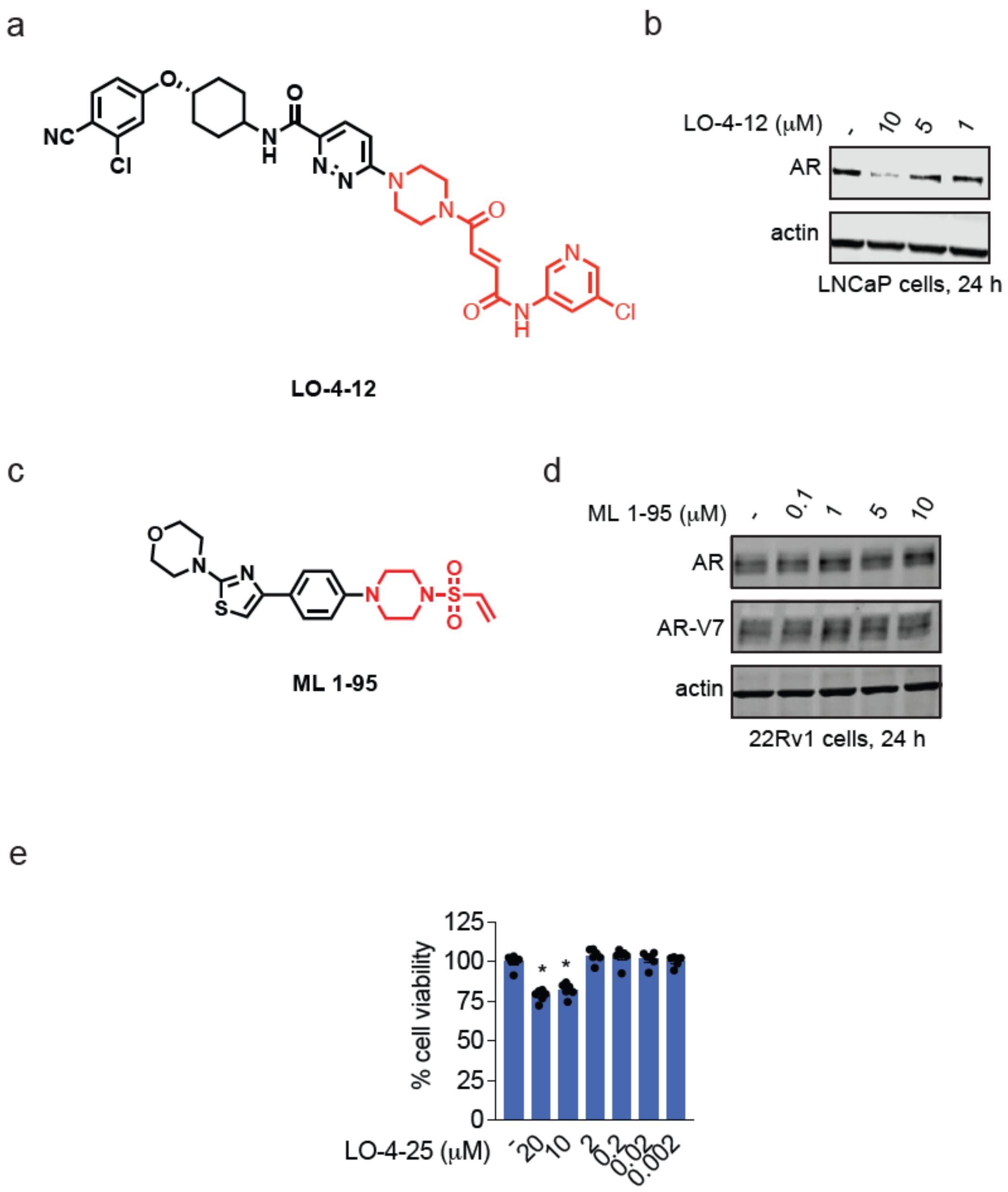


**Figure S5. Characterization of LO-4-12, ML 1-95, and LO-4-25. (a)** Structure of LO-4-12, an AR targeting ligand from the ARV110 PROTAC linked to the truncated fumaramide handle in red. **(b)** AR degradation in LNCaP cells. LNCaP cells were treated with DMSO vehicle or LO-4-12 for 24 h and AR and loading control actin levels were assessed by SDS/PAGE and Western blotting. **(c)** Structure of ML 1-95, an AR DNA binding domain ligand linked to our previously discovered DCAF16 C119-targeting vinyl sulfonylpiperazine handle in red. **(d)** ML 1-95 does not degrade AR or AR-V7. 22Rv1 cells were treated with DMSO vehicle or ML 1-95 for 24 h after which AR and AR-V7 and loading control actin levels were assessed by SDS/PAGE and Western blotting. **(e)** LO-4-25 effects on cell viability. 22Rv1 cells were treated with DMSO vehicle or LO-4-25 for 24 h, and cell viability was assessed with CellTiter-Glo. Blots in **(b,d)** are representative of n=3 biologically independent replicates per group. Data shown in **(e)** are from n=5 biologically independent replicates per group and show individual replicate values and average ± sem.

### Materials and Methods

#### Cell Culture

HEK293T and HEK293 cells were obtained from the UC Berkeley Cell Culture Facility and were cultured in Dulbecco's Modified Eagle Medium (DMEM) containing 10% (v/v) fetal bovine serum (FBS) and maintained at 37 °C with 5% $CO_2$. C33A cells were purchased from the American Type Culture Collection (ATCC) and were cultured in DMEM containing 10% (v/v) FBS and maintained at 37 °C with 5% $CO_2$. K562 cells were obtained from the UC Berkeley Cell Culture Facility and were cultured in Iscove's Modified Dulbecco's Medium (IMDM) containing 10% (v/v) FBS and maintained at 37 °C with 5% $CO_2$. MV-4-11 cells were obtained from the ATCC and were cultured in IMDM containing 10% (v/v) FBS and maintained at 37 °C with 5% $CO_2$. Mino cells were obtained from the ATCC and were cultured in RPMI-1640 Medium containing 10% (v/v) FBS and maintained at 37 °C with 5% $CO_2$. LNCaP cells were obtained from the UC Berkeley Cell Culture Facility and were cultured in DMEM containing 10% (v/v) FBS and maintained at 37 °C with 5% $CO_2$. 22Rv1 cells were obtained from the UC Berkeley Cell Culture Facility and were cultured in RPMI-1640 Medium containing 10% (v/v) FBS and maintained at 37 °C with 5% $CO_2$. HEK293 DCAF16 knockout cells were purchased from Ubigene Biosciences and were cultured in DMEM containing 10% (v/v) FBS and maintained at 37 °C with 5% $CO_2$. Unless otherwise specified, all cell culture materials were purchased from Gibco. It is not known whether HEK293T cells are from male or female origin.

#### Western Blotting

Cells were washed twice with cold PBS, scraped, and pelleted by centrifugation (1,200 g, 5 min, 4 °C). Pellets were resuspended in PBS, lysed by sonication or RIPA lysis buffer (Thermo Scientific), clarified by centrifugation (12,000 g, 10 min, 4 °C), and lysate was transferred to new low-adhesion microcentrifuge tubes. Proteome concentrations were determined using the BCA assay and lysate was diluted to appropriate working concentrations. Proteins were resolved by SDS/PAGE and transferred to nitrocellulose membranes using the Trans-Blot Turbo transfer system (Bio-Rad). Membranes were blocked with 5% BSA in Tris-buffered saline containing Tween 20 (TBS-T) solution for 1 hr at RT, washed in TBS-T, and probed with primary antibody diluted in recommended diluent per manufacturer overnight at 4 °C. After 3 washes with TBS-T, the membranes were incubated in the dark with IR680- or IR800-conjugated secondary antibodies at 1:10,000 dilution in 5 % BSA in TBS-T at RT for 1 h. After 3 additional washes with TBST, blots were visualized using an Odyssey Li-Cor fluorescent scanner. The membranes were stripped using ReBlot Plus Strong Antibody Stripping Solution (EMD Millipore, 2504) when additional primary antibody incubations were performed. Antibodies used in this study were BRD4 (Abcam ab128874), CDK4 (Abcam ab108357), CDK6 (Cell Signaling Technology DCS83), GAPDH (Cell Signaling Technology 14C10), Beta Actin (Cell Signaling Technology 13E5), SMARCA2 (Abcam ab240648), BRG1 (SMARCA4) (Cell Signaling Technology D1Q7F), Androgen Receptor (Cell Signaling Technology D6F11).

#### Bortezomib or MLN4924 Rescue Studies

2E6 of HEK293T cells per 3 mL of media were plated in 6-cm plates and left overnight to adhere. Cells were pretreated for 1 h with either Bortezomib (Cayman, C835F70) or MLN4924 (Tocris Bioscience, 649910) at a final concentration of 1 μM. Cells were then treated with degrader compound until desired time point. Cells from both the supernatant and on the plate were harvested and assessed via western blot.

### Cell Viability Assay

Cells were seeded in 96-well white plates overnight and then treated with DMSO vehicle control or degrader compounds and incubated at 37 °C for 24 h. Cell viability assay was performed using CellTiter-Glo® 2.0 reagent (Promega, G9241) according to manufacturer's protocol. Luminescent signals were measured using the Tecan Spark Plate reader (30086376).

### Generation of BRD4-GFP Reporter Cells

For lentivirus production, mGFP-tagged BRD4 (origene, RC216879L2), pMD2.G (Addgene, 12259) and psPAX2 (Addgene, 12260) were transfected into HEK293T cells using Lipofectamine 2000 (ThermoFisher, 11668027). The virus-containing medium was collected and filtered after 48 hours and was used to infect HEK293T cells with 1:1000 dilution of polybrene (Sigma-Adrich, TR-1003-G). After 48 hours, the infected cells were enriched for GFP signal with FACS.

### Generation of BRD4-GFP Stably Expressing Cas9

HEK293T cells were seeded into a 6-well plate in DMEM + 10% FBS and cultured to ~50% confluency after 24 hours. Cells were then transfected using Mirus LT1 transfection reagent with 500 ng of 3rd generation lentiviral packaging vector mix (equal parts pMDLg/pRRE [Addgene #12251], pRSV-Rev [Addgene #12253], and pMV2.g [Addgene #12259]) and 500 ng pLenti-Cas9-blast (Addgene #52962) at a 3:1 Mirus:DNA ratio.

BRD4-GFP reporter cells were seeded into a 6-well plate to achieve ~80% confluency the following day. After 24 hours, cells were treated with 0.5 mL fresh DMEM + 10% FBS containing 8 μg/mL polybrene and 0.5 mL pLenti-Cas9-blast lentiviral supernatant. Viral media was removed after 24 hours, and cells were expanded for another 24 hours before antibiotic selection with 4 μg/mL blasticidin (Thermo Fisher Scientific #A1113903). Surviving BRD4-GFP Cas9 reporter cells were frozen at −80°C and stored in liquid nitrogen. Cas9 activity was validated using the mCherry self-cutting system as previously described[1]. Briefly, cells were transfected with a plasmid expressing mCherry and a mCherry sgRNA, and mCherry fluorescence was assessed via flow cytometry after at least one week of growth.

### Flow Cytometry Fluorescence Intensity Measurements and Degradation Kinetics

Fluorescence intensities of BRD4-GFP cells were measured using an LSR Fortessa (BD Biosciences) at the UC Berkeley Flow Cytometry Core Facility. Approximately 500,000 cells were cultured in DMEM + 10% FBS, collected, and kept on ice. Cells were vortexed and analyzed at 1,000 events per second for a total of 10,000 events.

For degradation kinetic experiments, cells were treated with or without the degrader at specified times and doses. Live cells were gated using SSC-A vs. FSC-A, and single cells were gated using FSC-H vs. FSC-A. GFP signal was detected using the 488 nm laser, and mean fluorescence intensity (MFI) was calculated with FlowJo v10. To account for background fluorescence, 293T wild-type cells lacking the BRD4-GFP reporter were used as controls. The average MFI from control cells was subtracted from the BRD4-GFP reporter MFI at all time points and doses. Percent GFP remaining was calculated. Graphs were generated using GraphPad Prism (v8–10) with default settings.

**Lentiviral Transduction of the UBAL sgRNA Library into BRD4-GFP Cas9 HEK293T cells**

CRISPR-Cas9 screens were performed using the UBAL sgRNA library, which includes 20,710 elements: 18,710 sgRNAs targeting 1,871 genes (~10 sgRNAs per gene) and 2,000 negative control sgRNAs[2]. The library targets genes involved in ubiquitin conjugation, deubiquitination, ubiquitin-like conjugation, proteasome, autophagy, and lysosome pathways. The UBAL sgRNA library was lentivirally integrated into BRD4 reporter cells expressing Cas9. HEK293T cells were plated in 15-cm dishes and transfected the following day using the Mirus LT1 transfection reagent (Mirus Bio LLC) with the UBAL sgRNA library and lentiviral packaging vector mix (equal parts pVSV-G, pMDL, pRSV). Viral supernatant was collected, filtered through a 0.45 µm bottle-top filter (Nalgene, #124-0045), and stored at 4°C.

Prior to infecting BRD4-GFP Cas9 reporter cells, the viral titer was estimated by serial dilution. Cells were then infected with viral supernatant in the presence of 8µg/mL polybrene to achieve a 20%–50% mCherry-positive population, ensuring at least 200× coverage. Following viral transduction, cells were selected in the presence of 2 µg/mL puromycin (Thermo Fisher Scientific # A1113803). At least 1,000× coverage was maintained throughout the selection process. Surviving cells were frozen at −80°C and stored in liquid nitrogen.

**BRD4-GFP UBAL sgRNA Library CRISPR-Cas9 Screen**

UBAL library-infected BRD4-GFP Cas9 HEK293T cells were thawed and maintained at 1,000× coverage. On the day of sorting, cells were treated with 100 nM HRG038 or DMSO for 12 hr, dissociated using TrypLE™ Express Enzyme (Gibco #12605010), collected by centrifugation (500 × g for 5 minutes), washed with 1x DPBS (Gibco #1419-144), and resuspended in phenol red-free media (HyClone #16777-406) supplemented with 3% FBS and 1% fatty acid-free BSA. Cells were passed through 70-µm cell strainers (Falcon #352350) and kept on ice until FACS.

Cells were sorted using a BD FACS Aria Fusion Cell Sorter (BD Biosciences) at the UC Berkeley Flow Cytometry Core Facility. The top 5% and bottom 75% GFP+ cells were isolated using the following gating hierarchy: SSC-A v FSC-A polygonal gate for live cells, FSC-H v FSC-A polygonal gate for singlets, a histogram gate for mCherry+ cells using the 561 nm yellow-green laser, and a histogram gate for the top 5% and bottom 75% GFP+ cells using the 488 nm blue laser. Samples were kept at 4°C and sorting was performed using a 70 µm nozzle at a flow rate of 10-15,000 events/s, with four-way purity. Approximately $1.1 \times 10^6$ top 5% and $15\text{-}16 \times 10^6$ bottom 75% GFP+ cells were collected to ensure 1,000× coverage of the ~2,000-element UBAL library. Sorting was conducted in duplicate.

Genomic DNA was extracted using the QIAamp DNA Blood Midi Kit (Qiagen) as per the manufacturer's instructions. Guide sequence libraries were prepared from genomic DNA by two rounds of PCR using Herculase II Fusion DNA Polymerase (Agilent #600679)[1]. First, sgRNAs were amplified using the following reaction mix for a 100 µL reaction: 10 µg genomic DNA, 5x Herculase buffer (20 µL), 100 µM primers oMCB_1562 (1 µL) and oMCB_1563 (1 µL), 100 mM dNTPs (1 µL), 2 µL Herculase II Fusion DNA Polymerase, and nuclease-free water. PCR conditions were: 1x 98°C (2 min); 18x 98°C (30 sec), 59.1°C (30 sec), 72°C (45 sec); 1x 72°C (3 min). Next, amplicons were indexed using Illumina TruSeq LT adapters. The following reaction mix was used for indexing: 5 µL PCR1 product, 5x Herculase buffer (20 µL), 100 µM primers oMCB_1439 (0.8 µL) and barcoded oMCB_1440 (0.8 µL), 100 mM dNTPs (2 µL), 2 µL Herculase II Fusion DNA Polymerase, and nuclease-free water. PCR conditions were: 1x 98°C (2 min); 20x 98°C (30 sec), 59.1°C (30 sec), 72°C (45 sec); 1x 72°C (3 min).

PCR products were separated on a 2% TBE-agarose gel, purified using the QIAquick Gel Extraction Kit (Qiagen #28704), and assessed for quality using a Fragment Analyzer (Agilent). Amplicons from each library (GFPhigh and GFPlow) were pooled based on concentrations (as determined by Qubit Fluorometric Quantification) and the number of elements in each library. The sgRNA sequences were analyzed by deep sequencing on an Illumina NextSeq instrument at the UC Berkeley QB3 Functional Genomics Laboratory, using the standard Illumina indexing primer and custom sequencing primer oMCB_1672[1].

Sequence reads were aligned to the sgRNA reference library using Bowtie 2 software. For each gene, the gene effect and score (representing the likely maximum effect size and corresponding score) were calculated using the Cas9 high-throughput Maximum Likelihood Estimator (casTLE) statistical framework[3,4], and p-values were derived[3,4].

*Primer sequences*:

| Primer | Sequence |
|---|---|
| oMCB_1562 | aggcttggatttctataacttcgtatagcatacattatac |
| oMCB_1563 | acatgcatggcggtaatacggttatc |
| oMCB_1439 | caagcagaagacggcatacgagatgcacaaaagg aaactcaccct |
| oMCB_1440-AD001 | aatgatacggcgaccaccgagatctacacGATCGGAAGAGCACACGTCTGAACTCCAGTCA CATCACGCGACTCGGTGCCACTTTTTC |
| oMCB_1440-AD020 | aatgatacggcgaccaccgagatctacacGATCGGAAGAGCACACGTCTGAACTCCAGTCA CGTGGCCCGACTCGGTGCCACTTTTTC |
| oMCB_1440-AD008 | aatgatacggcgaccaccgagatctacacGATCGGAAGAGCACACGTCTGAACTCCAGTCA CACTTGACGACTCGGTGCCACTTTTTC |
| oMCB_1440-AD023 | aatgatacggcgaccaccgagatctacacGATCGGAAGAGCACACGTCTGAACTCCAGTCA CGAGTGGCGACTCGGTGCCACTTTTTC |
| oMCB_1440-AD011 | aatgatacggcgaccaccgagatctacacGATCGGAAGAGCACACGTCTGAACTCCAGTCA CGGCTACCGACTCGGTGCCACTTTTTC |
| oMCB_1440-AD027 | aatgatacggcgaccaccgagatctacacGATCGGAAGAGCACACGTCTGAACTCCAGTCA CATTCCTCGACTCGGTGCCACTTTTTC |

| | |
|---|---|
| oMCB_1440-AD022 | aatgatacggcgaccaccgagatctacacGATCGGAAGAGCACACGTCTGAACTCCAGTCACCGTACGCGACTCGGTGCCACTTTTTC |
| oMCB_1440-AD025 | aatgatacggcgaccaccgagatctacacGATCGGAAGAGCACACGTCTGAACTCCAGTCACACTGATCGACTCGGTGCCACTTTTTC |
| oMCB_1672 | GCCACTTTTTCAAGTTGATAACGGACTAGCCTTATTTAAACTTGCTATGCTGTTTCCAGCTTAGCTCTTAAAC |

**Purification of DCAF16-DDA-DDB1 Complex**

N-terminal ZZ-His-TEV tag full-length DCAF16, full-length untagged and DDA1 and untagged full-length DDB1 with an internal truncation of residues 285-689 were cloned into pFastBac and co-expressed in Sf21 at a 1:2:2 virus ratio. Cells expressing heterotrimer protein complex were resuspended and lysed in 100mM HEPES (pH 8.0), 250 mM NaCl, 20 mM Imidazole, 1mM TCEP with EDTA-free protease inhibitors by douncing on ice using 10-20 passes. The soluble protein was clarified via centrifugation, and the tagged complex was purified via Ni-NTA with a 3-hour incubation, followed by washing with 30 mM Imidazole and elution with 500 mM Imidazole. The purification tag was removed using TEV protease at 100 unit/mg. Subsequently, DCAF16 was dephosphorylated using Lambda Phosphatase with a 1:40 molar ratio at 4°C for 24 hours to reduce hyperphosphorylation. The treated heterotrimer was purified from modifying components via anion exchange by diluting the protein to 50 mM NaCl and performing a shallow gradient anion exchange. The protein complex was polished using a HiPrep S200 Superdex in 20mM HEPES pH 7.5, 150 mM NaCl, 1 mM TCEP, yielding 9 mg/L. Protein was concentrated to 20 mg/mL and stored at -80°C.

**Alkyne Probe Labeling**

Purified DCAF16-DDA1-DDB1 complex (5 µg/sample) was buffer exchanged into 50 µL of 1x PBS using Zeba™ Spin Desalting Columns, 7 MWCO (ThermoFisher, 89877). Samples were treated with either DMSO vehicle or alkyne probes, LO-3-44 or LO-4-06, at room termperature for 1 hr. To each sample, a cocktail of 0.25 µL 5 mM Carboxyrhodamine 110 Azide (Vector Labs, CCT-AZ105), 1 µL of 50 mM TCEP in $H_2O$, 1 µL of 50 mM $CuSO_4$ in $H_2O$, and 3 µL of TBTA ligand (1.7 mM in 1:4 DMSO/tBuOH, Cayman Chemical, 18816) was added and incubated at room temperature for 1 hr. The reaction was stopped by addition of 4 × reducing Laemmli SDS sample loading buffer (Alfa Aesar). After boiling at 95 °C for 5 min, the samples were separated on precast 4–20% Criterion TGX gels (Bio-Rad). Probe-labeled proteins were analyzed by in-gel fluorescence using a ChemiDoc MP (Bio-Rad). Imaged gels were stained using Pierce Silver Stain Kit (Thermo Scientific, 24612) following manufacturer's instructions.

**LO-361-Competed Targets from LO-406 Probe Pulldown Proteomics**

HEK293T cells were harvested, lysed, and the proteome concentration was adjusted to 5 mg/mL in 500 µL of PBS using the BCA assay. HEK293T cell lysate were pre-treated with DMSO vehicle or LO-361 (50 µM) for 1 h at room temperature prior to LO-406 probe labeling (10 µM) at room temperature for 1 h. To each tube containing cell lysate, the following reagents were added: 10 µL of 10 mM biotin picolyl azide (Sigma Aldrich, 900912) in DMSO, 10 µL of 50 mM TCEP in $H_2O$, 10 µL of 50 mM $CuSO_4$ in $H_2O$, and 30 µL of TBTA ligand (1.7 mM in 1:4 DMSO/tBuOH, Cayman Chemical, 18816). The reaction mixture was incubated at room temperature for 60 minutes, and the reaction was quenched by protein precipitation. Precipitated pellets were washed using 500 µL of MeOH and centrifuged again to yield white pellets. Samples were resuspended in 1.2% SDS-PBS (1 mL), completely dissolved, and heated to 90 °C for 5 minutes. The soluble proteome was then diluted with 5 mL of PBS and further incubated with high-capacity streptavidin-agarose beads (100 µL/sample, ThermoFisher Scientific, 20357). Beads and lysates were incubated overnight at 4 °C with rotation. On the following day, beads were suspended and washed three times with 0.1% SDS-PBS, PBS, and $H_2O$. Washed beads were resuspended in 6 M Urea/PBS (500 µL), and the samples were further treated with DTT and iodoacetamide. After removing the supernatant, beads were resuspended in 100 µL of 50 mM TEAB and enzymatically digested overnight using sequencing-grade trypsin (Promega, V5111). Digested peptides were eluted through centrifugation and labeled using commercially available TMTsixplex tags (ThermoFisher, 90061). After labeling, 35 µg of each labeled sample was combined and dried using a vacufuge. Dried samples were redissolved with 300 μL of 0.1% TFA in $H_2O$ and further fractionated using high-pH reversed-phase peptide fractionation kits (ThermoFisher, 84868) following the manufacturer's protocol. Dried fractions were then resuspended in 25 μL of 0.1 % Formic acid/$H_2O$ (w/v) to be analyzed by LC-MS/MS.

**Quantitative Global TMT Proteomics**

Cells were treated with either DMSO vehicle or compound (HRG038 (1 μM, 16 h) or LO361 (1 μM, 16 h)) and lysate was prepared as described above. Briefly, 25-100 μg protein from each sample was reduced, alkylated and tryptically digested overnight. Individual samples were then labeled with isobaric tags using commercially available TMTsixplex (Thermo Fisher Scientific, P/N 90061) kits, in accordance with the manufacturer's protocols. Tagged samples (20 μg per sample) were combined, dried using a vacuum concentrator at 30 °C, resuspended with 300 μL 0.1% TFA in $H_2O$, and fractionated using high pH reversed-phase peptide fractionation kits (Thermo Fisher Scientific, P/N 84868) according to the manufacturer's protocol. Fractions were dried using a vacuum concentrator at 30 °C, resuspended with 50 μL 0.1% FA in $H_2O$, and analyzed by LC-MS/MS as described below.

**TMT Proteomics Analysis**

Quantitative TMT-based proteomic analysis was performed as previously described using a Thermo Eclipse with FAIMS LC-MS/MS [5]. Acquired MS data was processed using ProLuCID search methodology in IP2 v.3-v.5 (Integrated Proteomics Applications, Inc.) [6]. Trypsin cleavage specificity (cleavage at K, R except if followed by P) allowed for up to 2 missed cleavages. Carbamidomethylation of cysteine was set as a fixed modification, methionine oxidation, and TMT-modification of N-termini and lysine residues were set as variable modifications. Reporter ion ratio calculations were performed using summed abundances with the most confident centroid selected from the 10 ppm window. Only peptide-to-spectrum matches that are unique assignments to a given identified protein within the total dataset are considered for protein quantitation. High confidence protein identifications were reported with a <1% false discovery rate (FDR) cut-off. Differential abundance significance was estimated using ANOVA with Benjamini-Hochberg correction to determine p-values.

**Knock Out Cell Line Generation**
The DCAF16 knock-out cell line was purchased from Ubigene with guide sequences AGAGGGGGCCATTCAGGAAT TGG and TTCTGACAAGTGGTCAGGAG AGG (catalog number YKO-H721).

**Site-Directed Mutagenesis on FLAG-tagged DCAF16 Plasmid**

Site-directed mutagenesis was performed on FLAG-tagged wild type DCAF16 plasmid (Origene, RC208716L3) using Q5® Site-Directed Mutagenesis Kit (NEB, E0552S) according to the manufacturer's protocol. The sequences of the primers used are shown below.

**C58S Primer (Forward)**: GCAGGTTAAGAGCCTTTTAAAATATTC

**C58S Primer (Reverse)**: CAGGCAAGACTCTCAAG

**C119S Primer (Forward)**: TCTGGCCTCTAGCGGAGTCCCAC
**C119S Primer (Reverse)**: GGGGGCCATTCAGGAATT

**C173S Primer (Forward):** CCCTAATTCAAGCGTTTCTGGGTG **(Reverse):** CACCCAGAAACGCTTGAATTAGGG

**C178S Primer (Forward):** TTCTGGGTGTAGCTGTGGCTGGC **(Reverse):** GCCAGCCACAGCTACACCCAGAA

**Plasmid Isolation**

E. coli containing desired plasmids were pelleted, lysed, and neutralized using QIAGEN Plasmid Plus Midi Kit (Qiagen, 12943) according to the manufacturer's protocol. The eluted plasmid concentrations were determined using Nanodrop quantification.

**Expression of FLAG-tagged Wild Type DCAF16 and Mutants in DCAF16 Knockout Cells**

For lentivirus production, FLAG-tagged wild type DCAF16 or FLAG-tagged DCAF16 mutant plasmids, pMD2.G (Addgene, 12259) and psPAX2 (Addgene, 12260) were transfected into HEK293T cells using Lipofectamine 2000 (ThermoFisher, 11668027). The virus-containing medium was collected and filtered after 48 hours and was used to infect HEK293 DCAF16 knockout cells with 1:1000 dilution of polybrene (Sigma-Adrich, TR-1003-G). After 48 hours, the infected cells were selected with puromycin (2 µg/mL).

**Mapping Site of Modification of BRD4 Degraders on the DCAF16-DDA1-DDB1 Complex**

25 µg of DCAF16-DDA1-DDB1 complex was buffer exchanged into 50 µL of 1X PBS using Zeba™ Spin Desalting Columns, 7 MWCO (ThermoFisher, 89877) and incubated with 50 µM HRG038 or LO-3-61 for 1 hr at room temperature. The samples were reduced and alkylated with DTT and iodoacetamide. Proteins were buffer exchanged again into 50 mM ammonium bicarbonate buffer, and then enzymatically digested for 4 hr at 37°C with agitation using sequencing-grade trypsin (Promega, V5111) and 0.01% ProteaseMAX™ Surfactant (Promega, V2071). The digestion was quenched by adding formic acid to give a concentration of 0.5% acid/$H_2O$ (w/v) and centrifuged at 21,000 xg for 30 minutes. The supernatant was then analyzed by LC-MS/MS.

**Site of Modification Proteomics Analysis**

Site of modification proteomic analysis was performed as previously described using a Thermo Eclipse with FAIMS LC-MS/MS[5,7,8] Acquired MS data was processed using MSFragger (v4.1) search methodology in FragPipe v19.1[9,10]. Semi-enzymatic cleavage was allowed, with trypsin cleavage specificity (cleavage at K, R) allowed for up to 2 missed cleavages. PSM validation was performed with PeptideProphet [11], protein interference was performed with ProteinProphet, and FDR filtering was performed with Philosopher (v5.1.0) [12]. Carbamidomethylation of cysteine, methionine oxidation, and modification of cysteine with HRG038 or LO-3-61, respectively, were set as variable modifications. Spectra of PSMs containing modifications corresponding to HRG038 or LO-3-61 were viewed with FP-PDV[13].

**DCAF16 Knockdown Studies**

ON-TARGETplus Human DCAF16 SMARTpool siRNA (Horizon, 54876) were transfected into 22RV1 cells using Lipofectamine™ RNAiMAX (ThermoFisher, 13778100). ON-TARGETplus Non-Targeting Control siRNA (Horizon, D-001810-01-20) was used as a control. After 48 hours, transfected cells were either harvested for RT-qPCR to measure knockdown or treated with degrader compounds for downstream western blotting.

**RT-qPCR Analysis**

Total RNA was extracted from cells using Monarch® Total RNA Miniprep Kit (NEB, T2010S) according to the manufacturer's protocol. cDNA was synthesized and gene expression was confirmed by qPCR using Luna® Universal One-Step RT-qPCR Kit (NEB, E3005S) following the manufacturer's protocol with the CFX Connect Real-Time PCR Detection System (BioRad). Relative DCAF16 gene expression was normalized to the GAPDH or ACTB gene. The sequences of the qPCR primers are shown below.

**GAPDH Primer (Forward):** GTCTCCTCTGACTTCAACAGCG
**GAPDH Primer (Reverse):** ACCACCCTGTTGCTGTAGC

**ACTB Primer (Forward):** CACCATTGGCAATGAGCGGTTC
**ACTB Primer (Reverse):** AGGTCTTTGCGGATGTCCACGT

**DCAF16 Primer #1 (Forward):** TGACCACTTGTCAGAATCAGAA
**DCAF16 Primer #1 (Reverse):** AGAGGCGATAAGTTGGGCAC

**DCAF16 Primer #2 (Forward):** TGGATCCAAGCACACCAGTC
**DCAF16 Primer #2 (Reverse):** TGGTTCCAGTTTGGGGACAC

**DCAF16 Primer #3 (Forward):** CAATTCCTGAATGGCCCCCT
**DCAF16 Primer #3 (Reverse):** GTGCTCCATTTAGAGTGGCA

**DCAF16 Primer #4 (Forward):** AGTCTTGCCTGGCAGGTTAAG
**DCAF16 Primer #4 (Reverse):** GGGACTTGTAAGAGGCTTTTGAA

**Data Availability Statement**

The datasets generated during and/or analyzed during the current study are available from the corresponding author on reasonable request. We would be happy also provide the raw Western blotting data upon request.

**Code Availability Statement**

Data processing and statistical analysis algorithms from our lab can be found on our lab’s Github site: https://github.com/NomuraRG, and we can make any further code from this study available at reasonable request.

**Synthetic Methods and Characterization**

All chemical reactions were carried out under a nitrogen atmosphere with dry solvents under anhydrous conditions, unless otherwise noted. Reagents were purchased at the highest commercial quality and used without further purification, unless otherwise stated. Room temperature is defined as between 21-25 °C. Reactions were stirred magnetically and monitored by thin layer chromatography (TLC) using TLC plates precoated with silica gel 60 F254 on aluminium (Merck KGaA). Detection was by UV (254 nm and 365 nm) or chemical stain ($KMnO_4$, ninhydrin, iodine). Solvents were removed *in vacuo* using either a Buchi R-300 Rotavapor (equipped with an I-300 Pro Interface, B-300 Base Heating Bath, Welch 2037B-01 DryFast pump, and VWR AD15R-40-V11B Circulating Bath). Solvents for silica gel chromatography were used as supplied by Sigma-Aldrich. Automated flash chromatography was performed on a Biotage Isolera instrument, equipped with a UV detector. Chromatograms were recorded at 254 and 280 nm. If additional purification is needed, compound was further purified using Thermo Scientific's semi-prep reversed phase high-performance liquid chromatography equipped (RP-HPLC: Ultimate 3000 HPLC) equipped with C18 column (Luna® 10 µm c18(2), 100 Å, Serial #:5293-0084). Elution of the sample was monitored using DIONEX UltiMate 3000. Eluting buffer A: 100% distilled water + 0.1% trifluoracetic acid (TFA). Eluting buffer B: 95% acetonitrile + 5% distilled water + 0.1% TFA. High-resolution mass spectra (HRMS) were obtained using Q Exactive™ Plus Hybrid Quadrupole-Orbitrap™ Mass Spectrometer. $^{1}H$ and $^{13}C$ Nuclear Magnetic Resonance (NMR) spectra were recorded on BRUKER AV (600 MHz and 700 MHz) and NEO (500 MHz) spectrometers. Measurements were carried out at ambient temperature. Chemical shifts (δ) are reported in ppm with the residual solvent signal as internal standard (chloroform at 7.26 and 77.2 ppm for $^{1}H$ NMR and $^{13}C$ NMR, respectively, methanol at 3.31 and 49.0, respectively and DMSO at 2.50 and 39.5, respectively). Multiplicity is reported as follows: singlet (s), doublet (d), doublet of doublet (dd) doublet of triplet (dt), triplet (t), triplet of doublet (td), quartet (q), and multiplet (m). Coupling constants (J) are reported in Hertz (Hz). $^{13}C$ NMR spectra were recorded with broadband 1 H decoupling.

**General Procedures**

**Amide Couplings**

**General Procedure A**

A mixture of the corresponding carboxylic acid (1.2 equiv.) and HATU (1.2 equiv.) was purged with $N_2$ for 5 minutes. The mixture was dissolved in N,N-dimethylformamide (DMF) (0.1 M), DIPEA (4 equiv.) was added and the reaction mixture was allowed to stir at ambient temperature for 30 minutes. The corresponding amine (1 equiv.) was dissolved in DMF (0.1 M) then added dropwise and the reaction mixture was stirred at ambient temperature overnight. The reaction was quenched with 5 times the reaction volume of 5% LiCl(aq) and extracted 3 times with dichloromethane (DCM). The organic extracts were washed once with brine, dried over $Na_2SO_4$, vacuum filtered, and concentrated *in vacuo*. The resultant residue was purified by silica gel flash chromatography to afford the title compound.

**Tert-butyloxycarbonyl Deprotection**

**General Procedure B**

The corresponding tert-butyloxycarbonyl protected amine (1 equiv.) was dissolved in DCM (0.1 M). Trifluoroacetic acid (32 equiv.) was added, and the reaction mixture was stirred at ambient temperature for 30 minutes to overnight. The volatiles were removed *in vacuo* and the crude residue was used without further purification, unless otherwise noted.

**Tert-butyl Ester Deprotection**

**General Procedure C**

The corresponding tert-butyl ester (1 equiv.) was dissolved in DCM (0.1 M). Trifluoroacetic acid (32 equiv.) was added, and the reaction mixture was stirred at ambient temperature for 30 minutes to overnight to afford the carboxylic acid. The volatiles were removed *in vacuo* and the crude residue was used without further purification, unless otherwise noted.

***tert*-butyl (*S*)-4-(2-(4-(4-chlorophenyl)-2,3,9-trimethyl-6*H*-thieno[3,2-*f*][1,2,4]triazolo[4,3-*a*][1,4]diazepin-6-yl)acetyl)piperazine-1-carboxylate (HY157)**

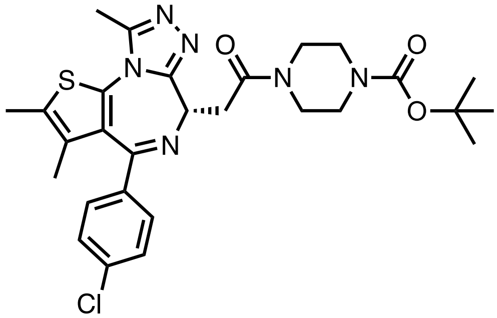

**General Procedure A** was followed with (S)-2-(4-(4-chlorophenyl)-2,3,9-trimethyl-6H-thieno [3,2-f][1,2,4]triazolo[4,3-a][1,4]diazepin-6-yl)acetic acid (JQ1-Acid) (257.9 mg, 0.64 mmol), HATU (245.1 mg, 0.64 mmol), DIPEA (0.37 mL, 2.1 mmol), and *tert*-butyl piperazine-1-carboxylate (100.0 mg, 0.54 mmol). The crude residue was purified by silica gel chromatography (0-10% MeOH in DCM) to afford 142.3 mg (47%) of the title compound as a yellow oil.

**$^1$H NMR** (500 MHz, $CDCl_3$) δ 7.41 (d, *J* = 8.2 Hz, 2H), 7.34 (d, *J* = 8.4 Hz, 2H), 4.82 (t, *J* = 6.6 Hz, 1H), 3.86 – 3.37 (m, 10H), 2.69 (s, 3H), 2.40 (s, 3H), 1.68 (s, 3H), 1.49 (s, 9H).

**$^{13}$C NMR** (126 MHz, $CDCl_3$) δ 169.09, 164.01, 155.69, 154.60, 149.96, 136.92, 136.45, 132.18, 131.03, 131.00, 130.49, 129.92, 128.75, 80.28, 54.30, 45.71, 41.70, 35.17, 28.40, 14.38, 13.11, 11.81.

**HRMS (ESI)** m/z calcd for $C_{28}H_{34}ClN_6O_3S^+$ [M+H]$^+$: 569.2102; found: 569.2072

**tert-butyl(S,E)-4-(4-(2-(4-(4-chlorophenyl)-2,3,9-trimethyl-6H-thieno[3,2-f][1,2,4]triazolo[4,3-a][1,4]diazepin-6-yl)acetyl)piperazin-1-yl)-4-oxobut-2-enoate (LO426)**

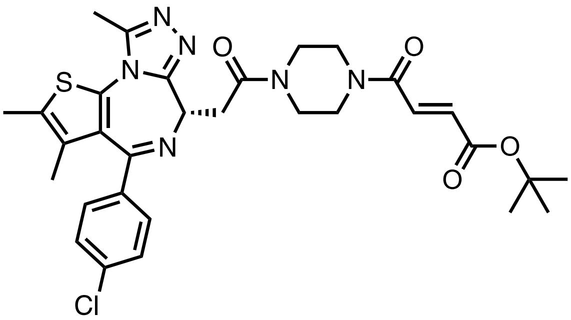

**General Procedure B** was followed with **HY157** (100.0 mg, 0.18 mmol), and TFA (0.43 mL, 5.62 mmol) for 2 hours. The crude material was used without further purification.

**General Procedure A** was followed with (E)-4-(tert-butoxy)-4-oxobut-2-enoic acid (36.3 mg, 0.21 mmol), HATU (80.2 mg, 0.21 mmol), DIPEA (0.12 mL, 0.70 mmol), and the amine from above (82.4 mg, 0.18 mmol). The crude residue was purified by silica gel chromatography (0-10% MeOH in DCM) to afford 45.1 mg (41%) of the title compound as a yellow-white powder.

**$^1$H NMR** (500 MHz, $CDCl_3$) δ 7.40 (d, *J* = 8.7 Hz, 2H), 7.36 – 7.28 (m, 3H), 6.73 (d, *J* = 15.2 Hz, 1H), 4.80 (t, *J* = 6.8 Hz, 1H), 4.05 – 3.47 (m, 10H), 2.67 (s, 3H), 2.41 (s, 3H), 1.68 (s, 3H), 1.52 (s, 9H).

**$^{13}$C NMR** (126 MHz, $CDCl_3$) δ 169.40, 169.24, 164.73, 164.05, 163.97, 163.85, 155.70, 149.94, 136.74, 133.80, 132.21, 130.92, 130.74, f130.46, 129.78, 128.73, 81.78, 54.61, 54.39, 46.16, 45.98, 45.75, 45.47, 42.05, 41.95, 41.87, 41.49, 35.32, 35.25, 31.66, 29.64, 28.02, 14.38, 13.09, 11.84.

**HRMS (ESI)** m/z calcd for $C_{31}H_{36}ClN_6O_4S^+$ [M+H]$^+$: 623.2202; found: 623.2142

**(*S*,*E*)-4-(4-(2-(4-(4-chlorophenyl)-2,3,9-trimethyl-6*H*-thieno[3,2-*f*][1,2,4]triazolo[4,3-*a*][1,4]diazepin-6-yl)acetyl)piperazin-1-yl)-*N*-((3-(4-fluorophenyl)-1,2,4-oxadiazol-5-yl)methyl)-4-oxobut-2-enamide (HRG034)**

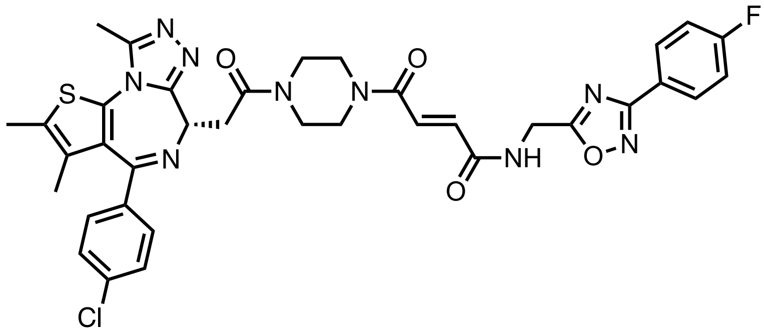

**General Procedure C** was followed with **LO426** (50.0 mg, 0.08 mmol), and TFA (0.20 mL, 2.57 mmol) for 2 hours. The crude material was used without further purification.
**General Procedure A** was followed with (3-(4-fluorophenyl)-1,2,4-oxadiazol-5-yl)methanamine (12.9 mg, 0.07 mmol), HATU (30.5 mg, 0.08 mmol), DIPEA (0.05 mL, 0.27 mmol), and the acid from above (45.5 mg, 0.08 mmol). The crude residue was purified by silica gel chromatography (0-10% MeOH in DCM) to afford 23.3 mg (47%) of the title compound as a yellow-white powder.
**$^{1}$H NMR** (500 MHz, $CDCl_3$) δ 8.08 – 8.00 (m, 2H), 7.73 (dd, *J* = 15.5, 9.7 Hz, 1H), 7.48 (d, *J* = 14.7 Hz, 1H), 7.39 (d, *J* = 8.4 Hz, 2H), 7.32 (d, *J* = 8.6 Hz, 2H), 7.20 – 7.08 (m, 3H), 4.85 (d, *J* = 5.6 Hz, 1H), 4.78 (td, *J* = 6.9, 2.8 Hz, 1H), 4.05 – 3.38 (m, 10H), 2.65 (s, 3H), 2.39 (s, 3H), 1.67 (s, 3H).
**$^{13}$C NMR** (126 MHz, $CDCl_3$) δ 175.70, 169.42, 167.65, 165.69, 164.39, 163.89, 163.68, 155.73, 149.97, 136.78, 136.70, 133.94, 132.16, 130.94, 129.79, 129.73, 129.66, 128.74, 122.54, 116.25, 116.07, 54.60, 54.41, 42.24, 41.91, 36.00, 35.26, 14.38, 13.10, 11.83.
**HRMS (ESI)** m/z calcd for $C_{36}H_{34}ClFN_9O_4S^+$ $[M+H]^+$: 742.2122; found: 742.2062

**(*S*,*E*)-*N*-(5-chloro-6-(2*H*-1,2,3-triazol-2-yl)pyridin-3-yl)-4-(4-(2-(4-(4-chlorophenyl)-2,3,9-trimethyl-6*H*-thieno[3,2-*f*][1,2,4]triazolo[4,3-*a*][1,4]diazepin-6-yl)acetyl)piperazin-1-yl)-4-oxobut-2-enamide (HRG038)**

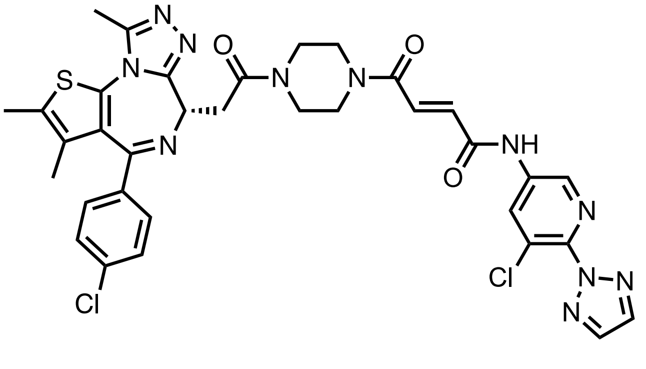

**General Procedure C** was followed with **LO426** (50.0 mg, 0.08 mmol), and TFA (0.20 mL, 2.57 mmol) for 2 hours. The crude material was used without further purification.
**General Procedure A** was followed with 5-chloro-6-(2*H*-1,2,3-triazol-2-yl)pyridin-3-amine (13.1 mg, 0.07 mmol), HATU (30.5 mg, 0.08 mmol), DIPEA (0.05 mL, 0.27 mmol), and the acid from above (45.5 mg, 0.08 mmol). The crude residue was purified by silica gel chromatography (0-10% MeOH in DCM) to afford 24.1 mg (48%) of the title compound as a yellow-white powder.
**$^{1}$H NMR** (500 MHz, $CDCl_3$) δ 10.29 (s, 1H), 8.76 – 8.63 (m, 2H), 7.94 – 7.90 (m, 2H), 7.57 (dd, *J* = 14.8, 9.9 Hz, 1H), 7.41 (d, *J* = 8.4 Hz, 2H), 7.33 (d, *J* = 6.4 Hz, 1H), 7.26 (d, *J* = 16.8 Hz, 1H), 4.81 (t, *J* = 6.8 Hz, 1H), 4.14 – 3.44 (m, 10H), 2.68 (s, 3H), 2.41 (s, 3H), 1.68 (s, 3H).

**$^{13}$C NMR** (126 MHz, $CDCl_3$) δ 169.55, 169.43, 164.53, 164.15, 163.94, 163.10, 155.76 (d, *J* = 17.5 Hz), 150.09 (d, *J* = 8.9 Hz), 143.65, 138.12, 137.98, 136.87, 136.81, 136.60, 135.98, 134.97, 134.29, 131.96, 131.64, 131.24, 131.10, 131.01, 130.95, 130.67, 130.55, 130.46, 130.33, 129.85, 128.76, 126.32, 54.64, 54.33, 45.87, 45.38, 42.38, 42.07, 41.83, 41.22, 35.35, 14.41, 13.13, 11.86.
**HRMS (ESI)** m/z calcd for $C_{34}H_{32}Cl_2N_{11}O_3S$ $[M+H]^+$: 744.1782; found: 744.1723

**(*S*,*E*)-4-(4-(2-(4-(4-chlorophenyl)-2,3,9-trimethyl-6*H*-thieno[3,2-*f*][1,2,4]triazolo[4,3-*a*][1,4]diazepin-6-yl)acetyl)piperazin-1-yl)-*N*-(4-(4-methylthiazol-5-yl)benzyl)-4-oxobut-2-enamide (HRG073)**

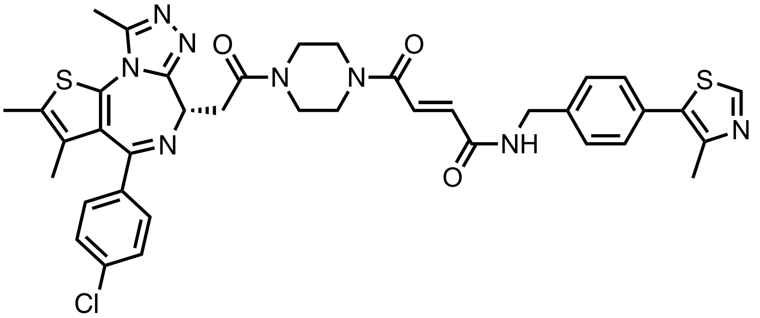

**General Procedure C** was followed with **LO426** (50.0 mg, 0.08 mmol), and TFA (0.20 mL, 2.57 mmol) for 2 hours. The crude material was used without further purification.
**General Procedure A** was followed with (4-(4-methylthiazol-5-yl)phenyl)methanamine (13.7 mg, 0.07 mmol), HATU (30.5 mg, 0.08 mmol), DIPEA (0.05 mL, 0.27 mmol), and the acid from above (45.5 mg, 0.08 mmol). The crude residue was purified by silica gel chromatography (0-10% MeOH in DCM) to afford 19.9 mg (47%) of the title compound as a yellow-white powder.
**$^1$H NMR** (500 MHz, $CDCl_3$) δ 8.66 (s, 1H), 7.47 (dd, *J* = 14.7, 2.2 Hz, 1H), 7.43 – 7.31 (m, 8H), 7.23 – 7.05 (m, 2H), 4.79 (q, *J* = 6.6 Hz, 1H), 4.60 (d, *J* = 5.9 Hz, 2H), 4.01 – 3.45 (m, 10H), 2.65 (s, 3H), 2.52 (s, 3H), 2.40 (s, 3H), 1.68 (s, 3H).
**$^{13}$C NMR** (126 MHz, $CDCl_3$) δ 169.38, 169.25, 164.23, 163.97, 163.84, 155.70, 150.42, 149.90, 148.60, 137.73, 136.76, 136.72, 135.33, 135.13, 132.18, 131.43, 131.26, 130.93, 130.79, 130.48, 129.80, 129.57, 129.41, 128.74, 128.15, 54.57, 54.41, 46.04, 45.87, 45.70, 45.41, 43.48, 42.12, 41.94, 41.36, 35.27, 16.12, 14.39, 13.11, 11.84, 11.82.
**HRMS (ESI)** m/z calcd for $C_{38}H_{38}ClN_8O_3S^+$ $[M+H]^+$: 753.2191; found: 753.2127

**(*S*,*E*)-4-(4-(2-(4-(4-chlorophenyl)-2,3,9-trimethyl-6*H*-thieno[3,2-*f*][1,2,4]triazolo[4,3-*a*][1,4]diazepin-6-yl)acetyl)piperazin-1-yl)-4-oxo-*N*-(4-phenoxybenzyl)but-2-enamide (HRG075)**

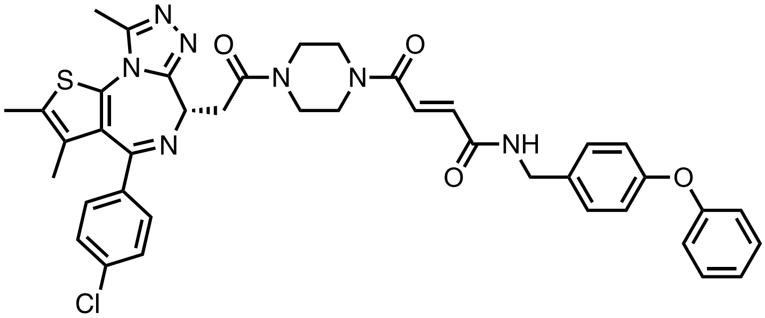

**General Procedure C** was followed with **LO426** (50.0 mg, 0.08 mmol), and TFA (0.20 mL, 2.57 mmol) for 2 hours. The crude material was used without further purification.
**General Procedure A** was followed with (4-phenoxyphenyl)methanamine (13.3 mg, 0.07 mmol), HATU (30.5 mg, 0.08 mmol), DIPEA (0.05 mL, 0.27 mmol), and the acid from above (45.5 mg, 0.08 mmol). The crude residue was purified by silica gel chromatography (0-10% MeOH in DCM) to afford 18.7 mg (37.4%) of the title compound as a yellow-white powder.

**¹H NMR** (500 MHz, $CDCl_3$) δ 7.45 (d, *J* = 14.1 Hz, 1H), 7.40 (d, *J* = 8.6 Hz, 2H), 7.36 – 7.28 (m, 4H), 7.28 (s, 3H), 7.10 (tt, *J* = 7.3, 1.1 Hz, 1H), 7.04 – 6.95 (m, 5H), 6.65 (s, 1H), 4.80 (t, *J* = 6.8 Hz, 1H), 4.53 (d, *J* = 5.1 Hz, 2H), 4.03 – 3.45 (m, 10H), 2.65 (s, 3H), 2.40 (s, 3H), 1.68 (s, 3H).
**¹³C NMR** (126 MHz, $CDCl_3$) δ 169.40, 169.26, 164.28, 164.22, 164.09, 163.96, 163.84, 157.07, 156.77, 155.73, 155.69, 149.93, 149.89, 136.73, 135.35, 135.21, 132.62, 132.20, 130.93, 130.78, 130.48, 129.82, 129.79, 129.45, 129.41, 129.32, 128.73, 123.37, 119.05, 118.90, 54.56, 54.45, 53.46, 46.02, 45.86, 45.68, 45.40, 43.35, 42.10, 41.91, 41.38, 35.28, 14.39, 13.11, 11.82.
**HRMS (ESI)** m/z calcd for $C_{40}H_{39}ClN_7O_4S^+$ $[M+H]^+$: 748.2467; found: 748.2402

**(*S*,*E*)-4-(4-(2-(4-(4-chlorophenyl)-2,3,9-trimethyl-6*H*-thieno[3,2-*f*][1,2,4]triazolo[4,3-*a*][1,4]diazepin-6-yl)acetyl)piperazin-1-yl)-*N*-(3-fluoro-4-thiomorpholinophenyl)-4-oxobut-2-enamide (HRG078)**

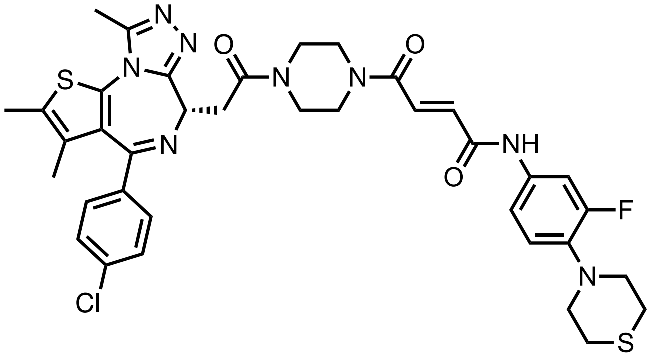

**General Procedure C** was followed with **LO426** (50.0 mg, 0.08 mmol), and TFA (0.20 mL, 2.57 mmol) for 2 hours. The crude material was used without further purification.
**General Procedure A** was followed with 3-fluoro-4-thiomorpholinoaniline (14.2 mg, 0.07 mmol), HATU (30.5 mg, 0.08 mmol), DIPEA (0.05 mL, 0.27 mmol), and the acid from above (45.5 mg, 0.08 mmol). The crude residue was purified by silica gel chromatography (0-10% MeOH in DCM) to afford 21.0 mg (41.3%) of the title compound as a yellow-white powder.
**¹H NMR** (500 MHz, $CDCl_3$) δ 9.39 (d, *J* = 6.2 Hz, 1H), 7.52 (dd, *J* = 27.2, 14.3 Hz, 2H), 7.39 (d, *J* = 7.5 Hz, 2H), 7.35 – 7.20 (m, 4H), 6.89 (t, *J* = 9.0 Hz, 1H), 4.80 (t, *J* = 6.8 Hz, 1H), 4.06 – 3.41 (m, 10H), 3.27 (t, 4H), 2.78 (t, 4H), 2.66 (s, 3H), 2.39 (s, 3H), 1.66 (s, 3H).
**¹³C NMR** (126 MHz, $CDCl_3$) δ 169.52, 169.44, 164.64, 164.53, 164.13, 164.07, 162.25, 156.55, 155.79, 154.60, 150.09, 137.92, 137.85, 136.90, 136.79, 136.21, 135.99, 133.63, 133.55, 132.26, 131.04, 130.96, 130.59, 129.91, 129.74, 128.85, 120.29, 120.26, 116.00, 109.21, 109.01, 54.72, 54.52, 53.49, 53.47, 46.13, 46.06, 45.94, 45.54, 42.42, 42.24, 42.03, 41.47, 35.38, 31.05, 29.80, 28.14, 14.49, 13.22, 11.95.
**HRMS (ESI)** m/z calcd for $C_{37}H_{39}ClFN_8O_3S_2^+$ $[M+H]^+$: 761.2254; found: 761.2199

**(*S*,*E*)-4-(4-(2-(4-(4-chlorophenyl)-2,3,9-trimethyl-6*H*-thieno[3,2-*f*][1,2,4]triazolo[4,3-*a*][1,4]diazepin-6-yl)acetyl)piperazin-1-yl)-*N*-(2-methyl-3-(trifluoromethyl)benzyl)-4-oxobut-2-enamide (HRG083)**

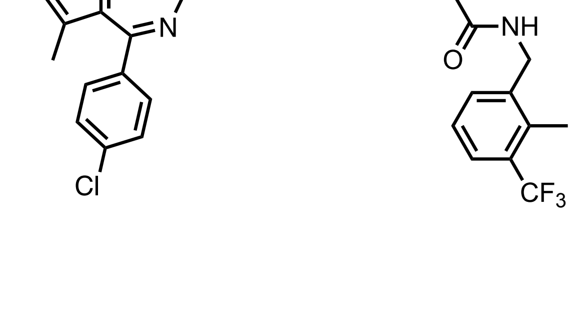

**General Procedure C** was followed with **LO426** (50.0 mg, 0.08 mmol), and TFA (0.20 mL, 2.57 mmol) for 2 hours. The crude material was used without further purification.
**General Procedure A** was followed with (2-methyl-3-(trifluoromethyl)phenyl)methanamine (12.6 mg, 0.07 mmol), HATU (30.5 mg, 0.08 mmol), DIPEA (0.05 mL, 0.27 mmol), and the acid from above (45.5 mg, 0.08 mmol). The crude residue was purified by silica gel chromatography (0-10% MeOH in DCM) to afford 17.2 mg (35%) of the title compound as a yellow-white powder.
**$^1$H NMR** (500 MHz, $CDCl_3$) δ 7.59 (d, *J* = 7.8 Hz, 1H), 7.48 – 7.37 (m, 4H), 7.33 (d, *J* = 8.8 Hz, 2H), 7.26 (t, *J* = 7.7 Hz, 2H), 7.12 (t, *J* = 14.9 Hz, 1H), 4.79 (td, *J* = 6.8, 2.9 Hz, 1H), 4.60 (d, *J* = 5.2 Hz, 2H), 4.04 – 3.32 (m, 10H), 2.63 (s, 3H), 2.41 (d, *J* = 9.0 Hz, 6H), 1.68 (s, 3H).
**$^{13}$C NMR** (126 MHz, $CDCl_3$) δ 169.48, 169.36, 164.42, 164.33, 164.19, 164.07, 163.96, 162.63, 155.81, 155.75, 150.00, 137.74, 136.86, 136.82, 135.62, 135.21, 135.07, 132.38, 132.32, 132.28, 131.02, 130.88, 130.58, 129.97, 129.89, 129.74, 129.59, 128.83, 125.99, 125.66, 125.58, 125.53, 123.48, 54.65, 54.52, 53.55, 46.06, 45.93, 45.78, 45.46, 42.22, 42.13, 41.97, 41.90, 41.36, 38.70, 36.59, 35.36, 31.53, 29.79, 14.90, 14.88, 14.86, 14.48, 13.19, 11.88.
**HRMS (ESI)** m/z calcd for $C_{36}H_{36}ClF_3N_7O_3S^+$ [M+H]$^+$: 738.2235; found: 738.2161.

**(*S*,*E*)-3-(4-(benzyloxy)phenyl)-1-(4-(2-(4-(4-chlorophenyl)-2,3,9-trimethyl-6*H*-thieno[3,2-*f*][1,2,4]triazolo[4,3-*a*][1,4]diazepin-6-yl)acetyl)piperazin-1-yl)prop-2-en-1-one (HRG049)**

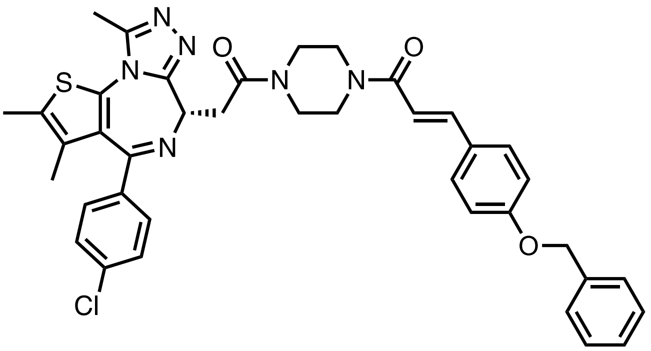


**General Procedure B** was followed with **HY157** (50.0 mg, 0.09 mmol), and TFA (0.22 mL, 2.81 mmol) for 2 hours. The crude material was used without further purification.
**General Procedure A** was followed with 3-(4-(benzyloxy)phenyl)acrylic acid (26.8 mg, 0.11 mmol), HATU (40.0 mg, 0.11 mmol), DIPEA (0.06 mL, 0.35 mmol), and the amine from above (41.2 mg, 0.09 mmol). The crude residue was purified by silica gel chromatography (0-10% MeOH in DCM) to afford 18.0 mg (29%) of the title compound as a yellow powder.
**$^1$H NMR** (500 MHz, CDCl3) δ 7.67 (d, *J* = 15.3 Hz, 1H), 7.49 (d, *J* = 8.4 Hz, 2H), 7.44 – 7.35 (m, 6H), 7.35 – 7.30 (m, 3H), 6.97 (d, *J* = 8.4 Hz, 2H), 6.75 (d, *J* = 15.3 Hz, 1H), 5.09 (s, 2H), 4.83 (t, *J* = 6.5 Hz, 1H), 3.93 – 3.53 (m, 10H), 2.69 (s, 3H), 2.40 (s, 3H), 1.67 (s, 3H).
**$^{13}$C NMR** (126 MHz, $CDCl_3$) δ 166.01, 160.21, 155.64, 149.98, 143.23, 137.00, 136.54, 136.39, 132.23, 131.05, 130.46, 129.94, 129.51, 128.79, 128.67, 128.14, 128.04, 127.49, 115.19, 114.17, 70.10, 54.39, 41.87, 35.23, 14.42, 13.14, 11.86.
**HRMS (ESI)** m/z calcd for $C_{39}H_{38}ClN_6O_3S^+$ [M+H]$^+$: 705.2409; found: 705.2334

**(*S*,*E*)-1-(4-(2-(4-(4-chlorophenyl)-2,3,9-trimethyl-6*H*-thieno[3,2-*f*][1,2,4]triazolo[4,3-*a*][1,4]diazepin-6-yl)acetyl)piperazin-1-yl)-3-(1-methyl-1*H*-pyrazol-4-yl)prop-2-en-1-one (HRG051)**

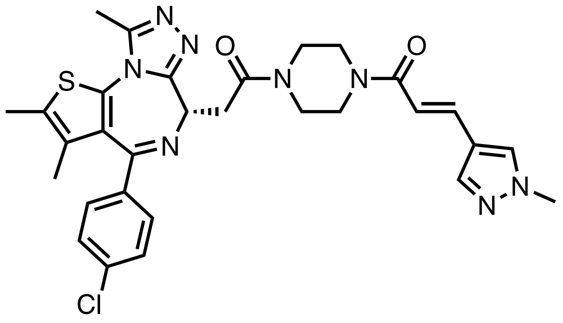

**General Procedure B** was followed with **HY157** (50.0 mg, 0.09 mmol), and TFA (0.22 mL, 2.81 mmol) for 2 hours. The crude material was used without further purification.

**General Procedure A** was followed with (*E*)-3-(1-methyl-1*H*-pyrazol-4-yl)acrylic acid (16.0 mg, 0.11 mmol), HATU (40.0 mg, 0.11 mmol), DIPEA (0.06 mL, 0.35 mmol), and the amine from above (41.2 mg, 0.09 mmol). The crude residue was purified by silica gel chromatography (0-10% MeOH in DCM) to afford 17.2 mg (32%) of the title compound as a yellow-white powder.

**$^1$H NMR** (500 MHz, $CDCl_3$) δ 7.70 (s, 1H), 7.63 – 7.51 (m, 2H), 7.38 (d, *J* = 8.5 Hz, 2H), 7.31 (d, *J* = 8.7 Hz, 2H), 6.60 (d, *J* = 15.2 Hz, 1H), 4.79 (t, *J* = 6.8 Hz, 1H), 3.99 – 3.52 (m, 13H), 2.65 (s, 3H), 2.38 (s, 3H), 1.66 (s, 3H).

**$^{13}$C NMR** (126 MHz, $CDCl_3$) δ 169.30, 165.93, 163.85, 155.73, 149.90, 138.27, 136.75, 136.70, 134.02, 132.20, 130.90, 130.74, 130.46, 130.34, 129.79, 128.72, 118.99, 113.97, 54.48, 53.49, 41.81, 39.13, 35.32, 29.68, 14.38, 13.10, 11.84.

**HRMS (ESI)** m/z calcd for $C_{30}H_{32}ClN_8O_2S^+$ [M+H]$^+$: 603.2052; found: 603.1990

**(*S*)-1-(4-(2-(4-(4-chlorophenyl)-2,3,9-trimethyl-6*H*-thieno[3,2-*f*][1,2,4]triazolo[4,3-*a*][1,4]diazepin-6-yl)acetyl)piperazin-1-yl)-3-(4-(dimethylamino)phenyl)prop-2-en-1-one (HRG052)**

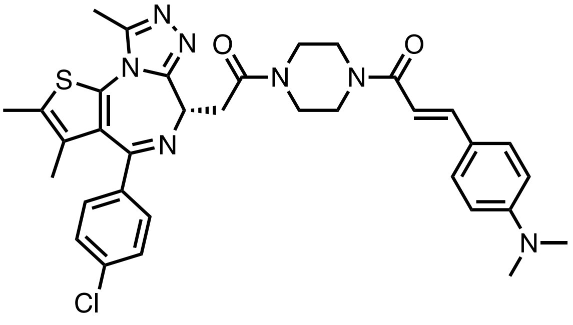

**General Procedure B** was followed with **HY157** (50.0 mg, 0.09 mmol), and TFA (0.22 mL, 2.81 mmol) for 2 hours. The crude material was used without further purification.

**General Procedure A** was followed with 3-(4-(dimethylamino)phenyl)acrylic acid (20.2 mg, 0.11 mmol), HATU (40.0 mg, 0.11 mmol), DIPEA (0.06 mL, 0.35 mmol), and the amine from above (41.2 mg, 0.09 mmol). The crude residue was purified by silica gel chromatography (0-10% MeOH in DCM) to afford 19.3 mg (34%, - $E{:}Z = 11{:}89$) of the title compound as a yellow-white powder.

**$^1$H NMR** (500 MHz, $CDCl_3$) δ 7.68 (d, *J* = 15.2 Hz, 0.1H, minor isomer), 7.45 – 7.34 (m, 2H), 7.31 (dd, *J* = 8.5, 3.8 Hz, 2H), 7.25 (dd, *J* = 8.7, 4.4 Hz, 2H), 6.69 – 6.58 (m, 3H), 5.79 (dd, *J* = 12.4, 5.9 Hz, 0.9H, major isomer), 4.78 (dt, *J* = 26.7, 6.4 Hz, 1H), 3.88 – 3.21 (m, 10H), 2.97 (s, 6H), 2.65 (d, *J* = 8.3 Hz, 3H), 2.39 (d, *J* = 5.7 Hz, 3H), 1.66 (d, *J* = 9.1 Hz, 3H).

**$^{13}$C NMR** (126 MHz, $CDCl_3$) δ 169.02, 168.72, 168.69, 166.57, 163.78, 155.77, 150.55, 149.88, 136.76, 136.69, 134.62, 134.58, 132.20, 130.93, 130.89, 130.73, 130.50, 129.80, 129.76, 129.49, 128.70, 123.22, 121.68, 117.98, 117.81, 111.88, 111.80, 110.88, 54.30, 53.46, 46.34, 46.05, 45.63, 45.31, 41.71, 41.25, 40.99, 40.22, 35.37, 35.28, 29.70, 14.37, 13.09, 11.84.

**HRMS (ESI)** m/z calcd for $C_{34}H_{37}ClN_7O_2S^+$ [M+H]$^+$: 642.2412; found: 642.2361

**(S)-1-(4-(2-(4-(4-chlorophenyl)-2,3,9-trimethyl-6H-thieno[3,2-f][1,2,4]triazolo[4,3-a][1,4]diazepin-6-yl)acetyl)piperazin-1-yl)-3-(naphthalen-1-yl)prop-2-en-1-one (HRG054)**

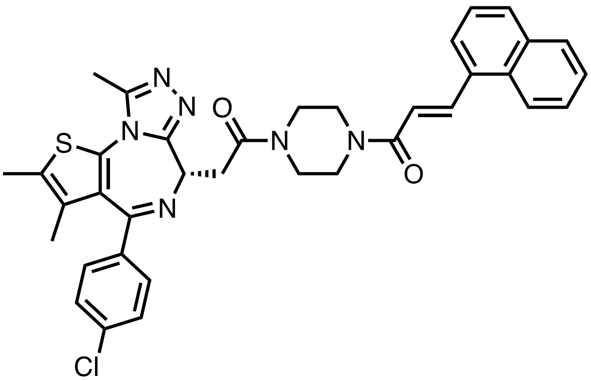


**General Procedure B** was followed with **HY157** (50.0 mg, 0.09 mmol), and TFA (0.22 mL, 2.81 mmol) for 2 hours. The crude material was used without further purification.
**General Procedure A** was followed with 3-(naphthalen-1-yl)acrylic acid (18.2 mg, 0.11 mmol), HATU (40.0 mg, 0.11 mmol), DIPEA (0.06 mL, 0.35 mmol), and the amine from above (41.2 mg, 0.09 mmol). The crude residue was purified by silica gel chromatography (0-10% MeOH in DCM) to afford 13.3 mg (24%, - $E{:}Z = 14{:}86$) of the title compound as a yellow-white powder.
**$^1$H NMR** (500 MHz, $CDCl_3$) δ 8.56 (d, *J* = 15.1 Hz, 0.9H, major isomer), 8.23 (d, *J* = 8.5 Hz, 0.9H, major isomer), 7.97 (dd, *J* = 12.3, 8.4 Hz, 0.1H, minor isomer), 7.93 – 7.84 (m, 2H), 7.74 (d, *J* = 7.2 Hz, 1H), 7.61 – 7.46 (m, 3H), 7.41 (d, *J* = 8.4 Hz, 2H), 7.37 – 7.27 (m, 2H), 6.96 (d, *J* = 15.2 Hz, 0.9H, major isomer), 6.30 (dd, *J* = 12.1, 3.5 Hz, 0.1H, minor isomer), 4.82 (t, *J* = 6.8 Hz, 0.9H, major isomer), 4.66 (t, *J* = 6.7 Hz, 0.1H, minor isomer), 4.08 – 3.08 (m, 10H), 2.66 (d, *J* = 20.4 Hz, 3H), 2.39 (d, *J* = 17.2 Hz, 3H), 1.68 (d, *J* = 9.0 Hz, 3H).
**$^{13}$C NMR** (126 MHz, $CDCl_3$) δ 165.58, 155.76, 149.92, 140.91, 136.78, 136.75, 133.67, 132.83, 132.25, 131.49, 130.94, 130.74, 130.49, 130.07, 129.80, 128.75, 128.64, 126.79, 126.24, 125.42, 124.69, 123.72, 119.62, 41.93, 35.36, 14.40, 13.11, 11.87.
**HRMS (ESI)** m/z calcd for $C_{36}H_{34}ClN_6O_2S^+$ $[M+H]^+$: 649.2147; found: 649.2243

**(*S*)-1-(4-(2-(4-(4-chlorophenyl)-2,3,9-trimethyl-6*H*-thieno[3,2-*f*][1,2,4]triazolo[4,3-*a*][1,4]diazepin-6-yl)acetyl)piperazin-1-yl)-3-(1*H*-indol-3-yl)prop-2-en-1-one (HRG060)**

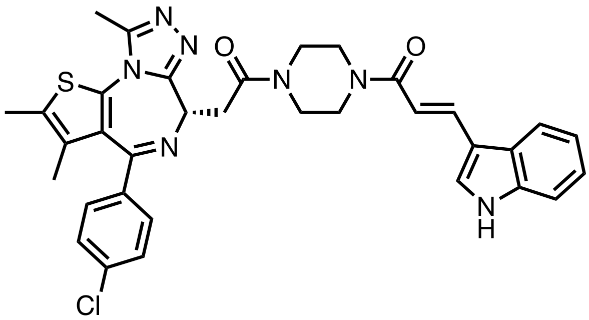


**General Procedure B** was followed with **HY157** (50.0 mg, 0.09 mmol), and TFA (0.22 mL, 2.81 mmol) for 2 hours. The crude material was used without further purification.
**General Procedure A** was followed with 3-(1*H*-indol-3-yl)acrylic acid (19.7 mg, 0.11 mmol), HATU (40.0 mg, 0.11 mmol), DIPEA (0.06 mL, 0.35 mmol), and the amine from above (41.2 mg, 0.09 mmol). The crude residue was purified by silica gel chromatography (0-10% MeOH in DCM) to afford 16.8 mg (30%, - $E{:}Z = 79{:}21$) of the title compound as a yellow powder.
**$^1$H NMR** (500 MHz, CDCl3) δ 10.39 (s, 1H), 9.35 (d, *J* = 18.0 Hz, 1H), 8.14 – 7.78 (m, 2H), 7.70 – 7.27 (m, 7H), 7.23 – 6.97 (m, 1H), 6.82 (d, *J* = 15.2 Hz, 0.8H, major isomer), 5.95 (d, *J* = 12.3 Hz, 0.2H, minor

isomer), 4.80 (dt, *J* = 30.2, 6.9 Hz, 1H), 3.54 (t, *J* = 114.7 Hz, 10H), 2.69 (d, *J* = 14.1 Hz, 3H), 2.40 (d, *J* = 10.6 Hz, 3H), 1.67 (d, *J* = 13.3 Hz, 3H).
**$^{13}$C NMR** (126 MHz, $CDCl_3$) δ 163.91, 149.98, 136.80, 136.69, 130.89, 129.85, 129.79, 128.77, 128.74, 126.11, 125.27, 123.18, 121.21, 120.30, 113.77, 54.28, 45.44, 41.73, 35.32, 14.39, 14.36, 13.12, 13.10, 11.87, 11.84.
**HRMS (ESI)** m/z calcd for $C_{34}H_{33}ClN_7O_2S^+$ [M+H]$^+$: 638.2099; found: 638.2048

**(*S*,*E*)-1-(4-(2-(4-(4-chlorophenyl)-2,3,9-trimethyl-6*H*-thieno[3,2-*f*][1,2,4]triazolo[4,3-*a*][1,4]diazepin-6-yl)acetyl)piperazin-1-yl)-3-(pyridin-3-yl)prop-2-en-1-one (HRG061)**

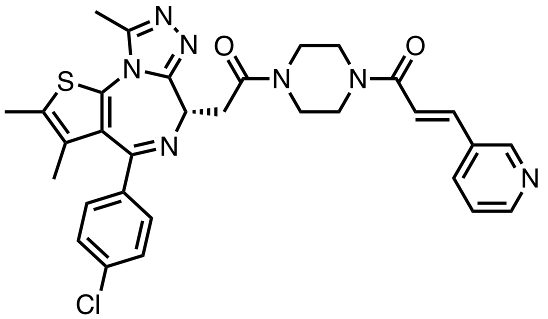

**General Procedure B** was followed with **HY157** (50.0 mg, 0.09 mmol), and TFA (0.22 mL, 2.81 mmol) for 2 hours. The crude material was used without further purification.
**General Procedure A** was followed with (*E*)-3-(pyridin-3-yl)acrylic acid (15.7 mg, 0.11 mmol), HATU (40.0 mg, 0.11 mmol), DIPEA (0.06 mL, 0.35 mmol), and the amine from above (41.2 mg, 0.09 mmol). The crude residue was purified by silica gel chromatography (0-10% MeOH in DCM) to afford 17.7 mg (34%) of the title compound as a yellow-white powder.
**$^1$H NMR** (500 MHz, $CDCl_3$) δ 8.78 (d, *J* = 2.2 Hz, 1H), 8.60 (dd, *J* = 4.8, 1.6 Hz, 1H), 7.86 (d, *J* = 7.8 Hz, 1H), 7.72 (d, *J* = 15.5 Hz, 1H), 7.46 – 7.31 (m, 5H), 6.98 (d, *J* = 15.5 Hz, 1H), 4.82 (t, *J* = 6.8 Hz, 1H), 4.09 – 3.51 (m, 10H), 2.68 (s, 3H), 2.41 (s, 3H), 1.69 (s, 3H).
**$^{13}$C NMR** (126 MHz, $CDCl_3$) δ 169.42, 165.00, 163.91, 155.71, 150.57, 149.92, 149.41, 139.85, 136.75, 136.74, 134.25, 132.23, 130.91, 130.83, 130.76, 130.45, 129.79, 128.73, 123.69, 118.81, 54.45, 46.15, 45.58, 41.91, 35.32, 14.39, 13.10, 11.85.
**HRMS (ESI)** m/z calcd for $C_{31}H_{31}ClN_7O_2S^+$ [M+H]$^+$: 600.1943; found: 600.2066

**ethyl (*S*,*E*)-4,6-dichloro-3-(3-(4-(2-(4-(4-chlorophenyl)-2,3,9-trimethyl-6*H*-thieno[3,2-*f*][1,2,4]triazolo[4,3-*a*][1,4]diazepin-6-yl)acetyl)piperazin-1-yl)-3-oxoprop-1-en-1-yl)-1*H*-indole-2-carboxylate (HRG064)**

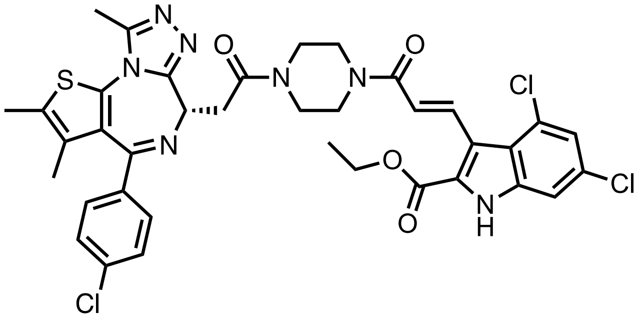

**General Procedure B** was followed with **HY157** (50.0 mg, 0.09 mmol), and TFA (0.22 mL, 2.81 mmol) for 2 hours. The crude material was used without further purification.
**General Procedure A** was followed with (*E*)-3-(4,6-dichloro-2-(ethoxycarbonyl)-1*H*-indol-3-yl)acrylic acid (34.6 mg, 0.11 mmol), HATU (40.0 mg, 0.11 mmol), DIPEA (0.06 mL, 0.35 mmol), and the amine from

above (41.2 mg, 0.09 mmol). The crude residue was purified by silica gel chromatography (0-10% MeOH in DCM) to afford 19.1 mg (28%) of the title compound as a yellow-white powder.
**$^1$H NMR** (500 MHz, $CDCl_3$) δ 10.18 (d, *J* = 57.5 Hz, 1H), 8.43 (d, *J* = 15.4 Hz, 1H), 7.44 – 7.33 (m, 5H), 7.20 – 7.11 (m, 2H), 4.85 (t, *J* = 6.7 Hz, 1H), 4.42 (q, *J* = 7.1 Hz, 2H), 4.04 – 3.57 (m, 11H), 2.69 (s, 3H), 2.41 (s, 3H), 1.67 (s, 3H), 1.41 (t, *J* = 7.1 Hz, 3H).
**$^{13}$C NMR** (126 MHz, $CDCl_3$) δ 166.05, 163.89, 160.84, 155.77, 149.95, 137.03, 136.76, 136.71, 133.36, 132.11, 131.20, 130.92, 130.52, 129.81, 128.73, 128.62, 125.98, 123.65, 123.35, 122.57, 118.94, 110.98, 61.69, 54.43, 53.46, 35.40, 29.70, 14.35, 13.11, 11.82.
**HRMS (ESI)** m/z calcd for $C_{37}H_{35}Cl_3N_7O_4S^+$ [M+H]$^+$: 778.1531; found: 778.1594

**(*S*,*E*)-1-(4-(2-(4-(4-chlorophenyl)-2,3,9-trimethyl-6*H*-thieno[3,2-*f*][1,2,4]triazolo[4,3-*a*][1,4]diazepin-6-yl)acetyl)piperazin-1-yl)-3-(3-(4-methoxyphenyl)-1-phenyl-1*H*-pyrazol-4-yl)prop-2-en-1-one (HRG069)**

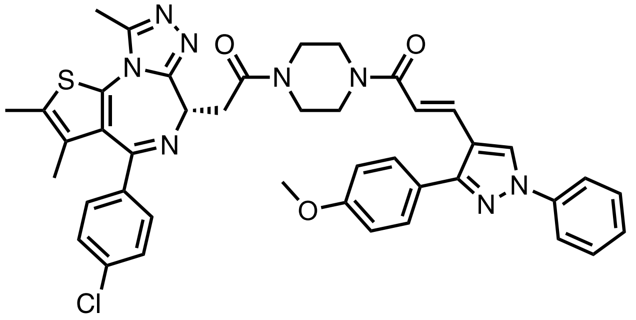

**General Procedure B** was followed with **HY157** (50.0 mg, 0.09 mmol), and TFA (0.22 mL, 2.81 mmol) for 2 hours. The crude material was used without further purification.
**General Procedure A** was followed with (*E*)-3-(3-(4-methoxyphenyl)-1-phenyl-1*H*-pyrazol-4-yl)acrylic acid (33.8 mg, 0.11 mmol), HATU (40.0 mg, 0.11 mmol), DIPEA (0.06 mL, 0.35 mmol), and the amine from above (41.2 mg, 0.09 mmol). The crude residue was purified by silica gel chromatography (0-10% MeOH in DCM) to afford 21.2 mg (31%) of the title compound as a yellow-white powder.
**$^1$H NMR** (500 MHz, $CDCl_3$) δ 8.25 (d, *J* = 5.6 Hz, 1H), 7.79 (dd, *J* = 11.6, 3.7 Hz, 3H), 7.70 – 7.58 (m, 2H), 7.54 – 7.46 (m, 2H), 7.44 – 7.38 (m, 2H), 7.37 – 7.31 (m, 3H), 7.07 – 6.98 (m, 2H), 6.71 (d, *J* = 15.3 Hz, 1H), 4.83 (t, *J* = 6.8 Hz, 1H), 4.03 – 3.50 (m, 13H), 2.69 (s, 3H), 2.42 (s, 3H), 1.70 (s, 3H).
**$^{13}$C NMR** (126 MHz, $CDCl_3$) δ 165.76, 163.90, 160.00, 155.75, 152.91, 149.94, 139.56, 136.76, 134.23, 132.20, 130.94, 130.78, 130.51, 129.99, 129.93, 129.81, 129.52, 129.46, 128.75, 126.95, 126.36, 125.00, 119.29, 119.22, 118.04, 115.77, 114.24, 114.21, 55.40, 54.49, 41.83, 35.34, 30.94, 14.39, 13.11, 11.86.
**HRMS (ESI)** m/z calcd for $C_{42}H_{40}ClN_8O_3S^+$ [M+H]$^+$: 771.2627; found: 771.2701

**(*S*,*E*)-1-(4-(3-(4-(2-(4-(4-chlorophenyl)-2,3,9-trimethyl-6*H*-thieno[3,2-*f*][1,2,4]triazolo[4,3-*a*][1,4]diazepin-6-yl)acetyl)piperazin-1-yl)-3-oxoprop-1-en-1-yl)phenyl)pyrrolidin-2-one (HRG070)**

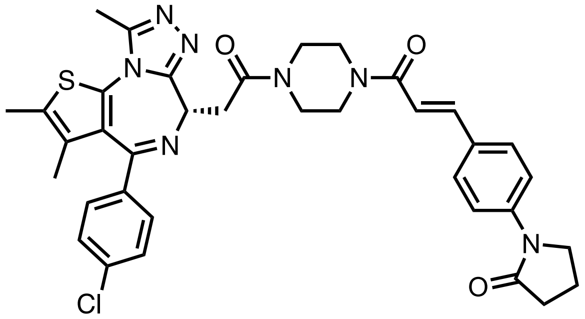

**General Procedure B** was followed with **HY157** (50.0 mg, 0.09 mmol), and TFA (0.22 mL, 2.81 mmol) for 2 hours. The crude material was used without further purification.
**General Procedure A** was followed with (*E*)-3-(4-(2-oxopyrrolidin-1-yl)phenyl)acrylic acid (24.4 mg, 0.11 mmol), HATU (40.0 mg, 0.11 mmol), DIPEA (0.06 mL, 0.35 mmol), and the amine from above (41.2 mg,

0.09 mmol). The crude residue was purified by silica gel chromatography (0-10% MeOH in DCM) to afford 17.7 mg (30%) of the title compound as a yellow-white powder.

**$^1$H NMR** (500 MHz, $CDCl_3$) δ 7.72 – 7.65 (m, 3H), 7.55 (d, *J* = 9.0 Hz, 2H), 7.41 (d, 2H), 7.33 (d, *J* = 8.6 Hz, 2H), 6.84 (d, *J* = 15.4 Hz, 1H), 4.81 (t, *J* = 6.7 Hz, 1H), 4.05 – 3.53 (m, 12H), 2.67 (s, 3H), 2.63 (t, *J* = 8.1 Hz, 2H), 2.41 (s, 3H), 2.18 (p, *J* = 7.4 Hz, 2H), 1.68 (s, 3H).

**$^{13}$C NMR** (126 MHz, $CDCl_3$) δ 174.44, 169.33, 165.79, 163.88, 155.73, 149.91, 142.86, 140.75, 136.73, 132.20, 130.92, 130.74, 130.48, 129.79, 128.73, 128.50, 119.61, 115.61, 54.47, 48.56, 45.61, 41.86, 35.32, 32.82, 17.91, 14.39, 13.10, 11.85.

**HRMS (ESI)** m/z calcd for $C_{36}H_{37}ClN_7O_3S^+$ $[M+H]^+$: 682.2362; found: 682.2487

**(*S*)-1-(4-(2-(4-(4-chlorophenyl)-2,3,9-trimethyl-6*H*-thieno[3,2-*f*][1,2,4]triazolo[4,3-*a*][1,4]diazepin-6-yl)acetyl)piperazin-1-yl)-3-(4-morpholinophenyl)prop-2-en-1-one (HRG071)**

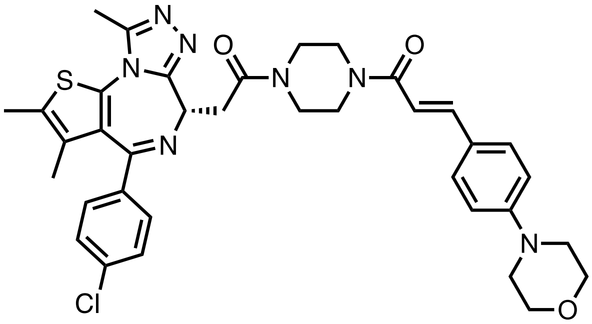

**General Procedure B** was followed with **HY157** (50.0 mg, 0.09 mmol), and TFA (0.22 mL, 2.81 mmol) for 2 hours. The crude material was used without further purification.

**General Procedure A** was followed with 3-(4-morpholinophenyl)acrylic acid (24.6 mg, 0.11 mmol), HATU (40.0 mg, 0.11 mmol), DIPEA (0.06 mL, 0.35 mmol), and the amine from above (41.2 mg, 0.09 mmol). The crude residue was purified by silica gel chromatography (0-10% MeOH in DCM) to afford 16.2 mg (27%, -*E:Z* = 58:42) of the title compound as a yellow-white powder.

**$^1$H NMR** (500 MHz, $CDCl_3$) δ 7.68 (d, *J* = 15.2 Hz, 0.6H, major isomer), 7.50 – 7.27 (m, 6H), 6.92 – 6.79 (m, 2H), 6.73 (d, *J* = 15.3 Hz, 0.6H, major isomer), 6.65 (t, *J* = 11.9 Hz, 0.4H, minor isomer), 5.89 (dd, *J* = 12.4, 7.0 Hz, 0.4H, minor isomer), 4.81 (t, *J* = 6.7 Hz, 0.6H, major isomer), 4.76 (td, *J* = 6.6, 1.4 Hz, 0.4H, minor isomer), 4.06 – 3.27 (m, 14H), 3.26 – 3.18 (m, 4H), 2.70 – 2.64 (m, 3H), 2.40 (dd, *J* = 5.0, 0.8 Hz, 3H), 1.67 (dd, *J* = 8.0, 0.8 Hz, 3H).

**$^{13}$C NMR** (126 MHz, $CDCl_3$) δ 169.05, 168.30, 166.21, 163.84, 155.76, 152.24, 151.27, 149.91, 143.50, 136.74, 134.02, 132.21, 130.94, 130.72, 130.50, 129.79 (d, *J* = 1.8 Hz), 129.65, 129.31, 128.74, 128.72, 126.55, 126.50, 126.22, 119.71, 119.54, 114.84, 112.91, 66.71, 54.29, 48.42, 48.30, 46.28, 46.02, 45.65, 45.30, 41.85, 41.70, 41.24, 41.19, 40.99, 35.34, 35.27, 14.39, 14.38, 13.10, 11.86, 11.85.

**HRMS (ESI)** m/z calcd for $C_{36}H_{38}ClN_7O_3S^+$ $[M+H]^+$: 684.2518; found: 684.2586

***tert-butyl (E)-4-((5-chloro-6-(2H-1,2,3-triazol-2-yl)pyridin-3-yl)amino)-4-oxobut-2-enoate*** **(LO314)**

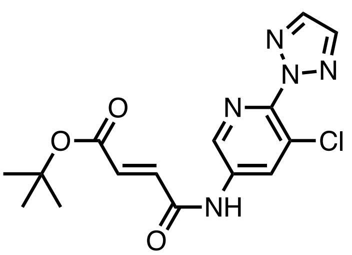

**General Procedure A** was followed with (*E*)-4-(*tert*-butoxy)-4-oxobut-2-enoic acid (105.6 mg, 0.61 mmol), HATU (233.3 mg, 0.51 mmol), DIPEA (0.36 mL, 2.04 mmol), and 5-chloro-6-(2*H*-1,2,3-triazol-2-yl)pyridin-3-

amine (100.0 mg, 0.51 mmol). The crude residue was purified by silica gel chromatography (0-30% EtOAc in hexanes) to afford 88.7 mg (44%) of the title compound as a beige powder.
**$^1$H NMR** (500 MHz, $CDCl_3$) δ 9.12 (s, 1H), 8.63 (d, *J* = 2.3 Hz, 1H), 8.41 (d, *J* = 2.4 Hz, 1H), 7.93 (s, 2H), 6.99 (d, *J* = 15.4 Hz, 1H), 6.88 (d, *J* = 15.3 Hz, 1H), 1.49 (s, 9H).
**$^{13}$C NMR** (126 MHz, $CDCl_3$) δ 169.05, 164.01, 155.68, 154.60, 149.92, 136.94, 136.41, 132.18, 131.03, 130.98, 130.48, 129.93, 128.75, 80.28, 54.31, 45.71, 41.70, 35.20, 28.40, 14.39, 13.11, 11.81.
**HRMS (ESI)** m/z calcd for $C_{15}H_{17}ClN_5O_3^+$ $[M+H]^+$: 350.1014; found: 350.1098

**_tert_-butyl (_E_)-4-(4-((5-chloro-6-(2_H_-1,2,3-triazol-2-yl)pyridin-3-yl)amino)-4-oxobut-2-enoyl)piperazine-1-carboxylate (LO340)**

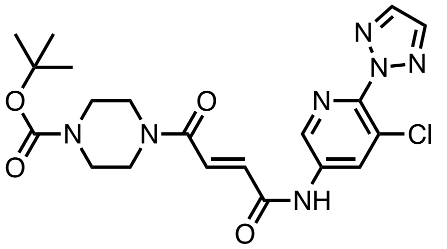

**General Procedure C** was followed with **LO314** (100.0 mg, 0.29 mmol), and TFA (0.7 mL, 9.15 mmol) for 2 hours. The crude material was used without further purification.
**General Procedure A** was followed with *tert*-butyl piperazine-1-carboxylate (44.4 mg, 0.24 mmol), HATU (108.7 mg, 0.29 mmol), DIPEA (0.17 mL, 0.95 mmol), and the acid from above (83.9 mg, 0.29 mmol). The crude residue was purified by silica gel chromatography (80-100% EtOAc in hexanes) to afford 49.2 mg (44.7%) of the title compound as a clear oil.
**$^1$H NMR** (500 MHz, $CDCl_3$) δ 10.10 (s, 1H), 8.72 (d, *J* = 2.3 Hz, 1H), 8.64 (d, *J* = 2.3 Hz, 1H), 7.93 (s, 2H), 7.60 (d, *J* = 14.9 Hz, 1H), 7.36 (d, *J* = 14.8 Hz, 1H), 3.75 (t, *J* = 5.3 Hz, 2H), 3.66 (d, *J* = 5.1 Hz, 2H), 3.56 – 3.49 (m, 4H), 1.48 (s, 9H).
**$^{13}$C NMR** (126 MHz, $CDCl_3$) δ 164.51, 162.78, 154.32, 143.76, 137.94, 136.03, 134.84, 131.52, 130.45, 126.34, 80.85, 46.21, 42.53, 28.36.
**HRMS (ESI)** m/z calcd for $C_{20}H_{25}ClN_7O_4^+$ $[M+H]^+$: 462.1651; found: 462.1641

**(_E_)-_N_-(5-chloro-6-(2_H_-1,2,3-triazol-2-yl)pyridin-3-yl)-4-(4-(hex-5-yn-1-yl)piperazin-1-yl)-4-oxobut-2-enamide (LO344)**

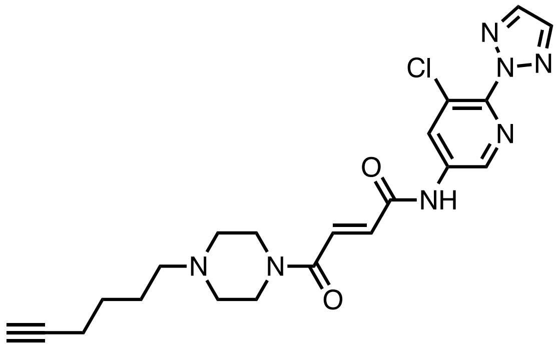

**General Procedure B** was followed with **LO340** (35.0 mg, 0.08 mmol), and TFA (0.22 mL, 2.91 mmol) for 2 hours. The crude material was used without further purification.
A combination of the amine from above (27.4 mg, 0.08 mmol) and potassium carbonate (31.4 mg, 0.23 mmol) was suspended in acetonitrile (3 mL) and stirred at ambient temperature for 10 minutes. 6-bromohex-1-yne (24.4 mL, 0.15 mmol) was added, and the reaction mixture was stirred at 60°C for 2 hours.

The reaction mixture was concentrated *in vacuo*, and the crude residue was purified by silica gel chromatography (0-30% EtOAc in hexanes) to afford 34.1 mg (87%) of the title compound as a yellow oil.
**$^1$H NMR** (500 MHz, $CDCl_3$) δ 10.30 (s, 1H), 8.69 (q, 2H), 7.93 (s, 2H), 7.62 (d, *J* = 14.9 Hz, 1H), 7.40 (d, *J* = 14.9 Hz, 1H), 3.78 (t, 2H), 3.70 (t, 2H), 2.51 (q, *J* = 5.4 Hz, 4H), 2.40 (t, *J* = 7.0 Hz, 2H), 2.23 (td, *J* = 6.8, 2.7 Hz, 2H), 1.96 (t, *J* = 2.6 Hz, 1H), 1.66 – 1.53 (m, 4H).
**$^{13}$C NMR** (126 MHz, $CDCl_3$) δ 164.23, 163.03, 143.68, 138.07, 136.82, 135.99, 134.60, 131.75, 130.42, 126.30, 84.15, 68.63, 57.53, 53.24, 52.51, 46.52, 42.82, 26.14, 25.64, 18.27.
**HRMS (ESI)** m/z calcd for $C_{20}H_{25}ClN_7O_2^+$ [M+H]$^+$: 442.1753; found: 442.1739

**_tert_-butyl (_R_)-4-(2-(4-(4-chlorophenyl)-2,3,9-trimethyl-6_H_-thieno[3,2-_f_][1,2,4]triazolo[4,3-_a_][1,4]diazepin-6-yl)acetyl)piperazine-1-carboxylate (LO359)**

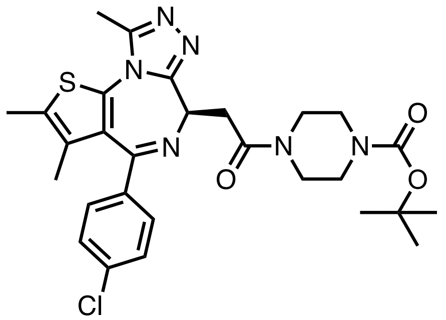

**General Procedure A** was followed with (R)-2-(4-(4-chlorophenyl)-2,3,9-trimethyl-6H-thieno [3,2-f][1,2,4]triazolo[4,3-a][1,4]diazepin-6-yl)acetic acid (R-JQ1-Acid) (129.0 mg, 0.32 mmol), HATU (122.6 mg, 0.32 mmol), DIPEA (0.19 mL, 1.1 mmol), and *tert*-butyl piperazine-1-carboxylate (50.0 mg, 0.27 mmol). The crude residue was purified by silica gel chromatography (0-10% MeOH in DCM) to afford 89.2 mg (58%) of the title compound as a yellow oil.
**$^1$H NMR** (500 MHz, $CDCl_3$) δ 7.40 (d, *J* = 8.5 Hz, 2H), 7.32 (d, *J* = 8.8 Hz, 2H), 4.81 (t, *J* = 6.7 Hz, 1H), 3.84 – 3.35 (m, 10H), 2.68 (s, 3H), 2.39 (s, 3H), 1.67 (s, 3H), 1.47 (s, 9H).
**$^{13}$C NMR** (126 MHz, $CDCl_3$) δ 169.05, 164.01, 155.68, 154.60, 149.92, 136.94, 136.41, 132.18, 131.01 (d, *J* = 6.5 Hz), 130.48, 129.93, 128.75, 80.28, 54.31, 45.71, 41.70, 35.20, 28.40, 14.39, 13.11, 11.81.
**HRMS (ESI)** m/z calcd for $C_{28}H_{34}ClN_6O_3S^+$ [M+H]$^+$: 569.2102; found: 569.2153

**(_R_,_E_)-_N_-(5-chloro-6-(2_H_-1,2,3-triazol-2-yl)pyridin-3-yl)-4-(4-(2-(4-(4-chlorophenyl)-2,3,9-trimethyl-6_H_-thieno[3,2-_f_][1,2,4]triazolo[4,3-_a_][1,4]diazepin-6-yl)acetyl)piperazin-1-yl)-4-oxobut-2-enamide (LO360)**

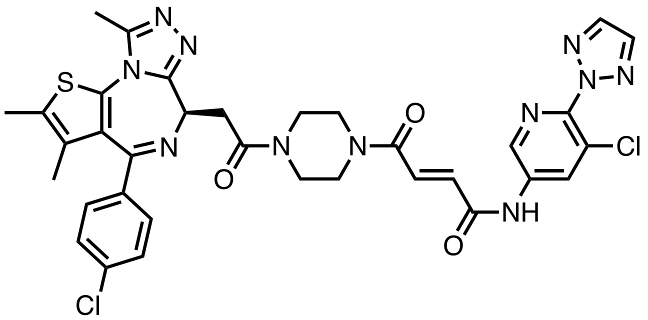

**General Procedure B** was followed with **LO359** (42.0 mg, 0.07 mmol), and TFA (0.18 mL, 2.36 mmol) for 2 hours. The crude material was used without further purification.
**General Procedure C** was followed with **LO314** (35.0 mg, 0.09 mmol), and TFA (0.22 mL, 2.84 mmol) for 2 hours. The crude material was used without further purification.
**General Procedure A** was followed with the amine from above (34.7 mg, 0.07 mmol), HATU (33.7 mg, 0.07 mmol), DIPEA (0.05 mL, 0.30 mmol), and the acid from above (26.0 mg, 0.09 mmol). The crude

residue was purified by silica gel chromatography (0-10% MeOH in DCM) to afford 28.2 mg (51%) of the title compound as a beige powder.

**$^{1}$H NMR** (500 MHz, $CDCl_3$) δ 10.33 (s, 1H), 8.77 – 8.62 (m, 2H), 7.92 (d, *J* = 2.7 Hz, 2H), 7.56 (dd, *J* = 14.9, 7.1 Hz, 1H), 7.40 (d, *J* = 8.4 Hz, 2H), 7.32 (d, *J* = 6.6 Hz, 2H), 7.26 – 7.21 (m, 1H), 4.81 (t, *J* = 6.8 Hz, 1H), 4.13 – 3.41 (m, 10H), 2.67 (d, *J* = 2.5 Hz, 3H), 2.39 (d, *J* = 4.7 Hz, 3H), 1.67 (d, *J* = 3.5 Hz, 3H).

**$^{13}$C NMR** (126 MHz, $CDCl_3$) δ 169.53, 169.42, 164.50, 164.15, 163.95, 163.08, 155.82, 150.06, 143.64, 138.11, 137.98, 136.82, 136.59, 135.97, 135.01, 134.34, 131.96, 131.57, 130.66, 130.53, 130.42, 130.30, 129.84, 128.73, 126.30, 54.62, 54.32, 45.86, 45.37, 42.39, 42.07, 41.83, 41.23, 29.71, 14.40, 13.12, 11.85.

**HRMS (ESI)** m/z calcd for $C_{34}H_{32}Cl_2N_{11}O_3S$ $[M+H]^+$: 744.1782; found: 744.1818

**(*E*)-2-((5-(4-(4-((5-chloro-6-(2*H*-1,2,3-triazol-2-yl)pyridin-3-yl)amino)-4-oxobut-2-enoyl)piperazin-1-yl)pyridin-2-yl)amino)-7-cyclopentyl-*N*,*N*-dimethyl-7*H*-pyrrolo[2,3-*d*]pyrimidine-6-carboxamide (LO320)**

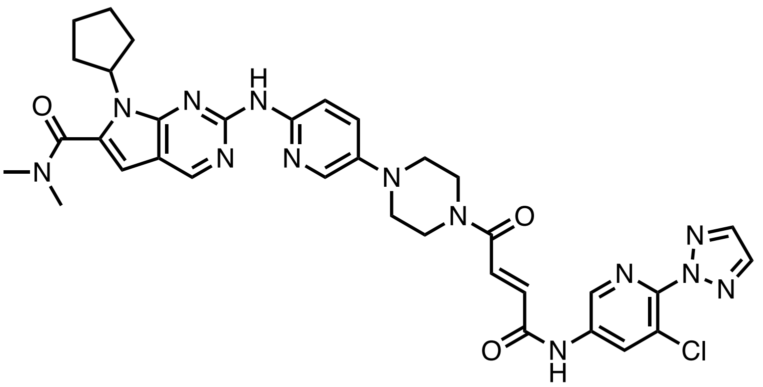

**General Procedure C** was followed with **LO314** (50.0 mg, 0.14 mmol), and TFA (0.35 mL, 4.57 mmol) for 2 hours. The crude material was used without further purification.

**General Procedure A** was followed with 7-cyclopentyl-N,N-dimethyl-2-((5-(piperazin-1-yl)pyridin-2-yl)amino)-7H-pyrrolo[2,3-d]pyrimidine-6-carboxamide (ribociclib) (51.8 mg, 0.12 mmol), HATU (54.4 mg, 0.14 mmol), DIPEA (0.08 mL, 0.48 mmol), and the acid from above (42.0 mg, 0.14 mmol). The crude residue was purified by silica gel chromatography (0-10% MeOH in DCM) to afford 29.1 mg (34%) of the title compound as a yellow-white powder.

**$^{1}$H NMR** (500 MHz, DMSO) δ 11.14 (s, 1H), 9.43 (s, 1H), 8.72 (s, 1H), 8.68 (d, *J* = 2.3 Hz, 1H), 8.56 (d, *J* = 2.2 Hz, 1H), 8.14 (d, *J* = 9.1 Hz, 1H), 8.01 (d, *J* = 3.0 Hz, 1H), 7.53 (d, *J* = 15.0 Hz, 1H), 7.44 (dd, *J* = 9.2, 3.0 Hz, 1H), 7.03 (d, *J* = 15.0 Hz, 1H), 6.54 (s, 1H), 4.68 (p, *J* = 8.9 Hz, 1H), 3.72 (dt, *J* = 20.7, 5.1 Hz, 4H), 3.12 (dt, *J* = 14.2, 5.5 Hz, 4H), 3.05 – 2.93 (m, 6H), 2.42 – 2.31 (m, 2H), 1.92 (dq, *J* = 12.5, 6.0 Hz, 4H), 1.58 (q, *J* = 6.5 Hz, 2H).

**$^{13}$C NMR** (126 MHz, DMSO) δ 163.81, 163.36, 163.31, 155.16, 152.58, 151.64, 147.13, 143.23, 142.36, 138.43, 138.20, 136.77 (d, *J* = 6.4 Hz), 134.03, 132.36, 132.24, 129.36, 126.53 (d, *J* = 7.4 Hz), 113.02, 112.32, 101.08, 57.42, 50.14, 49.37, 45.75, 42.01, 35.08, 30.19, 24.68.

**HRMS (ESI)** m/z calcd for $C_{34}H_{37}ClN_{13}O_3^+$ $[M+H]^+$: 710.2825; found: 710.2830

**(*E*)-4-(4-(3-amino-6-(2-hydroxyphenyl)pyridazin-4-yl)piperazin-1-yl)-*N*-(5-chloro-6-(2*H*-1,2,3-triazol-2-yl)pyridin-3-yl)-4-oxobut-2-enamide (LO325)**

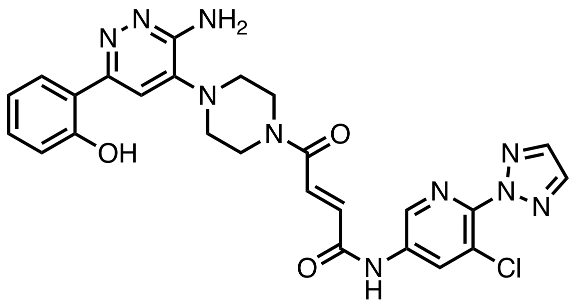

**General Procedure C** was followed with **LO314** (50.0 mg, 0.14 mmol), and TFA (0.35 mL, 4.57 mmol) for 2 hours. The crude material was used without further purification.
**General Procedure A** was followed with 2-(6-amino-5-(piperazin-1-yl)pyridazin-3-yl)phenol (32.3 mg, 0.12 mmol), HATU (54.4 mg, 0.14 mmol), DIPEA (0.08 mL, 0.48 mmol), and the acid from above (42.0 mg, 0.14 mmol). The crude residue was purified by silica gel chromatography (0-10% MeOH in DCM) to afford 22.1 mg (34%) of the title compound as a yellow-white powder.
**1H NMR** (500 MHz, DMSO) δ 14.17 (s, 1H), 11.06 (s, 1H), 8.67 (d, *J* = 2.3 Hz, 1H), 8.62 (d, *J* = 2.3 Hz, 1H), 8.17 (s, 2H), 7.93 (dd, *J* = 8.0, 1.6 Hz, 1H), 7.55 (s, 1H), 7.24 (ddd, *J* = 8.5, 7.3, 1.6 Hz, 1H), 6.88 (t, *J* = 7.7 Hz, 2H), 6.79 (d, *J* = 11.8 Hz, 1H), 6.42 (s, 2H), 6.34 (d, *J* = 11.8 Hz, 1H), 3.78 (t, 2H), 3.62 (t, 2H), 3.15 (t, *J* = 5.1 Hz, 2H), 3.10 (t, *J* = 5.0 Hz, 2H).
**$^{13}$C NMR** (126 MHz, DMSO) δ 166.25, 163.74, 158.96, 155.25, 153.60, 143.08, 140.56, 138.28, 138.21, 137.82, 136.76, 130.60, 129.10, 126.66, 126.51, 125.31, 118.95, 118.25, 117.84, 111.38, 48.92, 48.66, 45.56.
**HRMS (ESI)** m/z calcd for $C_{25}H_{24}ClN_{10}O_3^+$ [M+H]$^+$: 547.1716; found: 547.1712

**(*E*)-*N*-(4-(3-chloro-4-cyanophenoxy)cyclohexyl)-6-(4-(4-((5-chloro-6-(2*H*-1,2,3-triazol-2-yl)pyridin-3-yl)amino)-4-oxobut-2-enoyl)piperazin-1-yl)pyridazine-3-carboxamide (LO348)**

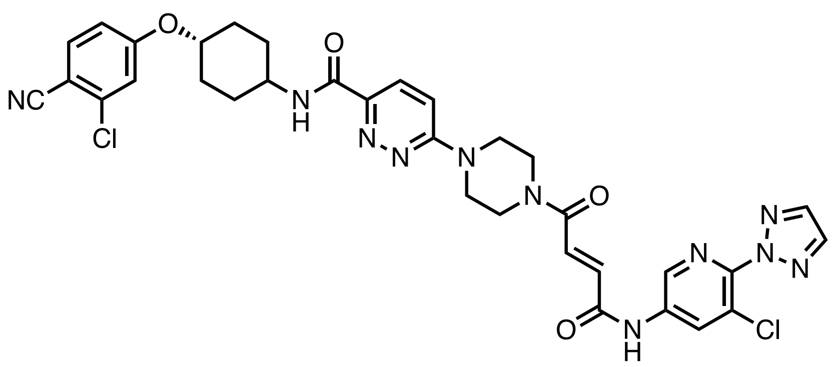

**General Procedure C** was followed with **LO314** (50.0 mg, 0.14 mmol), and TFA (0.35 mL, 4.57 mmol) for 2 hours. The crude material was used without further purification.
**General Procedure A** was followed with *N*-(4-(3-chloro-4-cyanophenoxy)cyclohexyl)-6-(piperazin-1-yl)pyridazine-3-carboxamide (52.5 mg, 0.12 mmol), HATU (54.4 mg, 0.14 mmol), DIPEA (0.08 mL, 0.48 mmol), and the acid from above (42.0 mg, 0.14 mmol). The crude residue was purified by silica gel chromatography (0-10% MeOH in DCM) to afford 50.8 mg (60%) of the title compound as a white powder.
**$^{1}$H NMR** (500 MHz, DMSO) δ 11.22 (s, 1H), 8.75 (d, *J* = 2.2 Hz, 1H), 8.65 (d, *J* = 8.2 Hz, 1H), 8.62 (d, *J* = 2.2 Hz, 1H), 8.18 (s, 2H), 7.91 – 7.88 (m, 1H), 7.86 (d, *J* = 8.8 Hz, 1H), 7.60 (d, *J* = 15.0 Hz, 1H), 7.42 – 7.35 (m, 3H), 7.14 (dd, *J* = 8.8, 2.4 Hz, 1H), 7.10 (d, *J* = 15.0 Hz, 1H), 4.54 (dt, *J* = 10.5, 5.7 Hz, 1H), 3.90 – 3.74 (m, 10H), 2.13 – 2.08 (m, 2H), 1.91 (d, *J* = 12.2 Hz, 2H), 1.65 (q, *J* = 12.4 Hz, 2H), 1.52 (q, *J* = 12.0 Hz, 2H).
**$^{13}$C NMR** (126 MHz, DMSO) δ 164.18, 163.79, 163.53, 162.89, 162.27, 160.40, 145.57, 143.24, 138.43, 138.19, 137.49, 136.81, 136.23, 134.10, 132.23, 131.64, 129.35, 126.96, 126.50, 117.28, 116.90, 115.93, 113.40, 103.68, 76.05, 47.59, 45.18, 44.95, 44.38, 41.65, 30.34, 29.89.

**HRMS (ESI)** m/z calcd for $C_{33}H_{31}Cl_2N_{11}NaO_4^+$ [M+H]$^+$: 738.1830; found: 738.1772

**(*S*,*E*)-4-(4-(2-(4-(4-chlorophenyl)-2,3,9-trimethyl-6*H*-thieno[3,2-*f*][1,2,4]triazolo[4,3-*a*][1,4]diazepin-6-yl)acetyl)piperazin-1-yl)-*N*-(5-chloropyridin-3-yl)-4-oxobut-2-enamide (LO361)**

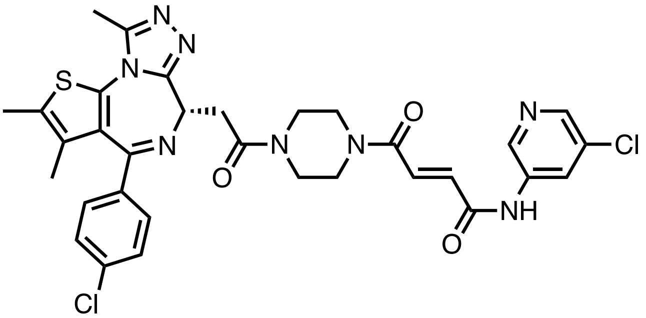

**General Procedure C** was followed with **LO426** (50.0 mg, 0.08 mmol), and TFA (0.20 mL, 2.57 mmol) for 2 hours. The crude material was used without further purification.

**General Procedure A** was followed with 5-chloropyridin-3-amine (8.6 mg, 0.07 mmol), HATU (30.5 mg, 0.08 mmol), DIPEA (0.05 mL, 0.27 mmol), and the acid from above (45.5 mg, 0.08 mmol). The crude residue was purified by silica gel chromatography (0-10% MeOH in DCM) to afford 11.9 mg (26%) of the title compound as a beige powder.

**$^1$H NMR** (500 MHz, $CDCl_3$) δ 9.63 (d, *J* = 43.7 Hz, 1H), 8.62 (dd, *J* = 14.8, 2.2 Hz, 1H), 8.41 (d, *J* = 11.3 Hz, 1H), 8.32 (q, *J* = 2.2 Hz, 1H), 7.54 (dd, *J* = 14.8, 11.1 Hz, 1H), 7.41 (d, *J* = 8.5 Hz, 2H), 7.37 – 7.30 (m, 2H), 7.21 (d, *J* = 15.0 Hz, 1H), 4.81 (td, *J* = 6.5, 2.3 Hz, 1H), 4.16 – 3.40 (m, 10H), 2.68 (d, *J* = 1.9 Hz, 3H), 2.44 – 2.37 (m, 3H), 1.68 (s, 3H).

**$^{13}$C NMR** (126 MHz, $CDCl_3$) δ 169.51, 169.40, 164.57, 164.24, 164.13, 163.96, 162.89, 162.52, 155.74, 155.66, 150.03 (d, *J* = 9.6 Hz), 144.10, 139.05, 138.93, 136.84, 136.62, 135.74, 135.06, 134.48, 132.10, 132.02, 131.30, 130.97, 130.60, 129.82, 128.76, 127.01, 126.92, 54.68, 54.37, 45.88, 45.41, 42.37, 42.06, 41.85, 41.22, 35.29, 14.40, 13.13, 11.83.

**HRMS (ESI)** m/z calcd for $C_{32}H_{31}Cl_2N_8O_3S^+$ [M+H]$^+$: 677.1611; found: 677.1554

***tert*-butyl (*E*)-4-((5-chloropyridin-3-yl)amino)-4-oxobut-2-enoate (LO358)**

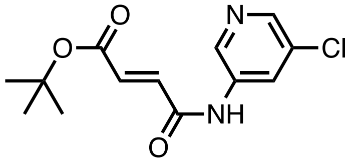

**General Procedure A** was followed with (*E*)-4-(*tert*-butoxy)-4-oxobut-2-enoic acid (160.7 mg, 0.93 mmol), HATU (354.9 mg, 0.93 mmol), DIPEA (0.54 mL, 3.11 mmol), and 5-chloropyridin-3-amine (100.0 mg, 0.78 mmol). The crude residue was purified by silica gel chromatography (0-60% EtOAc in hexanes) to afford 83.2 mg (37%) of the title compound as a beige powder.

**$^1$H NMR** (500 MHz, MeOD) δ 8.66 (d, *J* = 2.1 Hz, 1H), 8.39 (t, *J* = 2.2 Hz, 1H), 8.30 (d, *J* = 2.2 Hz, 1H), 7.08 (d, *J* = 15.4 Hz, 1H), 6.79 (d, *J* = 15.4 Hz, 1H), 1.53 (s, 9H).

**$^{13}$C NMR** (126 MHz, MeOD) δ 164.40, 163.31, 142.98, 138.58, 136.40, 134.82, 133.09, 131.88, 126.59, 81.60, 26.82.

**HRMS (ESI)** m/z calcd for $C_{13}H_{16}ClN_2O_3^+$ [M+H]$^+$: 283.0844; found: 283.0818

***tert*-butyl (*E*)-4-(4-((5-chloropyridin-3-yl)amino)-4-oxobut-2-enoyl)piperazine-1-carboxylate (LO405)**

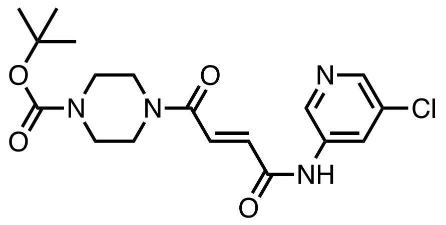

**General Procedure C** was followed with **LO358** (100.0 mg, 0.25 mmol), and TFA (0.87 mL, 11.32 mmol) for 2 hours. The crude material was used without further purification.
**General Procedure A** was followed with *tert*-butyl piperazine-1-carboxylate (54.9 mg, 0.29 mmol), HATU (134.5 mg, 0.35 mmol), DIPEA (0.21 mL, 0.1.18 mmol), and the acid from above (80.2 mg, 0.35 mmol). The crude residue was purified by silica gel chromatography (0-60% EtOAc in hexanes) to afford 55.1 mg (47%) of the title compound as a beige powder.
**$^1$H NMR** (400 MHz, $CDCL_3$) δ 8.59 (d, *J* = 2.0 Hz, 1H), 8.37 (d, *J* = 2.5 Hz, 1H), 8.32 (d, *J* = 2.2 Hz, 1H), 7.56 (d, *J* = 14.8 Hz, 1H), 7.33 (d, *J* = 14.8 Hz, 1H), 3.69 (dt, *J* = 29.5, 4.8 Hz, 4H), 3.59 – 3.45 (m, 4H), 1.48 (s, 9H).**$^{13}$C NMR** (101 MHz, $CDCL_3$) δ 164.60, 162.80, 154.43, 144.26, 139.17, 139.10, 135.81, 135.28, 132.14, 131.04, 126.98, 80.89, 46.23, 42.55, 28.43.
**HRMS (ESI)** m/z calcd for $C_{18}H_{24}ClN_4O_4^+$ $[M+H]^+$: 395.1481; found: 395.1423

**(E)-N-(5-chloropyridin-3-yl)-4-(4-(hex-5-yn-1-yl)piperazin-1-yl)-4-oxobut-2-enamide (LO406)**

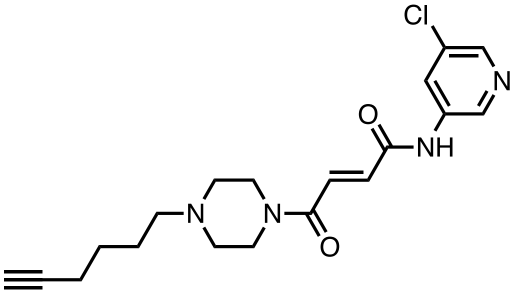

**General Procedure B** was followed with **LO405** (35.0 mg, 0.09 mmol), and TFA (0.26 mL, 3.4 mmol) for 2 hours. The crude material was used without further purification.
A combination of the amine from above (26.1 mg, 0.09 mmol) and potassium carbonate (36.8, 0.27 mmol) was suspended in acetonitrile (3 mL) and stirred at ambient temperature for 10 minutes. 6-bromohex-1-yne (0.02 mL, 0.18 mmol) was added, and the reaction mixture was stirred at 60°C for 2 hours. The reaction mixture was concentrated *in vacuo*, and the crude residue was purified by silica gel chromatography (0-30% EtOAc in hexanes) to afford 26.1 mg (79%) of the title compound as a yellow oil.
**$^1$H NMR** (400 MHz, $CDCL_3$) δ 9.90 (s, 1H), 8.62 (d, *J* = 2.2 Hz, 1H), 8.35 (d, *J* = 2.3 Hz, 1H), 8.30 (d, *J* = 2.2 Hz, 1H), 7.56 (d, *J* = 14.9 Hz, 1H), 7.35 (d, *J* = 14.9 Hz, 1H), 3.76 (t, *J* = 5.0 Hz, 2H), 3.67 (t, *J* = 5.0 Hz, 2H), 2.48 (q, *J* = 4.5 Hz, 4H), 2.38 (t, *J* = 6.9 Hz, 2H), 2.22 (td, *J* = 6.7, 2.7 Hz, 2H), 1.95 (t, *J* = 2.6 Hz, 1H), 1.58 (dtt, *J* = 21.4, 8.8, 4.9 Hz, 4H).
**$^{13}$C NMR** (101 MHz, $CDCL_3$) δ 164.25, 163.06, 144.13, 139.28, 135.94, 135.00, 132.06, 131.25, 126.97, 84.23, 68.69, 57.65, 53.35, 52.63, 46.50, 42.81, 26.24, 25.74, 18.35.
**HRMS (ESI)** m/z calcd for $C_{19}H_{24}ClN_4O_2^+$ $[M+H]^+$: 375.1582; found 375.1546

**(*E*)-4-(4-(3-amino-6-(2-hydroxyphenyl)pyridazin-4-yl)piperazin-1-yl)-*N*-(5-chloropyridin-3-yl)-4-oxobut-2-enamide (LO362)**

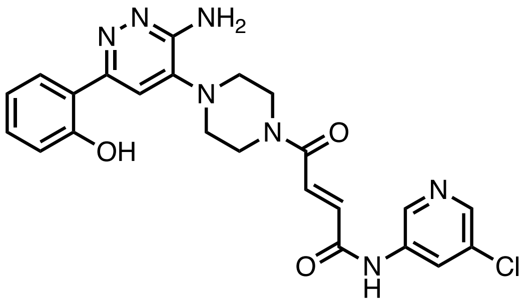

**General Procedure C** was followed with **LO358** (35.0 mg, 0.12 mmol), and TFA (0.30 mL, 3.96 mmol) for 2 hours. The crude material was used without further purification.
**General Procedure A** was followed with 2-(6-amino-5-(piperazin-1-yl)pyridazin-3-yl)phenol (28.0 mg, 0.10 mmol), HATU (47.1 mg, 0.12 mmol), DIPEA (0.07 mL, 0.41 mmol), and the acid from above (28.1 mg, 0.12 mmol). The crude residue was purified by silica gel chromatography (0-10% MeOH in DCM) to afford 32.5 mg (66%) of the title compound as a yellow-white powder.
**1H NMR** (500 MHz, DMSO) δ 14.11 (s, 1H), 10.93 (s, 1H), 8.71 (d, *J* = 2.1 Hz, 1H), 8.38 (d, *J* = 2.3 Hz, 1H), 8.33 (t, *J* = 2.2 Hz, 1H), 7.91 (dd, *J* = 8.4, 1.7 Hz, 1H), 7.59 – 7.52 (m, 2H), 7.27 – 7.22 (m, 1H), 7.09 (d, *J* = 15.0 Hz, 1H), 6.90 (ddd, *J* = 6.8, 6.0, 1.4 Hz, 2H), 6.52 (d, *J* = 22.6 Hz, 2H), 3.85 (dt, *J* = 22.9, 4.9 Hz, 4H), 3.14 (q, *J* = 5.5 Hz, 4H).
$^{13}$C NMR (126 MHz, DMSO) δ 163.59, 163.41, 158.88, 155.23, 153.46, 143.40, 140.44, 139.59, 136.79, 134.46, 131.66, 131.16, 130.68, 126.71, 125.98, 118.99, 118.25, 117.84, 111.77, 55.39, 49.54, 48.88, 45.59, 41.81.
**HRMS (ESI)** m/z calcd for $C_{23}H_{23}ClN_7O_3^+$ [M+H]$^+$: 480.1545; found: 480.1503

**(*E*)-2-((5-(4-(4-((5-chloropyridin-3-yl)amino)-4-oxobut-2-enoyl)piperazin-1-yl)pyridin-2-yl)amino)-7-cyclopentyl-*N*,*N*-dimethyl-7*H*-pyrrolo[2,3-*d*]pyrimidine-6-carboxamide (LO363)**

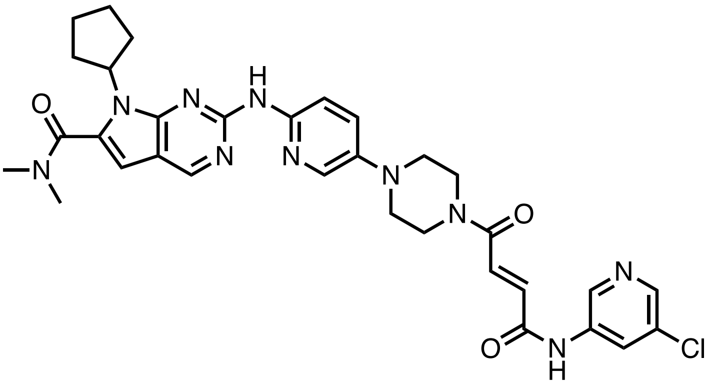

**General Procedure C** was followed with **LO358** (35.0 mg, 0.12 mmol), and TFA (0.30 mL, 3.96 mmol) for 2 hours. The crude material was used without further purification.
**General Procedure A** was followed with 7-cyclopentyl-*N*,*N*-dimethyl-2-((5-(piperazin-1-yl)pyridin-2-yl)amino)-7*H*-pyrrolo[2,3-*d*]pyrimidine-6-carboxamide (ribociclib) (44.8 mg, 0.10 mmol), HATU (47.1 mg, 0.12 mmol), DIPEA (0.07 mL, 0.41 mmol), and the acid from above (28.1 mg, 0.12 mmol). The crude residue was purified by silica gel chromatography (0-10% MeOH in DCM) to afford 28.0 mg (42%) of the title compound as a yellow powder.
**$^{1}$H NMR** (500 MHz, DMSO) δ 10.92 (s, 1H), 9.62 (s, 1H), 8.80 (s, 1H), 8.71 (d, *J* = 2.1 Hz, 1H), 8.41 – 8.30 (m, 2H), 8.13 (d, *J* = 9.1 Hz, 1H), 8.04 (d, *J* = 3.0 Hz, 1H), 7.59 – 7.49 (m, 2H), 7.07 (d, *J* = 15.0 Hz, 1H), 6.63 (s, 1H), 4.75 (p, *J* = 8.9 Hz, 1H), 3.78 (dt, *J* = 20.0, 5.0 Hz, 4H), 3.19 (dt, *J* = 14.0, 5.3 Hz, 4H), 3.06 (s, 6H), 2.48 – 2.38 (m, 2H), 1.99 (dh, *J* = 14.3, 4.3 Hz, 4H), 1.71 – 1.59 (m, 2H).

**$^{13}$C NMR** (126 MHz, DMSO) δ 163.62, 163.39, 163.30, 151.70, 146.84, 143.38, 142.36, 139.60, 136.80, 134.35, 131.71, 131.15, 125.98, 113.32, 112.55, 101.11, 57.43, 50.02, 49.27, 45.69, 41.96, 35.07, 30.24, 24.68.
**HRMS (ESI)** m/z calcd for $C_{32}H_{36}ClN_{10}O_3^+$ [M+H]$^+$: 643.2655; found: 643.2598

**(*E*)-*N*-(4-(3-chloro-4-cyanophenoxy)cyclohexyl)-6-(4-(4-((5-chloropyridin-3-yl)amino)-4-oxobut-2-enoyl)piperazin-1-yl)pyridazine-3-carboxamide (LO412)**

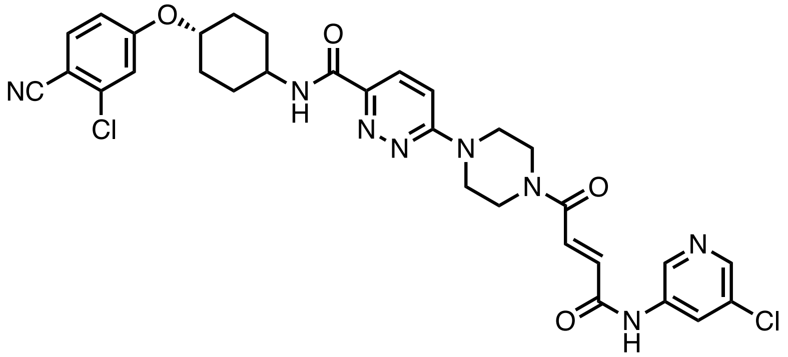

**General Procedure C** was followed with **LO358** (35.0 mg, 0.12 mmol), and TFA (0.30 mL, 3.96 mmol) for 2 hours. The crude material was used without further purification.
**General Procedure A** was followed with *N*-(4-(3-chloro-4-cyanophenoxy)cyclohexyl)-6-(piperazin-1-yl)pyridazine-3-carboxamide (45.5 mg, 0.10 mmol), HATU (47.1 mg, 0.12 mmol), DIPEA (0.07 mL, 0.41 mmol), and the acid from above (28.1 mg, 0.12 mmol). The crude residue was purified by silica gel chromatography (0-10% MeOH in DCM) to afford 35.8 mg (53%) of the title compound as a white powder.
**$^{1}$H NMR** (500 MHz, DMSO) δ 11.02 (s, 1H), 8.72 (d, *J* = 2.2 Hz, 1H), 8.66 (d, *J* = 8.2 Hz, 1H), 8.38 (d, *J* = 2.2 Hz, 1H), 8.34 (t, *J* = 2.2 Hz, 1H), 7.88 (dd, *J* = 12.5, 9.1 Hz, 2H), 7.55 (d, *J* = 15.0 Hz, 1H), 7.43 – 7.38 (m, 2H), 7.14 (dd, *J* = 8.7, 2.4 Hz, 1H), 7.08 (d, *J* = 14.9 Hz, 1H), 4.54 (tt, *J* = 10.4, 4.2 Hz, 1H), 3.92 – 3.71 (m, 9H), 2.14 – 2.07 (m, 3H), 1.95 – 1.86 (m, 2H), 1.70 – 1.60 (m, 2H), 1.57 – 1.47 (m, 2H).
**$^{13}$C NMR** (126 MHz, DMSO) δ 163.62, 162.90, 162.28, 160.40, 145.56, 143.38, 139.62, 137.49, 136.83, 136.24, 134.42, 131.69, 131.15, 126.96, 125.99, 117.29, 116.91, 115.95, 113.40, 103.67, 76.05, 47.59, 45.05 (d, *J* = 27.3 Hz), 44.38, 41.62, 31.18, 30.34, 29.88.
**HRMS (ESI)** m/z calcd for $C_{31}H_{31}Cl_2N_8O_4^+$ [M+H]$^+$: 649.1840; found: 649.1774

**4-(4-(4-(4-(vinylsulfonyl)piperazin-1-yl)phenyl)thiazol-2-yl)morpholine (ML1-95)**

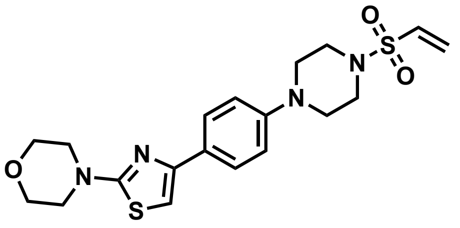

**General Procedure B** was followed with *tert*-butyl 4-(4-(2-morpholinothiazol-4-yl)phenyl)piperazine-1-carboxylate (20.0 mg, 0.05 mmol), and TFA (0.11 mL, 1.49 mmol) for 2 hours. The crude material was used without further purification.
The above product (15.3 mg, 0.05 mmol) was dissolved in dichloromethane (DCM) (0.3 mL) and triethylamine (0.03 mL, 0.21 mmol) was added at 0 °C. 2-chloroethanesulfonyl chloride (0.01 mL, 0.07 mmol) in DCM (0.1 mL) was added dropwise and the resulting reaction mixture was stirred at ambient temperature overnight. The reaction was quenched with 5 times the reaction volume of water and extracted 3 times with DCM. The organic extracts were washed once with brine, dried over $Na_2SO_4$, vacuum filtered, and concentrated *in vacuo*.

The crude residue was purified by silica gel chromatography (0-50% EtOAc in hexanes) to afford 14.1 mg (72.2%) of the title compound as a yellow powder.

**$^1$H NMR** (500 MHz, $CDCl_3$) δ 7.77 – 7.71 (m, 2H), 6.94 – 6.88 (m, 2H), 6.66 (s, 1H), 6.46 (dd, *J* = 16.6, 9.9 Hz, 1H), 6.29 (d, *J* = 16.6 Hz, 1H), 6.09 (d, *J* = 10.0 Hz, 1H), 3.85 – 3.82 (m, 5H), 3.54 – 3.50 (m, 4H), 3.34 – 3.27 (m, 9H).

**$^{13}$C NMR** (126 MHz, $CDCl_3$) δ 171.19, 151.59, 150.02, 132.13, 129.30, 127.86, 127.10, 116.62, 99.97, 66.26, 49.09, 48.57, 45.49.

**HRMS (ESI)** m/z calcd for $C_{19}H_{25}N_4O_3S_2^+$ [M+H]$^+$: 421.1363; found: 421.1354

**Ethyl 6-(4-(2-morpholinothiazol-4-yl)phenoxy)hexanoate (LO422)**

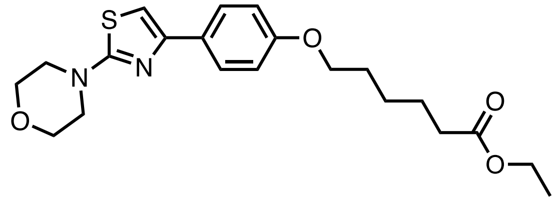

A combination of 4-(2-morpholinothiazol-4-yl)phenol (VPC-14228) (100.0 mg, 0.38 mmol) and potassium carbonate (158.1 mg, 1.14 mmol) was suspended in DMF (8 mL) and stirred at ambient temperature for 10 minutes. Ethyl 4-bromobutanoate (0.14 mL, 0.76 mmol) was added, and the reaction mixture was stirred at 90°C for 16 hours. . The reaction was quenched with 5 times the reaction volume of 5% LiCl(aq) and extracted 3 times with dichloromethane (DCM). The organic extracts were washed once with brine, dried over $Na_2SO_4$, vacuum filtered, and concentrated *in vacuo*. The crude residue was purified by silica gel chromatography (0-30% EtOAc in hexanes) to afford 110.8 mg (72%) of the title compound as a clear oil.

**$^1$H NMR** (500 MHz, $CDCl_3$) δ 7.77 – 7.70 (m, 2H), 6.91 – 6.85 (m, 2H), 6.65 (s, 1H), 4.13 (q, *J* = 7.1 Hz, 2H), 3.97 (t, *J* = 6.5 Hz, 2H), 3.86 – 3.79 (m, 4H), 3.52 (t, 4H), 2.33 (t, *J* = 7.5 Hz, 2H), 1.85 – 1.77 (m, 2H), 1.71 (d, *J* = 7.8 Hz, 2H), 1.56 – 1.47 (m, 2H), 1.26 (t, *J* = 7.1 Hz, 3H).

$^{13}$C NMR (126 MHz, $CDCl_3$) δ 173.67, 171.16, 158.75, 151.70, 127.29, 114.45, 99.82, 67.67, 66.24, 60.26, 48.57, 34.26, 29.06, 28.97, 25.66, 24.74, 14.26.

**HRMS (ESI)** m/z calcd for $C_{21}H_{29}N_2O_4S^+$ [M+H]$^+$: 405.1843 ; found: 405.1805

**(*E*)-*N*-(5-chloropyridin-3-yl)-4-(4-(6-(4-(2-morpholinothiazol-4-yl)phenoxy)hexanoyl)piperazin-1-yl)-4-oxobut-2-enamide (LO425)**

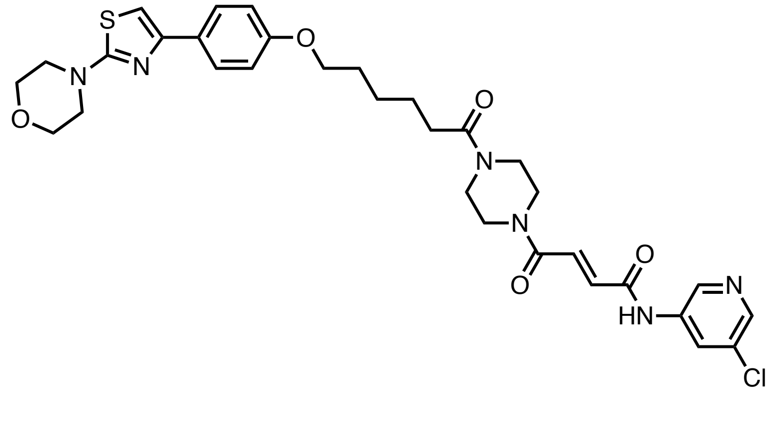

**General Procedure B** was followed with **LO405** (50.0 mg, 0.13 mmol), and TFA (0.37 mL, 4.86 mmol) for 2 hours. The afforded crude amine was used without further purification.

A solution of **LO422** (61.5 mg, 0.15 mmol) in 2M lithium hydroxide (1 mL, 2 mmol), ethanol (3 mL), and THF (3 mL) was stirred overnight. The reaction quenched by acidifying to pH 2 using 1 M HCL, and then the volatiles were removed *in vacuo*. The resultant aqueous solution was extracted 3 times with DCM. The organic extracts were washed once with brine, dried over $Na_2SO_4$, vacuum filtered, and concentrated *in vacuo*. The afforded crude acid was then used without further purification.

**General Procedure A** was followed with the amine from above (37.3 mg, 0.13 mmol), HATU (57.8 mg, 0.15 mmol), DIPEA (0.09 mL, 0.51 mmol), and the acid from above (57.2 mg, 0.15 mmol). The crude residue was purified by silica gel chromatography (0-10% MeOH in DCM) to afford 21.9 mg (26%) of the title compound as a white powder.

**$^1$H NMR** (500 MHz, $CDCl_3$) δ 9.55 (d, *J* = 14.2 Hz, 1H), 8.59 (s, 1H), 8.38 (d, *J* = 9.0 Hz, 1H), 8.33 (d, *J* = 2.2 Hz, 1H), 7.77 – 7.71 (m, 2H), 7.53 (t, *J* = 15.0 Hz, 1H), 7.34 – 7.27 (m, 1H), 6.91 – 6.85 (m, 2H), 6.65 (s, 1H), 3.99 (t, *J* = 6.3 Hz, 2H), 3.87 – 3.80 (m, 4H), 3.78 – 3.64 (m, 6H), 3.61 – 3.49 (m, 6H), 2.39 (t, *J* = 7.5 Hz, 2H), 1.82 (p, *J* = 6.6 Hz, 2H), 1.73 (p, *J* = 7.7 Hz, 2H), 1.55 (p, *J* = 7.4 Hz, 2H).

**$^{13}$C NMR** (126 MHz, $CDCl_3$) δ 171.77, 171.20, 164.40, 162.61, 158.69, 151.62, 144.21, 139.01, 135.62, 135.22, 132.09, 130.75, 127.93, 127.32, 126.90, 114.43, 99.89, 67.59, 66.23, 48.56, 45.99, 44.92, 42.52, 41.57, 33.09, 29.07, 25.89, 24.83.

**HRMS (ESI)** m/z calcd for $C_{32}H_{38}ClN_6O_5S^+$ $[M+H]^+$: 653.2307; found: 653.2309

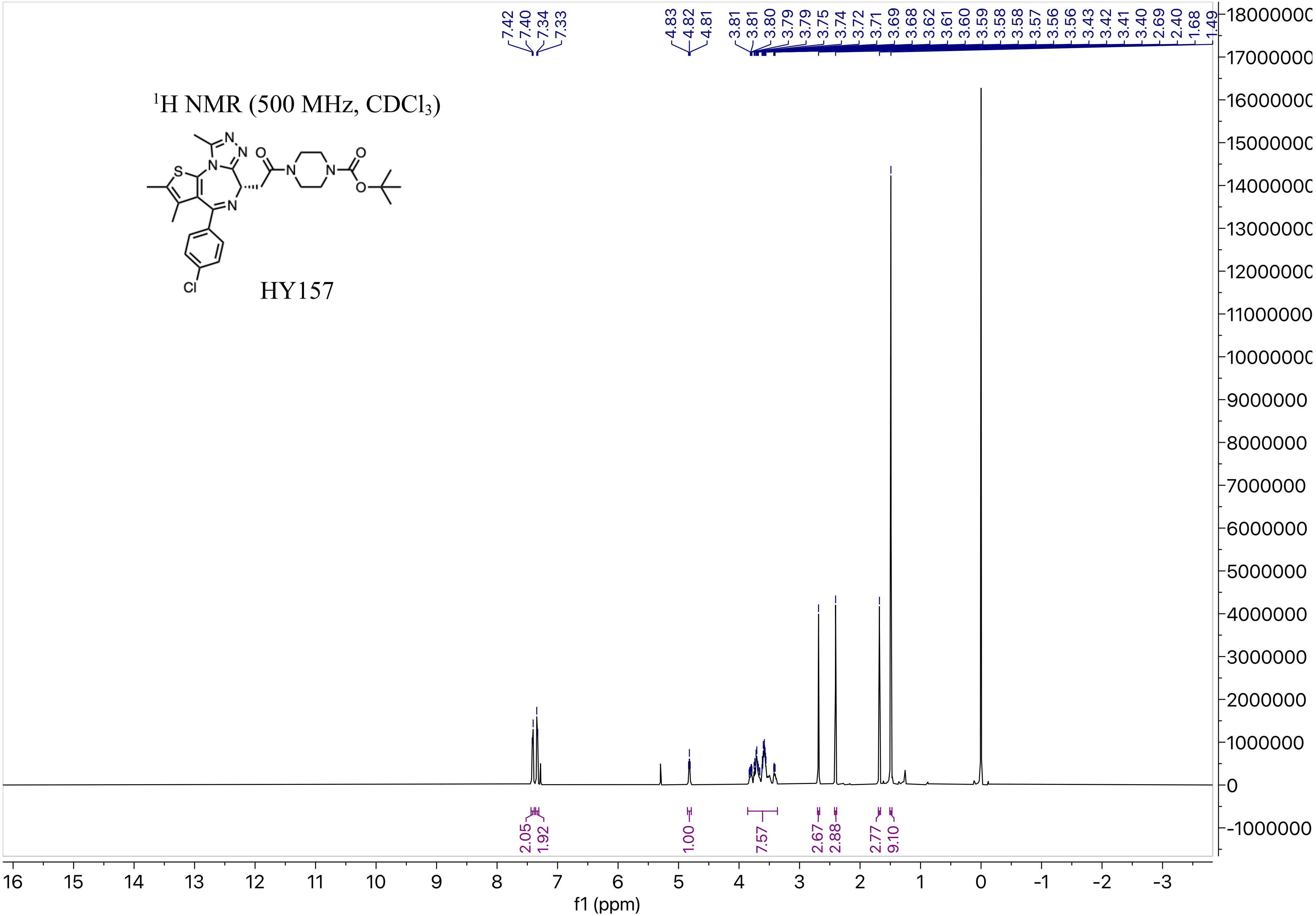


$^1$H NMR (500 MHz, $CDCl_3$)

HY157

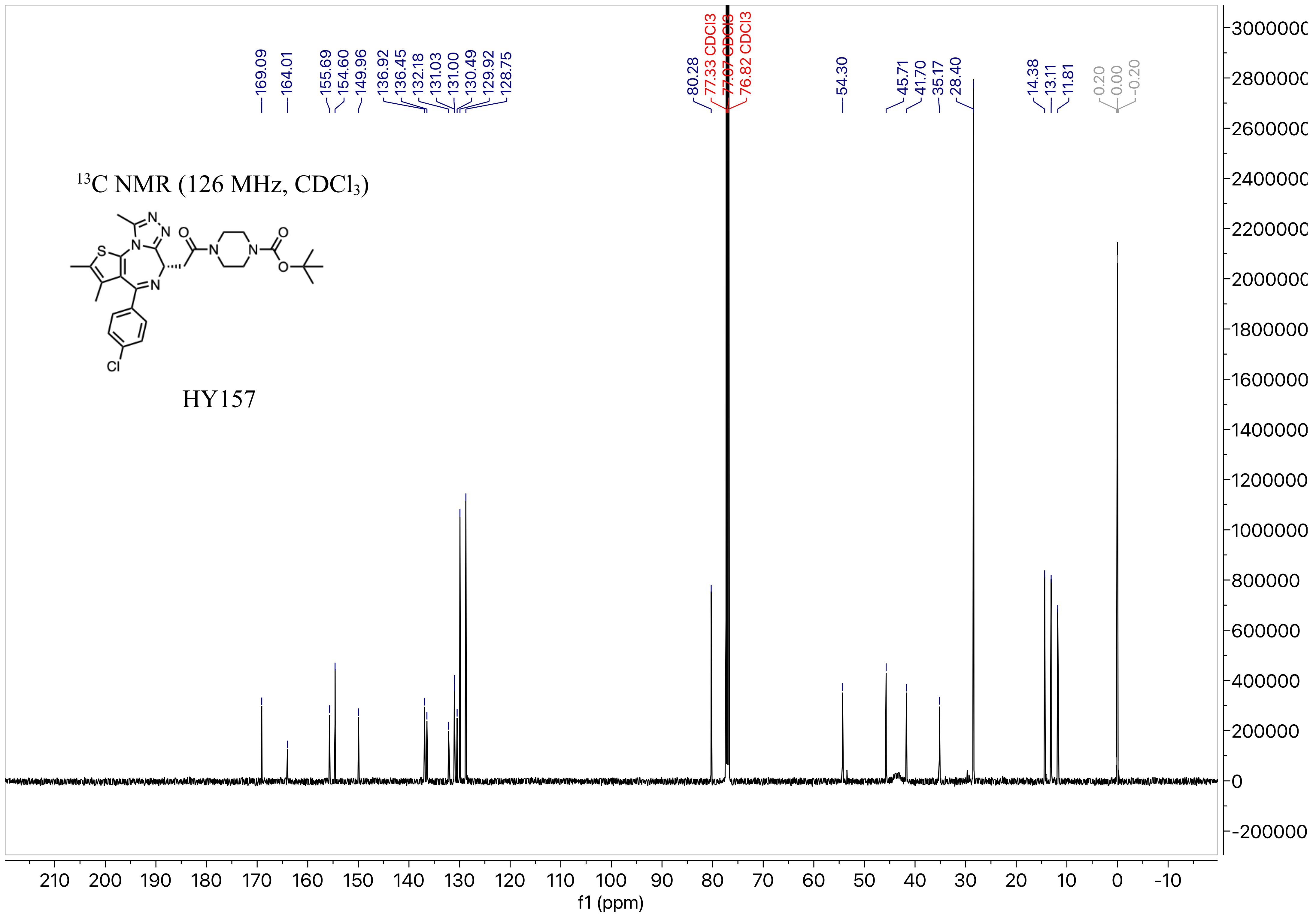

169.09
164.01
155.69
154.60
149.96
136.92
136.45
132.18
131.03
131.00
130.49
129.92
128.75
80.28
77.33 CDCl3
77.07 CDCl3
76.82 CDCl3
54.30
45.71
41.70
35.17
28.40
14.38
13.11
11.81
0.20
0.00
-0.20
13C NMR (126 MHz, CDCl3)
HY157
f1 (ppm)

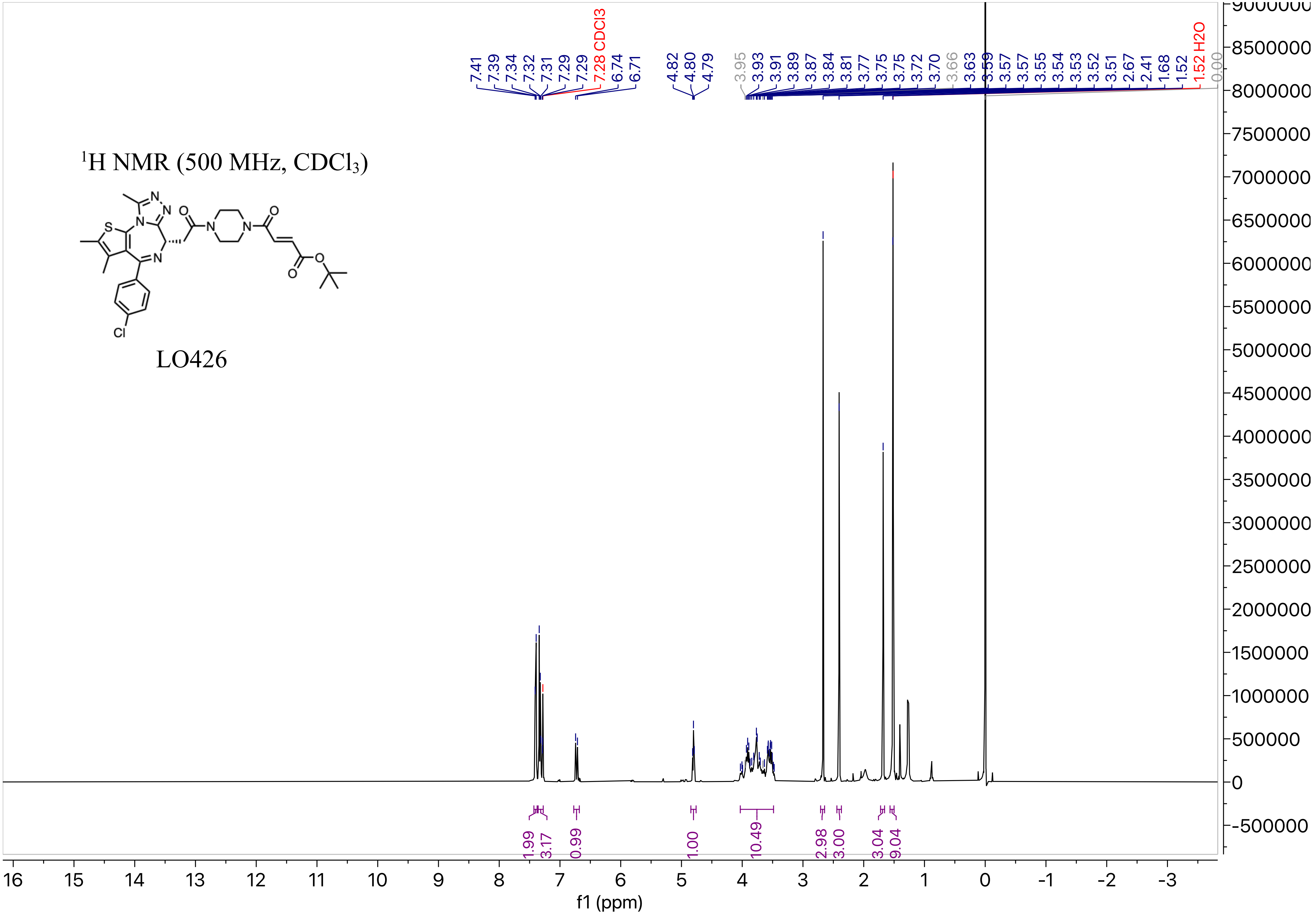


$^{1}H$ NMR (500 MHz, $CDCl_3$)

LO426

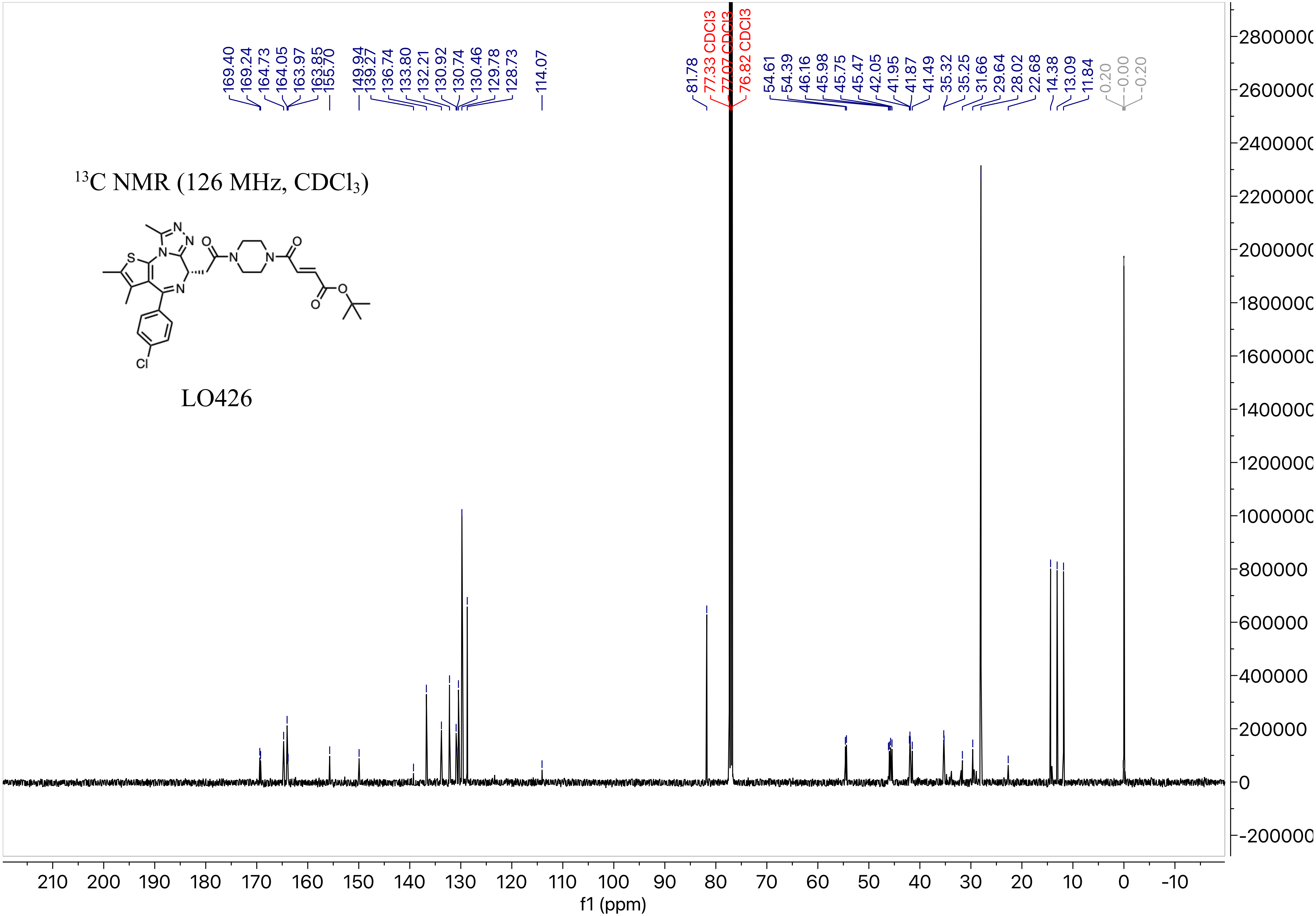

$^{13}$C NMR (126 MHz, CDCl$_3$)
LO426
169.40
169.24
164.73
164.05
163.97
163.85
155.70
149.94
139.27
136.74
133.80
132.21
130.92
130.74
130.46
129.78
128.73
114.07
81.78
77.33 CDCl3
77.07 CDCl3
76.82 CDCl3
54.61
54.39
46.16
45.98
45.75
45.47
42.05
41.95
41.87
41.49
35.32
35.25
31.66
29.64
28.02
22.68
14.38
13.09
11.84
0.20
-0.00
-0.20
f1 (ppm)

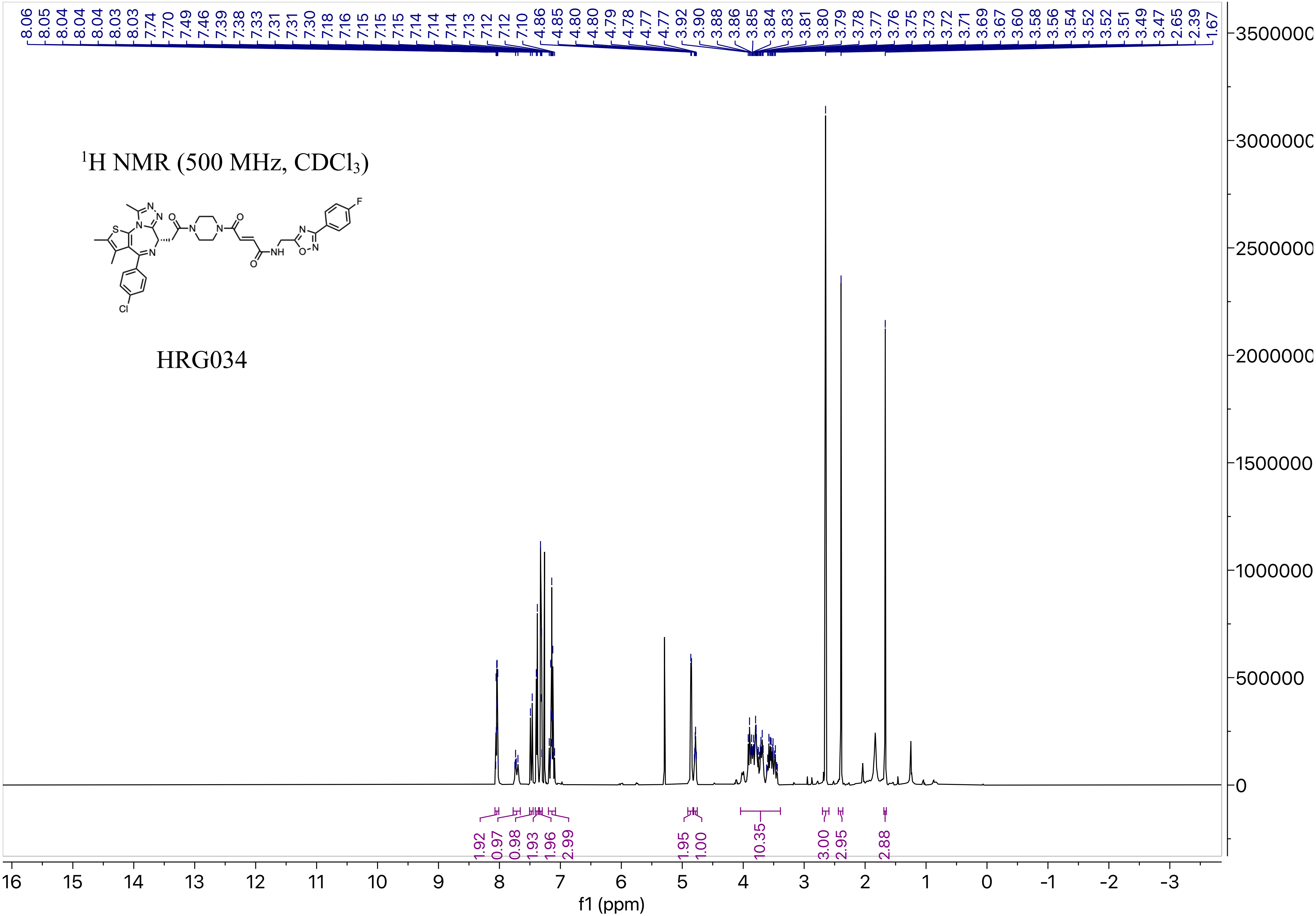

1H NMR (500 MHz, CDCl3)
HRG034
8.06 8.05 8.04 8.04 8.04 8.03 8.03 7.74 7.70 7.49 7.46 7.39 7.38 7.33 7.31 7.31 7.30 7.18 7.16 7.15 7.15 7.15 7.14 7.14 7.14 7.13 7.12 7.12 7.10 4.86 4.85 4.80 4.80 4.79 4.78 4.77 4.77 3.92 3.90 3.88 3.86 3.85 3.84 3.83 3.81 3.80 3.79 3.78 3.77 3.76 3.75 3.73 3.72 3.71 3.69 3.67 3.60 3.58 3.56 3.54 3.52 3.52 3.51 3.49 3.47 2.65 2.39 1.67
1.92 0.97 0.98 1.93 1.96 2.99 1.95 1.00 10.35 3.00 2.95 2.88
f1 (ppm)

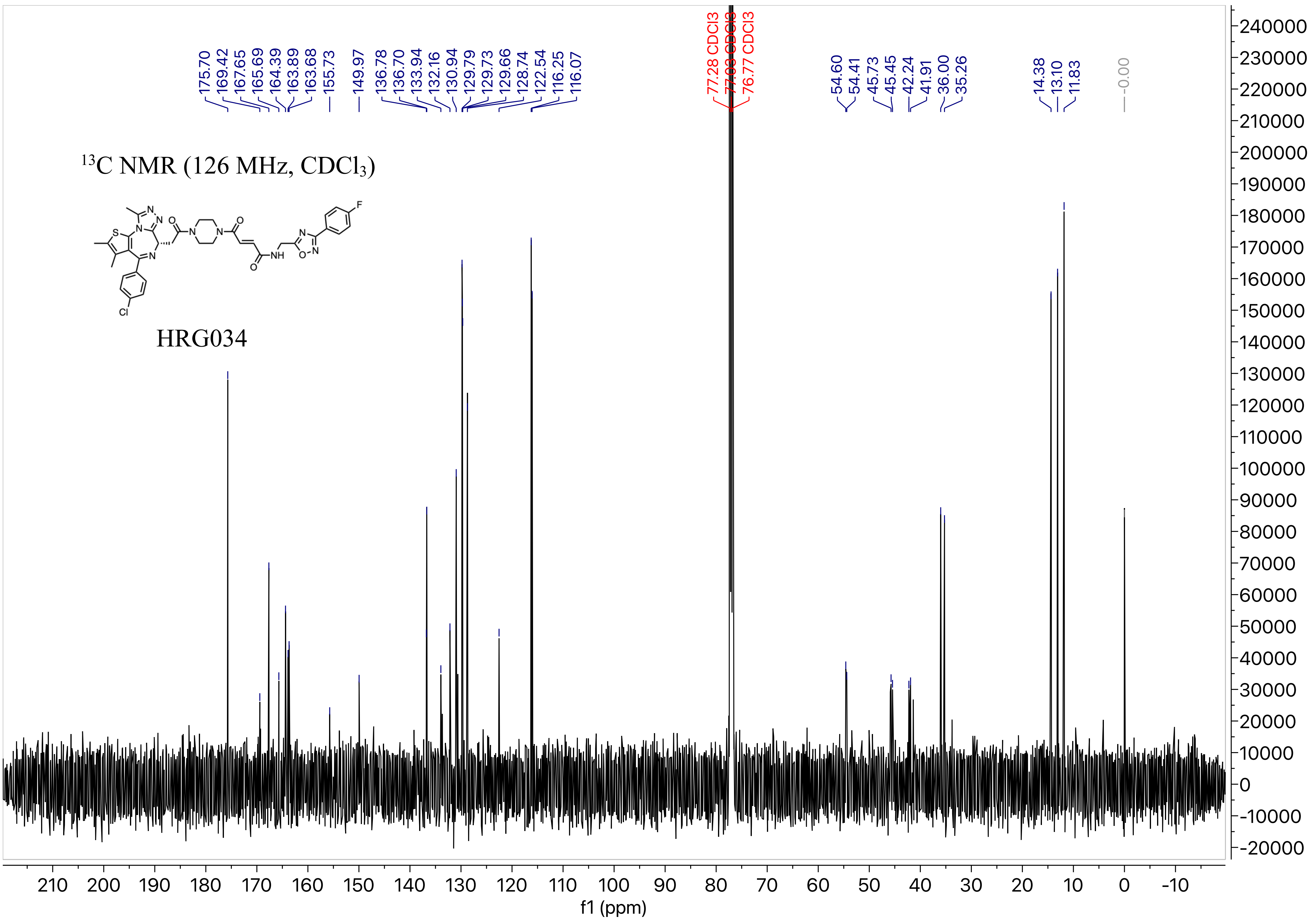

175.70
169.42
167.65
165.69
164.39
163.89
163.68
155.73
149.97
136.78
136.70
133.94
132.16
130.94
129.79
129.73
129.66
128.74
122.54
116.25
116.07
77.28 CDCl3
77.03 CDCl3
76.77 CDCl3
54.60
54.41
45.73
45.45
42.24
41.91
36.00
35.26
14.38
13.10
11.83
-0.00
13C NMR (126 MHz, CDCl3)
HRG034
f1 (ppm)

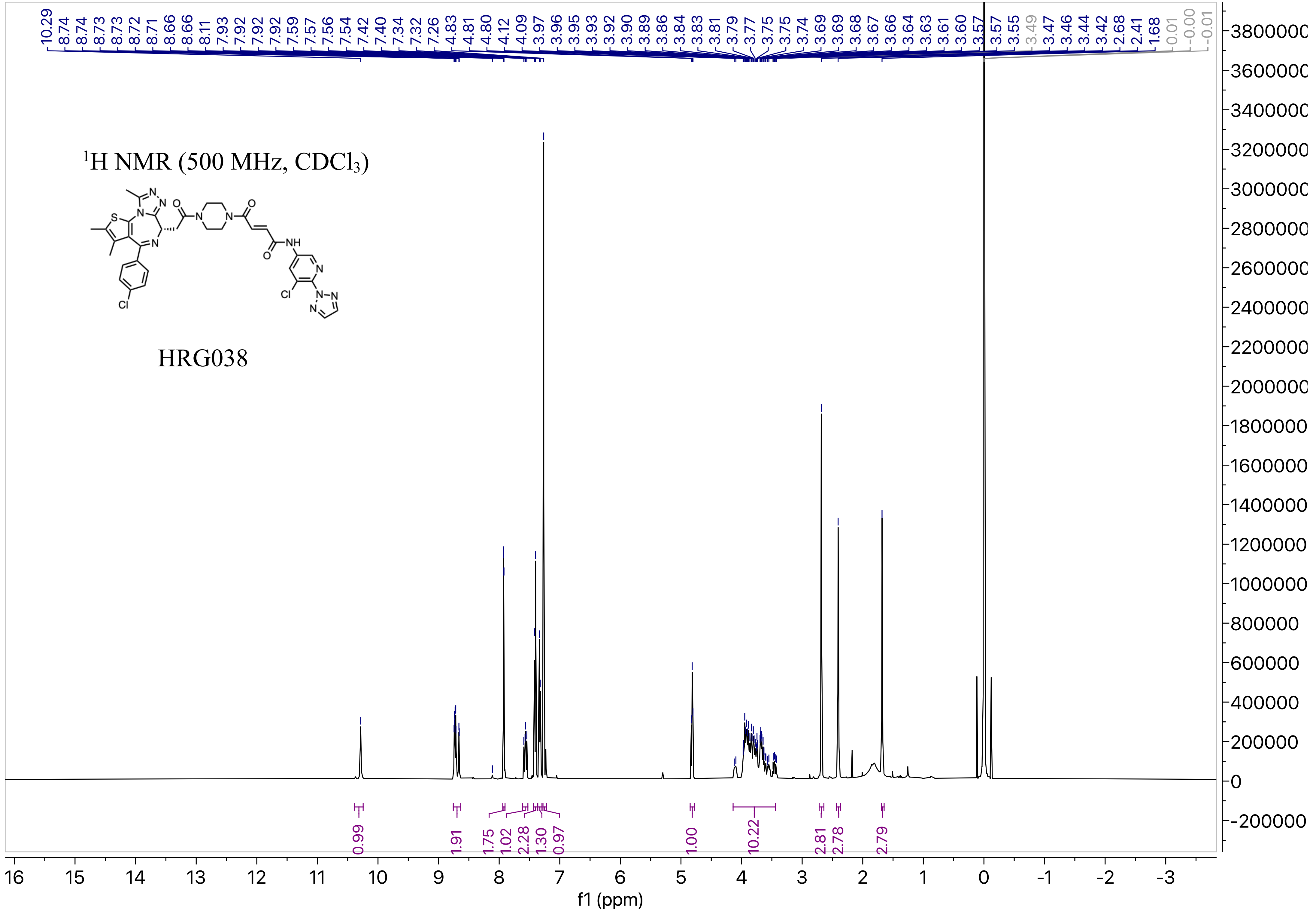

1H NMR (500 MHz, CDCl3)
HRG038
f1 (ppm)

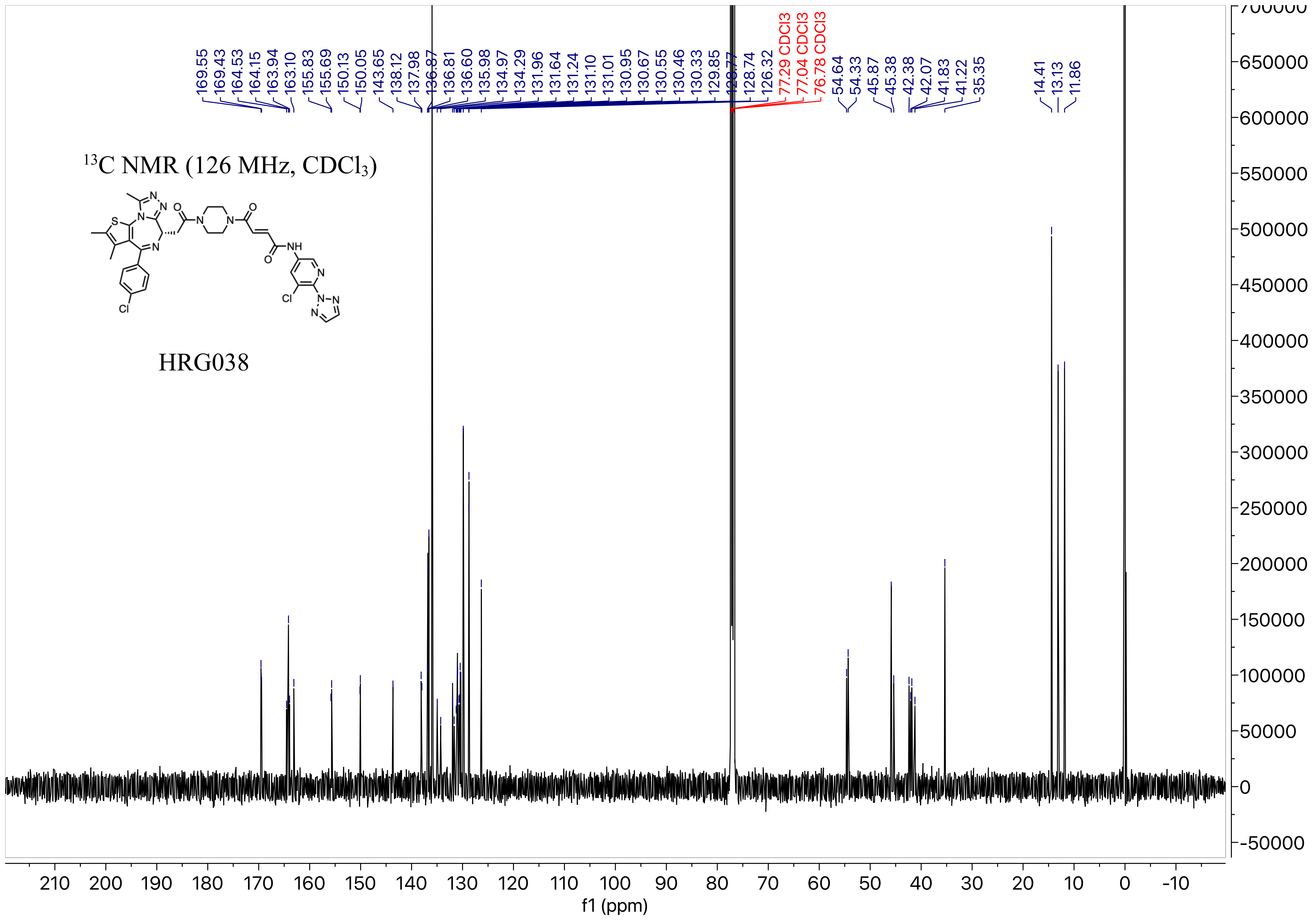

$^{13}C$ NMR (126 MHz, $CDCl_3$)
HRG038
169.55
169.43
164.53
164.15
163.94
163.10
155.83
155.69
150.13
150.05
143.65
138.12
137.98
136.87
136.81
136.60
135.98
134.97
134.29
131.96
131.64
131.24
131.10
131.01
130.95
130.67
130.55
130.46
130.33
129.85
128.77
128.74
126.32
77.29 CDCl3
77.04 CDCl3
76.78 CDCl3
54.64
54.33
45.87
45.38
42.38
42.07
41.83
41.22
35.35
14.41
13.13
11.86
f1 (ppm)

$^{1}$H NMR (500 MHz, $CDCl_3$)

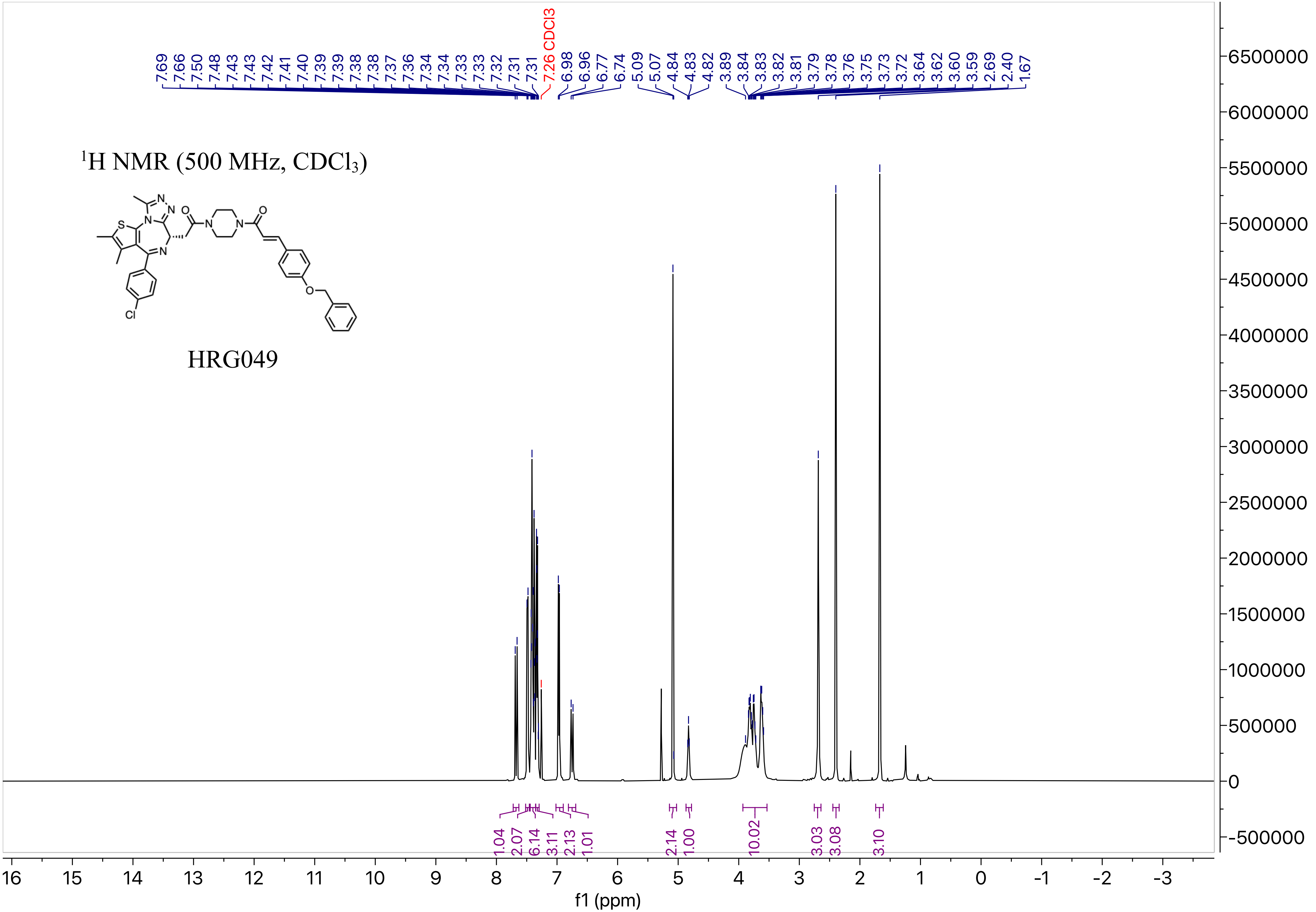


HRG049

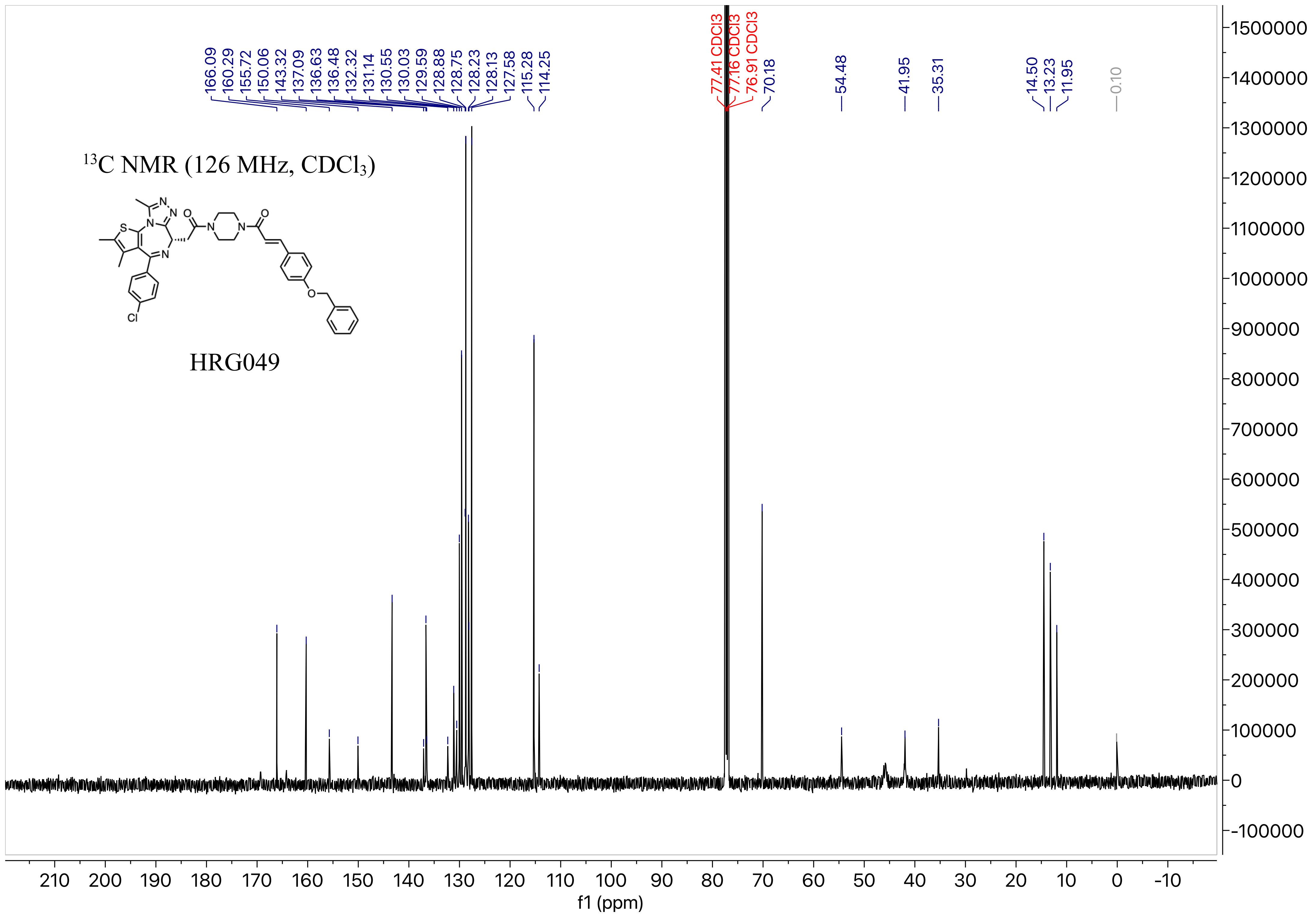


$^{13}C$ NMR (126 MHz, $CDCl_3$)

HRG049

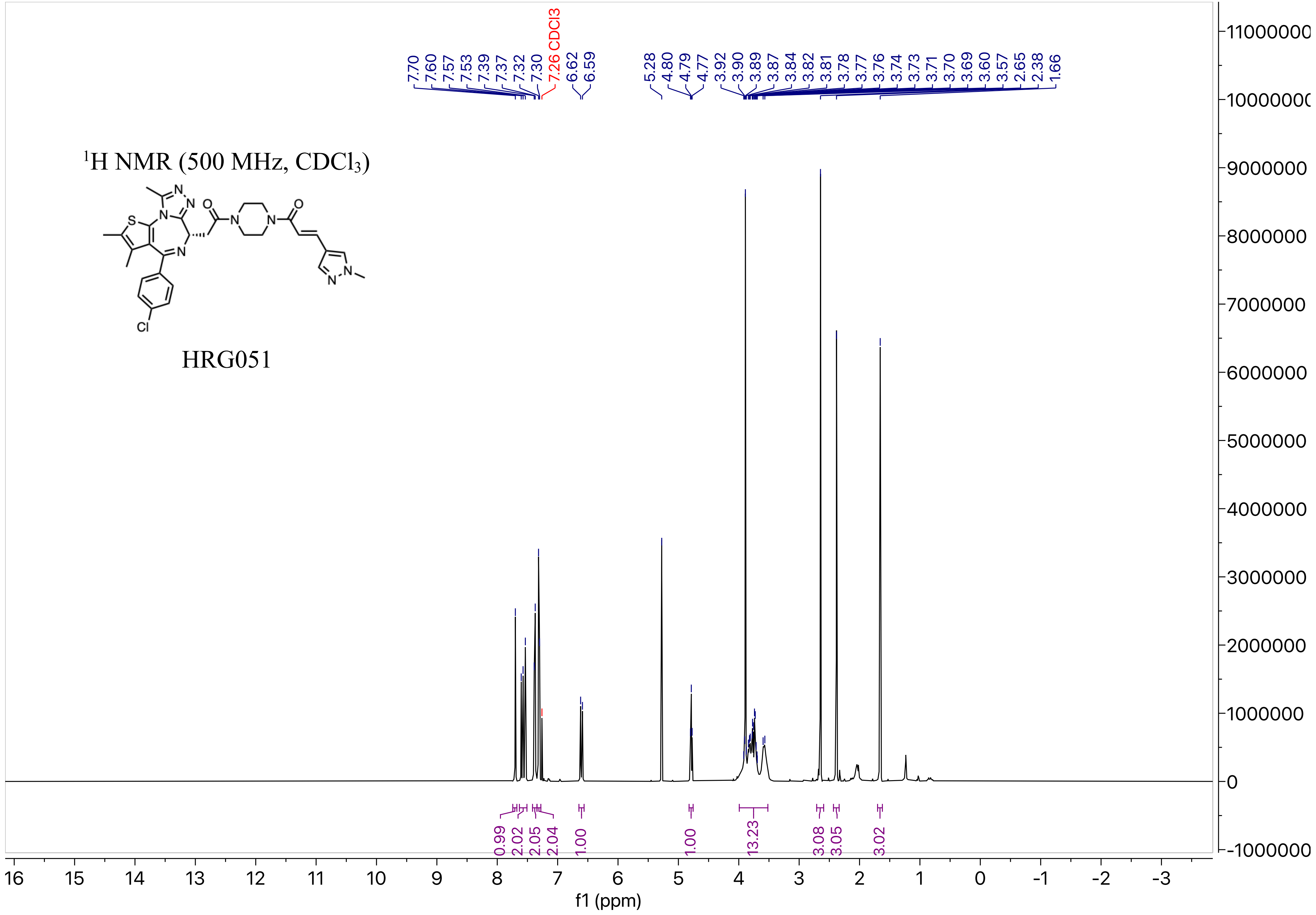

$^{1}H$ NMR (500 MHz, $CDCl_3$)
HRG051
7.70 7.60 7.57 7.53 7.39 7.37 7.32 7.30 7.26 CDCl3 6.62 6.59
5.28 4.80 4.79 4.77 3.92 3.90 3.89 3.87 3.84 3.82 3.81 3.78 3.77 3.76 3.74 3.73 3.71 3.70 3.69 3.60 3.57 2.65 2.38 1.66
0.99 2.02 2.05 2.04 1.00 1.00 13.23 3.08 3.05 3.02
f1 (ppm)

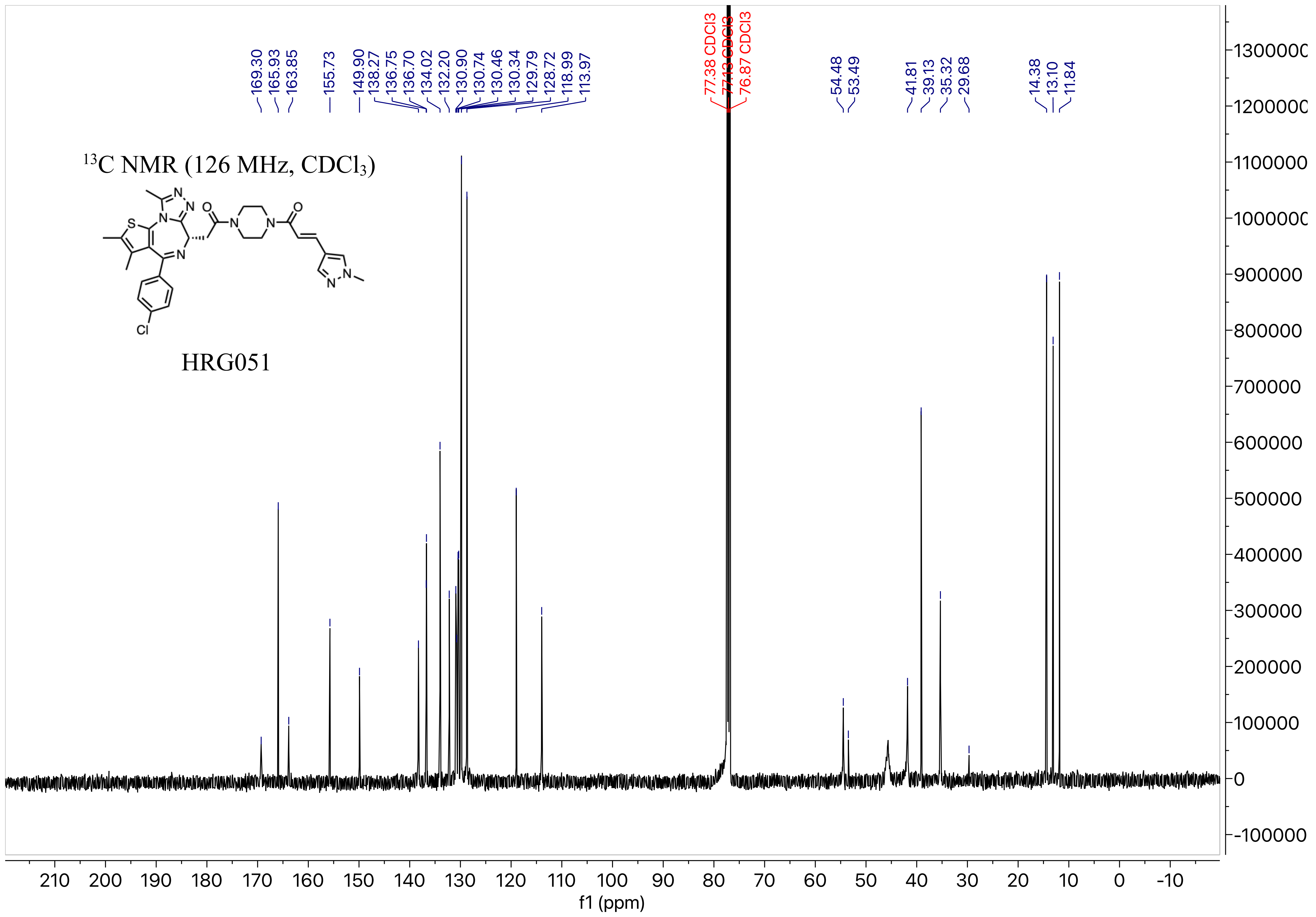

13C NMR (126 MHz, CDCl3)
HRG051
169.30
165.93
163.85
155.73
149.90
138.27
136.75
136.70
134.02
132.20
130.90
130.74
130.46
130.34
129.79
128.72
118.99
113.97
77.38 CDCl3
77.13 CDCl3
76.87 CDCl3
54.48
53.49
41.81
39.13
35.32
29.68
14.38
13.10
11.84
f1 (ppm)

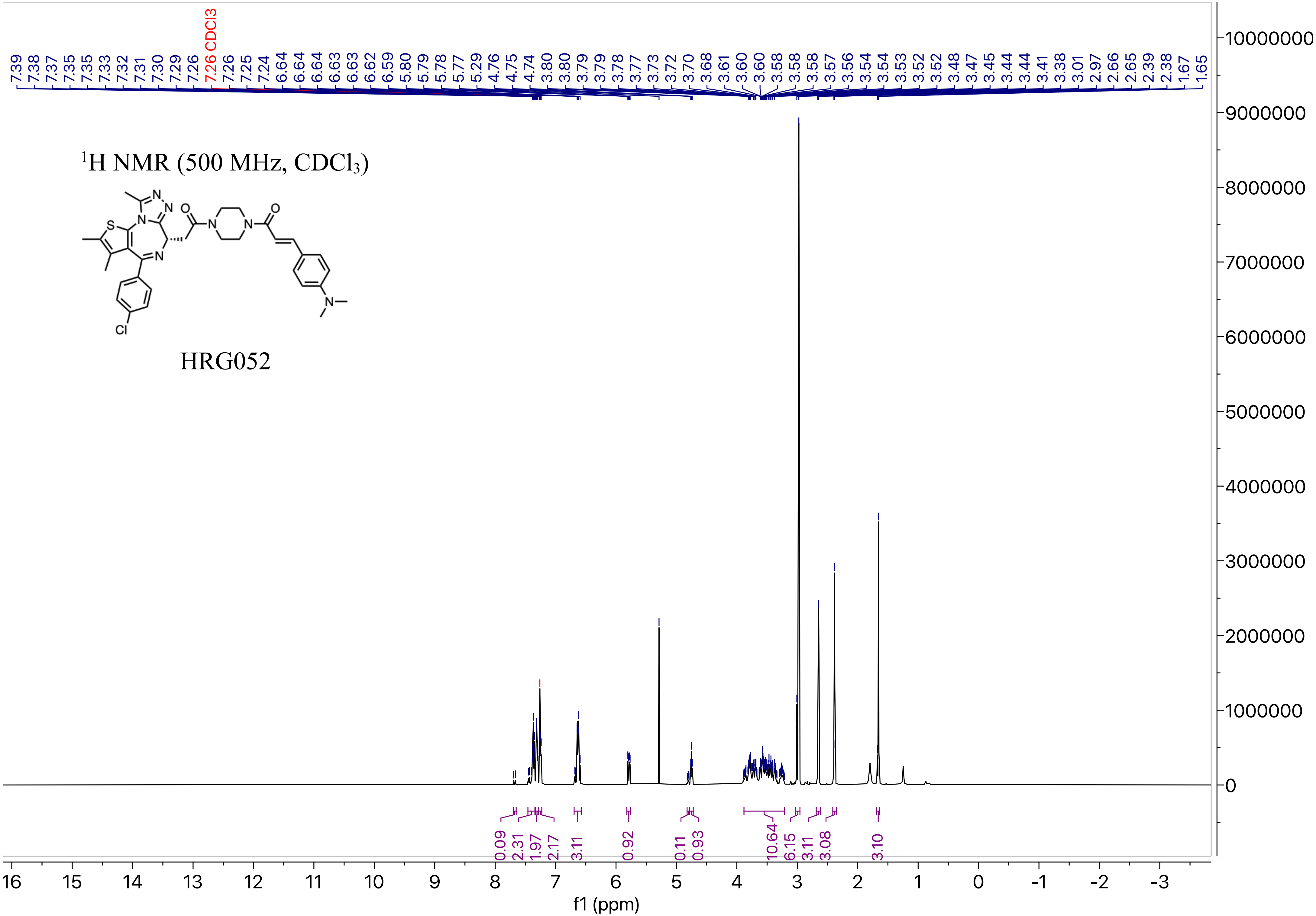

$^1$H NMR (500 MHz, $CDCl_3$)
HRG052
7.39 7.38 7.37 7.35 7.35 7.33 7.32 7.31 7.30 7.29 7.26 7.26 CDCl3 7.26 7.25 7.24 6.64 6.64 6.64 6.63 6.63 6.62 6.59 5.80 5.79 5.78 5.77 5.29 4.76 4.75 4.74 3.80 3.80 3.79 3.79 3.78 3.77 3.73 3.72 3.70 3.68 3.61 3.60 3.60 3.58 3.58 3.58 3.57 3.56 3.54 3.54 3.53 3.52 3.52 3.48 3.47 3.45 3.44 3.44 3.41 3.38 3.01 2.97 2.66 2.65 2.39 2.38 1.67 1.65
0.09 2.31 1.97 2.17 3.11 0.92 0.11 0.93 10.64 6.15 3.11 3.08 3.10
f1 (ppm)

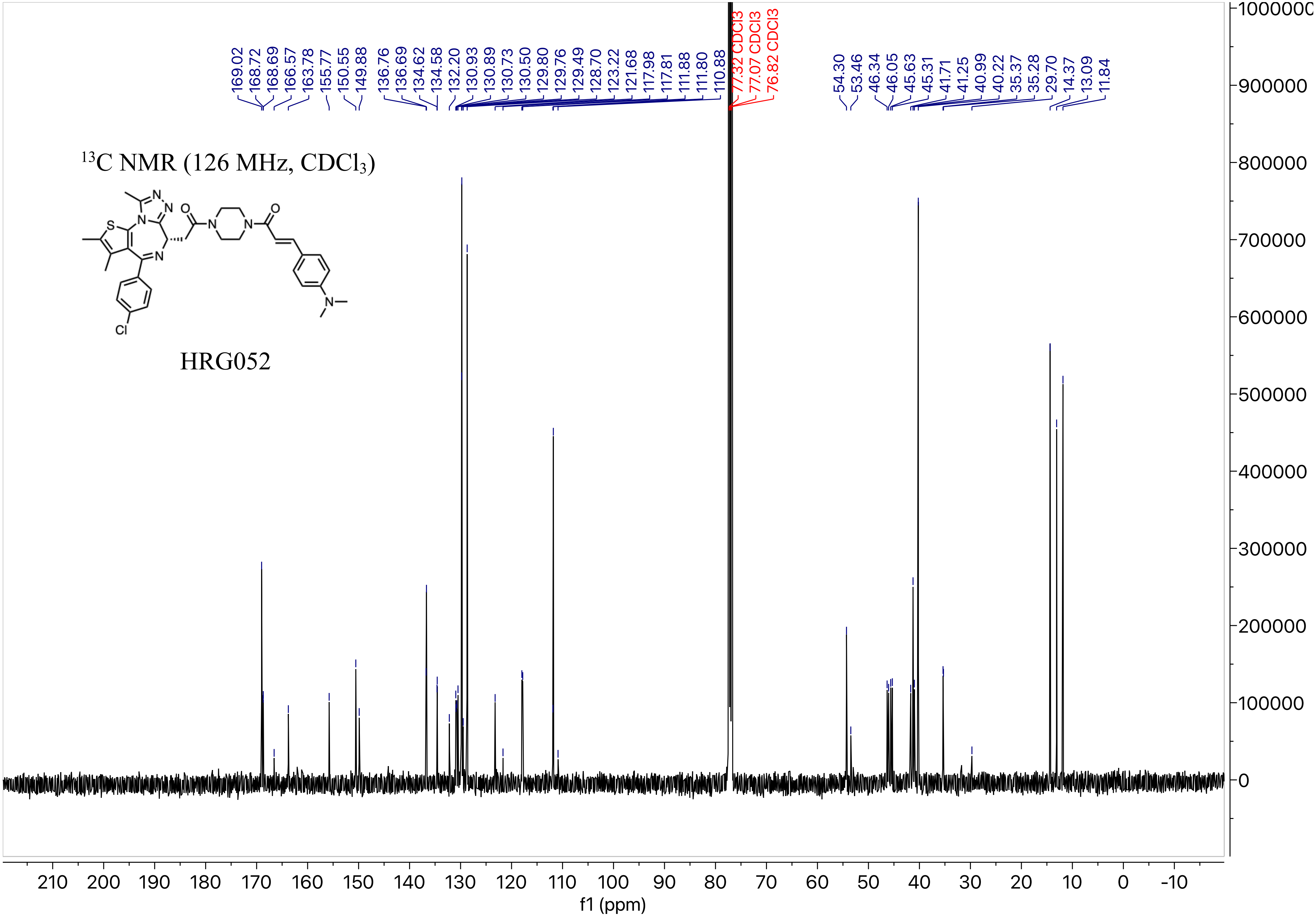
169.02
168.72
168.69
166.57
163.78
155.77
150.55
149.88
136.76
136.69
134.62
134.58
132.20
130.93
130.89
130.73
130.50
129.80
129.76
129.49
128.70
123.22
121.68
117.98
117.81
111.88
111.80
110.88
77.32 CDCl3
77.07 CDCl3
76.82 CDCl3
54.30
53.46
46.34
46.05
45.63
45.31
41.71
41.25
40.99
40.22
35.37
35.28
29.70
14.37
13.09
11.84
$^{13}$C NMR (126 MHz, CDCl$_3$)
HRG052
f1 (ppm)

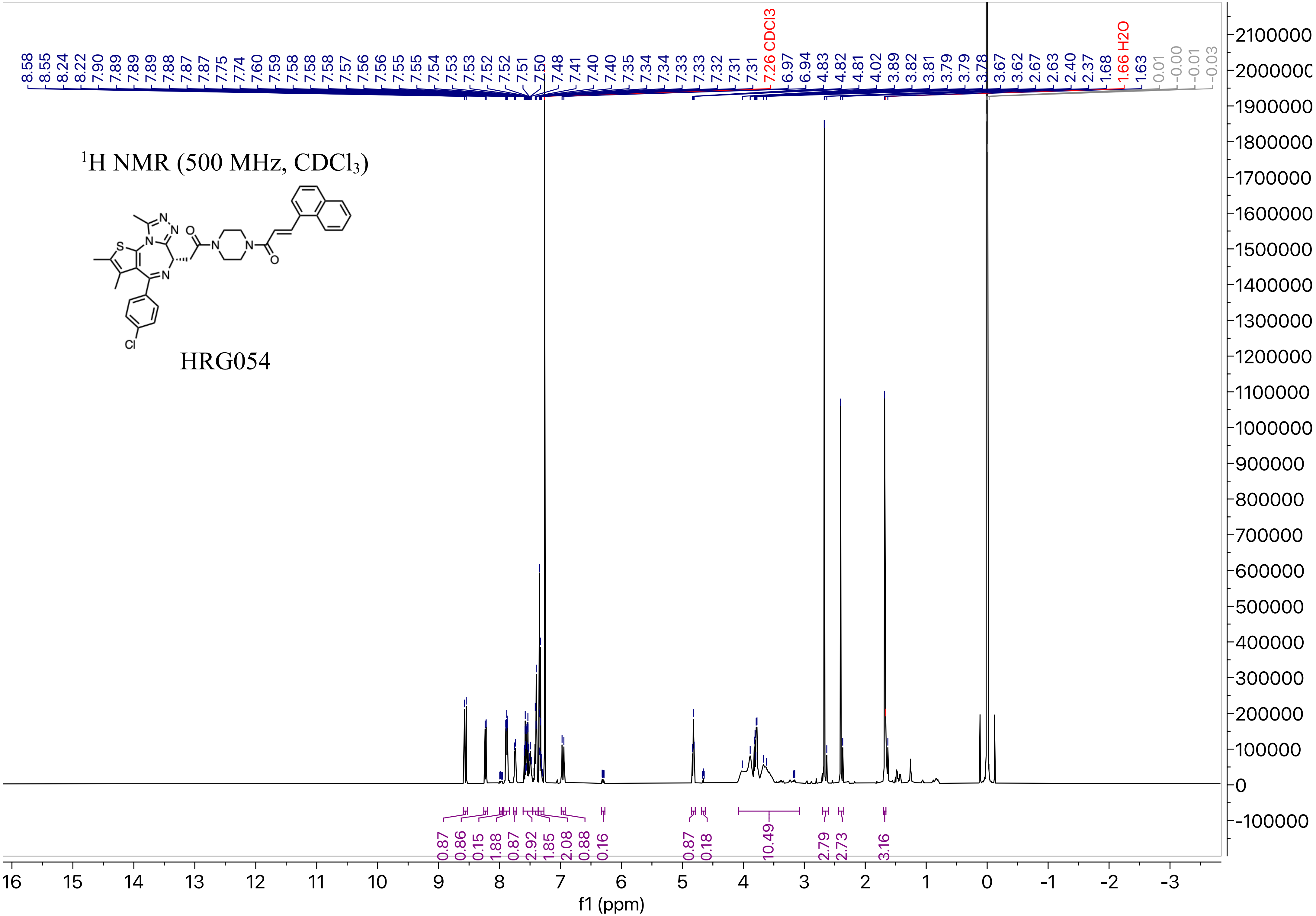

$^1$H NMR (500 MHz, CDCl$_3$)
HRG054
8.58 8.55 8.24 8.22 7.90 7.89 7.89 7.89 7.88 7.87 7.87 7.75 7.74 7.60 7.59 7.58 7.58 7.58 7.57 7.56 7.56 7.55 7.55 7.54 7.53 7.53 7.52 7.52 7.51 7.50 7.48 7.41 7.40 7.40 7.35 7.34 7.34 7.33 7.33 7.32 7.31 7.31
7.26 CDCl3
6.97 6.94 4.83 4.82 4.81 4.02 3.89 3.82 3.81 3.79 3.79 3.78 3.67 3.62 2.67 2.63 2.40 2.37 1.68
1.66 H2O
1.63 0.01 -0.00 -0.01 -0.03
0.87 0.86 0.15 1.88 0.87 2.92 1.85 2.08 0.88 0.16 0.87 0.18 10.49 2.79 2.73 3.16
f1 (ppm)

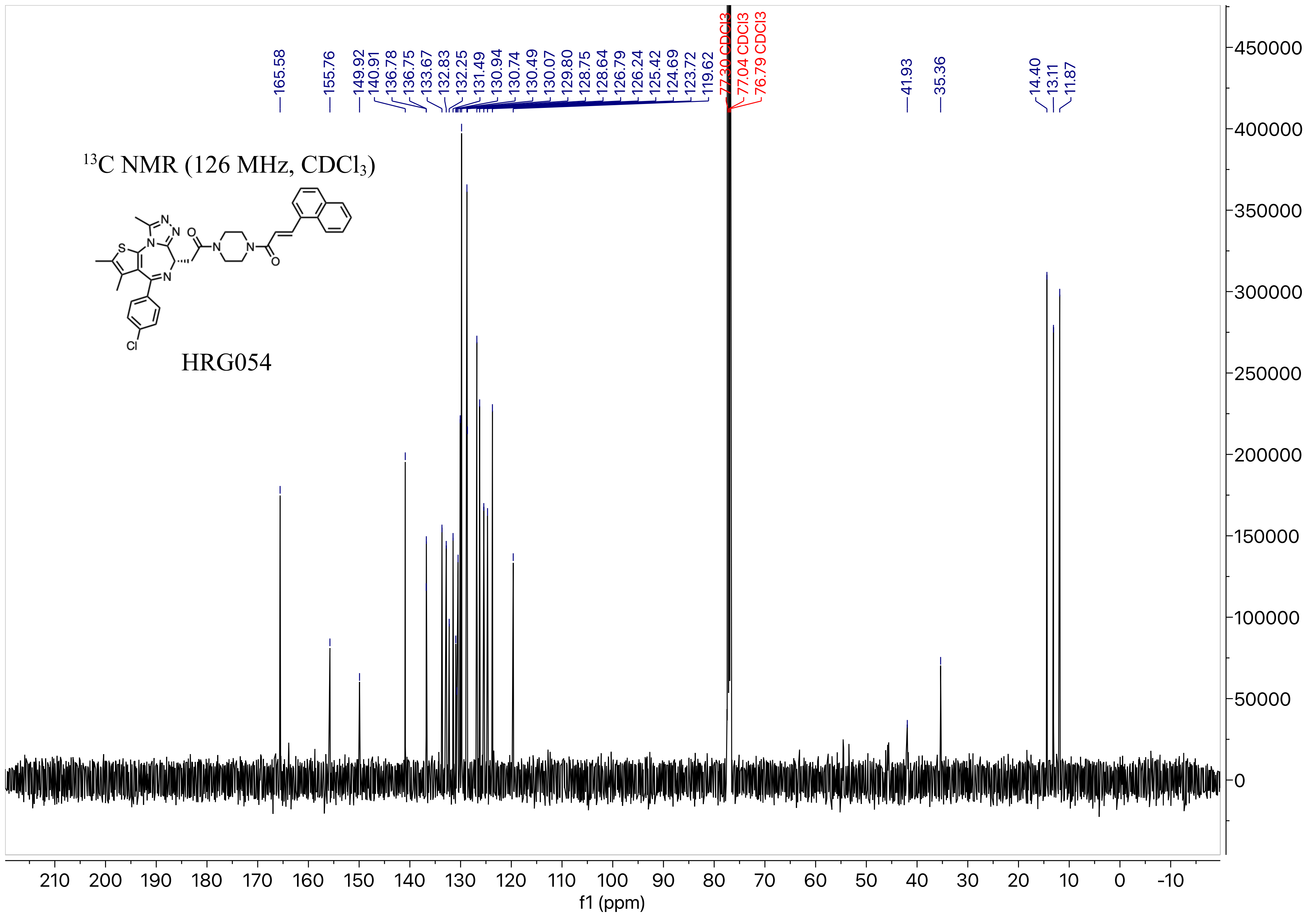

13C NMR (126 MHz, CDCl3)
HRG054
165.58
155.76
149.92
140.91
136.78
136.75
133.67
132.83
132.25
131.49
130.94
130.74
130.49
130.07
129.80
128.75
128.64
126.79
126.24
125.42
124.69
123.72
119.62
77.30 CDCl3
77.04 CDCl3
76.79 CDCl3
41.93
35.36
14.40
13.11
11.87
f1 (ppm)

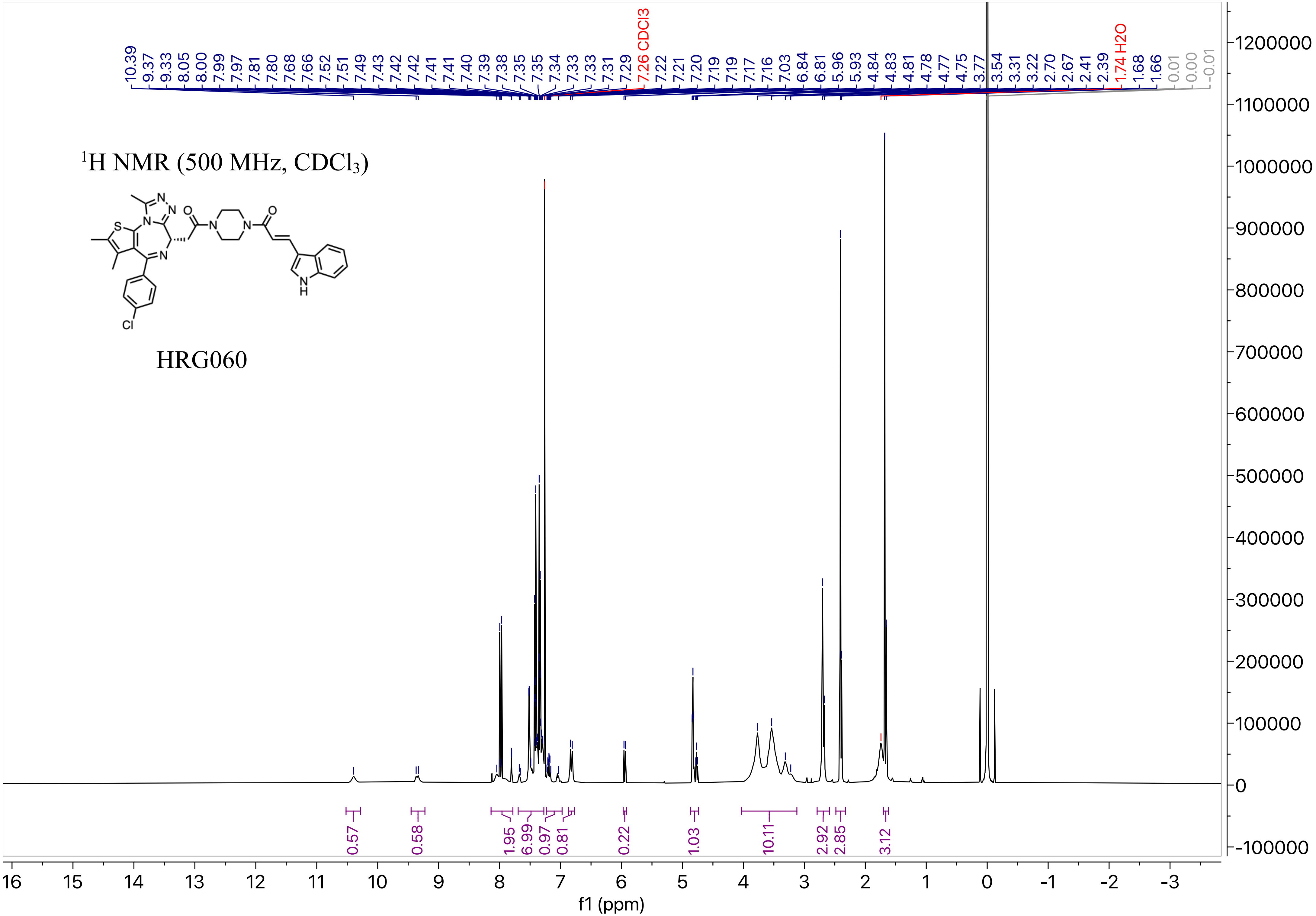

1H NMR (500 MHz, CDCl3)
HRG060
10.39 9.37 9.33 8.05 8.00 7.99 7.97 7.81 7.80 7.68 7.66 7.52 7.51 7.49 7.43 7.42 7.42 7.41 7.41 7.40 7.39 7.38 7.35 7.35 7.34 7.33 7.33 7.31 7.29 7.26 CDCl3 7.22 7.21 7.20 7.19 7.19 7.17 7.16 7.03 6.84 6.81 5.96 5.93 4.84 4.83 4.81 4.78 4.77 4.75 3.77 3.54 3.31 3.22 2.70 2.67 2.41 2.39 1.74 H2O 1.68 1.66 0.01 0.00 -0.01
0.57 0.58 1.95 6.99 0.97 0.81 0.22 1.03 10.11 2.92 2.85 3.12
f1 (ppm)

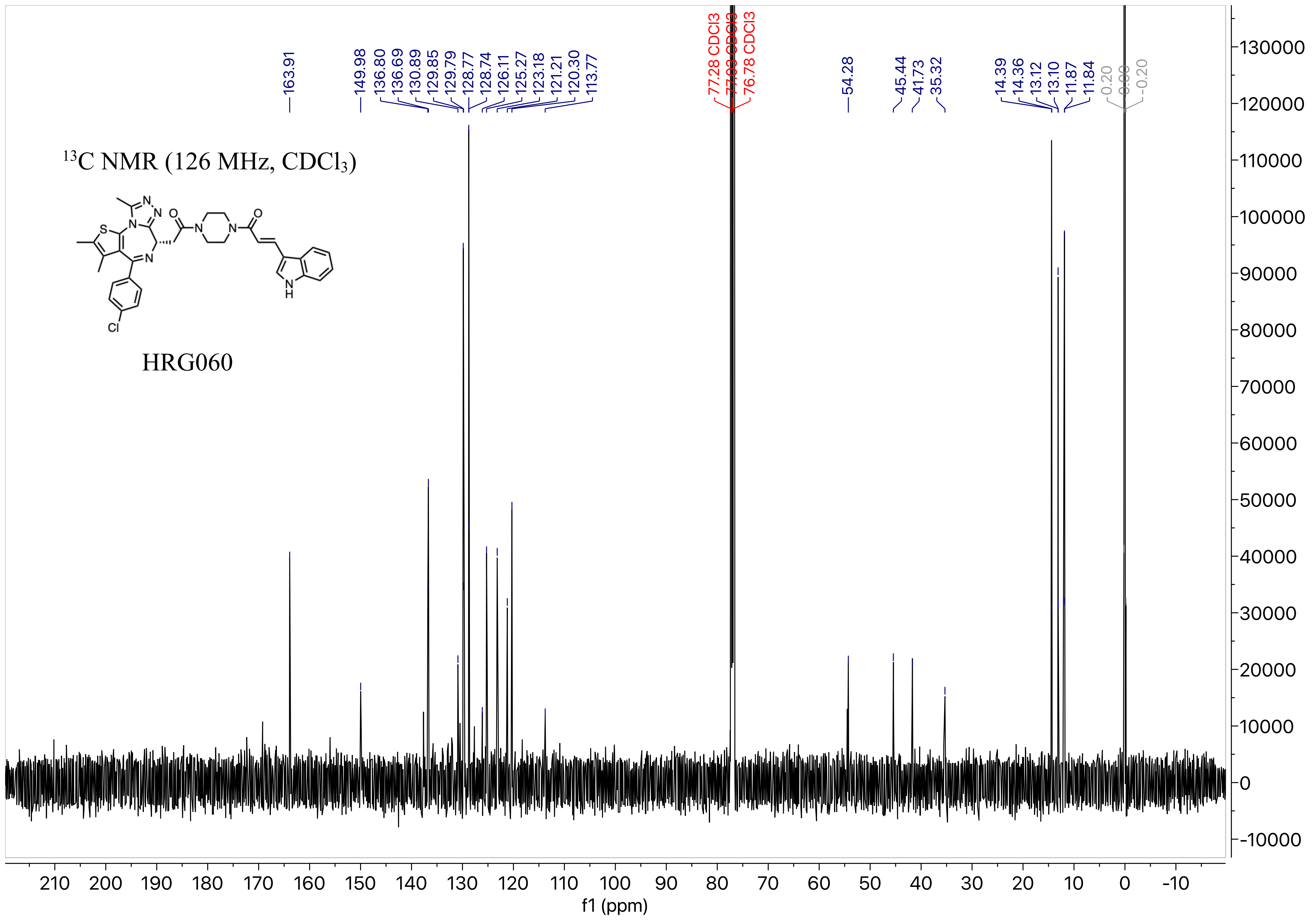

$^{13}$C NMR (126 MHz, CDCl$_3$)
HRG060
163.91
149.98
136.80
136.69
130.89
129.85
129.79
128.77
128.74
126.11
125.27
123.18
121.21
120.30
113.77
77.28 CDCl3
77.03 CDCl3
76.78 CDCl3
54.28
45.44
41.73
35.32
14.39
14.36
13.12
13.10
11.87
11.84
0.20
0.00
-0.20
f1 (ppm)

[1]H NMR (500 MHz, $CDCl_3$)

HRG061

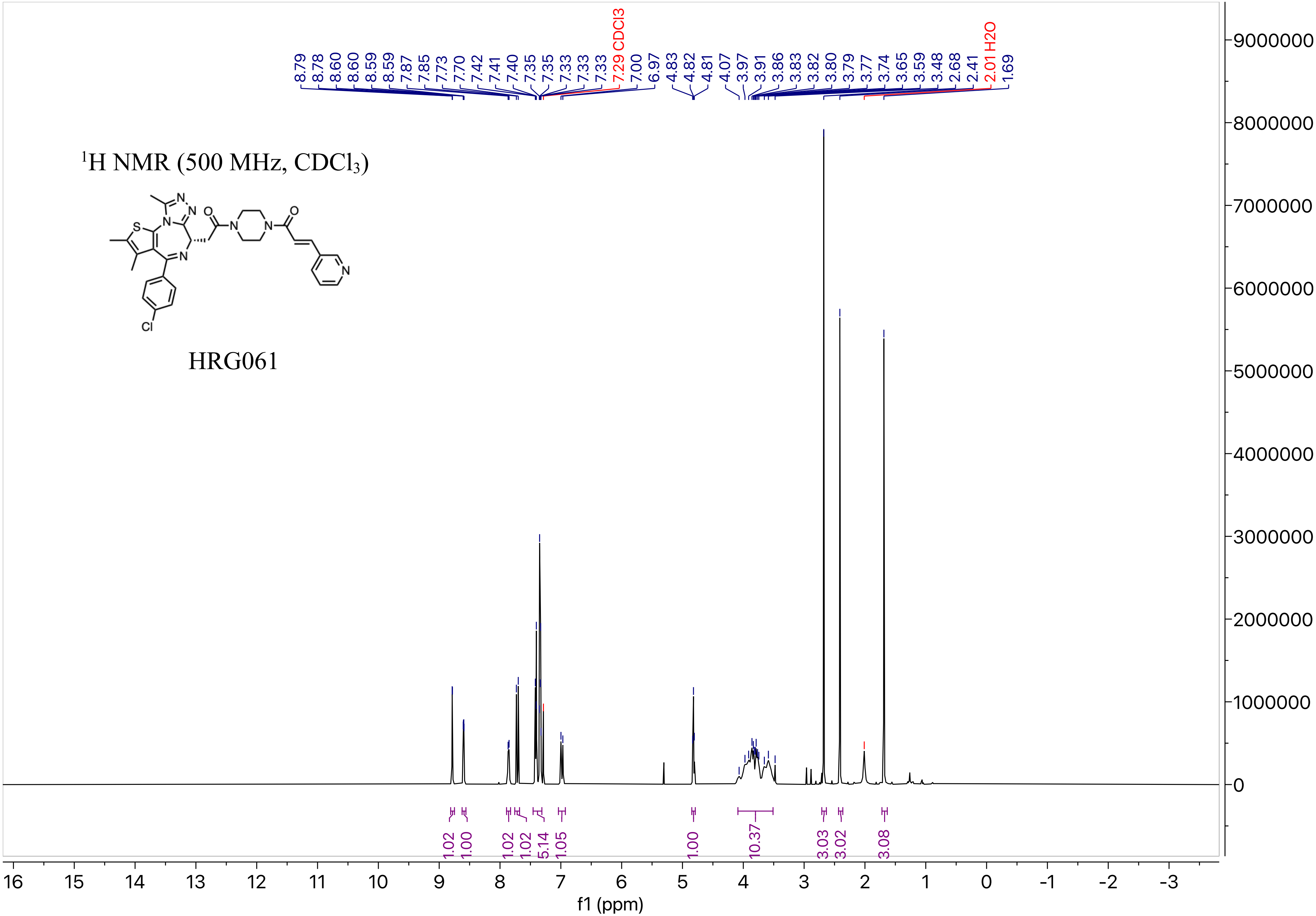

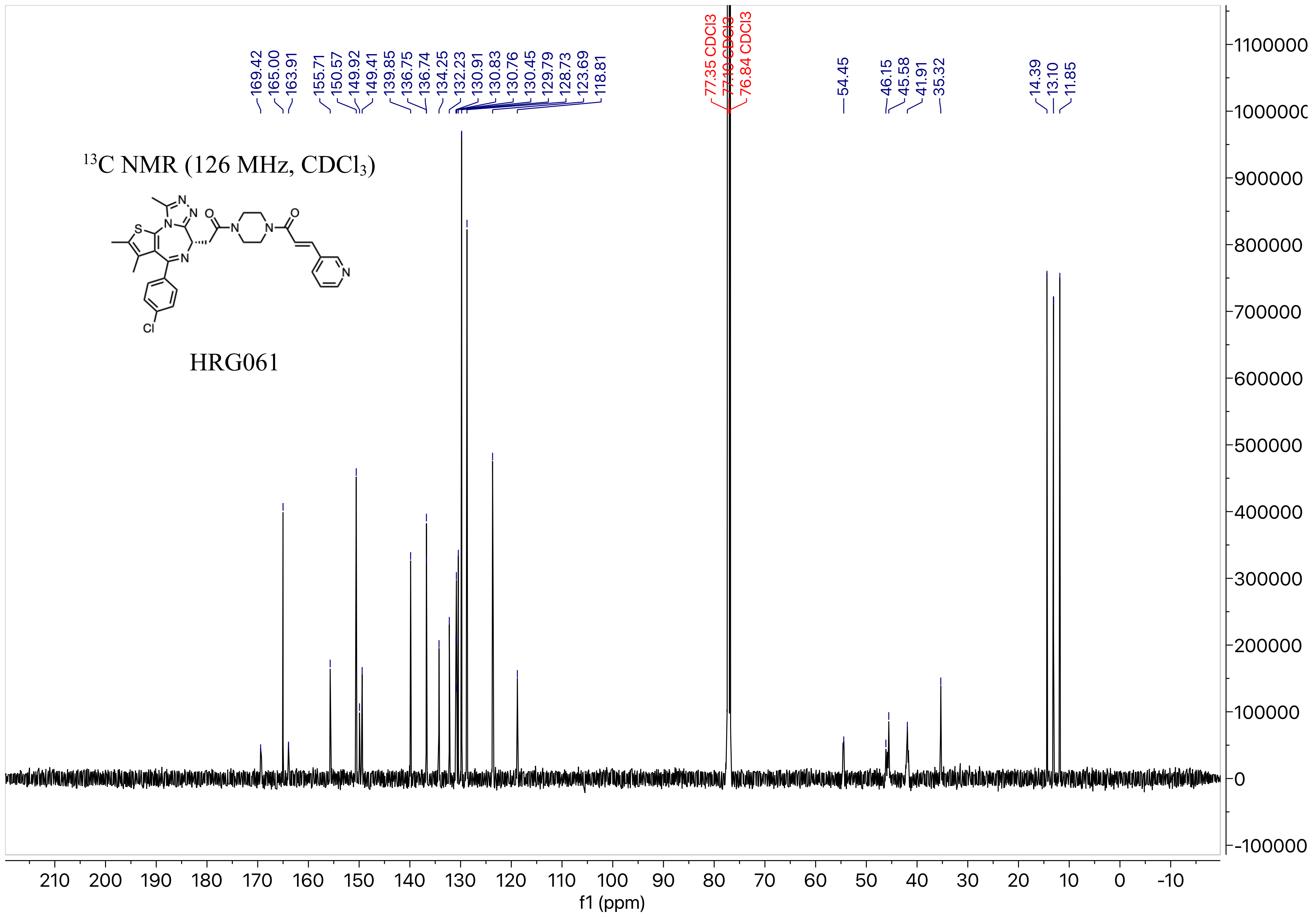

169.42
165.00
163.91
155.71
150.57
149.92
149.41
139.85
136.75
136.74
134.25
132.23
130.91
130.83
130.76
130.45
129.79
128.73
123.69
118.81
77.35 CDCl3
77.10 CDCl3
76.84 CDCl3
54.45
46.15
45.58
41.91
35.32
14.39
13.10
11.85
13C NMR (126 MHz, CDCl3)
HRG061
f1 (ppm)

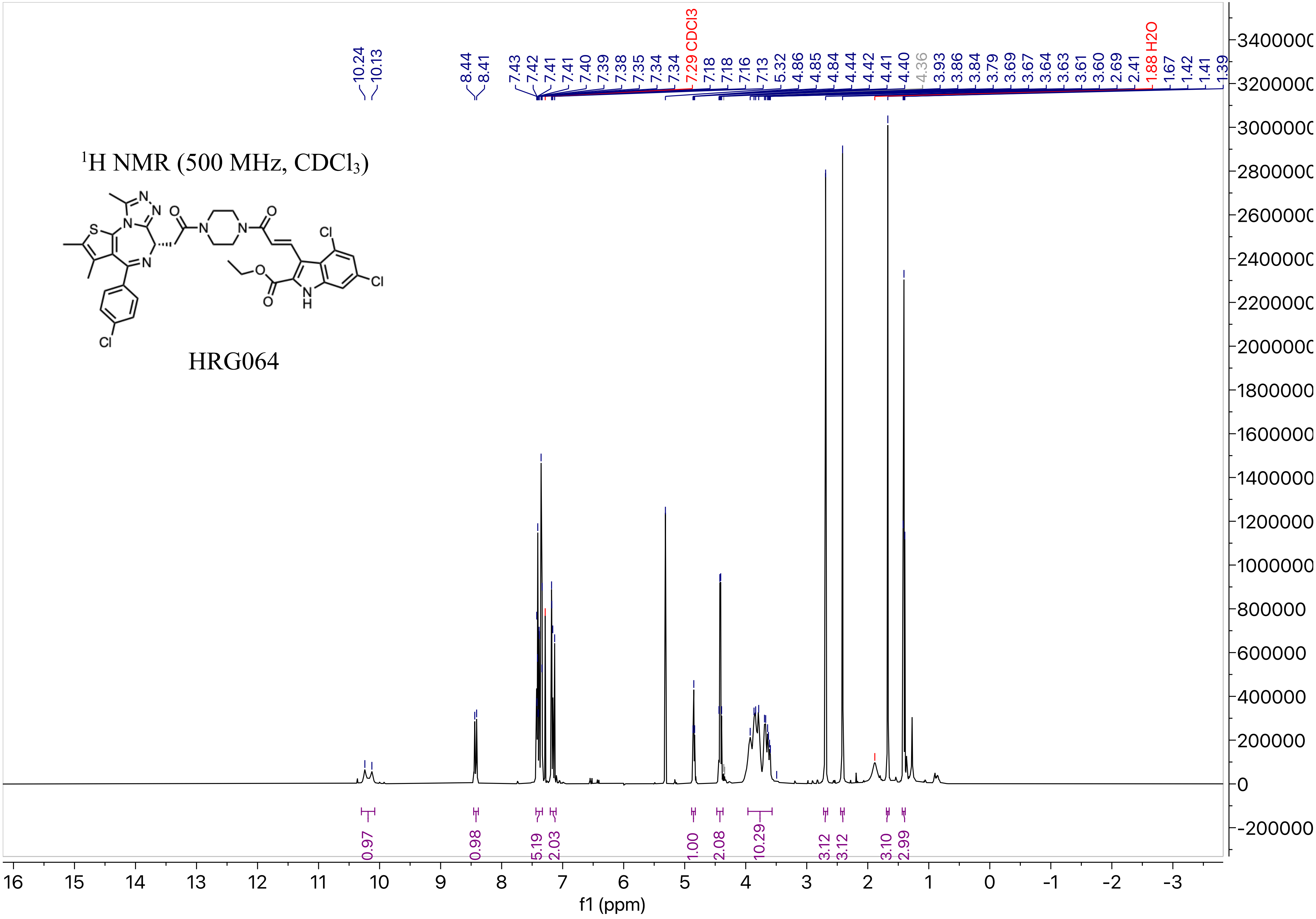

¹H NMR (500 MHz, CDCl₃)
HRG064
10.24 10.13 8.44 8.41 7.43 7.42 7.41 7.41 7.40 7.39 7.38 7.35 7.34 7.34 7.29 CDCl3 7.18 7.18 7.16 7.13 5.32 4.86 4.85 4.84 4.44 4.42 4.41 4.40 4.36 3.93 3.86 3.84 3.79 3.69 3.67 3.64 3.63 3.61 3.60 2.69 2.41 1.88 H2O 1.67 1.42 1.41 1.39
0.97 0.98 5.19 2.03 1.00 2.08 10.29 3.12 3.12 3.10 2.99
f1 (ppm)

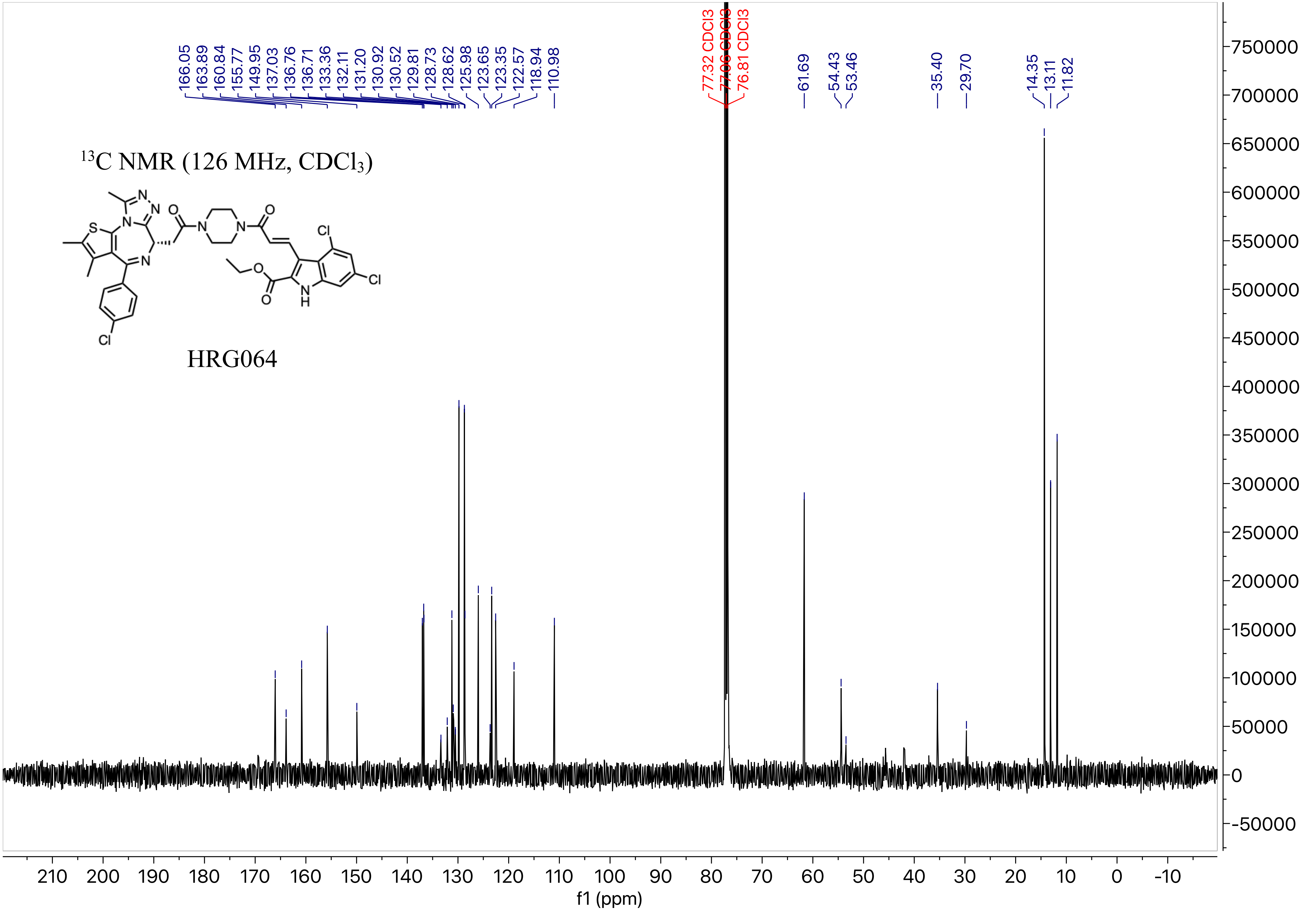

13C NMR (126 MHz, CDCl3)
HRG064
166.05
163.89
160.84
155.77
149.95
137.03
136.76
136.71
133.36
132.11
131.20
130.92
130.52
129.81
128.73
128.62
125.98
123.65
123.35
122.57
118.94
110.98
77.32 CDCl3
77.06 CDCl3
76.81 CDCl3
61.69
54.43
53.46
35.40
29.70
14.35
13.11
11.82
f1 (ppm)

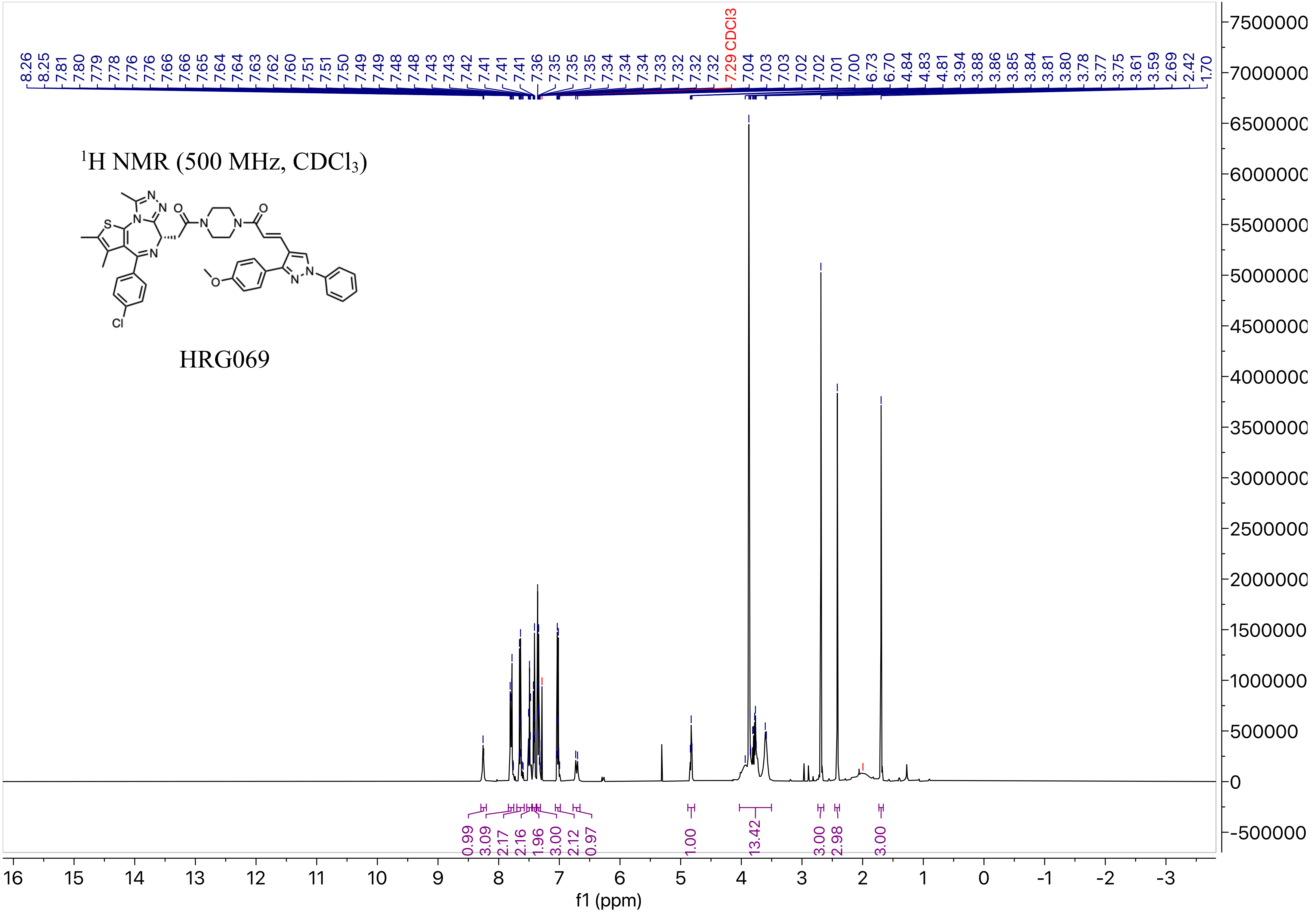
1H NMR (500 MHz, CDCl3)
HRG069
8.26 8.25 7.81 7.80 7.79 7.78 7.76 7.76 7.66 7.66 7.65 7.64 7.64 7.63 7.62 7.60 7.51 7.51 7.50 7.49 7.49 7.48 7.48 7.43 7.43 7.42 7.41 7.41 7.41 7.36 7.35 7.35 7.35 7.34 7.34 7.34 7.33 7.32 7.32 7.32 7.29 CDCl3 7.04 7.03 7.03 7.02 7.02 7.01 7.00 6.73 6.70 4.84 4.83 4.81 3.94 3.88 3.86 3.85 3.84 3.81 3.80 3.78 3.77 3.75 3.61 3.59 2.69 2.42 1.70
0.99 3.09 2.17 2.16 1.96 3.00 2.12 0.97 1.00 13.42 3.00 2.98 3.00
f1 (ppm)

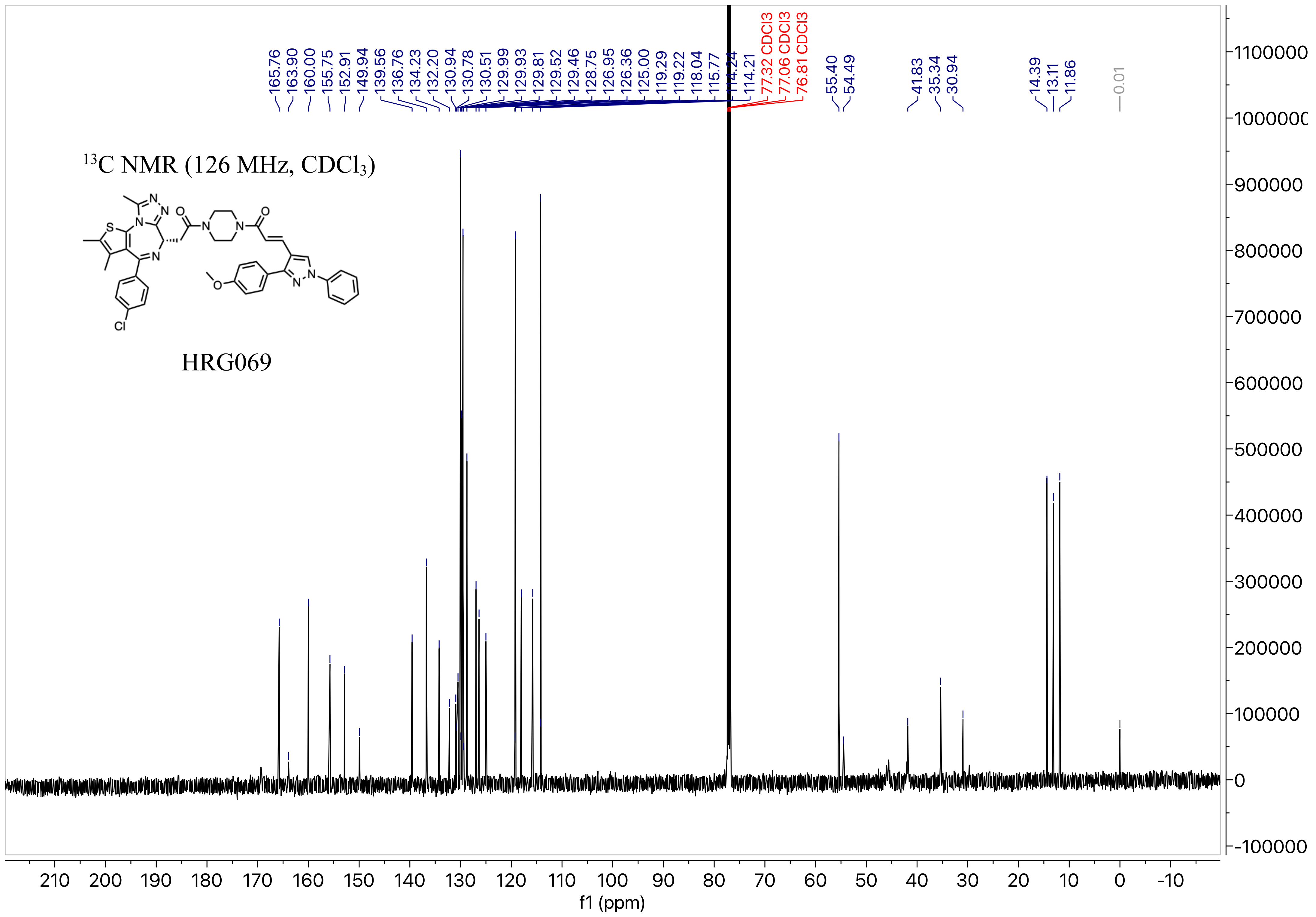

13C NMR (126 MHz, CDCl3)
HRG069
165.76
163.90
160.00
155.75
152.91
149.94
139.56
136.76
134.23
132.20
130.94
130.78
130.51
129.99
129.93
129.81
129.52
129.46
128.75
126.95
126.36
125.00
119.29
119.22
118.04
115.77
114.24
114.21
77.32 CDCl3
77.06 CDCl3
76.81 CDCl3
55.40
54.49
41.83
35.34
30.94
14.39
13.11
11.86
-0.01
f1 (ppm)

[1]H NMR (500 MHz, $CDCl_3$)

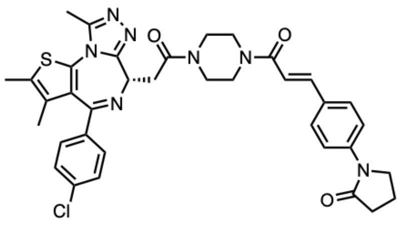

HRG070

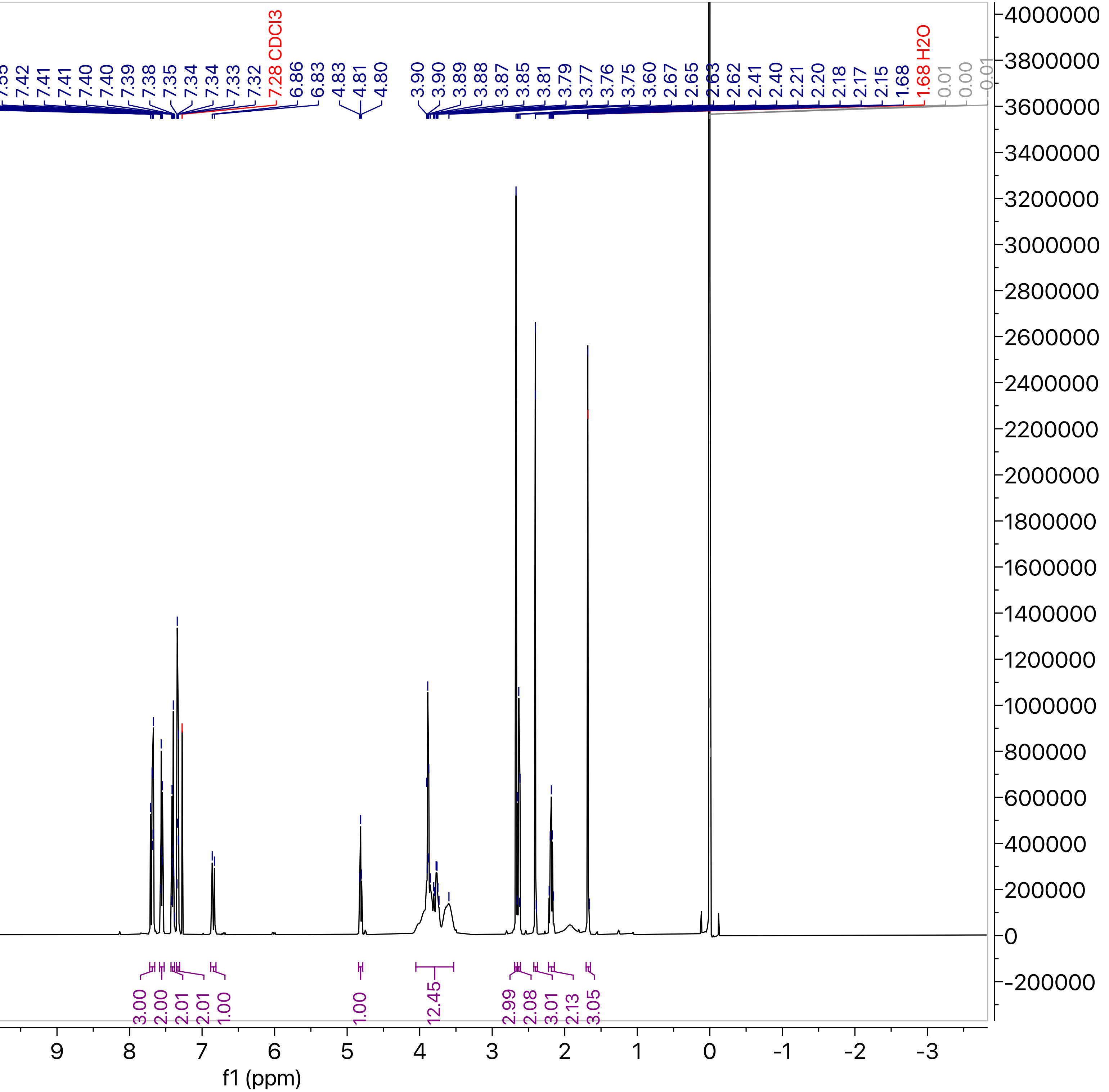

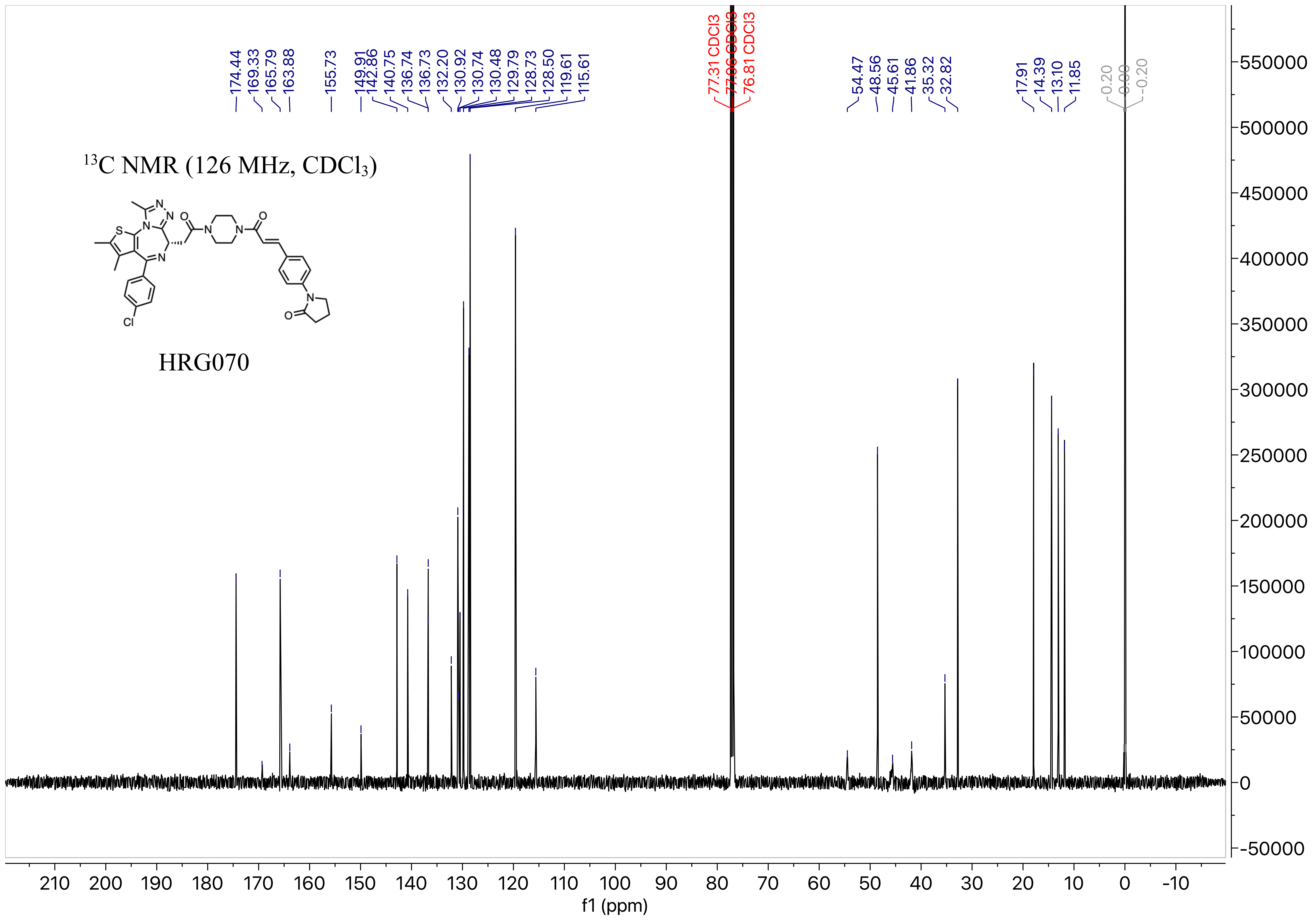

13C NMR (126 MHz, CDCl3)
HRG070
174.44
169.33
165.79
163.88
155.73
149.91
142.86
140.75
136.74
136.73
132.20
130.92
130.74
130.48
129.79
128.73
128.50
119.61
115.61
77.31 CDCl3
77.06 CDCl3
76.81 CDCl3
54.47
48.56
45.61
41.86
35.32
32.82
17.91
14.39
13.10
11.85
0.20
0.00
-0.20
f1 (ppm)

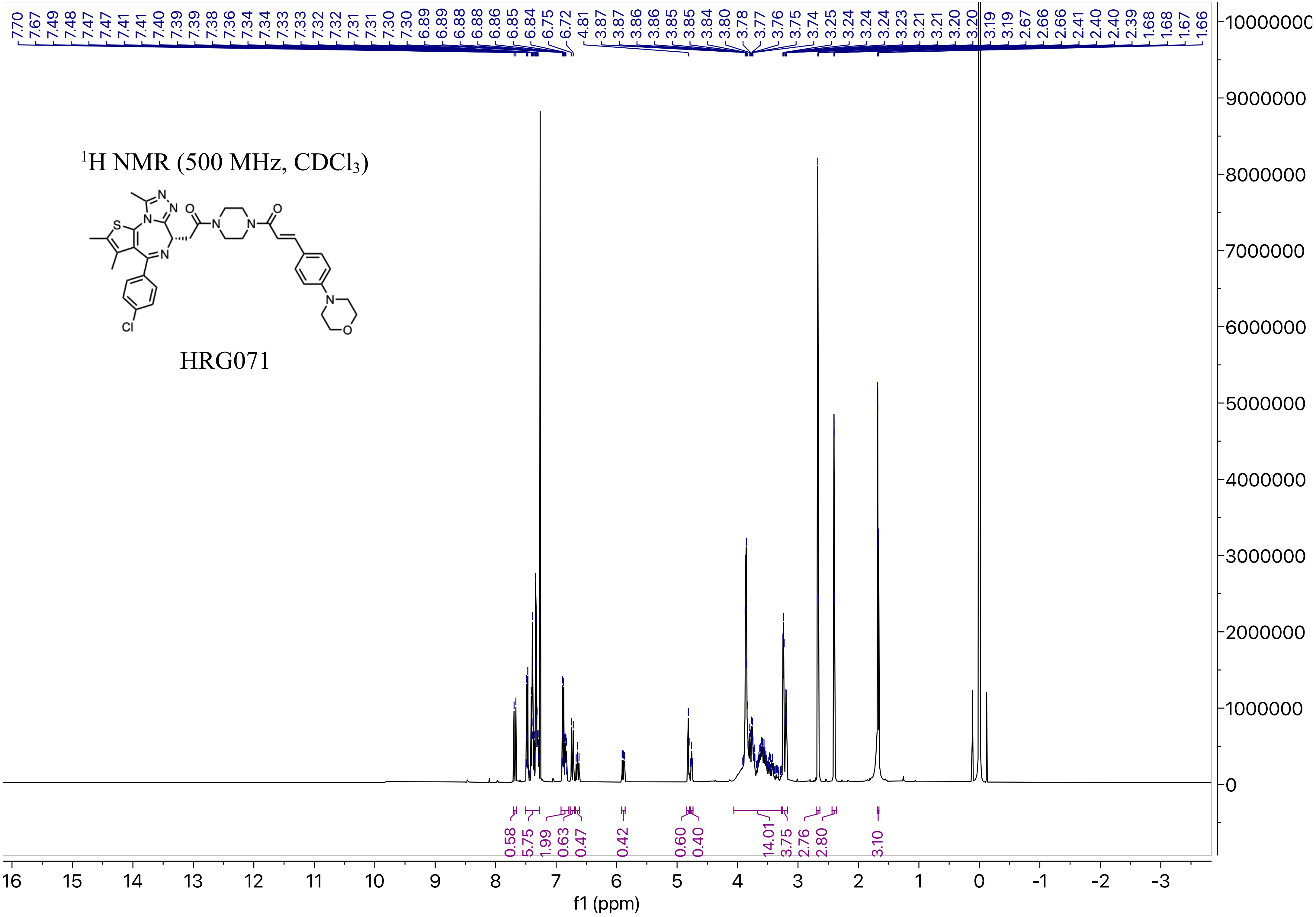

1H NMR (500 MHz, CDCl3)
HRG071
f1 (ppm)

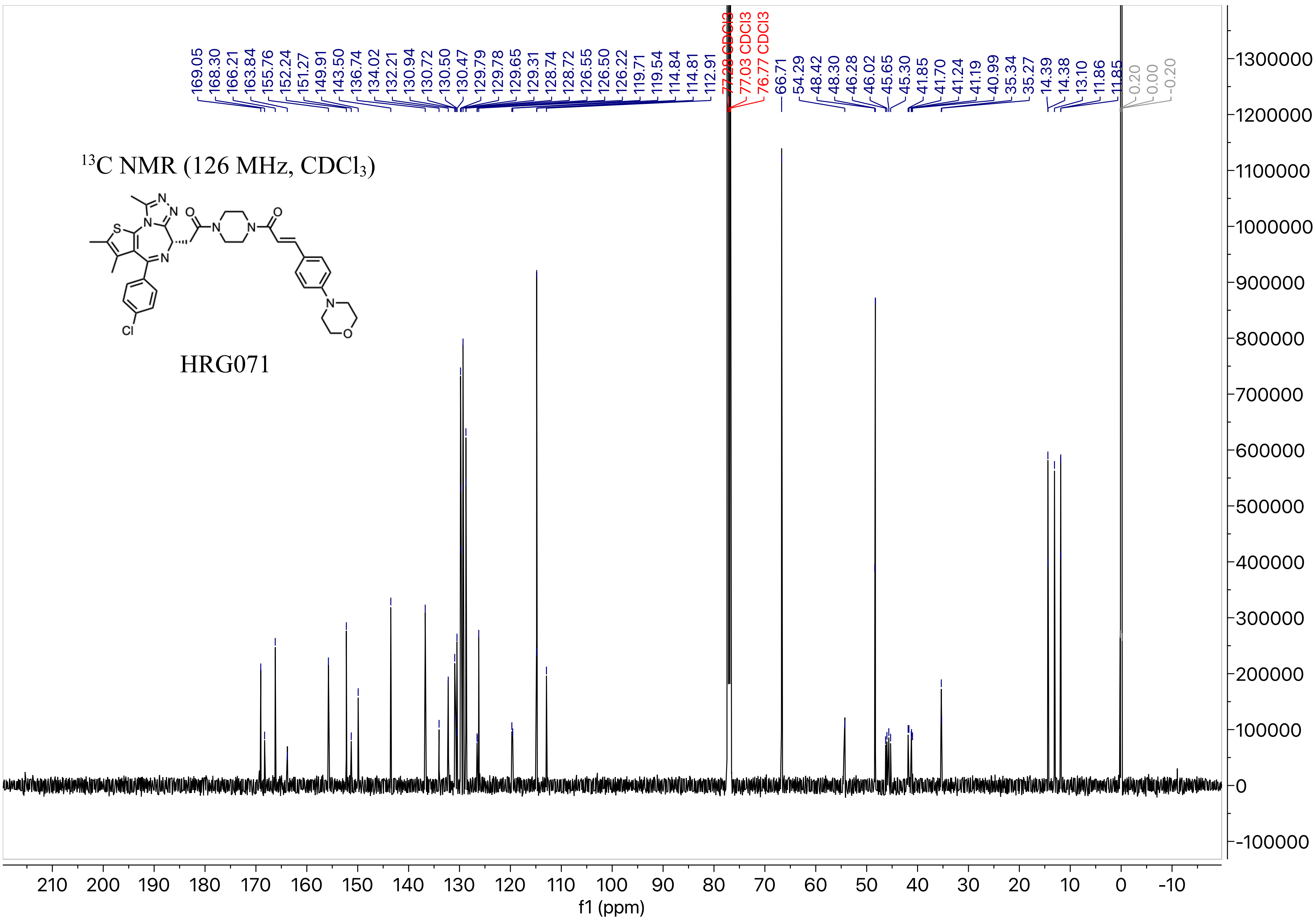

13C NMR (126 MHz, CDCl3)
HRG071
169.05
168.30
166.21
163.84
155.76
152.24
151.27
149.91
143.50
136.74
134.02
132.21
130.94
130.72
130.50
130.47
129.79
129.78
129.65
129.31
128.74
128.72
126.55
126.50
126.22
119.71
119.54
114.84
114.81
112.91
77.28 CDCl3
77.03 CDCl3
76.77 CDCl3
66.71
54.29
48.42
48.30
46.28
46.02
45.65
45.30
41.85
41.70
41.24
41.19
40.99
35.34
35.27
14.39
14.38
13.10
11.86
11.85
0.20
0.00
-0.20
f1 (ppm)

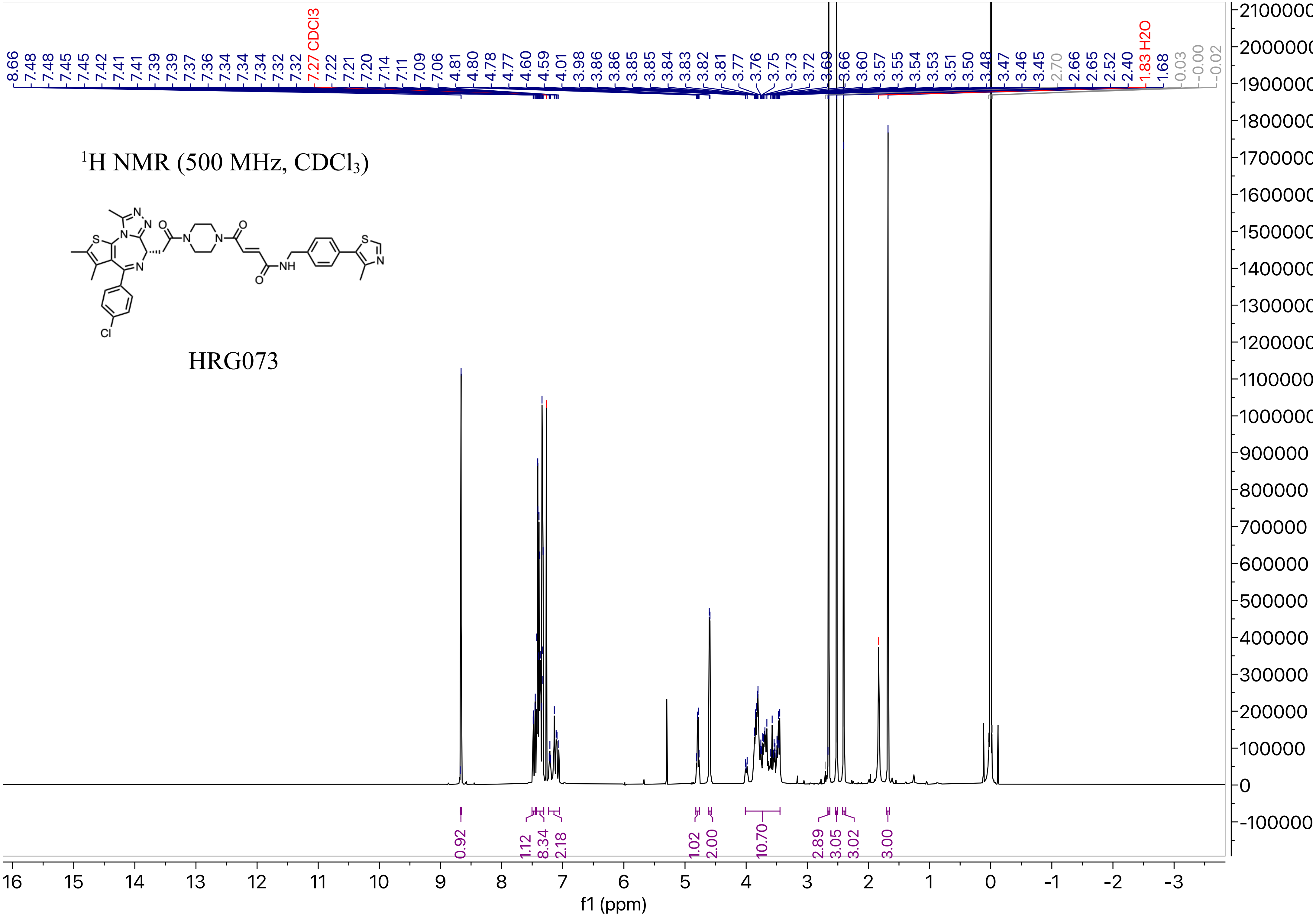

$^1H$ NMR (500 MHz, $CDCl_3$)
HRG073
f1 (ppm)
7.27 CDCl3
1.83 H2O

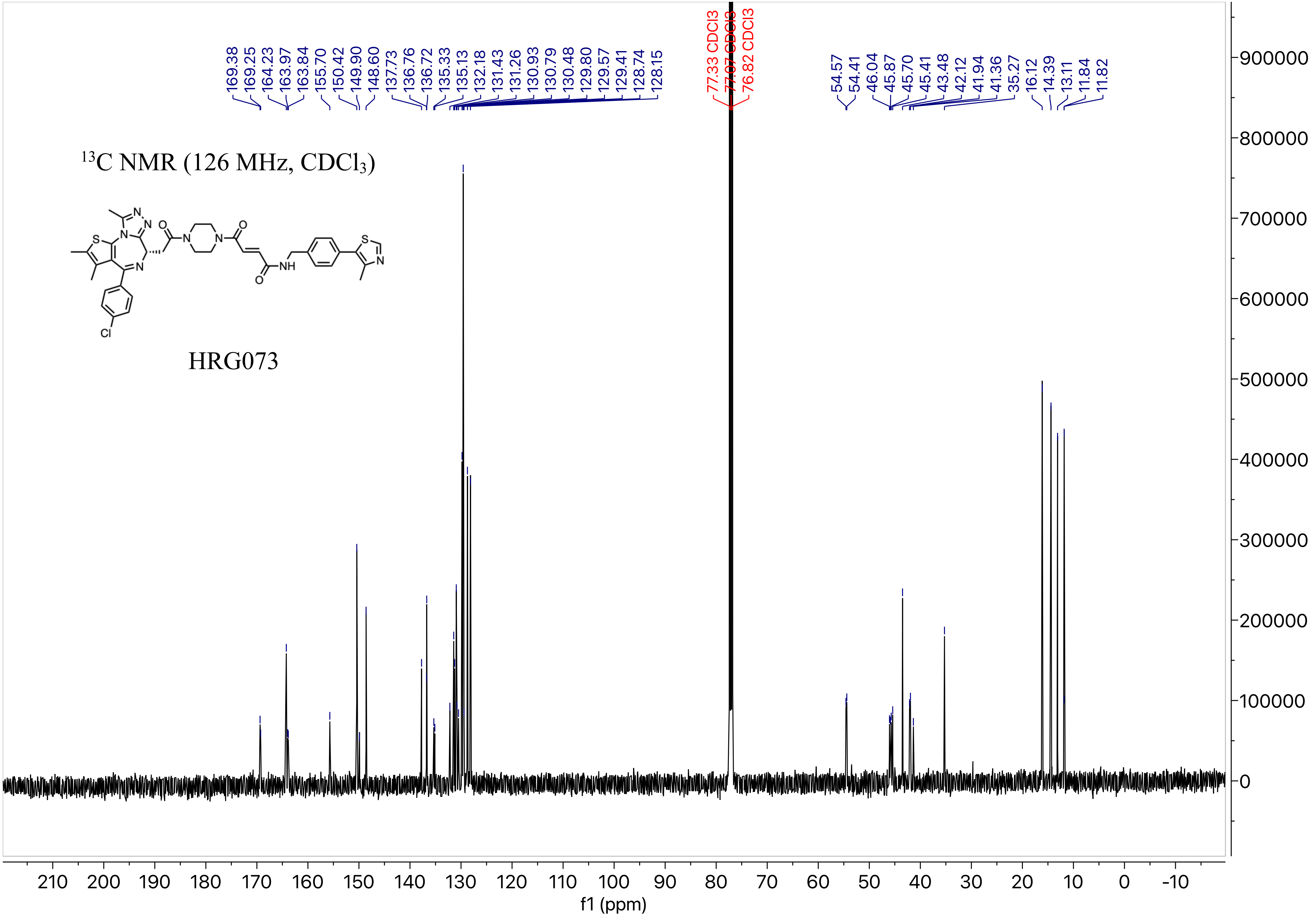

$^{13}$C NMR (126 MHz, CDCl$_3$)
HRG073
169.38
169.25
164.23
163.97
163.84
155.70
150.42
149.90
148.60
137.73
136.76
136.72
135.33
135.13
132.18
131.43
131.26
130.93
130.79
130.48
129.80
129.57
129.41
128.74
128.15
77.33 CDCl3
77.07 CDCl3
76.82 CDCl3
54.57
54.41
46.04
45.87
45.70
45.41
43.48
42.12
41.94
41.36
35.27
16.12
14.39
13.11
11.84
11.82
f1 (ppm)

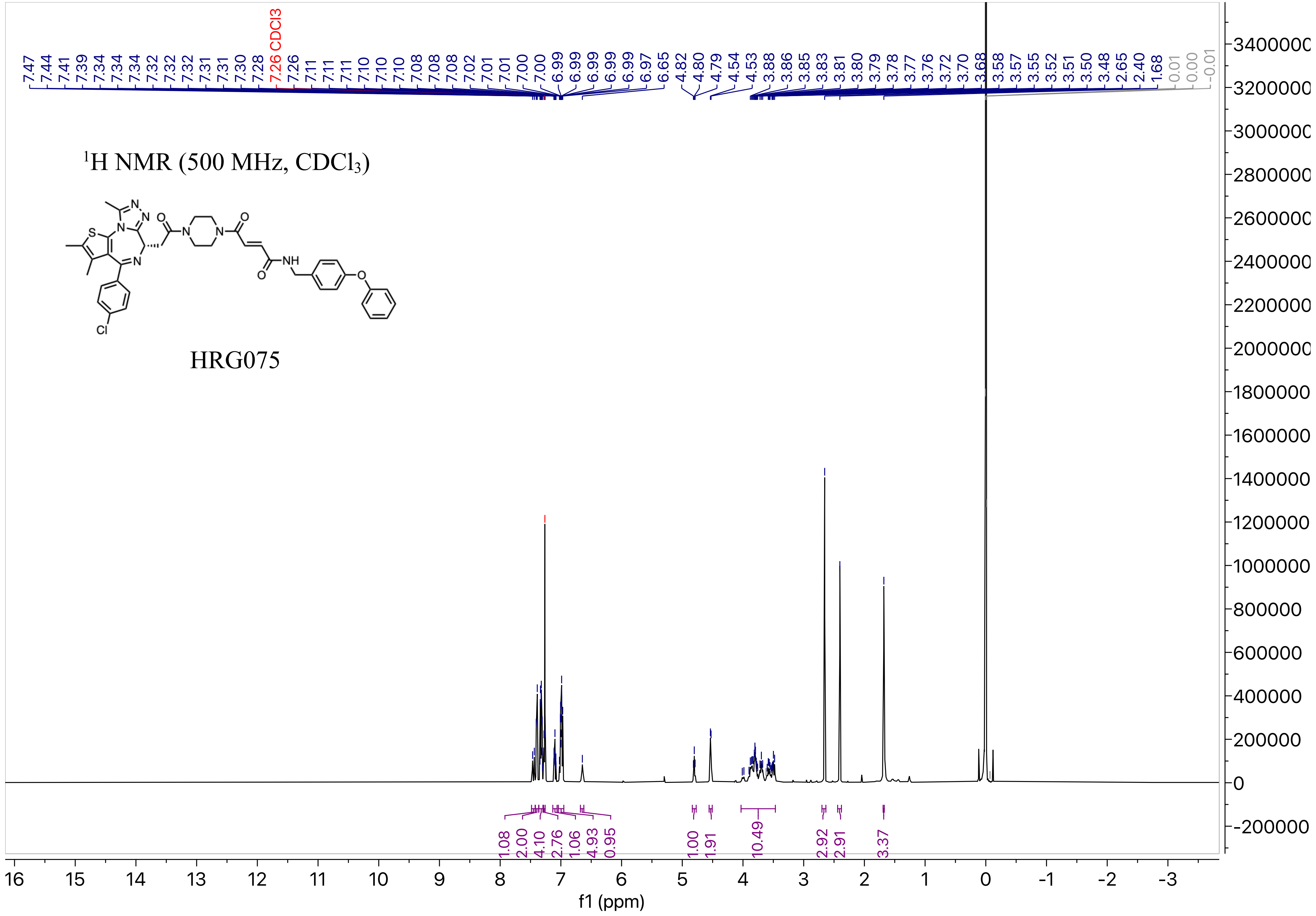

1H NMR (500 MHz, CDCl3)
HRG075
S
N
N
N
O
O
N
N
N
NH
O
O
Cl
7.47 7.44 7.41 7.39 7.34 7.34 7.34 7.32 7.32 7.32 7.31 7.31 7.30 7.28 7.26 CDCl3 7.26 7.11 7.11 7.11 7.10 7.10 7.10 7.08 7.08 7.08 7.02 7.01 7.01 7.00 7.00 6.99 6.99 6.99 6.99 6.99 6.97 6.65 4.82 4.80 4.79 4.54 4.53 3.88 3.86 3.85 3.83 3.81 3.80 3.79 3.78 3.77 3.76 3.72 3.70 3.68 3.58 3.57 3.55 3.52 3.51 3.50 3.48 2.65 2.40 1.68 0.01 0.00 -0.01
1.08 2.00 4.10 2.76 1.06 4.93 0.95 1.00 1.91 10.49 2.92 2.91 3.37
3400000 3200000 3000000 2800000 2600000 2400000 2200000 2000000 1800000 1600000 1400000 1200000 1000000 800000 600000 400000 200000 0 -200000
16 15 14 13 12 11 10 9 8 7 6 5 4 3 2 1 0 -1 -2 -3
f1 (ppm)

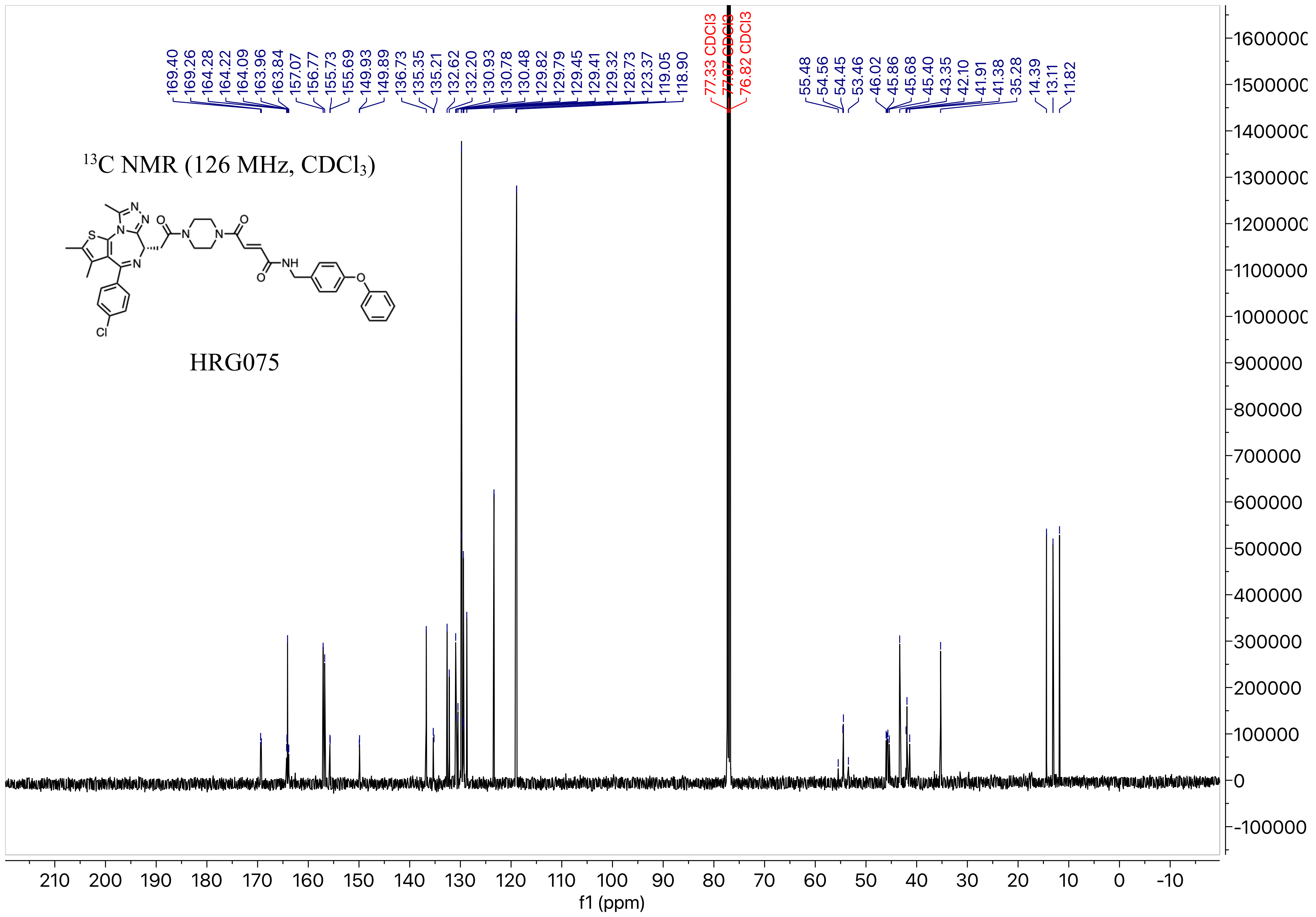

169.40
169.26
164.28
164.22
164.09
163.96
163.84
157.07
156.77
155.73
155.69
149.93
149.89
136.73
135.35
135.21
132.62
132.20
130.93
130.78
130.48
129.82
129.79
129.45
129.41
129.32
128.73
123.37
119.05
118.90
77.33 CDCl3
77.07 CDCl3
76.82 CDCl3
55.48
54.56
54.45
53.46
46.02
45.86
45.68
45.40
43.35
42.10
41.91
41.38
35.28
14.39
13.11
11.82
13C NMR (126 MHz, CDCl3)
HRG075
f1 (ppm)

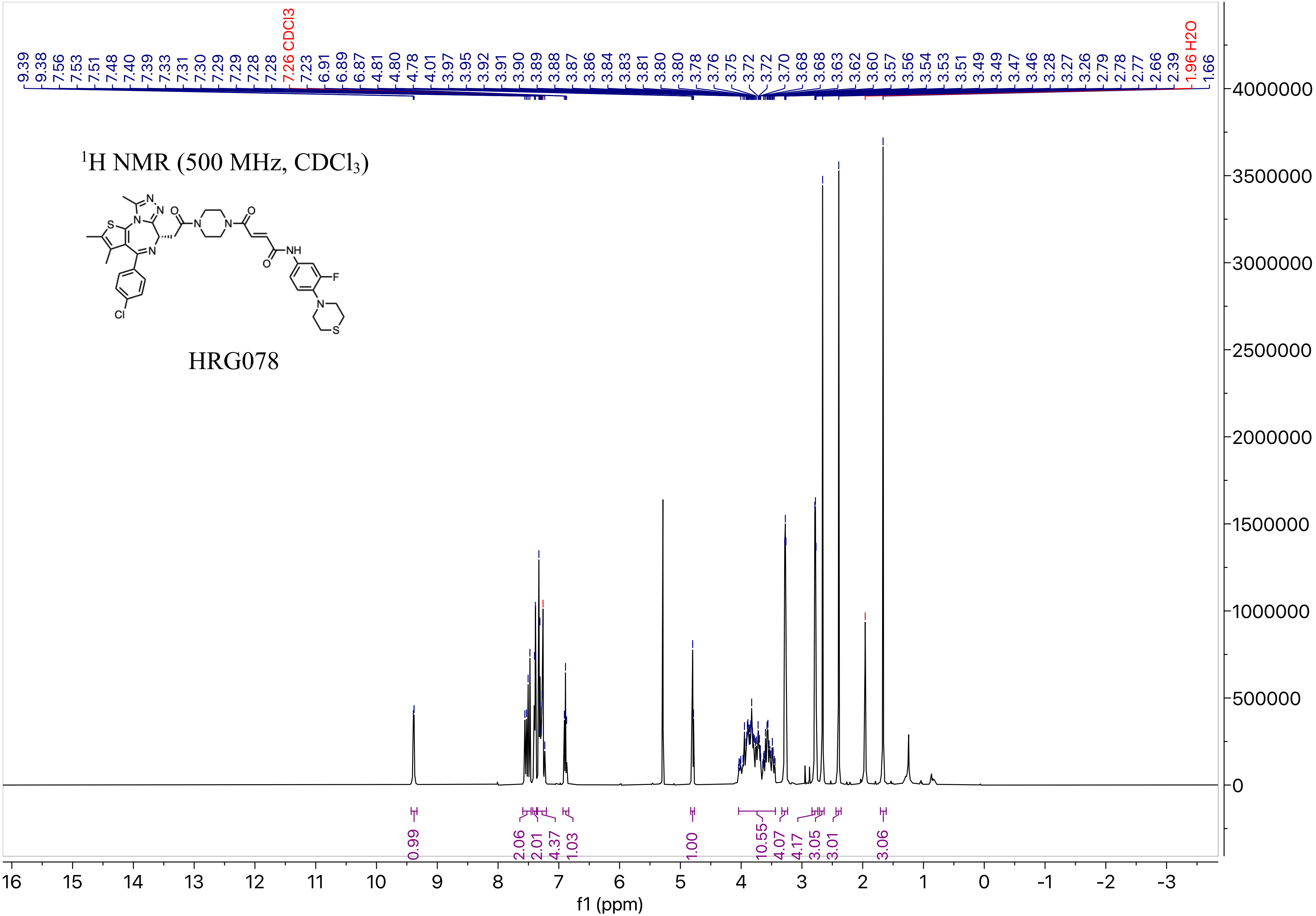

9.39 9.38 7.56 7.53 7.51 7.48 7.40 7.39 7.33 7.31 7.30 7.29 7.29 7.28 7.28 7.26 CDCl3 7.23 6.91 6.89 6.87 4.81 4.80 4.78 4.01 3.97 3.95 3.92 3.91 3.90 3.89 3.88 3.87 3.86 3.84 3.83 3.81 3.80 3.80 3.78 3.76 3.75 3.72 3.72 3.70 3.68 3.68 3.63 3.62 3.60 3.57 3.56 3.54 3.53 3.51 3.49 3.49 3.47 3.46 3.28 3.27 3.26 2.79 2.78 2.77 2.66 2.39 1.96 H2O 1.66
1H NMR (500 MHz, CDCl3)
HRG078
0.99 2.06 2.01 4.37 1.03 1.00 10.55 4.07 4.17 3.05 3.01 3.06
f1 (ppm)

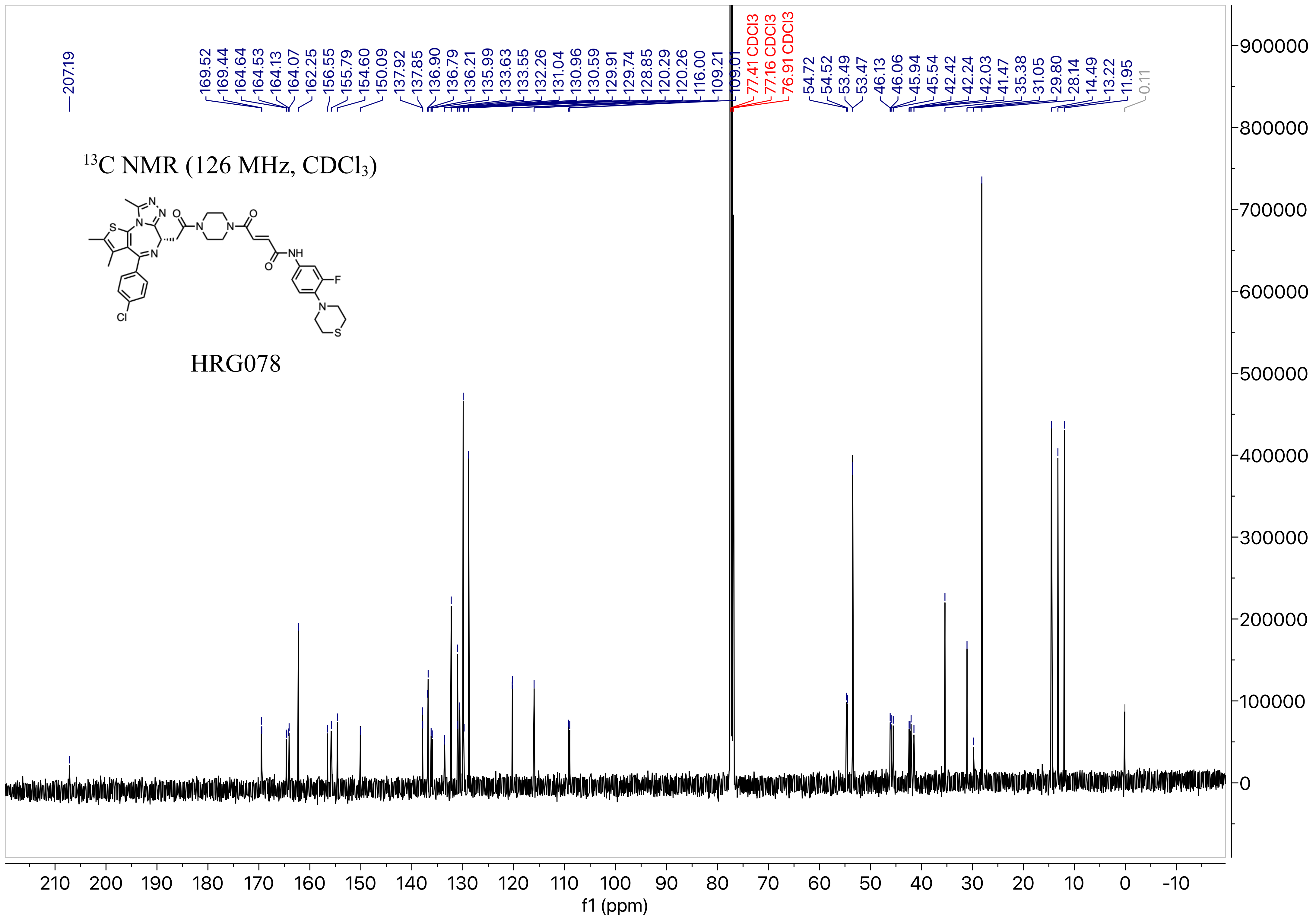

207.19
169.52
169.44
164.64
164.53
164.13
164.07
162.25
156.55
155.79
154.60
150.09
137.92
137.85
136.90
136.79
136.21
135.99
133.63
133.55
132.26
131.04
130.96
130.59
129.91
129.74
128.85
120.29
120.26
116.00
109.21
109.01
77.41 CDCl3
77.16 CDCl3
76.91 CDCl3
54.72
54.52
53.49
53.47
46.13
46.06
45.94
45.54
42.42
42.24
42.03
41.47
35.38
31.05
29.80
28.14
14.49
13.22
11.95
0.11
$^{13}$C NMR (126 MHz, CDCl$_3$)
HRG078
f1 (ppm)

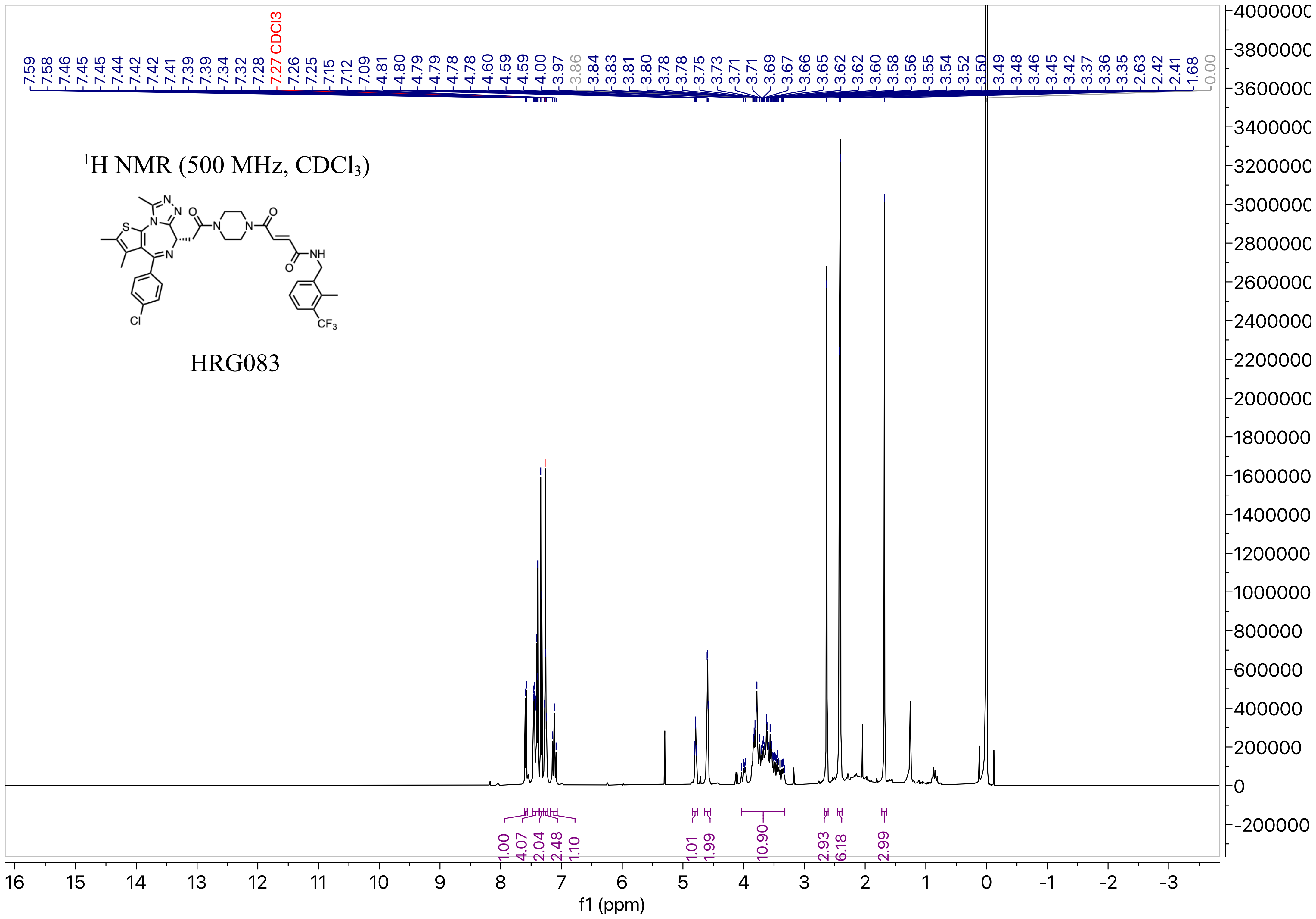

$^1$H NMR (500 MHz, CDCl$_3$)
HRG083
f1 (ppm)

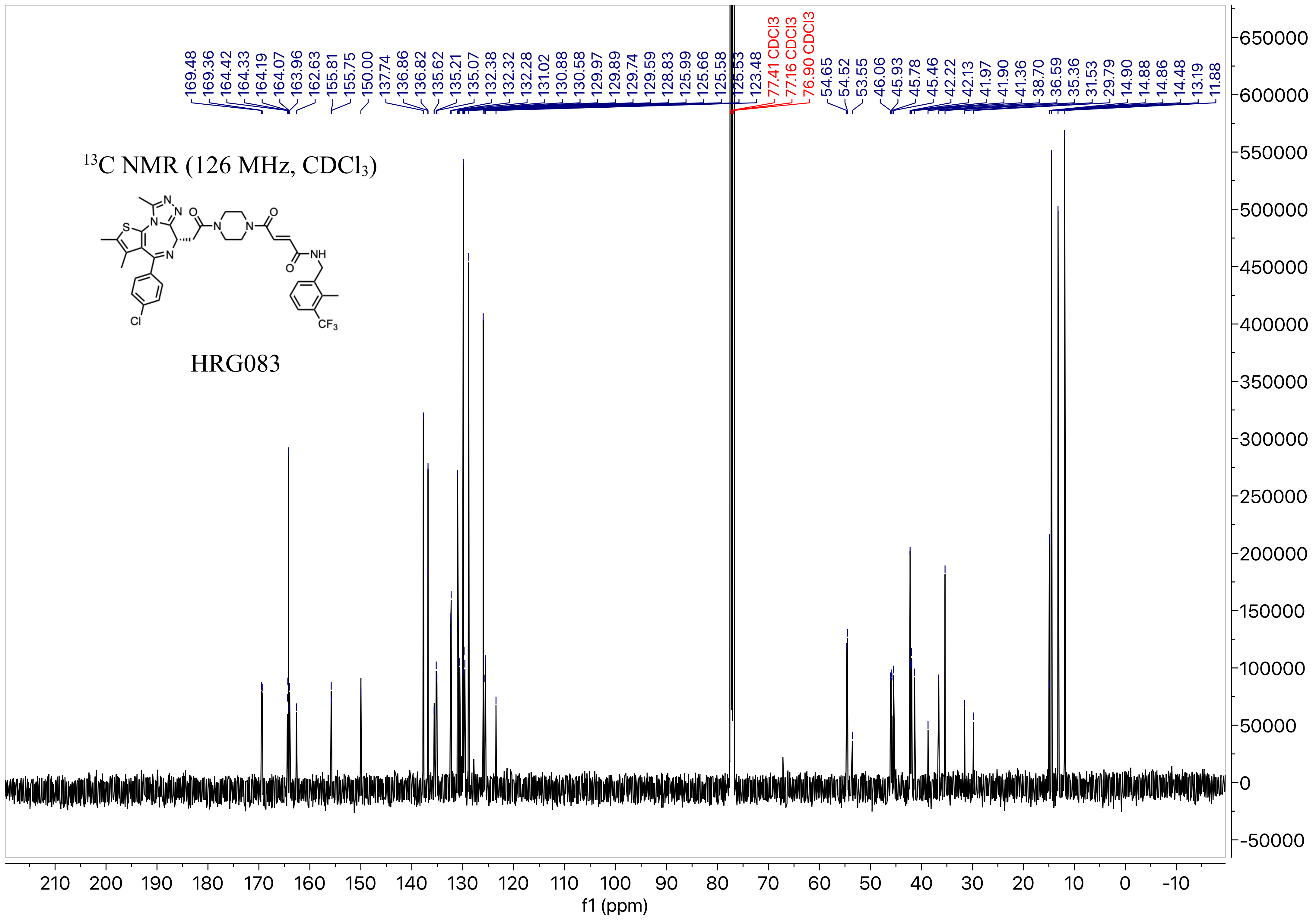

13C NMR (126 MHz, CDCl3)
HRG083
169.48
169.36
164.42
164.33
164.19
164.07
163.96
162.63
155.81
155.75
150.00
137.74
136.86
136.82
135.62
135.21
135.07
132.38
132.32
132.28
131.02
130.88
130.58
129.97
129.89
129.74
129.59
128.83
125.99
125.66
125.58
125.53
123.48
77.41 CDCl3
77.16 CDCl3
76.90 CDCl3
54.65
54.52
53.55
46.06
45.93
45.78
45.46
42.22
42.13
41.97
41.90
41.36
38.70
36.59
35.36
31.53
29.79
14.90
14.88
14.86
14.48
13.19
11.88
f1 (ppm)

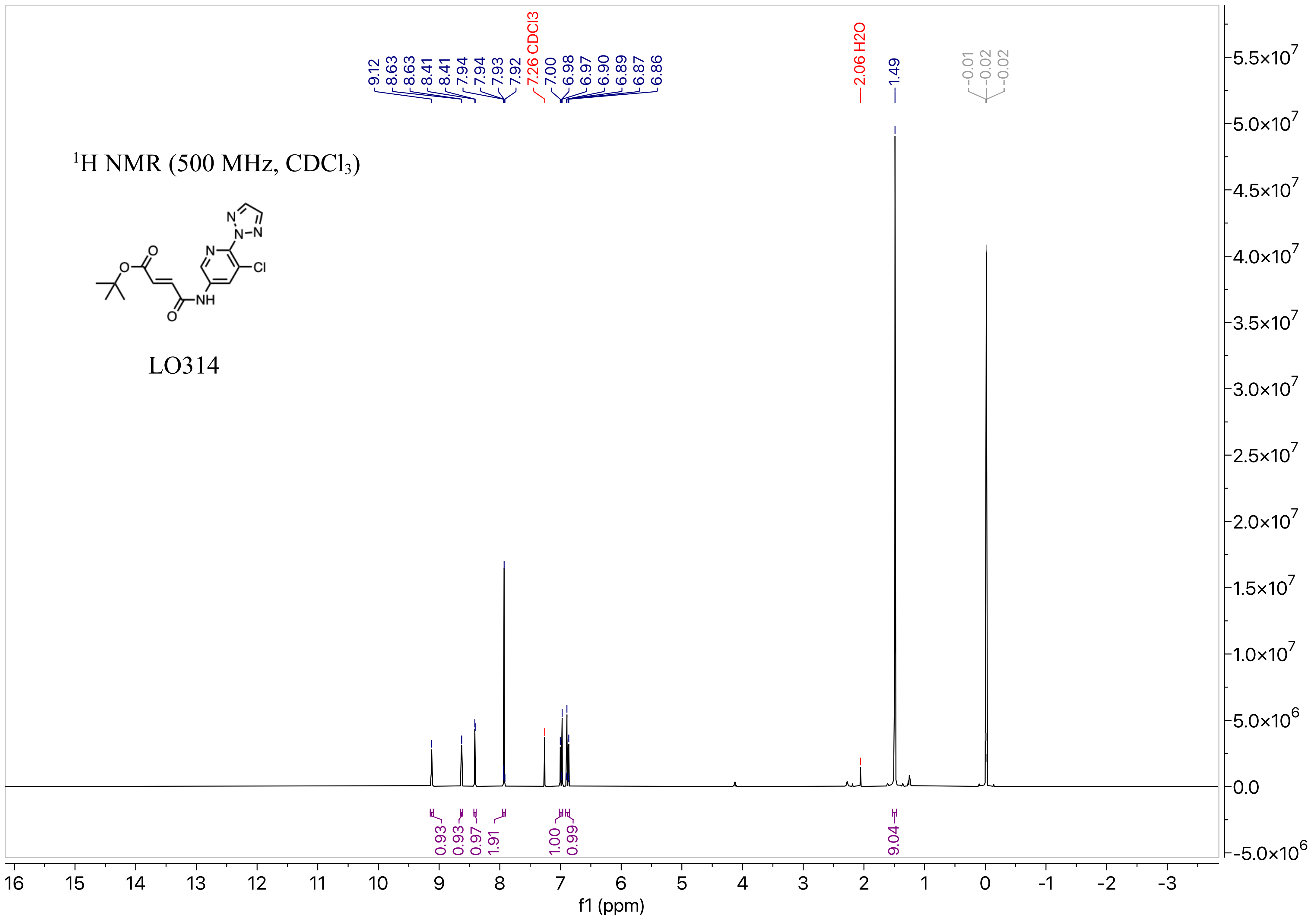

$^1$H NMR (500 MHz, CDCl$_3$)
LO314
9.12
8.63
8.63
8.41
8.41
7.94
7.94
7.93
7.92
7.26 CDCl3
7.00
6.98
6.97
6.90
6.89
6.87
6.86
2.06 H2O
1.49
-0.01
-0.02
-0.02
0.93
0.93
0.97
1.91
1.00
0.99
9.04
f1 (ppm)

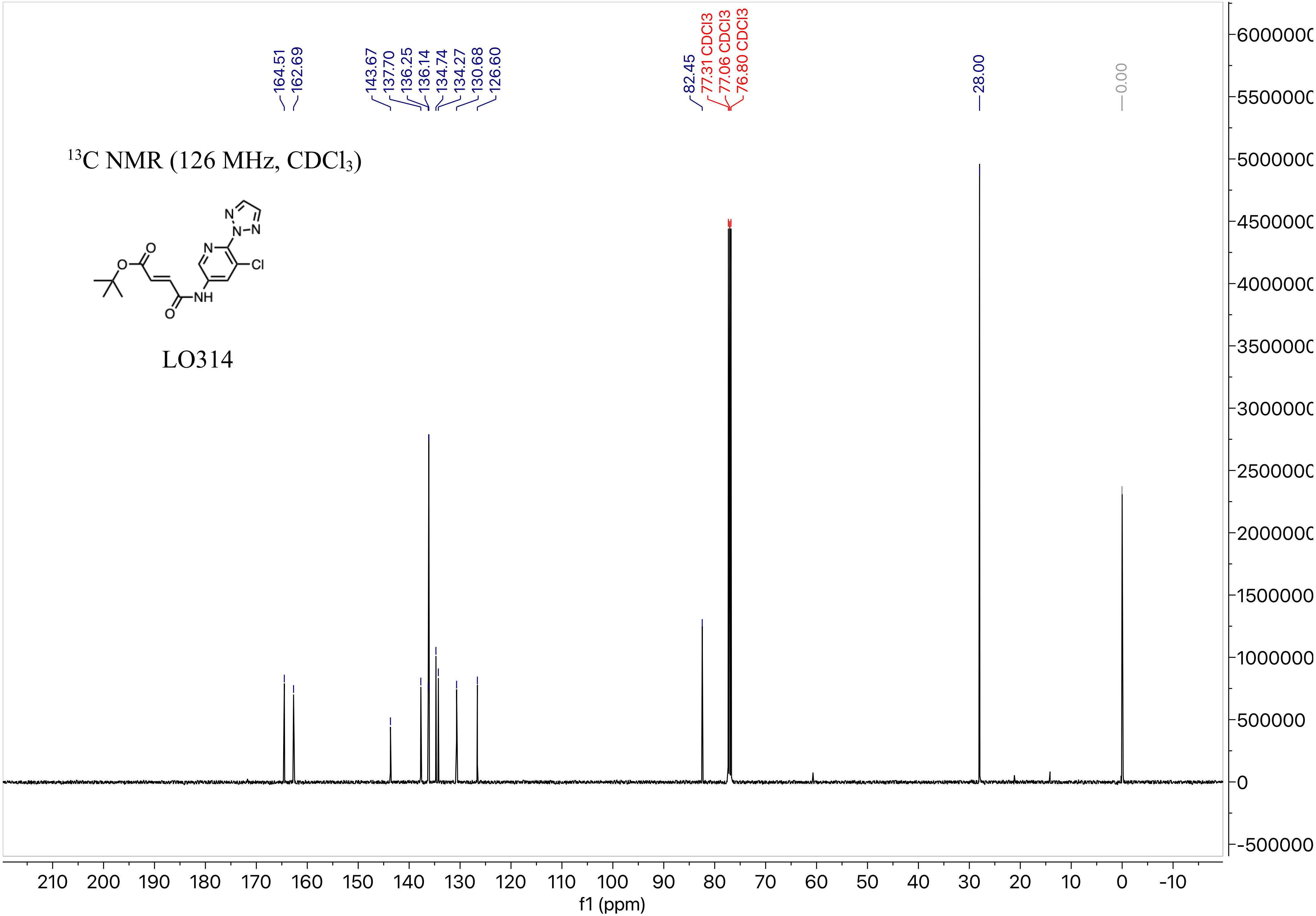

13C NMR (126 MHz, CDCl3)
LO314
164.51
162.69
143.67
137.70
136.25
136.14
134.74
134.27
130.68
126.60
82.45
77.31 CDCl3
77.06 CDCl3
76.80 CDCl3
28.00
0.00
f1 (ppm)

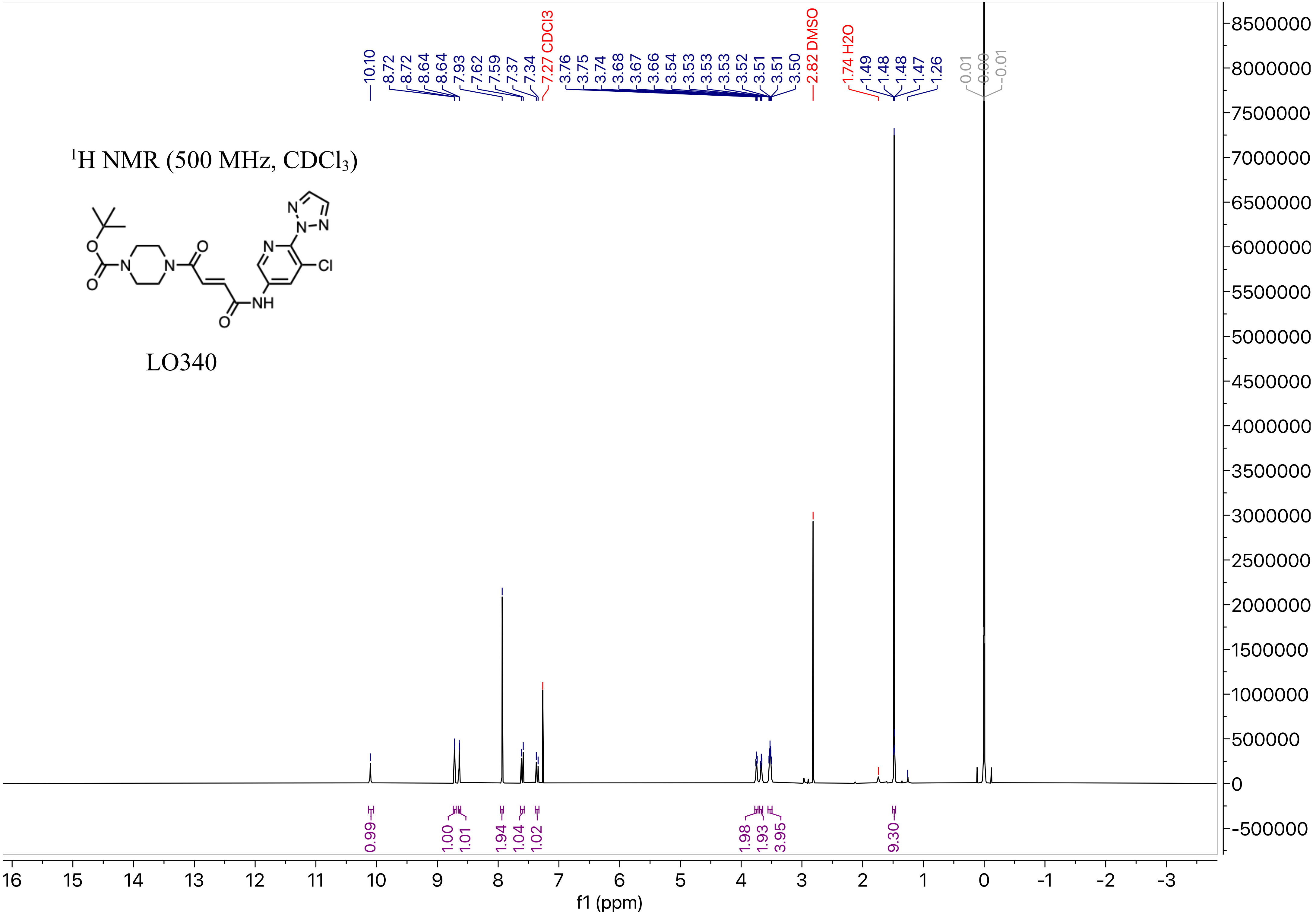

1H NMR (500 MHz, CDCl3)
LO340
10.10
8.72
8.72
8.64
8.64
7.93
7.62
7.59
7.37
7.34
7.27 CDCl3
3.76
3.75
3.74
3.68
3.67
3.66
3.54
3.53
3.53
3.53
3.52
3.51
3.51
3.50
2.82 DMSO
1.74 H2O
1.49
1.48
1.48
1.47
1.26
0.01
0.00
-0.01
0.99
1.00
1.01
1.94
1.04
1.02
1.98
1.93
3.95
9.30
f1 (ppm)

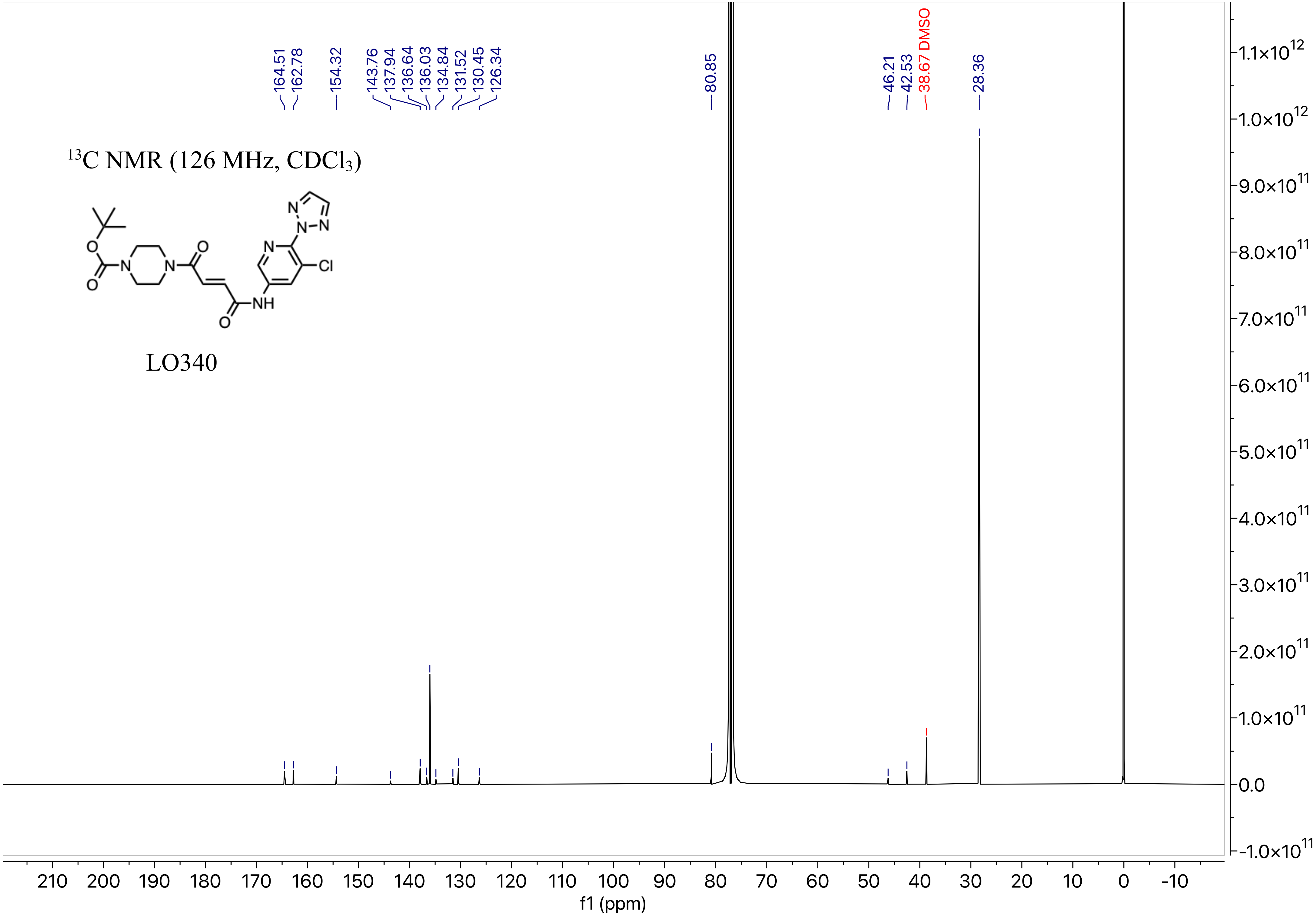

$^{13}$C NMR (126 MHz, CDCl$_3$)
LO340
164.51
162.78
154.32
143.76
137.94
136.64
136.03
134.84
131.52
130.45
126.34
80.85
46.21
42.53
38.67 DMSO
28.36
f1 (ppm)

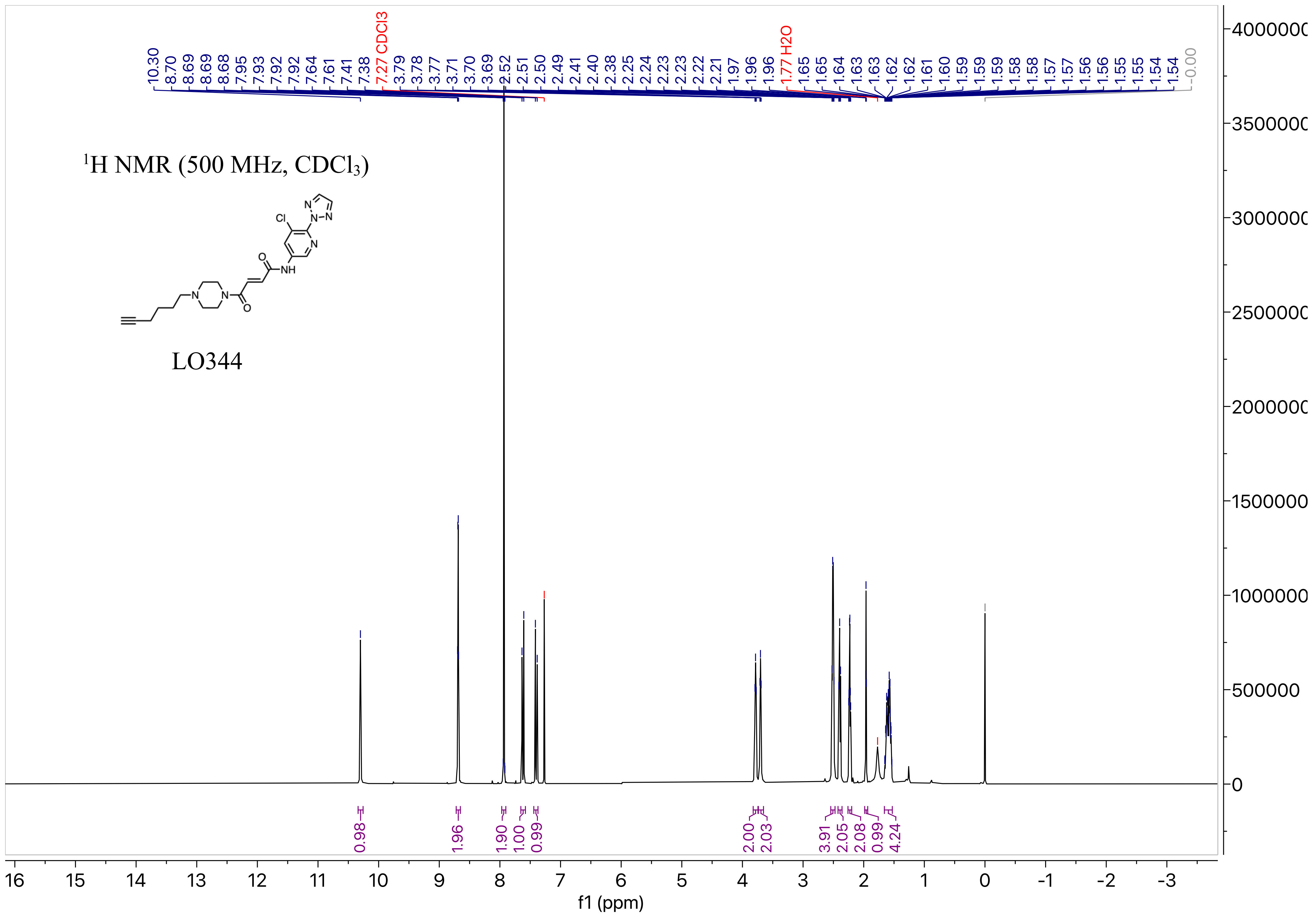

$^1$H NMR (500 MHz, CDCl$_3$)
LO344
10.30 8.70 8.69 8.69 8.68 7.95 7.93 7.92 7.92 7.64 7.61 7.41 7.38 7.27 CDCl3 3.79 3.78 3.77 3.71 3.70 3.69 2.52 2.51 2.50 2.49 2.41 2.40 2.38 2.25 2.24 2.23 2.23 2.22 2.21 1.97 1.96 1.96 1.77 H2O 1.65 1.65 1.64 1.63 1.63 1.62 1.62 1.61 1.60 1.59 1.59 1.59 1.58 1.58 1.57 1.57 1.56 1.56 1.55 1.55 1.54 1.54 -0.00
0.98 1.96 1.90 1.00 0.99 2.00 2.03 3.91 2.05 2.08 0.99 4.24
f1 (ppm)

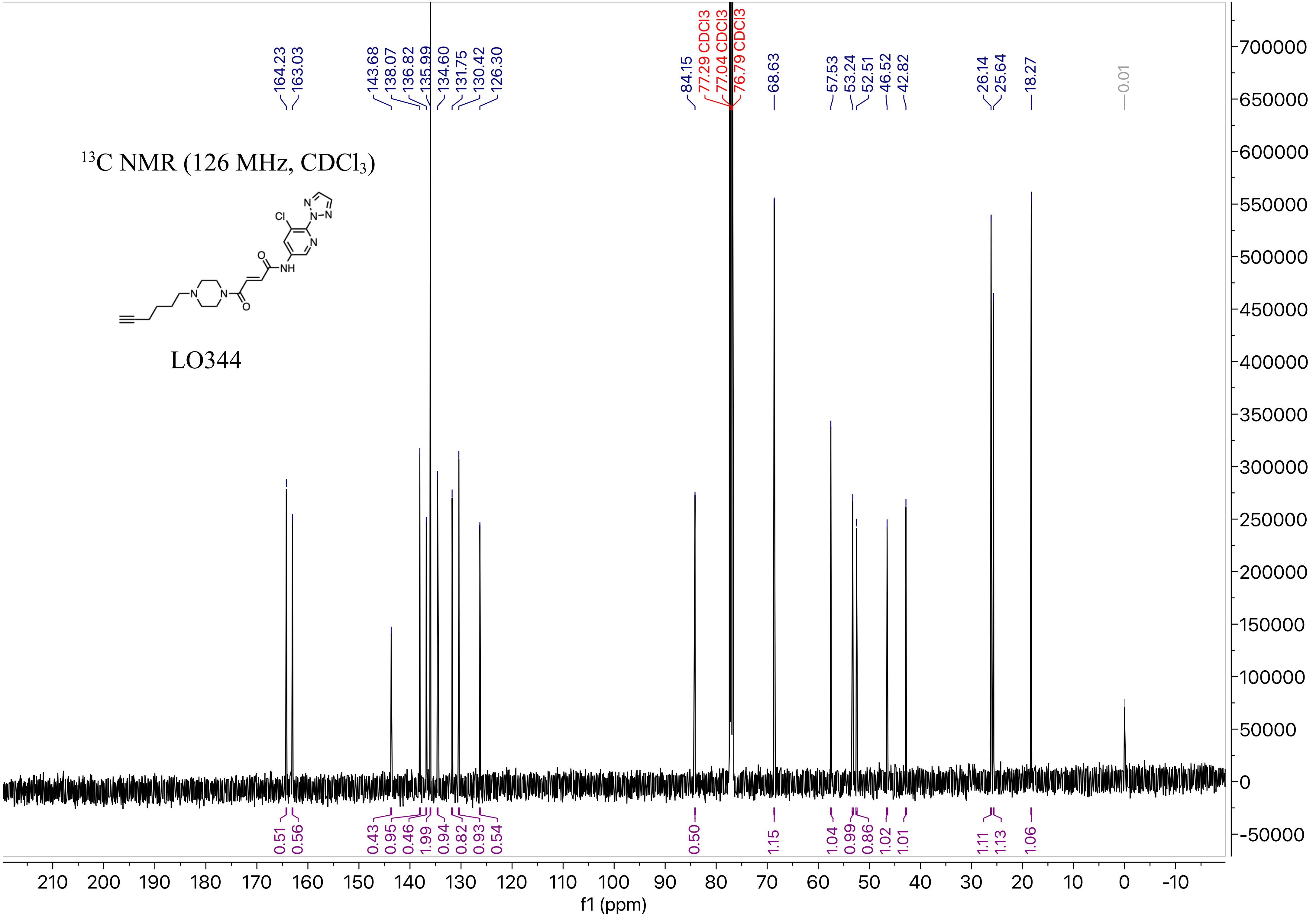
13C NMR (126 MHz, CDCl3)
LO344
164.23
163.03
143.68
138.07
136.82
135.99
134.60
131.75
130.42
126.30
84.15
77.29 CDCl3
77.04 CDCl3
76.79 CDCl3
68.63
57.53
53.24
52.51
46.52
42.82
26.14
25.64
18.27
0.01
f1 (ppm)

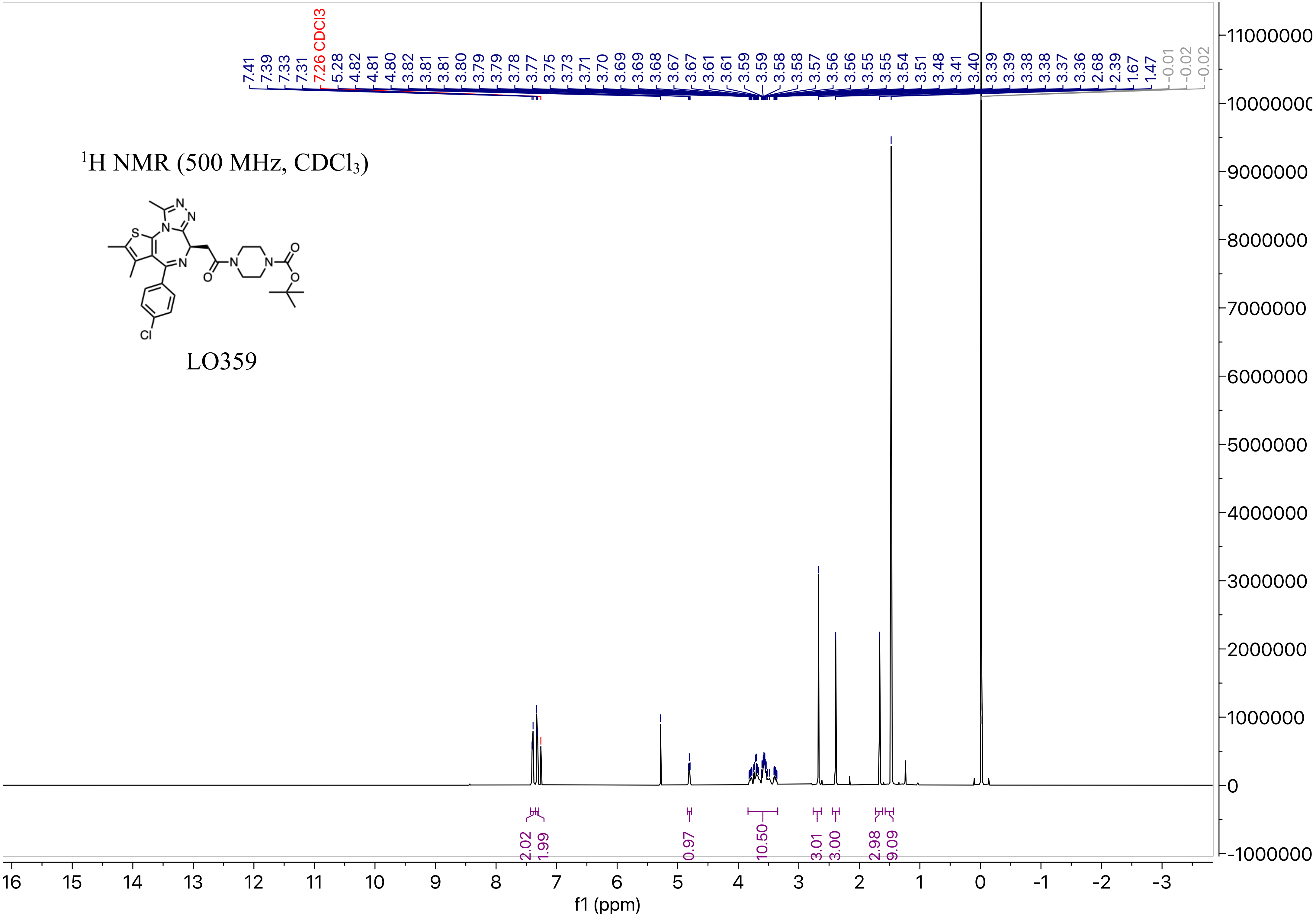
7.41 7.39 7.33 7.31 7.26 CDCl3 5.28 4.82 4.81 4.80 3.82 3.81 3.81 3.80 3.79 3.79 3.78 3.77 3.75 3.73 3.71 3.70 3.69 3.69 3.68 3.67 3.67 3.61 3.61 3.59 3.59 3.58 3.58 3.57 3.56 3.56 3.55 3.55 3.54 3.51 3.48 3.41 3.40 3.39 3.39 3.38 3.38 3.37 3.36 2.68 2.39 1.67 1.47 -0.01 -0.02 -0.02
1H NMR (500 MHz, CDCl3)
LO359
2.02 1.99 0.97 10.50 3.01 3.00 2.98 9.09
f1 (ppm)

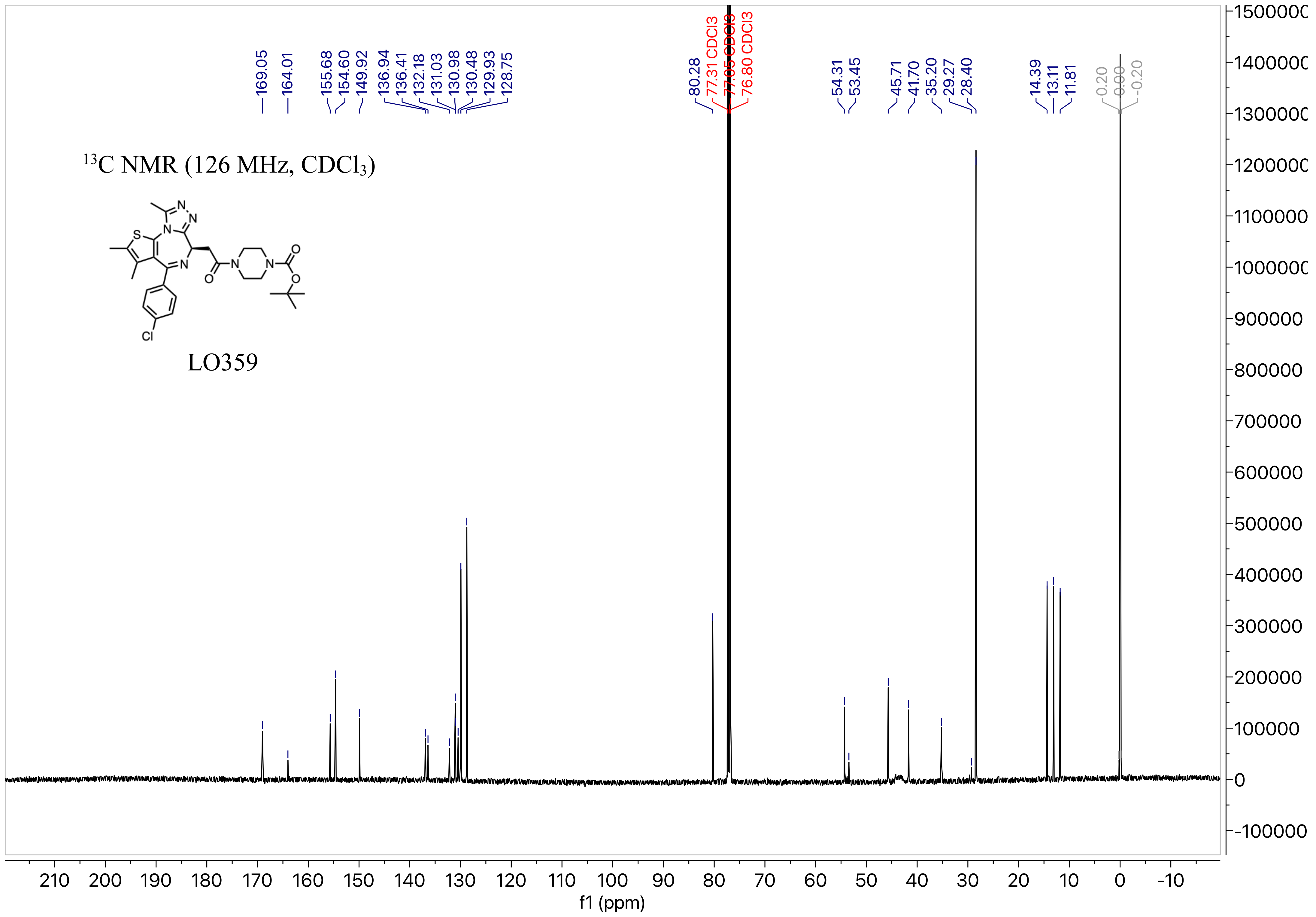

$^{13}$C NMR (126 MHz, CDCl$_3$)
LO359
169.05
164.01
155.68
154.60
149.92
136.94
136.41
132.18
131.03
130.98
130.48
129.93
128.75
80.28
77.31 CDCl3
77.05 CDCl3
76.80 CDCl3
54.31
53.45
45.71
41.70
35.20
29.27
28.40
14.39
13.11
11.81
0.20
0.00
-0.20
f1 (ppm)

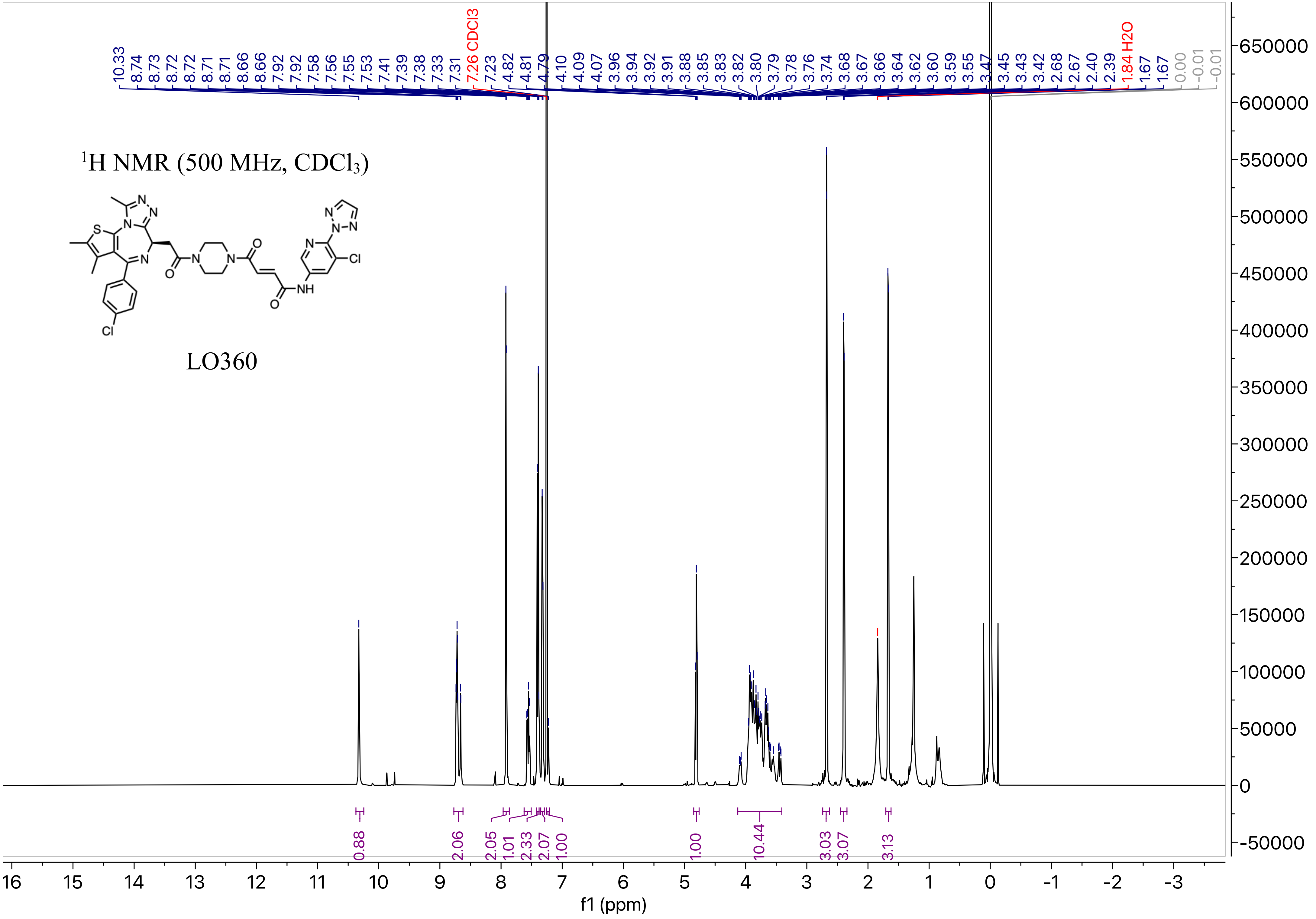

1H NMR (500 MHz, CDCl3)
LO360
10.33 8.74 8.73 8.72 8.72 8.71 8.71 8.66 8.66 7.92 7.92 7.58 7.56 7.55 7.53 7.41 7.39 7.38 7.33 7.31 7.26 CDCl3 7.23 4.82 4.81 4.79 4.10 4.09 4.07 3.96 3.94 3.92 3.91 3.88 3.85 3.83 3.82 3.80 3.79 3.78 3.76 3.74 3.68 3.67 3.66 3.64 3.62 3.60 3.59 3.55 3.47 3.45 3.43 3.42 2.68 2.67 2.40 2.39 1.84 H2O 1.67 1.67 0.00 -0.01 -0.01
0.88 2.06 2.05 1.01 2.33 2.07 1.00 1.00 10.44 3.03 3.07 3.13
f1 (ppm)

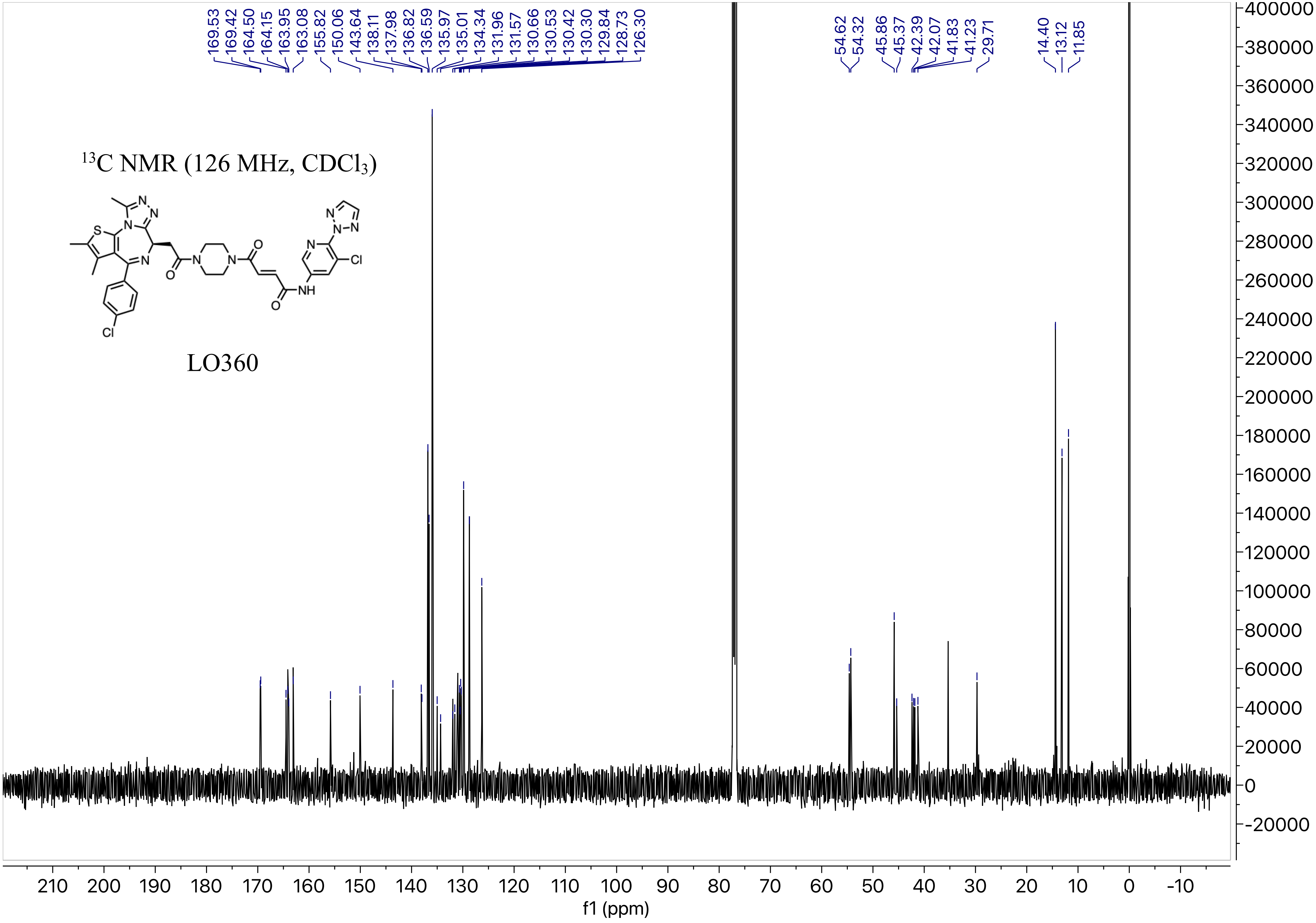
13C NMR (126 MHz, CDCl3)
LO360
169.53
169.42
164.50
164.15
163.95
163.08
155.82
150.06
143.64
138.11
137.98
136.82
136.59
135.97
135.01
134.34
131.96
131.57
130.66
130.53
130.42
130.30
129.84
128.73
126.30
54.62
54.32
45.86
45.37
42.39
42.07
41.83
41.23
29.71
14.40
13.12
11.85
f1 (ppm)

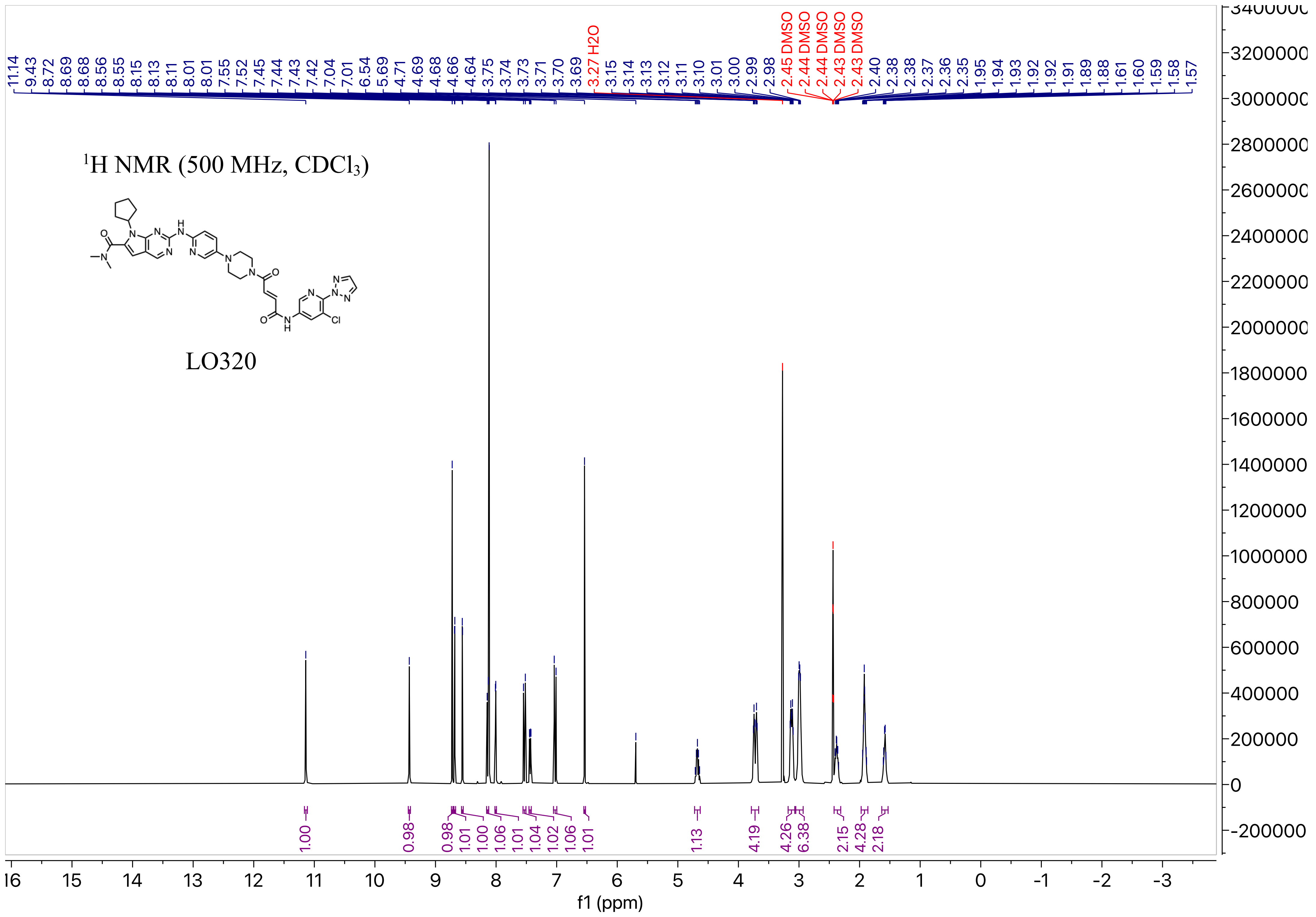

¹H NMR (500 MHz, CDCl₃)
LO320
f1 (ppm)

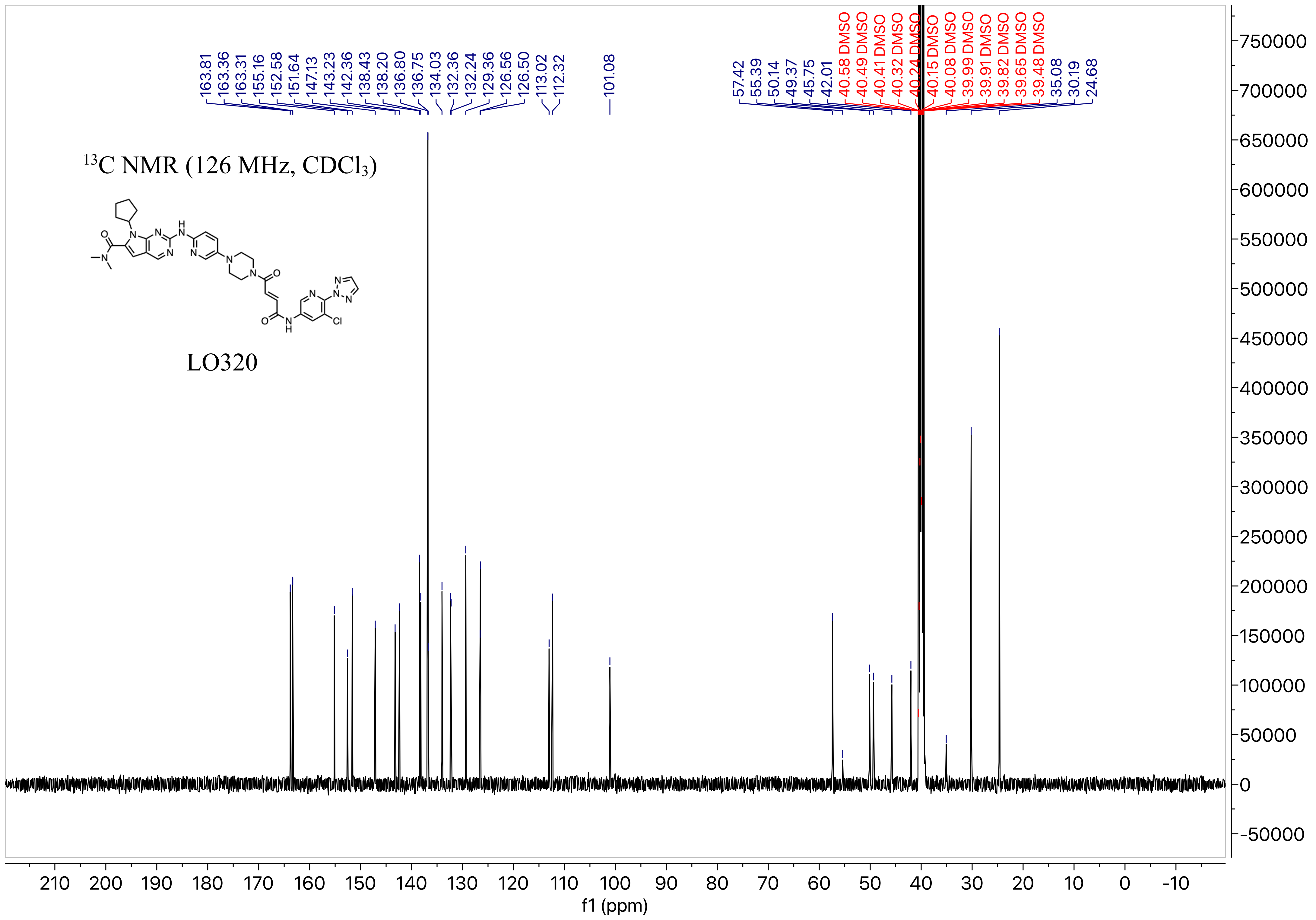

13C NMR (126 MHz, CDCl3)
LO320
163.81
163.36
163.31
155.16
152.58
151.64
147.13
143.23
142.36
138.43
138.20
136.80
136.75
134.03
132.36
132.24
129.36
126.56
126.50
113.02
112.32
101.08
57.42
55.39
50.14
49.37
45.75
42.01
40.58 DMSO
40.49 DMSO
40.41 DMSO
40.32 DMSO
40.24 DMSO
40.15 DMSO
40.08 DMSO
39.99 DMSO
39.91 DMSO
39.82 DMSO
39.65 DMSO
39.48 DMSO
35.08
30.19
24.68
f1 (ppm)

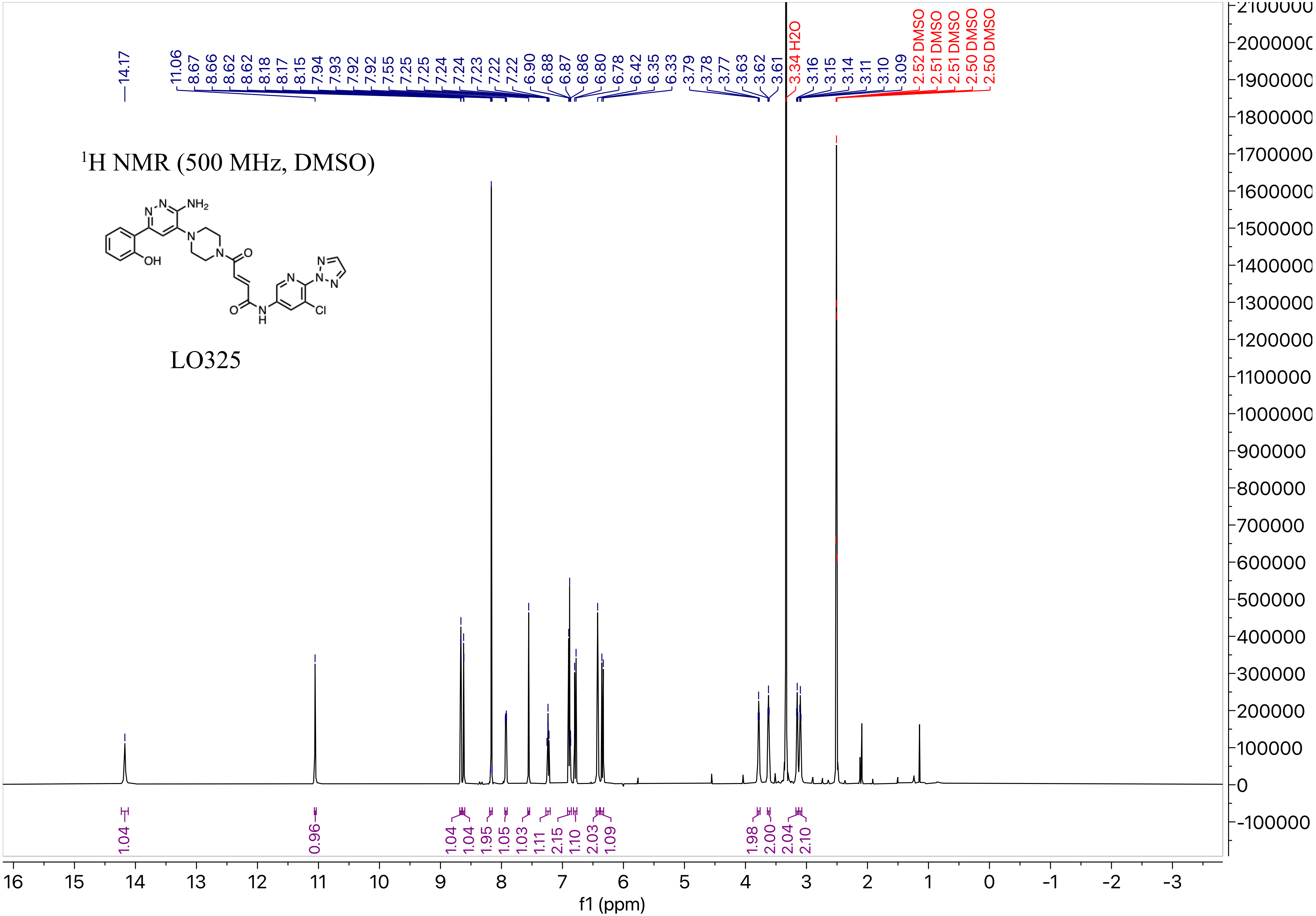

$^1$H NMR (500 MHz, DMSO)
LO325
14.17
11.06 8.67 8.66 8.62 8.62 8.18 8.17 8.15 7.94 7.93 7.92 7.92 7.55 7.25 7.25 7.24 7.24 7.23 7.22 7.22 6.90 6.88 6.87 6.86 6.80 6.78 6.42 6.35 6.33 3.79 3.78 3.77 3.63 3.62 3.61
3.34 H2O
3.16 3.15 3.14 3.11 3.10 3.09
2.52 DMSO 2.51 DMSO 2.51 DMSO 2.50 DMSO 2.50 DMSO
1.04 0.96 1.04 1.04 1.95 1.05 1.03 1.11 2.15 1.10 2.03 1.09 1.98 2.00 2.04 2.10
f1 (ppm)

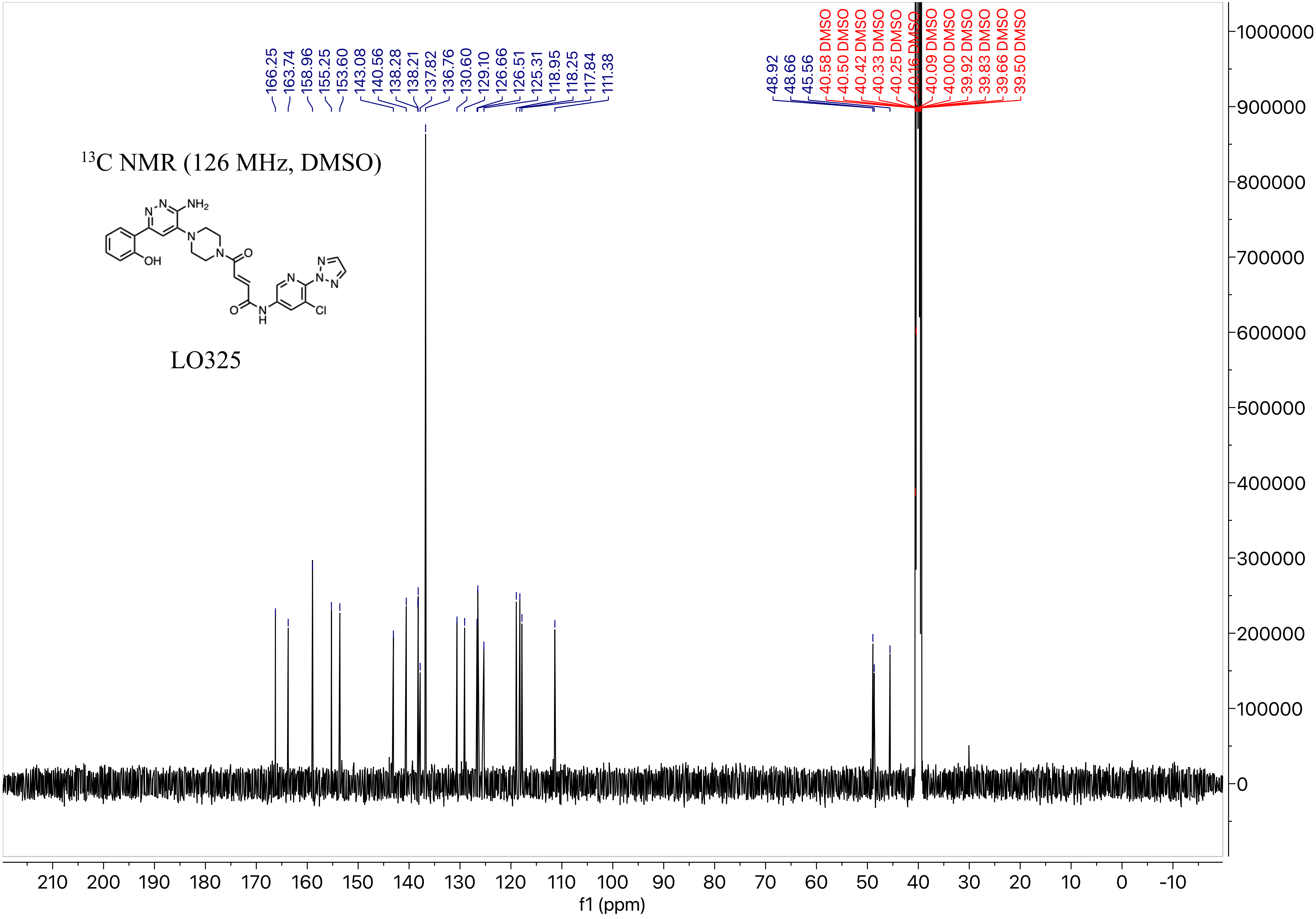

$^{13}$C NMR (126 MHz, DMSO)
LO325
166.25
163.74
158.96
155.25
153.60
143.08
140.56
138.28
138.21
137.82
136.76
130.60
129.10
126.66
126.51
125.31
118.95
118.25
117.84
111.38
48.92
48.66
45.56
40.58 DMSO
40.50 DMSO
40.42 DMSO
40.33 DMSO
40.25 DMSO
40.16 DMSO
40.09 DMSO
40.00 DMSO
39.92 DMSO
39.83 DMSO
39.66 DMSO
39.50 DMSO
f1 (ppm)

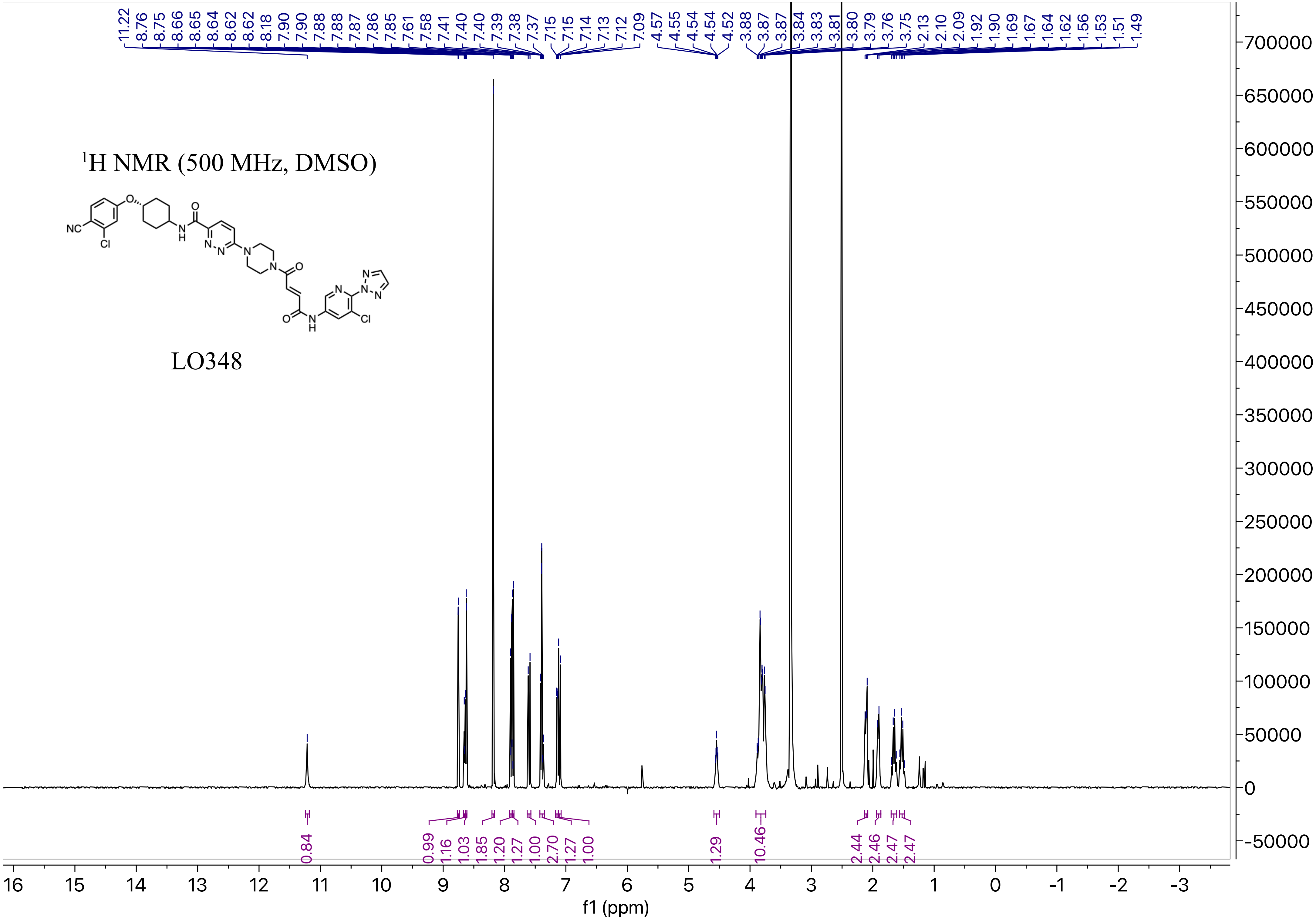

1H NMR (500 MHz, DMSO)
LO348
11.22 8.76 8.75 8.66 8.65 8.64 8.62 8.62 8.18 7.90 7.90 7.88 7.88 7.87 7.86 7.85 7.61 7.58 7.41 7.40 7.40 7.39 7.38 7.37 7.15 7.15 7.14 7.13 7.12 7.09 4.57 4.55 4.54 4.54 4.52 3.88 3.87 3.87 3.84 3.83 3.81 3.80 3.79 3.76 3.75 2.13 2.10 2.09 1.92 1.90 1.69 1.67 1.64 1.62 1.56 1.53 1.51 1.49
0.84 0.99 1.16 1.03 1.85 1.20 1.27 1.00 2.70 1.27 1.00 1.29 10.46 2.44 2.46 2.47 2.47
f1 (ppm)

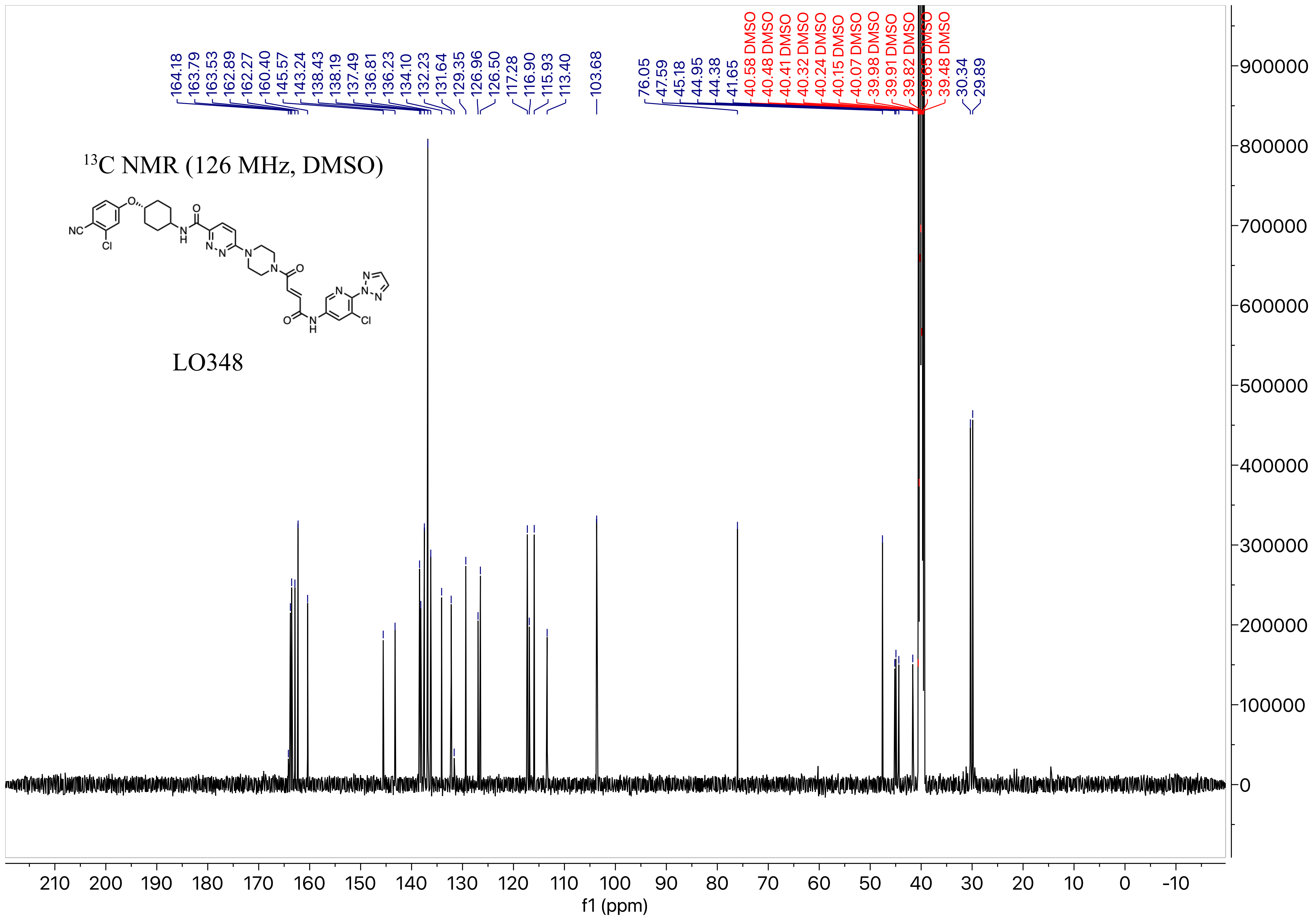
164.18
163.79
163.53
162.89
162.27
160.40
145.57
143.24
138.43
138.19
137.49
136.81
136.23
134.10
132.23
131.64
129.35
126.96
126.50
117.28
116.90
115.93
113.40
103.68
76.05
47.59
45.18
44.95
44.38
41.65
40.58 DMSO
40.48 DMSO
40.41 DMSO
40.32 DMSO
40.24 DMSO
40.15 DMSO
40.07 DMSO
39.98 DMSO
39.91 DMSO
39.82 DMSO
39.65 DMSO
39.48 DMSO
30.34
29.89
$^{13}$C NMR (126 MHz, DMSO)
LO348
f1 (ppm)

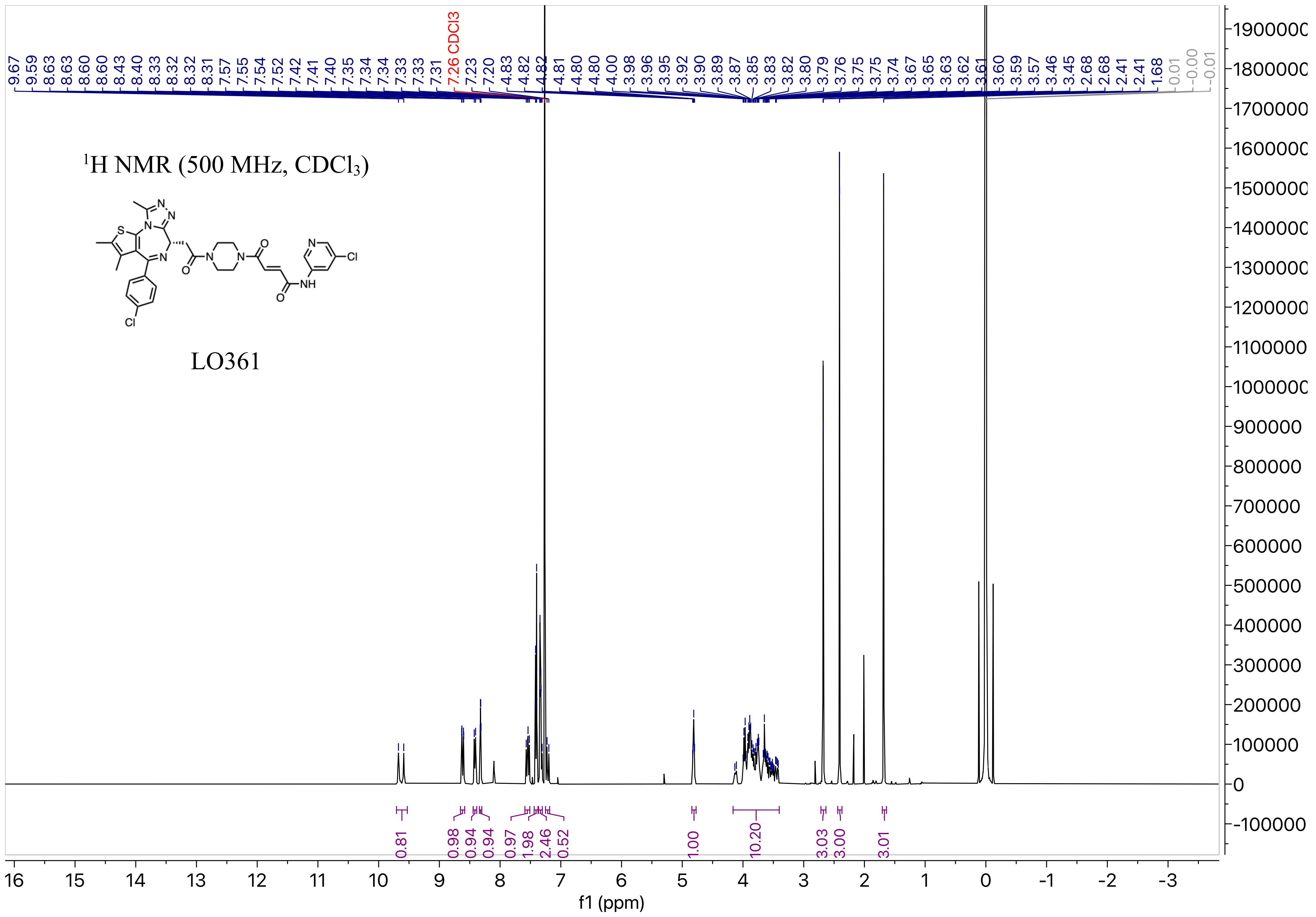

1H NMR (500 MHz, CDCl3)
LO361
f1 (ppm)

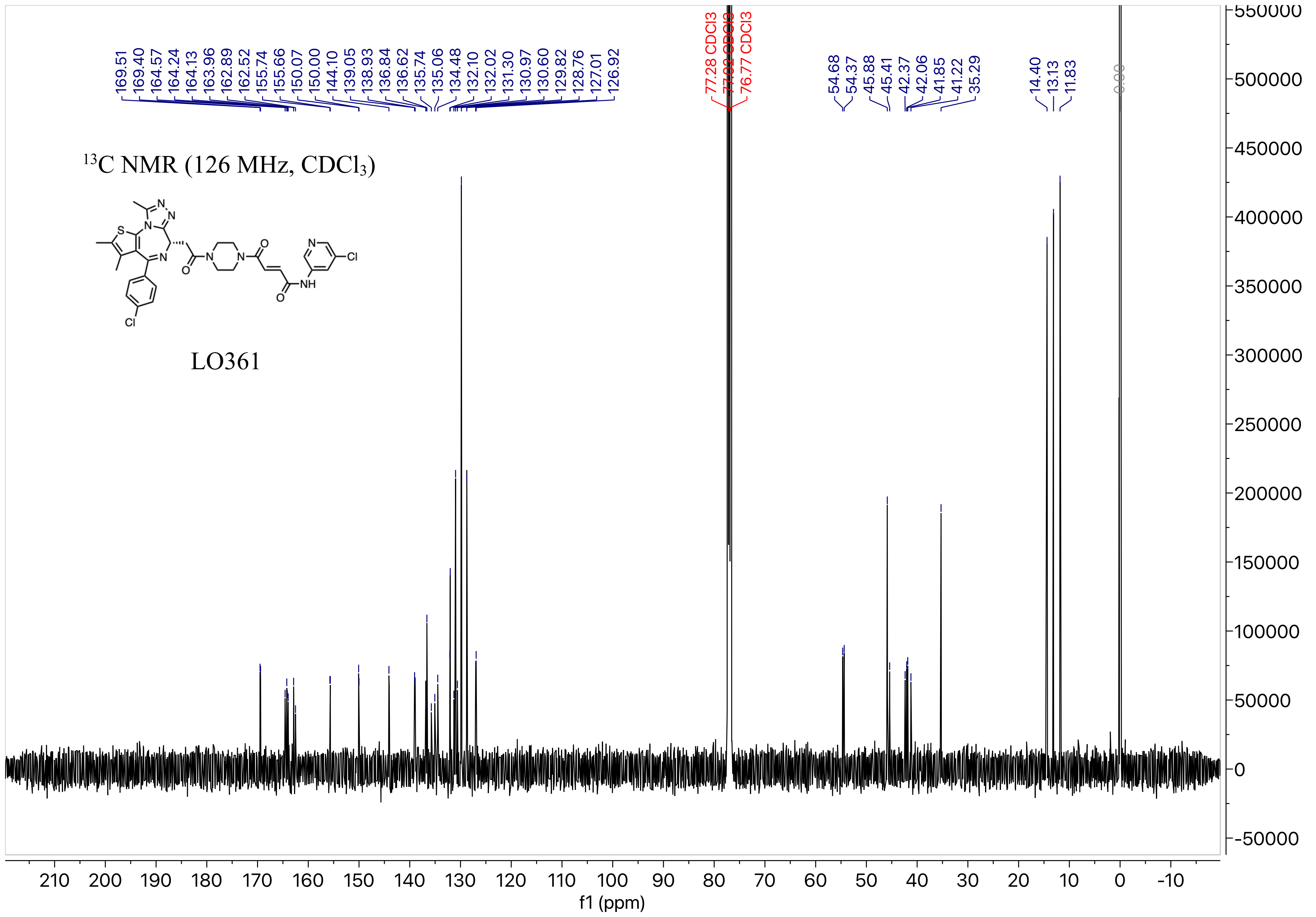

169.51
169.40
164.57
164.24
164.13
163.96
162.89
162.52
155.74
155.66
150.07
150.00
144.10
139.05
138.93
136.84
136.62
135.74
135.06
134.48
132.10
132.02
131.30
130.97
130.60
129.82
128.76
127.01
126.92
77.28 CDCl3
77.02 CDCl3
76.77 CDCl3
54.68
54.37
45.88
45.41
42.37
42.06
41.85
41.22
35.29
14.40
13.13
11.83
0.00
$^{13}$C NMR (126 MHz, CDCl$_3$)
LO361
f1 (ppm)

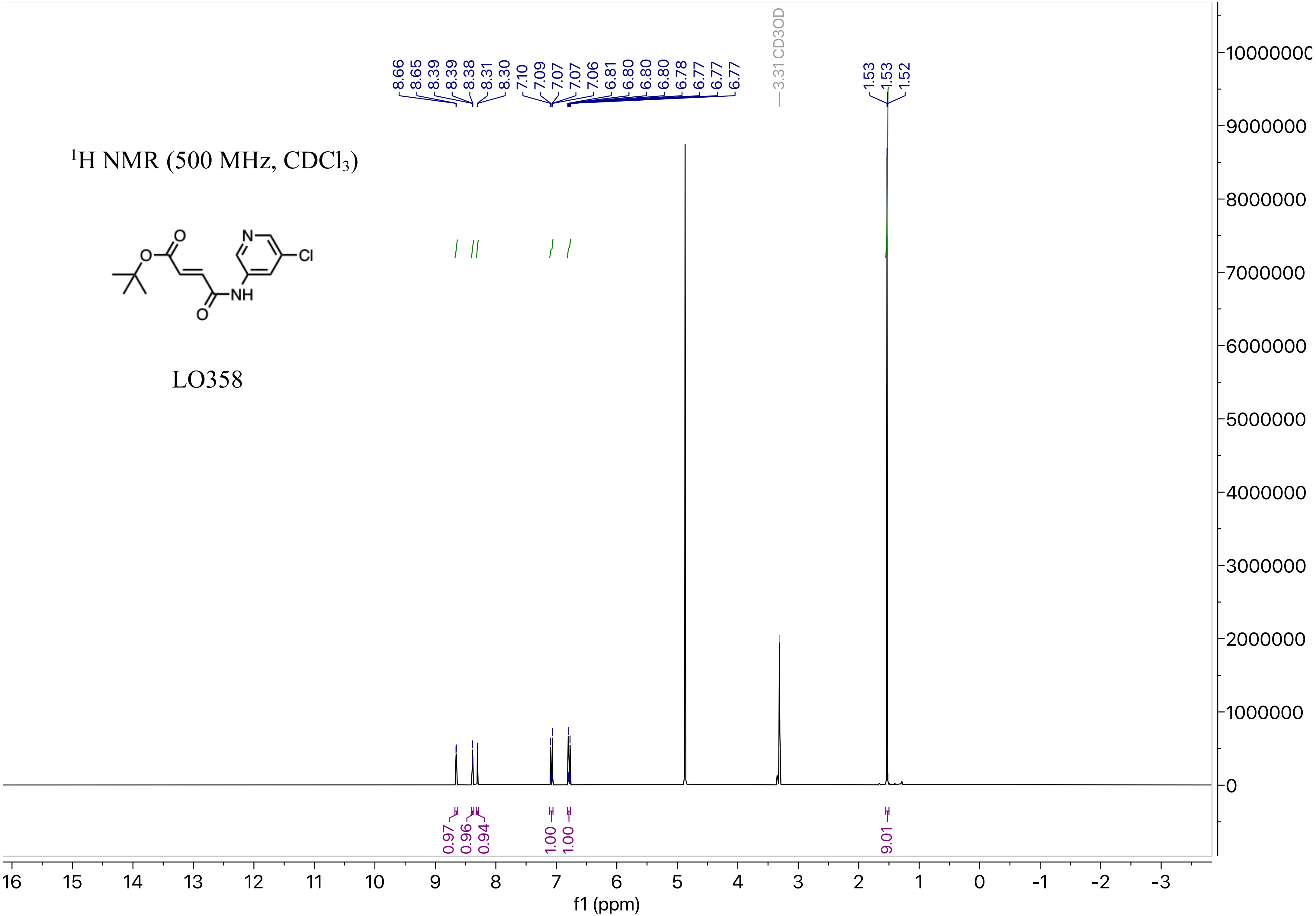

$^{1}H$ NMR (500 MHz, $CDCl_3$)
LO358
O
O
N
Cl
NH
O
8.66
8.65
8.39
8.39
8.38
8.31
8.30
7.10
7.09
7.07
7.07
7.06
6.81
6.80
6.80
6.80
6.78
6.77
6.77
6.77
3.31 CD3OD
1.53
1.53
1.52
0.97
0.96
0.94
1.00
1.00
9.01
f1 (ppm)

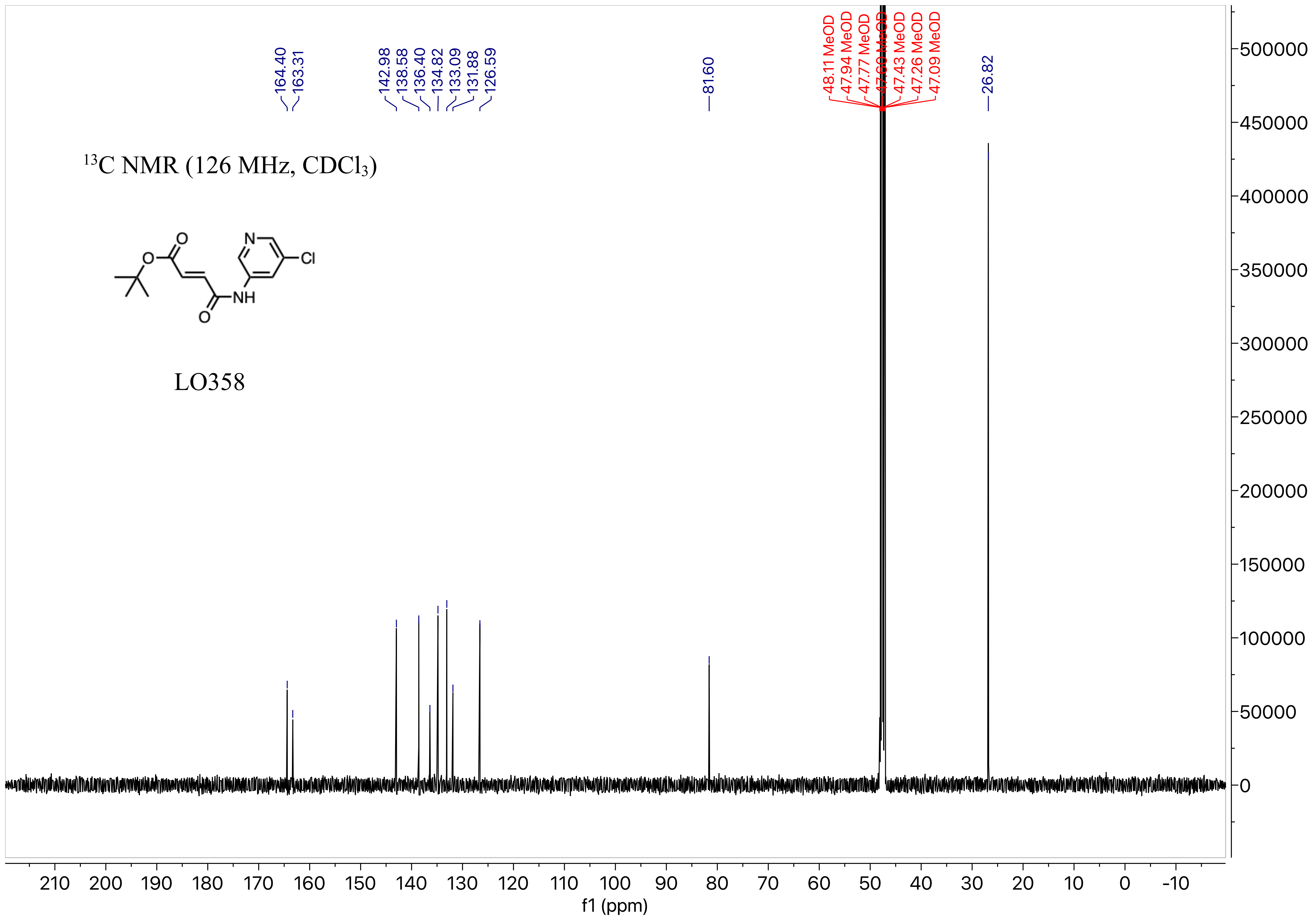

164.40
163.31
142.98
138.58
136.40
134.82
133.09
131.88
126.59
81.60
48.11 MeOD
47.94 MeOD
47.77 MeOD
47.43 MeOD
47.26 MeOD
47.09 MeOD
26.82
$^{13}$C NMR (126 MHz, CDCl$_3$)
O
O
N
Cl
NH
O
LO358
f1 (ppm)

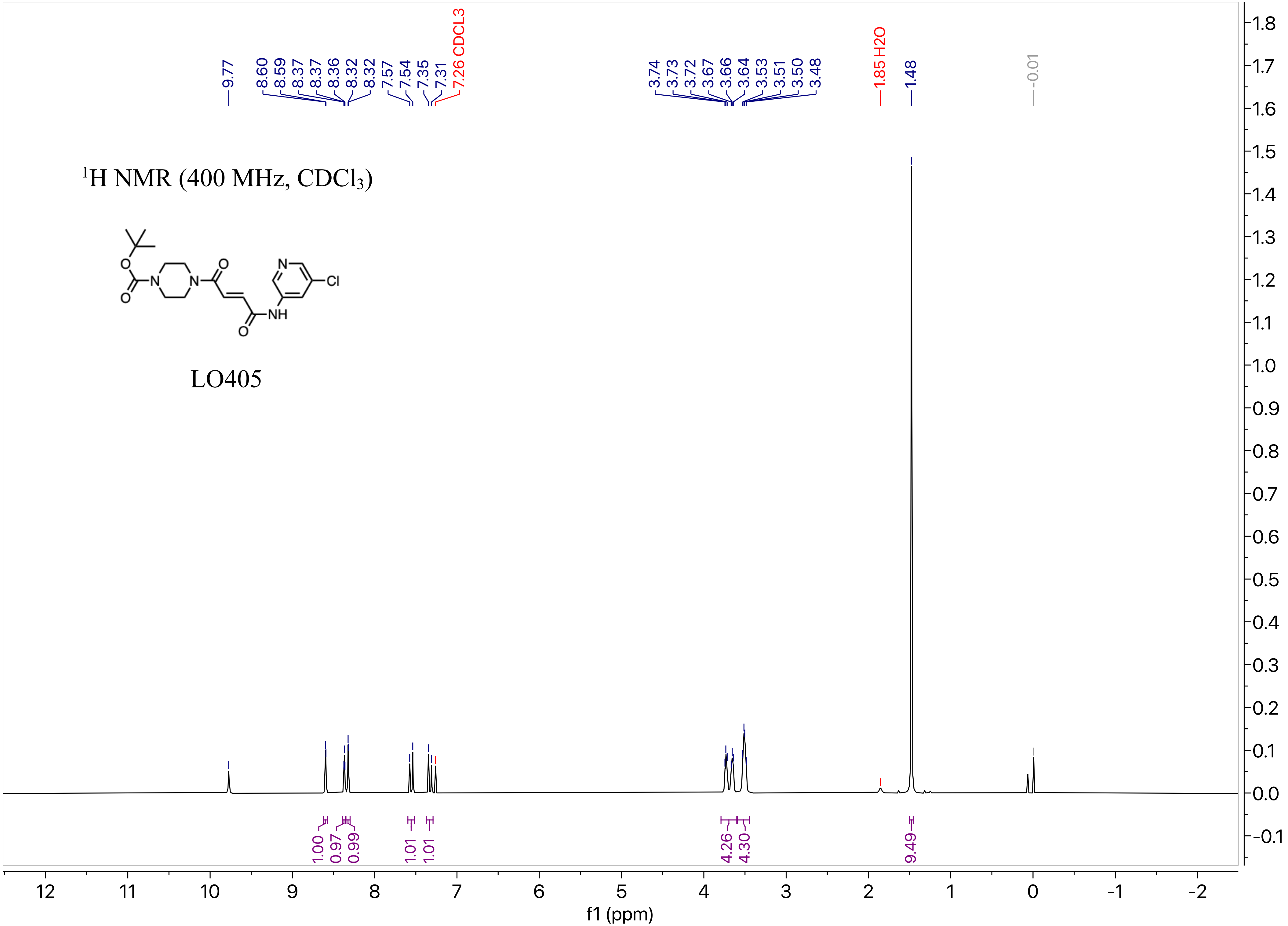

9.77
8.60
8.59
8.37
8.37
8.36
8.32
8.32
7.57
7.54
7.35
7.31
7.26 CDCL3
3.74
3.73
3.72
3.67
3.66
3.64
3.53
3.51
3.50
3.48
1.85 H2O
1.48
-0.01
1H NMR (400 MHz, CDCl3)
O
N
N
O
O
N
Cl
NH
O
LO405
1.00
0.97
0.99
1.01
1.01
4.26
4.30
9.49
f1 (ppm)

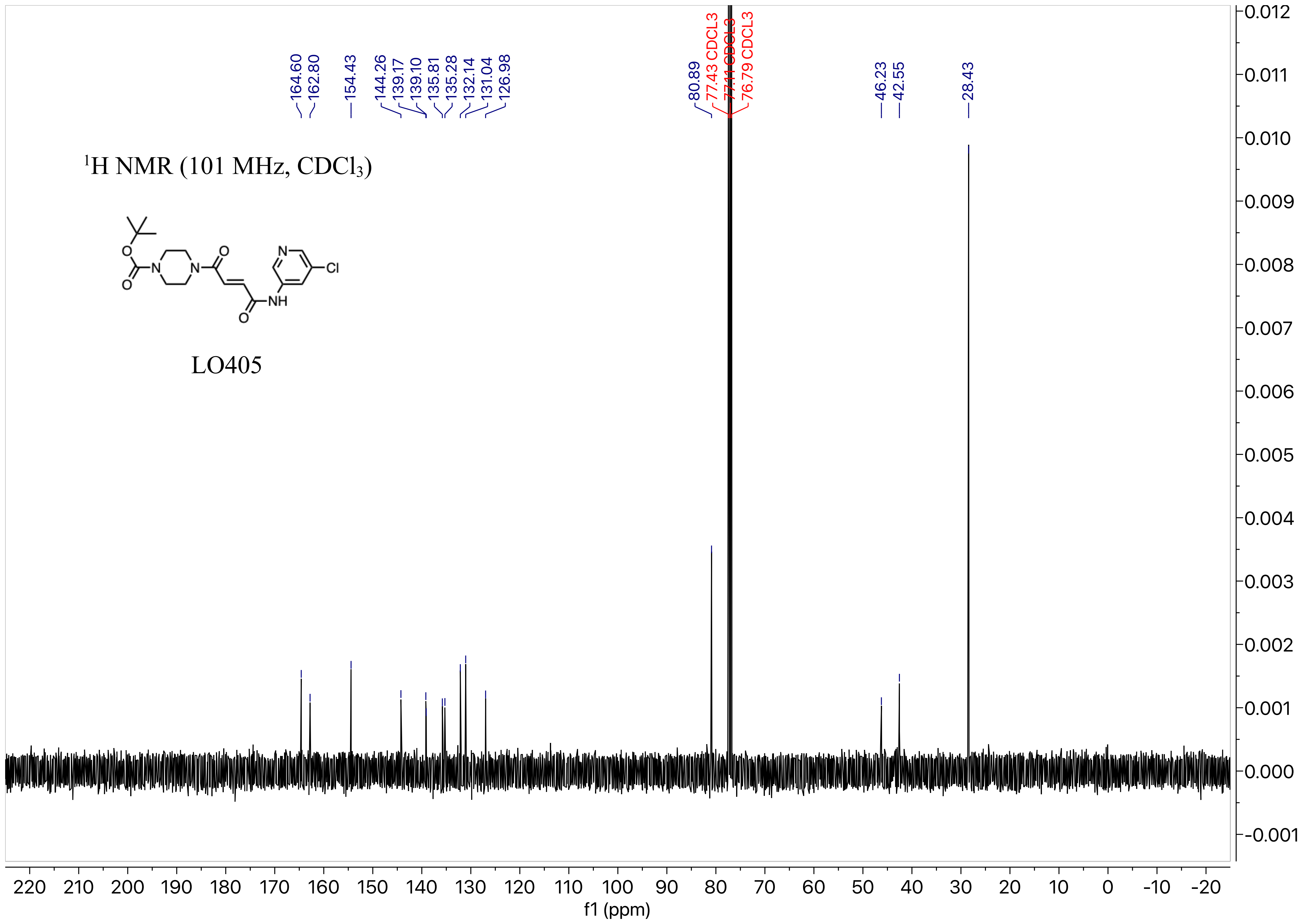

164.60
162.80
154.43
144.26
139.17
139.10
135.81
135.28
132.14
131.04
126.98
80.89
77.43 CDCL3
77.11 CDCL3
76.79 CDCL3
46.23
42.55
28.43
$^{1}$H NMR (101 MHz, $CDCl_3$)
O
N
N
O
N
Cl
NH
O
LO405
220 210 200 190 180 170 160 150 140 130 120 110 100 90 80 70 60 50 40 30 20 10 0 -10 -20
f1 (ppm)
0.012 0.011 0.010 0.009 0.008 0.007 0.006 0.005 0.004 0.003 0.002 0.001 0.000 -0.001

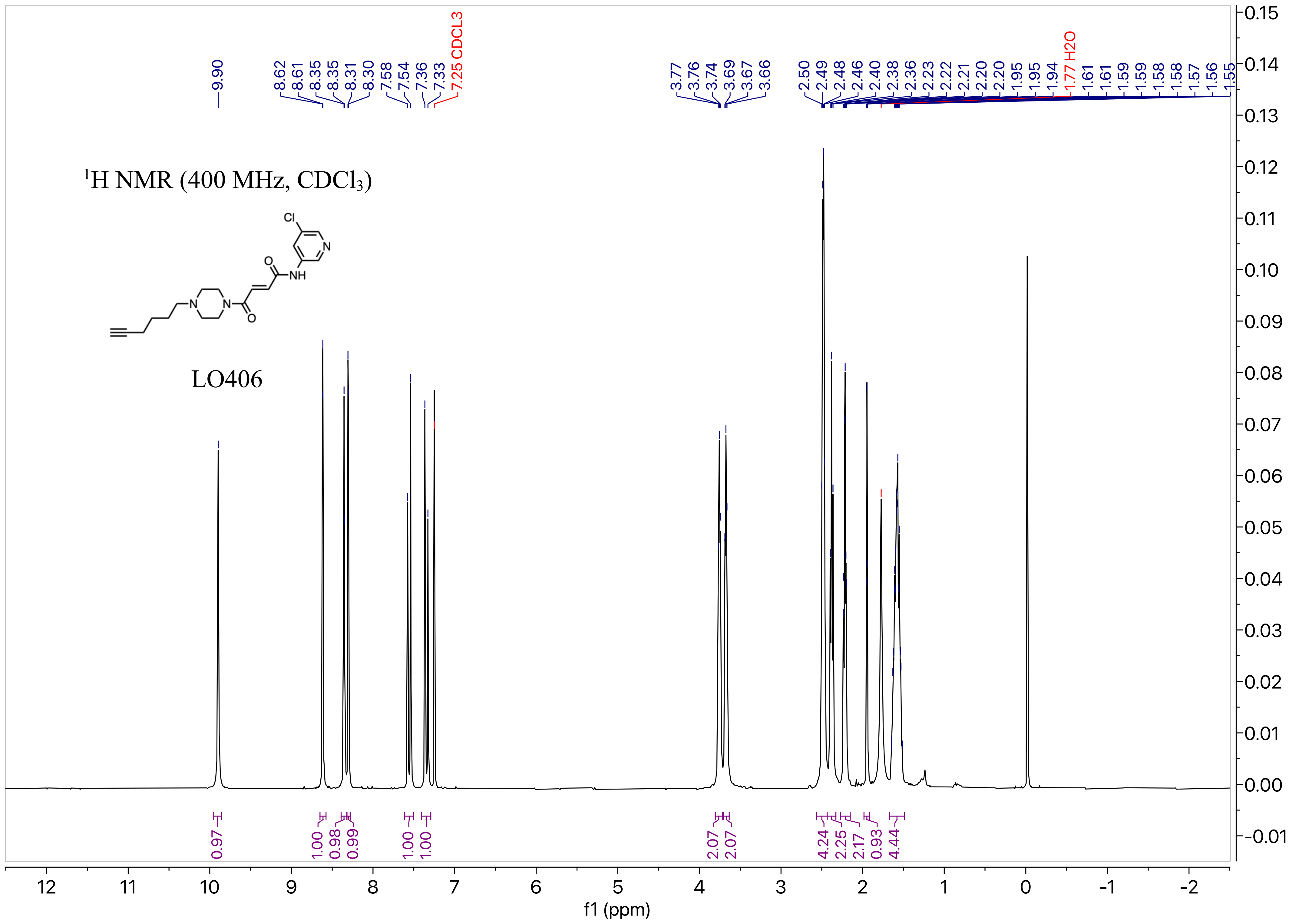

1H NMR (400 MHz, CDCl3)
LO406
Cl
N
O
NH
N
N
O
9.90
8.62
8.61
8.35
8.35
8.31
8.30
7.58
7.54
7.36
7.33
7.25 CDCL3
3.77
3.76
3.74
3.69
3.67
3.66
2.50
2.49
2.48
2.46
2.40
2.38
2.36
2.23
2.22
2.21
2.20
2.20
1.95
1.95
1.94
1.77 H2O
1.61
1.61
1.59
1.59
1.58
1.58
1.57
1.56
1.55
0.97
1.00
0.98
0.99
1.00
1.00
2.07
2.07
4.24
2.25
2.17
0.93
4.44
f1 (ppm)

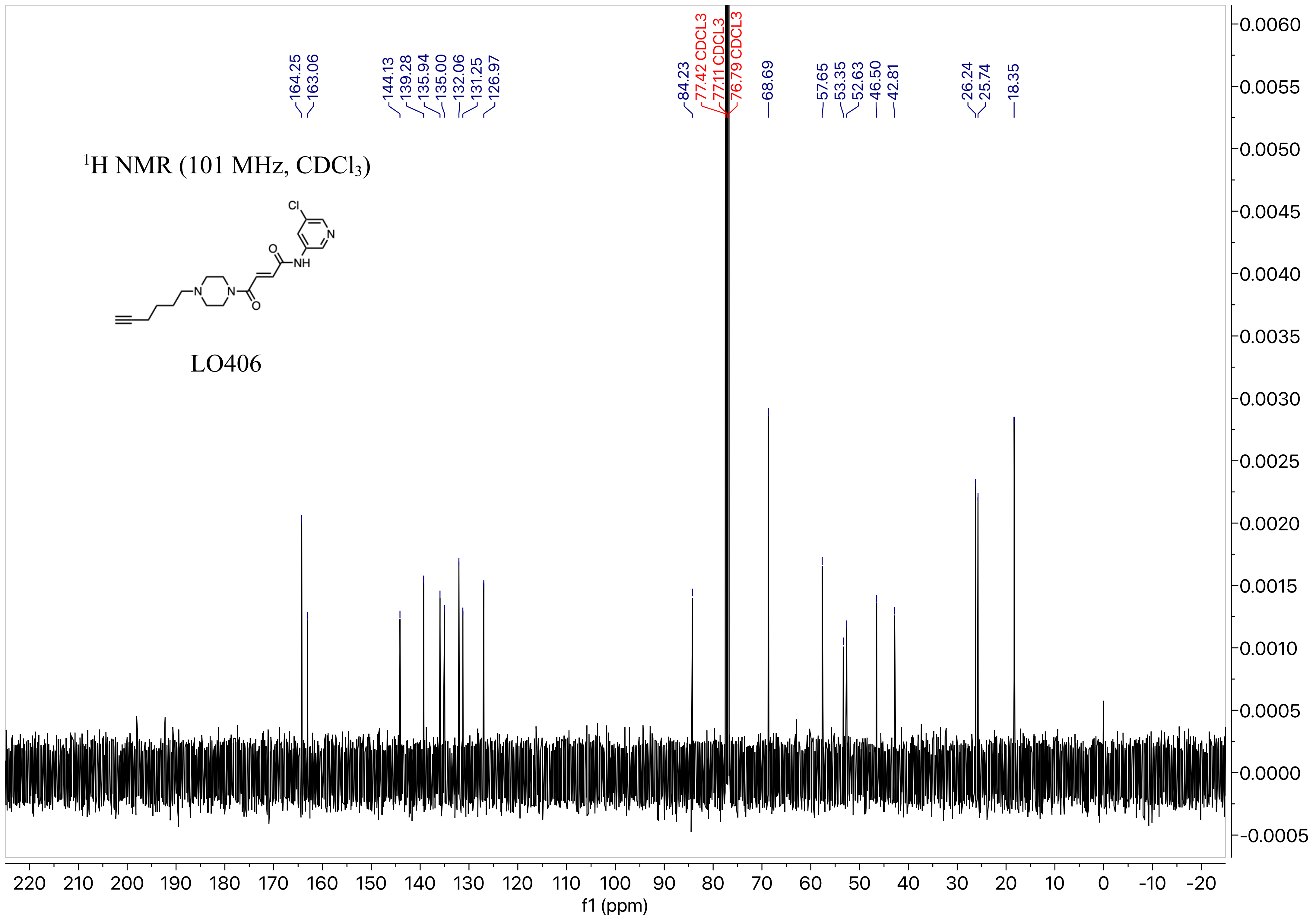

1H NMR (101 MHz, CDCl3)
LO406
Cl
N
O
NH
N
N
O
164.25
163.06
144.13
139.28
135.94
135.00
132.06
131.25
126.97
84.23
77.42 CDCL3
77.11 CDCL3
76.79 CDCL3
68.69
57.65
53.35
52.63
46.50
42.81
26.24
25.74
18.35
f1 (ppm)

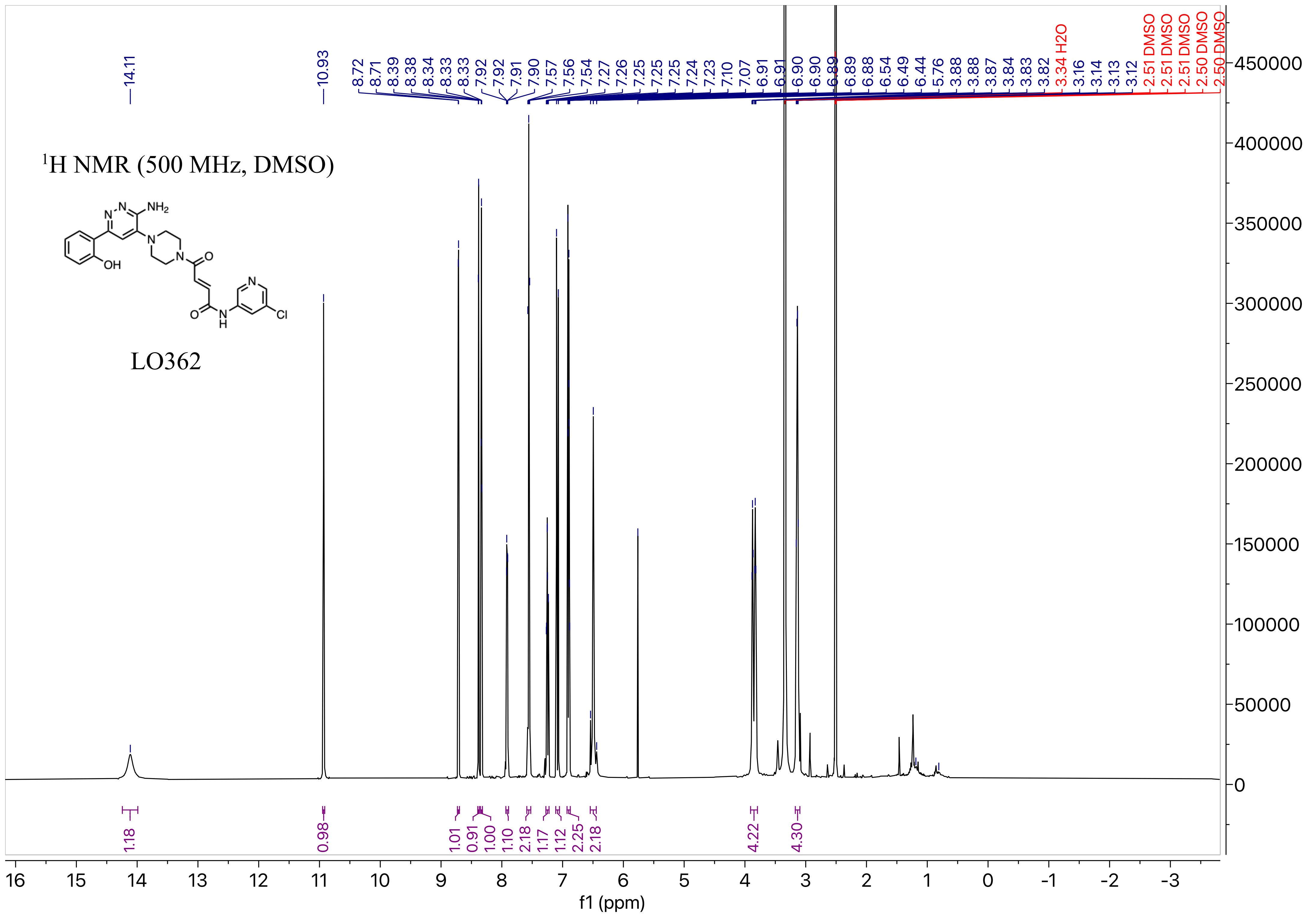

1H NMR (500 MHz, DMSO)
NH2
N
N
OH
N
N
O
O
N
H
N
Cl
LO362
14.11
10.93
8.72 8.71 8.39 8.38 8.34 8.33 8.33 7.92 7.92 7.91 7.90 7.57 7.56 7.54 7.27 7.26 7.25 7.25 7.25 7.24 7.23 7.10 7.07 6.91 6.91 6.90 6.90 6.89 6.89 6.88 6.54 6.49 6.44 5.76 3.88 3.88 3.87 3.84 3.83 3.82
3.34 H2O
3.16 3.14 3.13 3.12
2.51 DMSO 2.51 DMSO 2.51 DMSO 2.50 DMSO 2.50 DMSO
1.18 0.98 1.01 0.91 1.00 1.10 2.18 1.17 1.12 2.25 2.18 4.22 4.30
f1 (ppm)

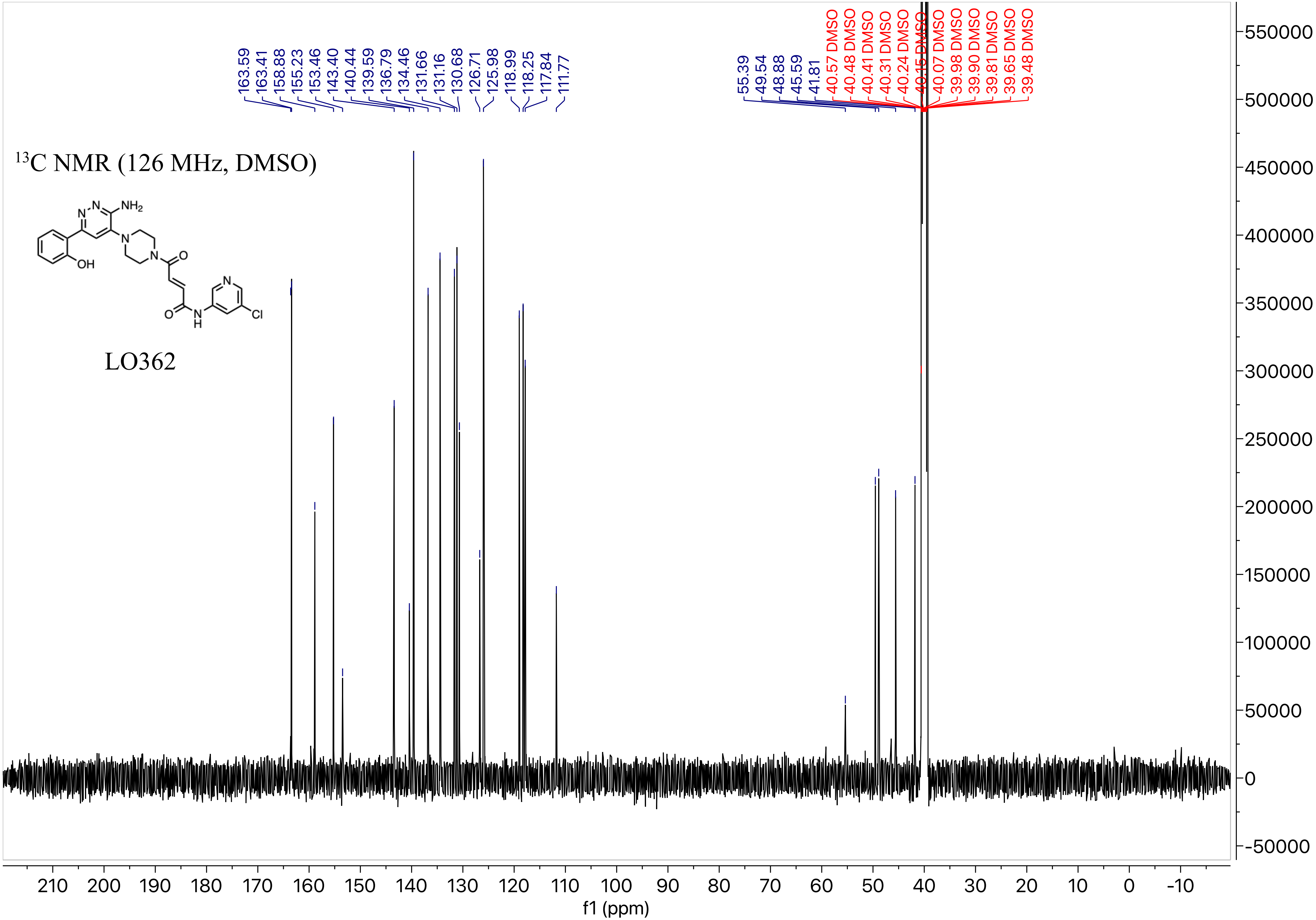

13C NMR (126 MHz, DMSO)
LO362
163.59
163.41
158.88
155.23
153.46
143.40
140.44
139.59
136.79
134.46
131.66
131.16
130.68
126.71
125.98
118.99
118.25
117.84
111.77
55.39
49.54
48.88
45.59
41.81
40.57 DMSO
40.48 DMSO
40.41 DMSO
40.31 DMSO
40.24 DMSO
40.15 DMSO
40.07 DMSO
39.98 DMSO
39.90 DMSO
39.81 DMSO
39.65 DMSO
39.48 DMSO
NH2
N
OH
O
H
Cl
f1 (ppm)

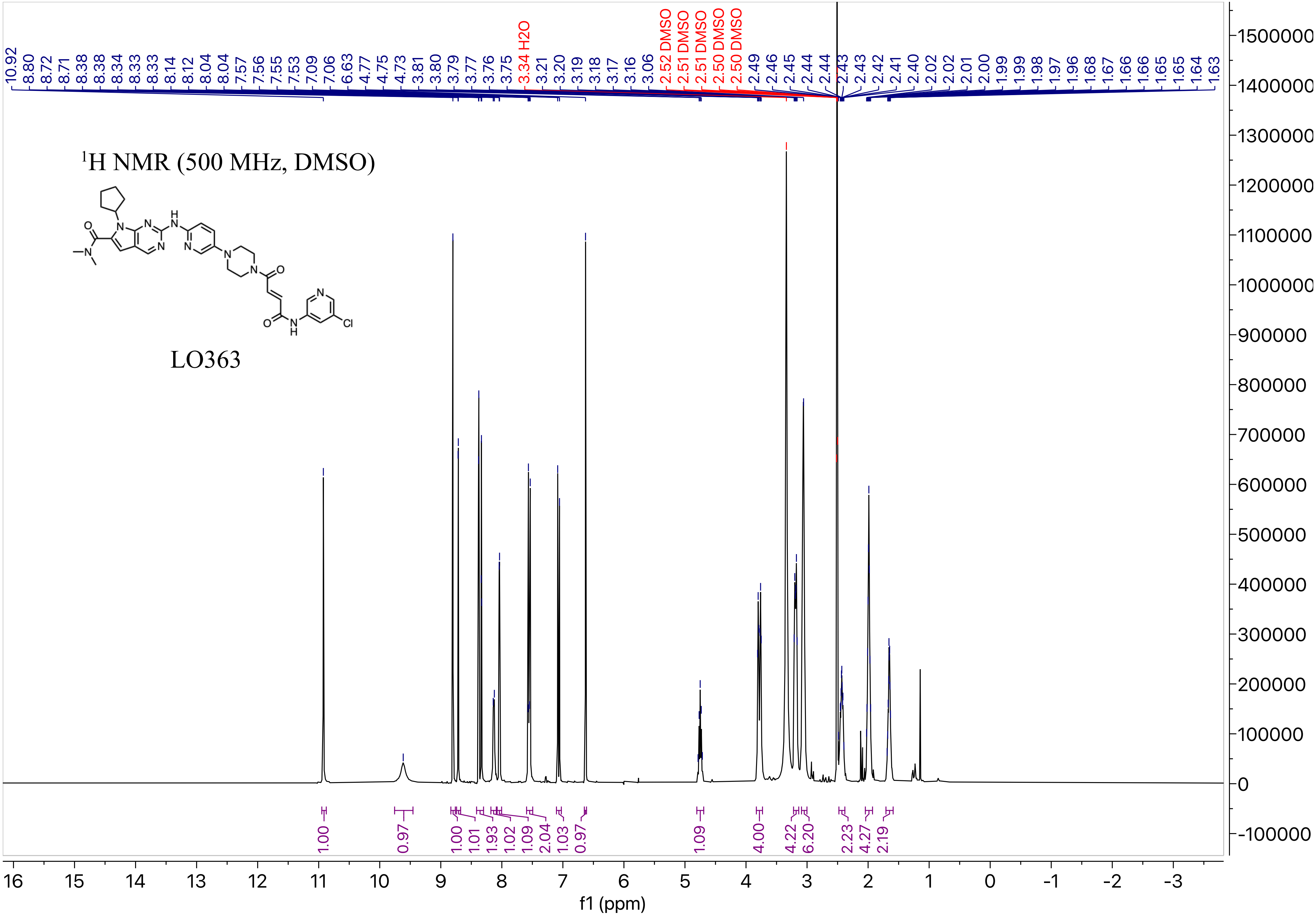
[1]H NMR (500 MHz, DMSO)
LO363
f1 (ppm)

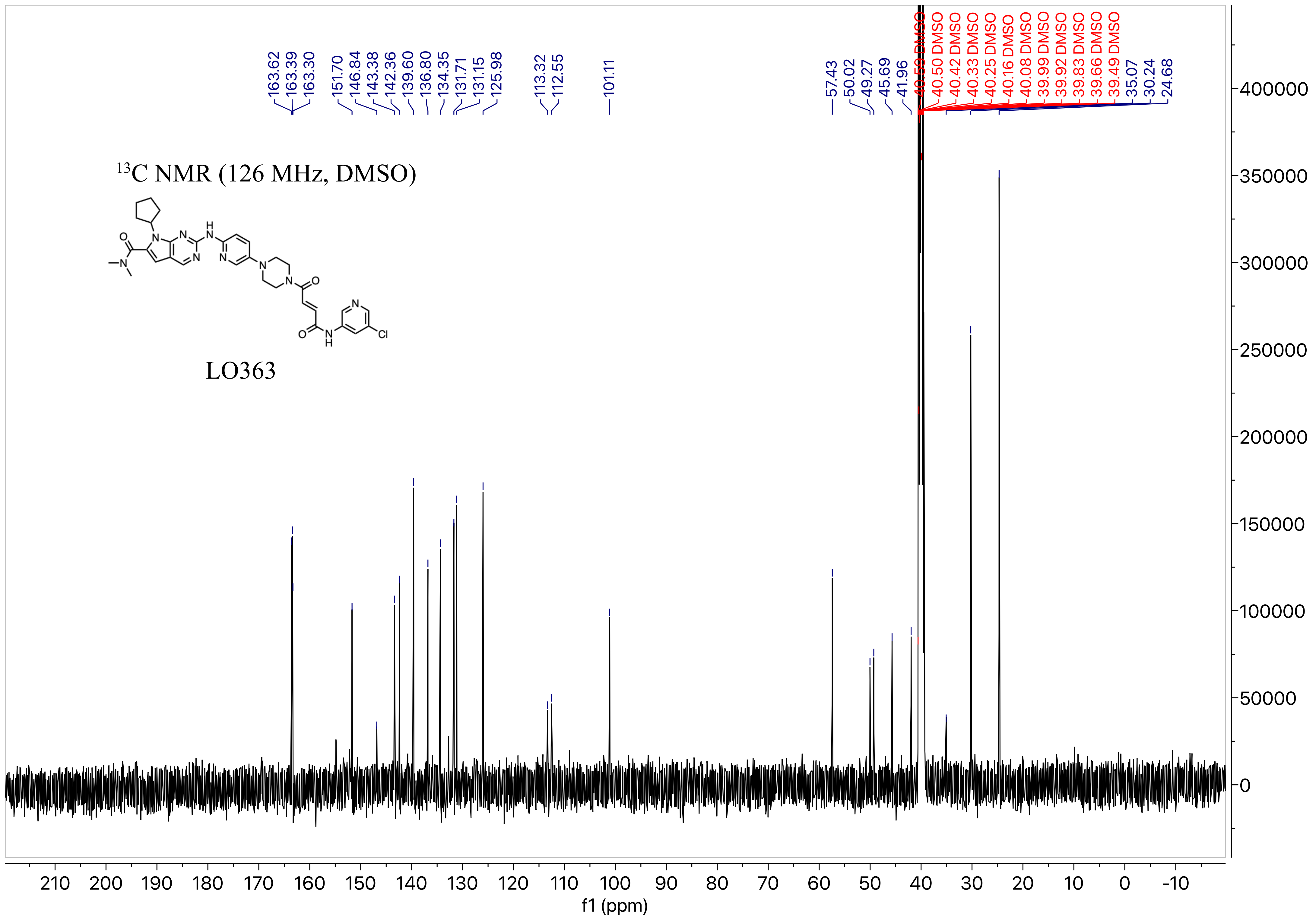

163.62
163.39
163.30
151.70
146.84
143.38
142.36
139.60
136.80
134.35
131.71
131.15
125.98
113.32
112.55
101.11
57.43
50.02
49.27
45.69
41.96
40.50 DMSO
40.42 DMSO
40.33 DMSO
40.25 DMSO
40.16 DMSO
40.08 DMSO
39.99 DMSO
39.92 DMSO
39.83 DMSO
39.66 DMSO
39.49 DMSO
35.07
30.24
24.68
13C NMR (126 MHz, DMSO)
LO363
f1 (ppm)

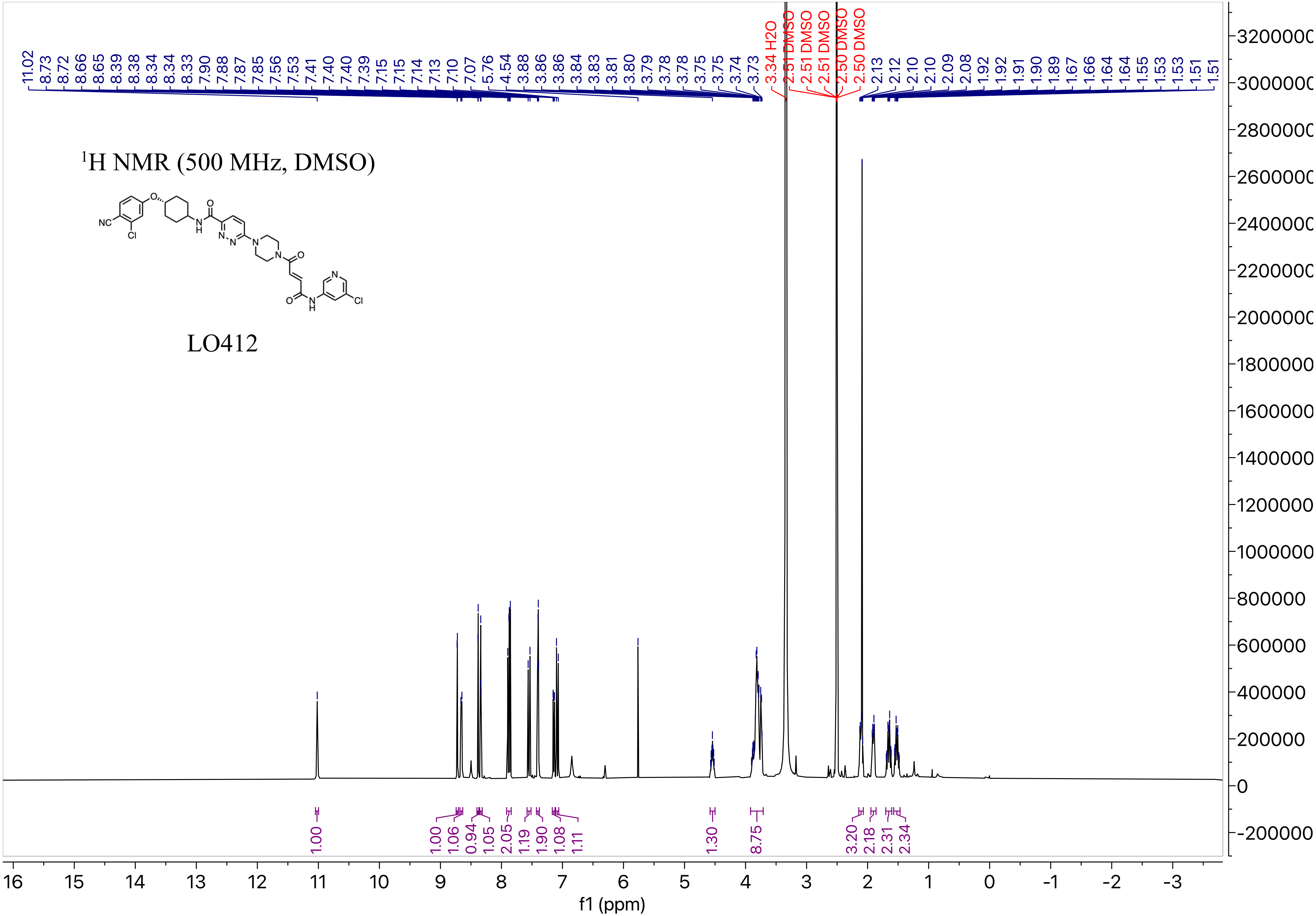

1H NMR (500 MHz, DMSO)
LO412
11.02 8.73 8.72 8.66 8.65 8.39 8.38 8.34 8.34 8.33 7.90 7.88 7.87 7.85 7.56 7.53 7.41 7.40 7.40 7.39 7.15 7.15 7.14 7.13 7.10 7.07 5.76 4.54 3.88 3.86 3.86 3.84 3.83 3.81 3.80 3.79 3.78 3.78 3.75 3.75 3.74 3.73
3.34 H2O
2.51 DMSO
2.51 DMSO
2.51 DMSO
2.50 DMSO
2.50 DMSO
2.13 2.12 2.10 2.10 2.09 2.08 1.92 1.92 1.91 1.90 1.89 1.67 1.66 1.64 1.64 1.55 1.53 1.53 1.51 1.51
1.00 1.00 1.06 0.94 1.05 2.05 1.19 1.90 1.08 1.11 1.30 8.75 3.20 2.18 2.31 2.34
f1 (ppm)

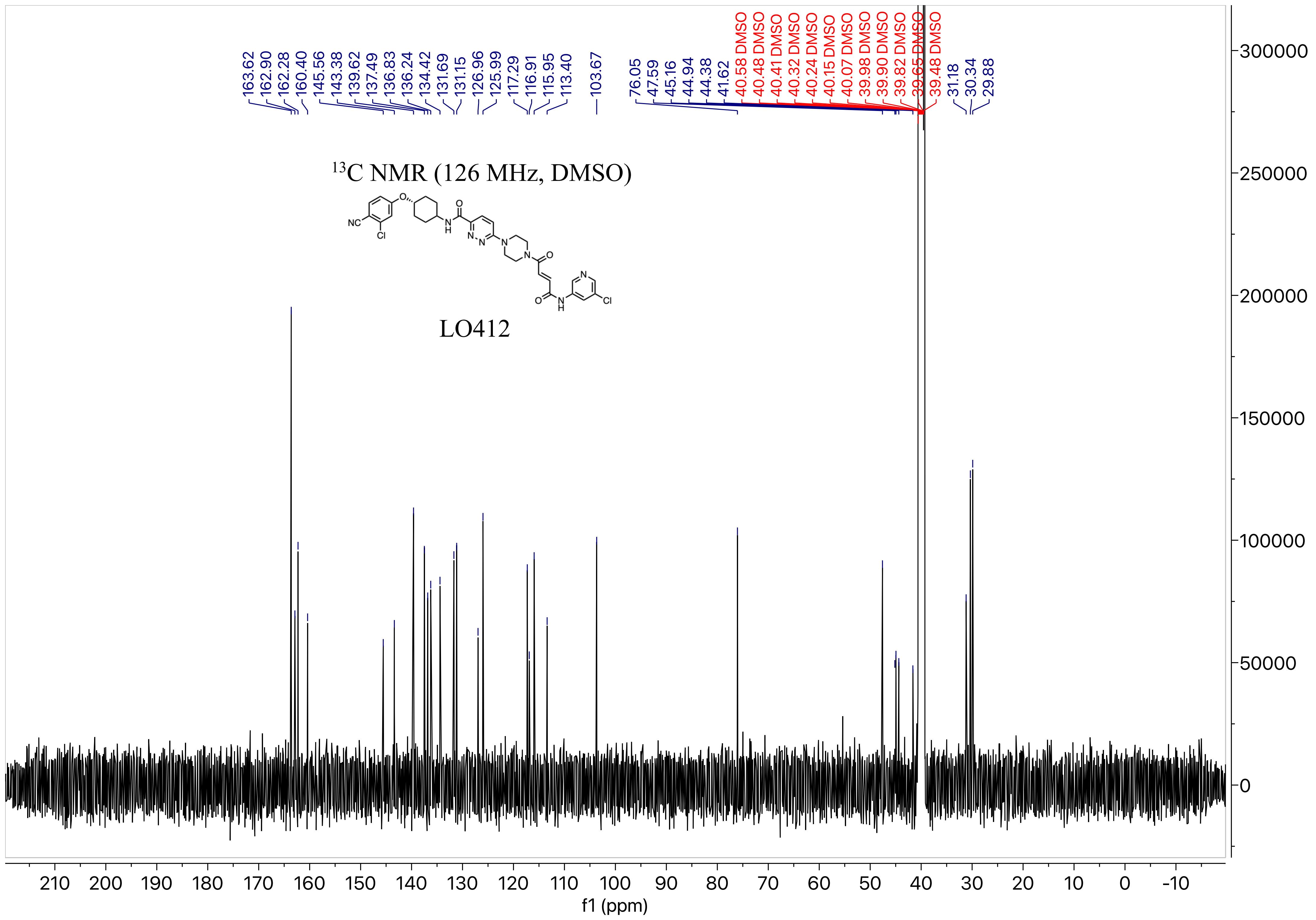

13C NMR (126 MHz, DMSO)
LO412
NC
Cl
O
N
H
N
N
N
N
O
O
N
H
N
Cl
163.62
162.90
162.28
160.40
145.56
143.38
139.62
137.49
136.83
136.24
134.42
131.69
131.15
126.96
125.99
117.29
116.91
115.95
113.40
103.67
76.05
47.59
45.16
44.94
44.38
41.62
40.58 DMSO
40.48 DMSO
40.41 DMSO
40.32 DMSO
40.24 DMSO
40.15 DMSO
40.07 DMSO
39.98 DMSO
39.90 DMSO
39.82 DMSO
39.65 DMSO
39.48 DMSO
31.18
30.34
29.88
f1 (ppm)

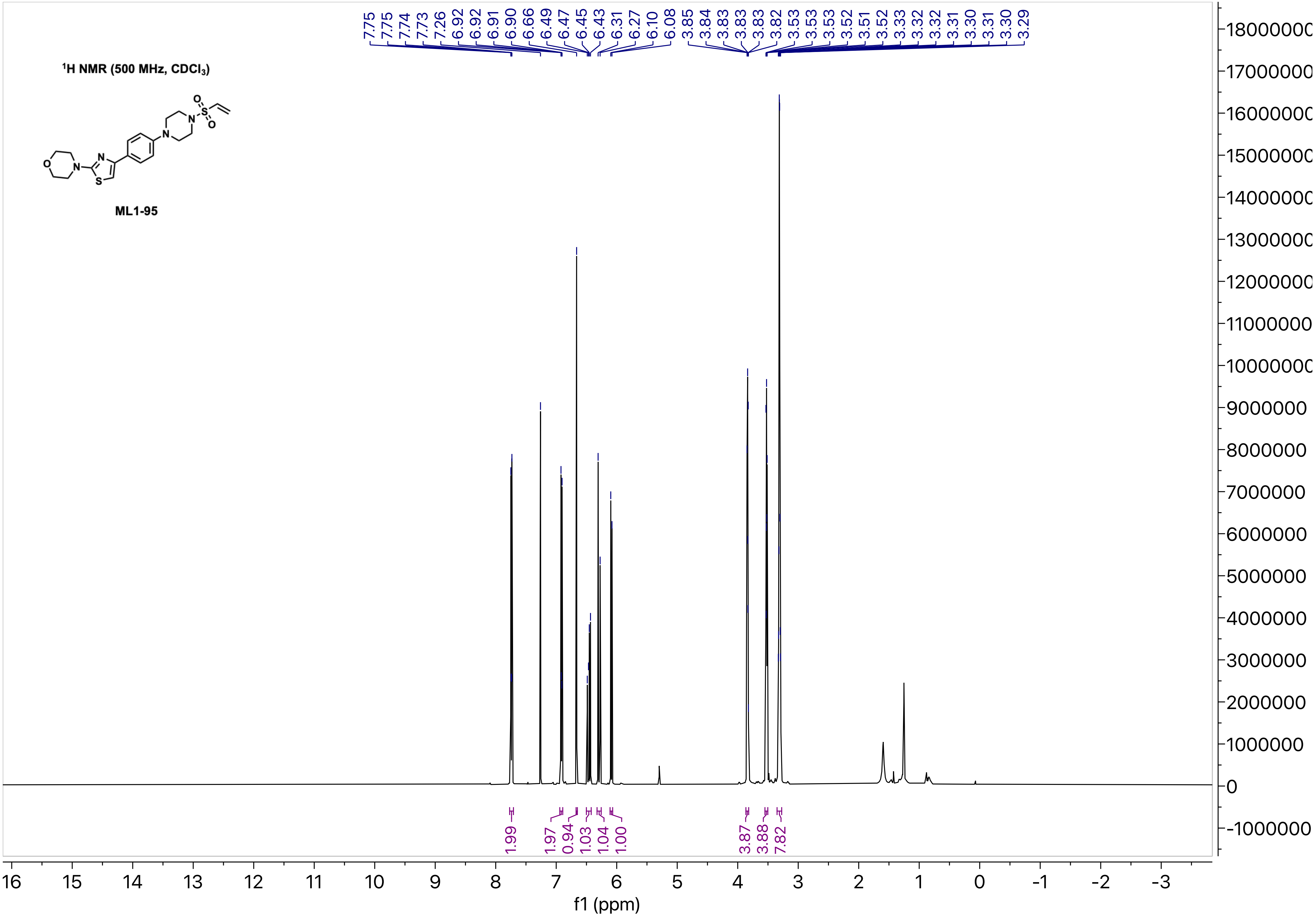

$^1$H NMR (500 MHz, $CDCl_3$)
ML1-95
7.75 7.75 7.74 7.73 7.26 6.92 6.92 6.91 6.90 6.66 6.49 6.47 6.45 6.43 6.31 6.27 6.10 6.08 3.85 3.84 3.83 3.83 3.83 3.82 3.53 3.53 3.53 3.52 3.51 3.52 3.33 3.32 3.32 3.31 3.30 3.31 3.30 3.29
1.99 1.97 0.94 1.03 1.04 1.00 3.87 3.88 7.82
f1 (ppm)

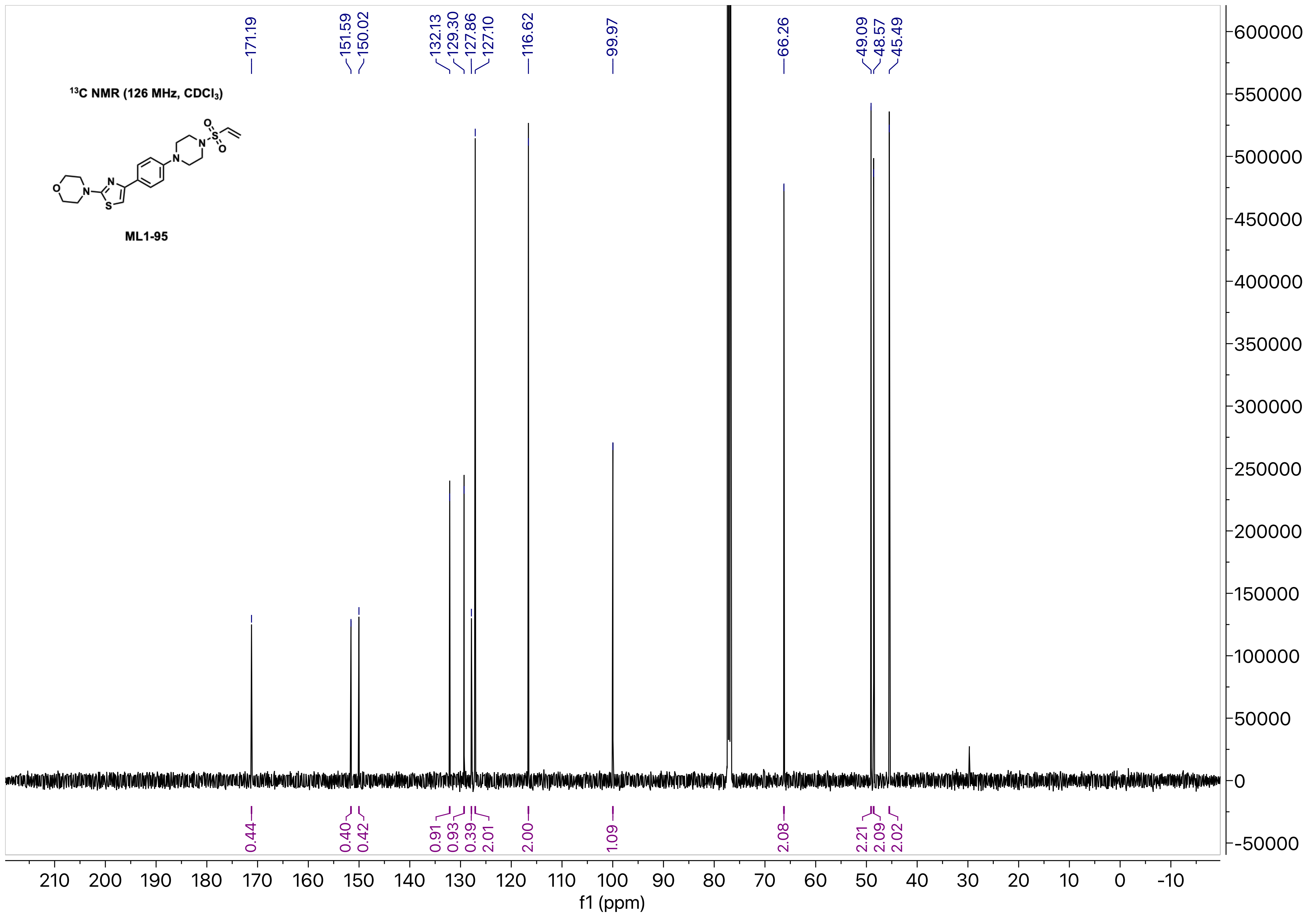

13C NMR (126 MHz, CDCl3)
ML1-95
171.19
151.59
150.02
132.13
129.30
127.86
127.10
116.62
99.97
66.26
49.09
48.57
45.49
0.44
0.40
0.42
0.91
0.93
0.39
2.01
2.00
1.09
2.08
2.21
2.09
2.02
f1 (ppm)

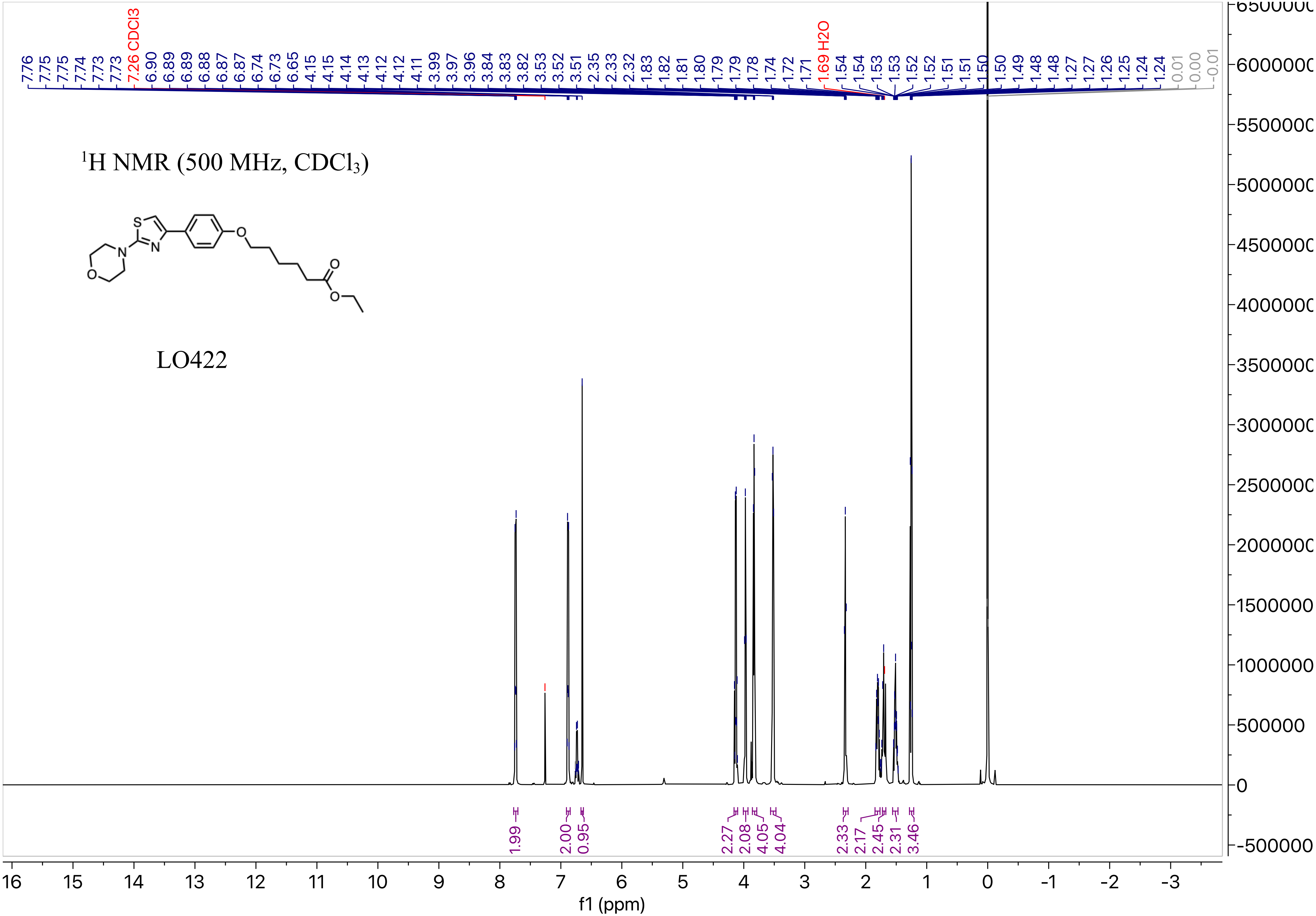

$^1H$ NMR (500 MHz, $CDCl_3$)
LO422
7.76 7.75 7.75 7.74 7.73 7.73 7.26 CDCl3 6.90 6.89 6.89 6.88 6.87 6.87 6.74 6.73 6.65 4.15 4.15 4.14 4.13 4.12 4.12 4.11 3.99 3.97 3.96 3.84 3.83 3.82 3.53 3.52 3.51 2.35 2.33 2.32 1.83 1.82 1.81 1.80 1.79 1.79 1.78 1.74 1.72 1.71 1.69 H2O 1.54 1.54 1.53 1.53 1.52 1.52 1.51 1.51 1.50 1.50 1.49 1.48 1.48 1.27 1.27 1.26 1.25 1.24 1.24 0.01 0.00 -0.01
1.99 2.00 0.95 2.27 2.08 4.05 4.04 2.33 2.17 2.45 2.31 3.46
f1 (ppm)

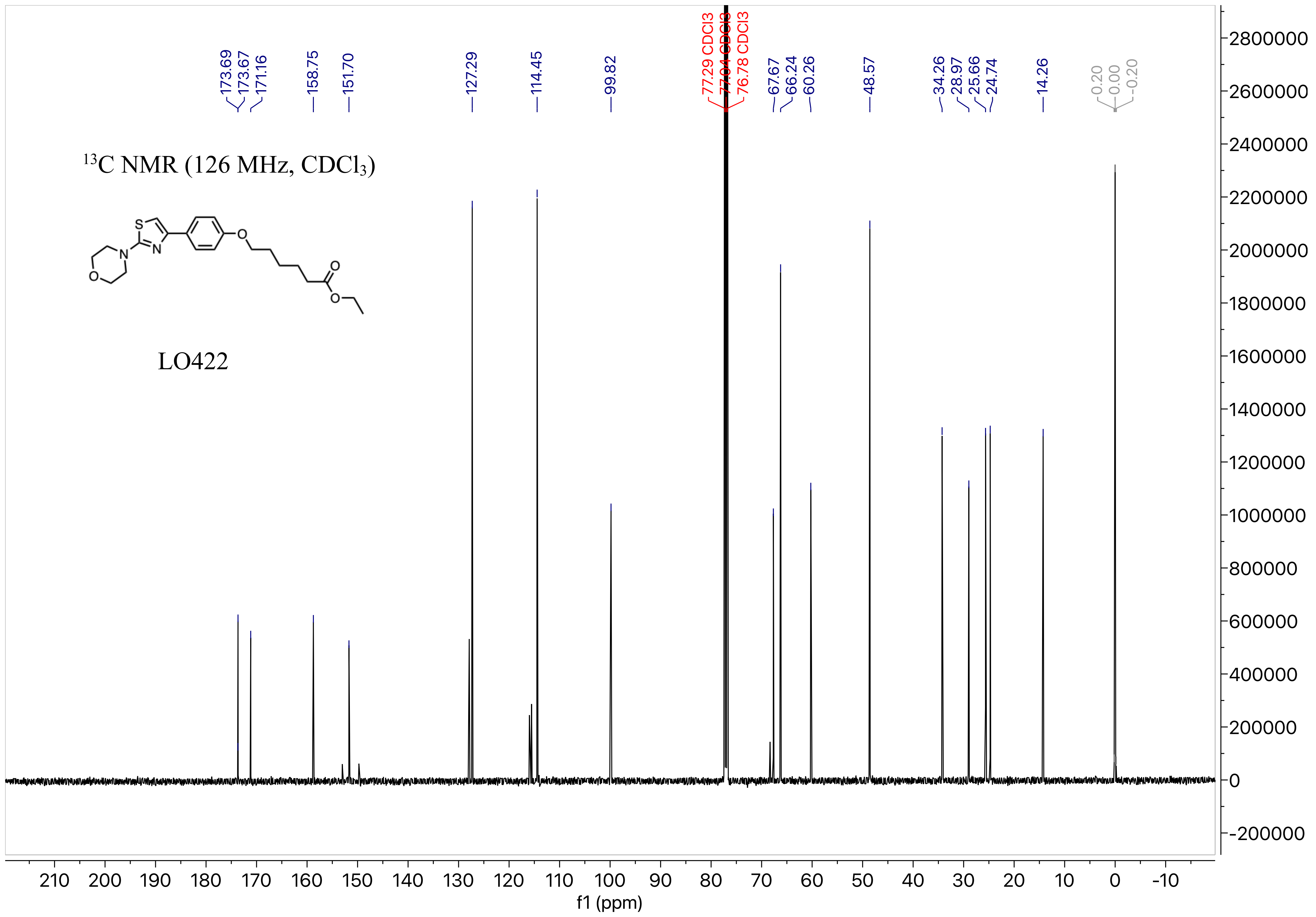
173.69
173.67
171.16
158.75
151.70
127.29
114.45
99.82
77.29 CDCl3
77.04 CDCl3
76.78 CDCl3
67.67
66.24
60.26
48.57
34.26
28.97
25.66
24.74
14.26
0.20
0.00
-0.20
13C NMR (126 MHz, CDCl3)
S
N
N
O
O
O
O
LO422
f1 (ppm)

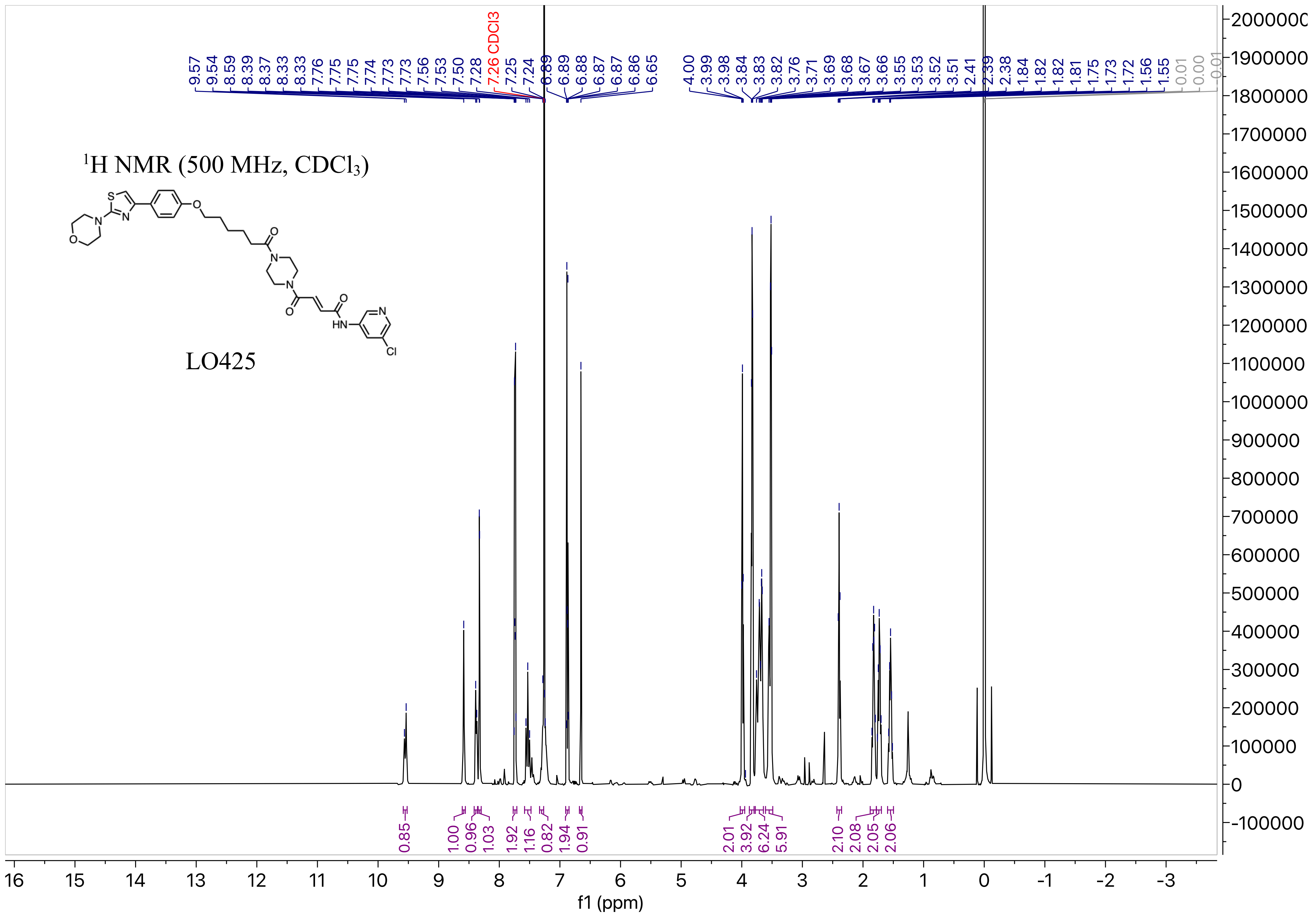
1H NMR (500 MHz, CDCl3)
LO425
9.57 9.54 8.59 8.39 8.37 8.33 8.33 7.76 7.75 7.75 7.74 7.73 7.73 7.56 7.53 7.50 7.28 7.26 CDCl3 7.25 7.24 6.89 6.89 6.88 6.87 6.87 6.86 6.65
4.00 3.99 3.98 3.84 3.83 3.82 3.76 3.71 3.69 3.68 3.67 3.66 3.55 3.53 3.52 3.51 2.41 2.39 2.38 1.84 1.82 1.82 1.81 1.75 1.73 1.72 1.56 1.55 0.01 0.00 -0.01
0.85 1.00 0.96 1.03 1.92 1.16 0.82 1.94 0.91 2.01 3.92 6.24 5.91 2.10 2.08 2.05 2.06
f1 (ppm)

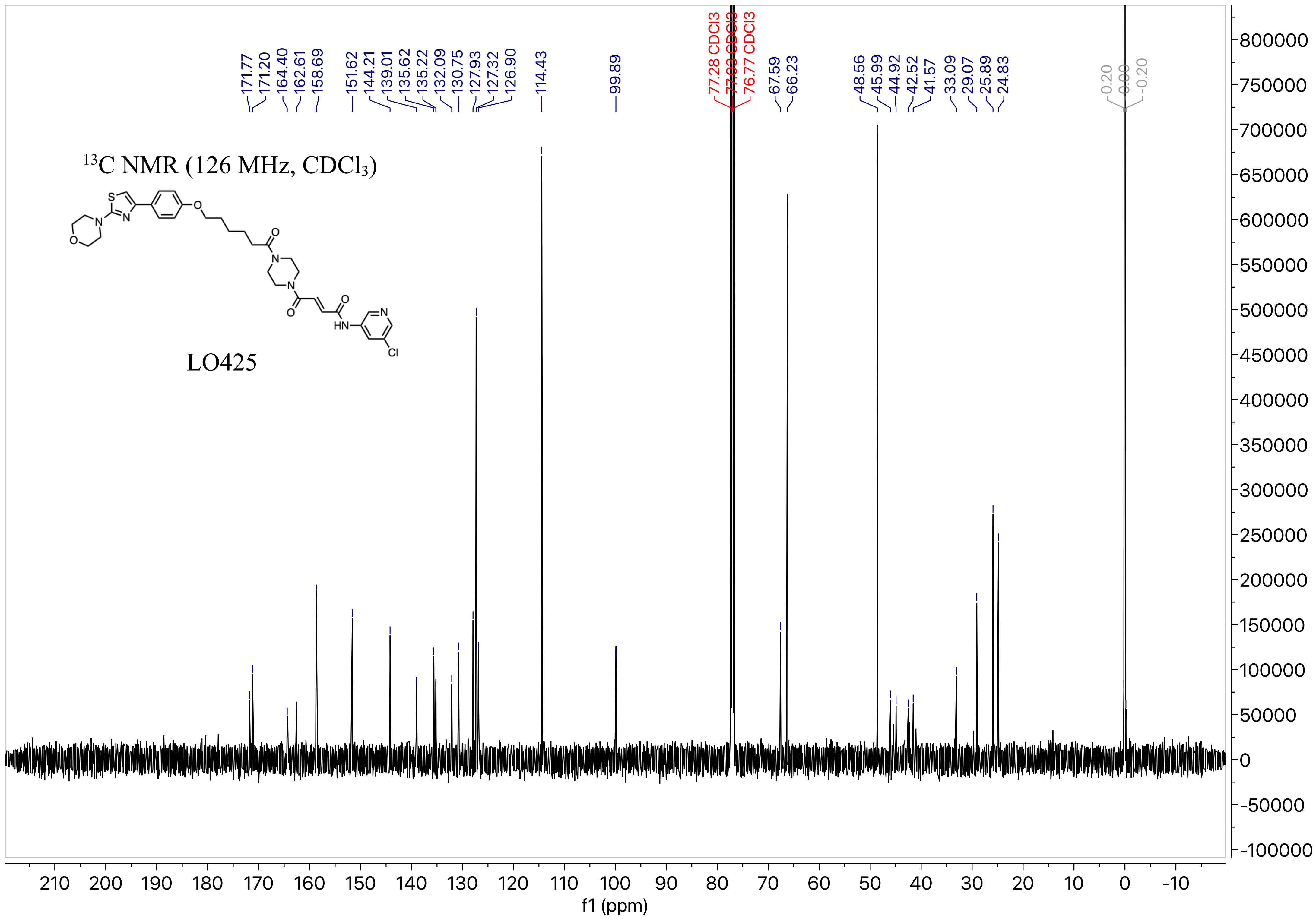
13C NMR (126 MHz, CDCl3)
LO425
171.77
171.20
164.40
162.61
158.69
151.62
144.21
139.01
135.62
135.22
132.09
130.75
127.93
127.32
126.90
114.43
99.89
77.28 CDCl3
77.03 CDCl3
76.77 CDCl3
67.59
66.23
48.56
45.99
44.92
42.52
41.57
33.09
29.07
25.89
24.83
0.20
0.00
-0.20
f1 (ppm)
